# A population-scale Y chromosome reference resource resolves structural diversity and reduces reference bias

**DOI:** 10.64898/2026.06.03.729890

**Authors:** Pille Hallast, Arang Rhie, Mark Loftus, Mariateresa Mazzetto, Peter Ebert, Shenghan Gao, Gianni V. Martino, Peter A. Audano, Hufsah Ashraf, Karol Pal, Rebecca Siford, Jana Ebler, Sergey Koren, DongAhn Yoo, Kwondo Kim, Yunzhe Jiang, Nancy F. Hansen, Prajna Hebbar, Tomoya Kanno, Oliver Purnoch, Carolina Montano, Jiadong Lin, Keisuke K. Oshima, Luyao Ren, Andrea Guarracino, Matthew Jensen, Brendan J. Pinto, David Porubsky, Samuel Kuziel, Lingbin Ni, Jiaqi Li, Parithi Balachandran, Feyza Yilmaz, HGSVC, HPRC, Jan O. Korbel, Benedict Paten, Mark Gerstein, Kateryna D. Makova, Xinghua Shi, Evan E. Eichler, Christine R. Beck, Melissa A. Wilson, Miriam K. Konkel, Tobias Marschall, Monika Cechova, Glennis A. Logsdon, Adam M. Phillippy, Charles Lee

## Abstract

The human Y chromosome remains underrepresented in genomic resources because its repetitive architecture is difficult to resolve and poorly represented by any single reference. Here, we present 142 curated near-complete *de novo* Y chromosome assemblies spanning 98 terminal-marker-defined lineages across 17 haplogroup clades and 180,000 years of evolution, establishing a population-scale sequence and annotation resource for the chromosome. Across 100 quality-filtered *AZFc* intervals, recurrent inversions, deletions and complex rearrangements resolve into 12 structural haplotypes, while *DAZ*, *RBMY* and *TSPY* families show distinct paralogue- and array-specific evolution. Complete centromere and Yq12 sequences reveal extensive lineage-specific change within comparatively stable higher-order architectures, and haplotype-resolved methylomes show distinct epigenetic states among similar repeat copies. The reference panel reduces analytical bias: in testis Iso-Seq data, the best-matching assembly increased median *DAZ* transcript alignment identity from 91.7% to 99.9% and improved isoform annotation. Together, this resource enables analysis of Y-chromosome structural, evolutionary and reproductive variation.

## Introduction

The human Y chromosome is one of the most structurally unusual chromosomes in the genome^1,2^. In addition to its central roles in sex determination and spermatogenesis, several retained Y-linked genes are broadly expressed across non-reproductive tissues and function as dosage-sensitive regulators of cellular processes^3,4^. Yet the Y was the last human chromosome to be completely deciphered because much of its sequence is exceptionally difficult to assemble and interpret^1,2^. It contains megabases of satellite-rich heterochromatin, extensive segmental duplications, large palindromes, and multicopy ampliconic gene families (**Fig. 1a)**. These same features also make the Y exceptionally structurally dynamic: recurrent inversions, deletions, duplications and copy-number changes within the azoospermia factor (*AZF)* c region, *TSPY*, Yq heterochromatin and other repeat-rich regions have been documented for decades^5–9^. Thus, the existence of extensive Y-chromosome structural polymorphism is well established. What has remained much harder to determine is its sequence-resolved architecture: which repeat units and gene copies are altered, how often similar structures independently arise in different Y lineages, and which features remain constrained despite rapid sequence turnover.

**Figure 1.**
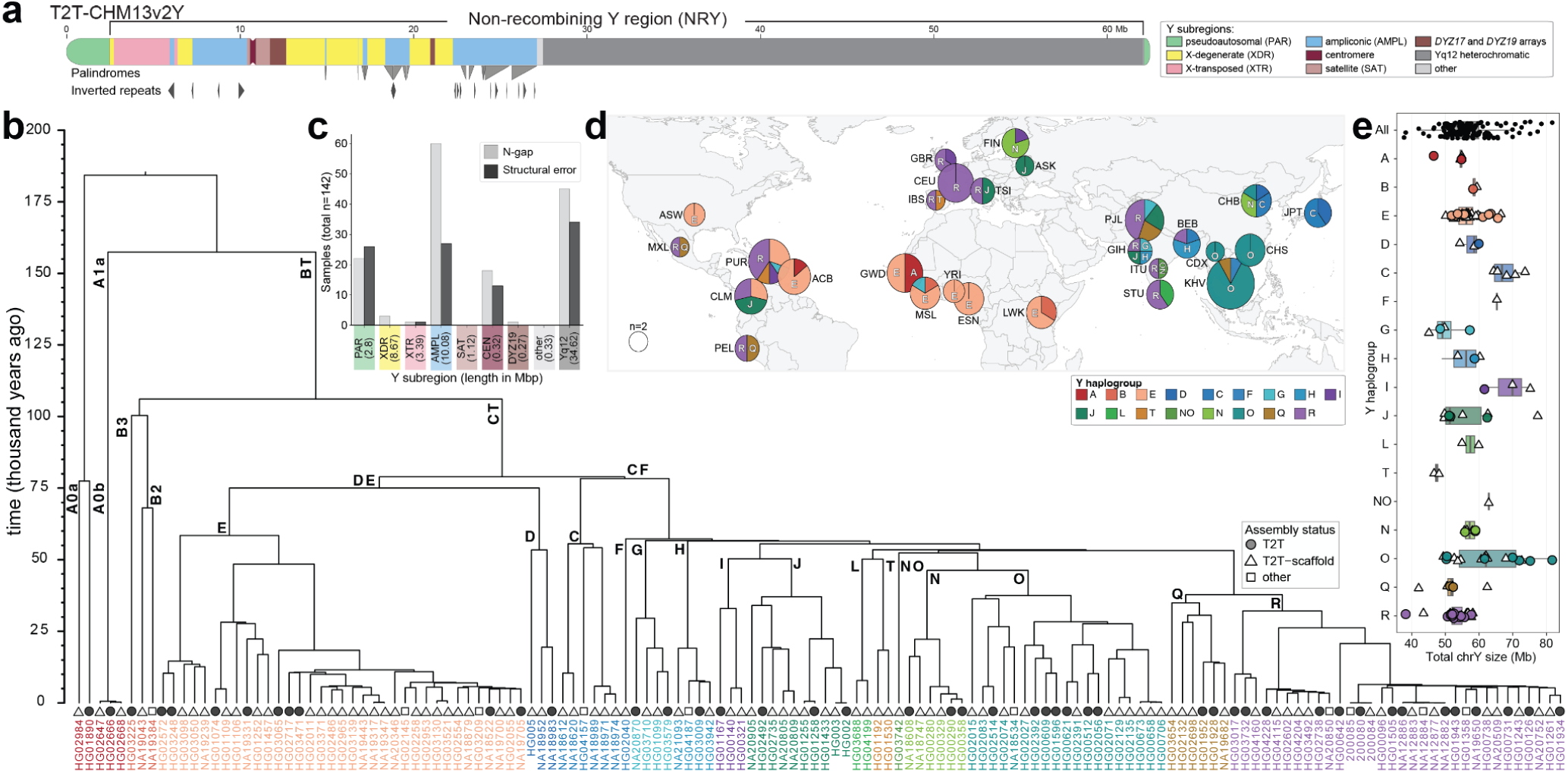
Phylogenetic framework, geographic sampling, and assembly characteristics of 142 human Y chromosomes. **a**, Schematic of the T2T-CHM13v2Y reference showing the Y chromosome organization into major sequence classes and subregions, including the pseudoautosomal region, X-degenerate, X-transposed, ampliconic, centromeric, and heterochromatic compartments, with palindromes and inverted repeats indicated below. **b**, Time-calibrated phylogeny of the 142 assembled Y chromosomes spanning the major branches of human Y chromosome diversity. Branch lengths are scaled in thousands of years before present, and major haplogroup clades are indicated on the tree. Symbols at the tips denote assembly status (telomere to telomere - T2T as a circle, T2T-scaffold as a triangle, or other as a square) and sample names are coloured according to Y haplogroup. **c**, Number of assemblies with flagged issues across major T2T-CHM13v2Y subregion groups. Bar plot summarizing the number of Y-chromosome assemblies (out of 142 total) with at least one flagged issue in each major sequence class group defined using the T2T-CHM13v2Y annotation. T2T-CHM13v2Y subregion group sizes are shown below in Mb. Note: the *DYZ17* array is considered part of the satellite class (SAT). **d**, A world map indicating for each of the populations the geographic source, sample size, and haplogroup distribution. **e**, Total assembled Y chromosome size across all samples and across major haplogroups, with each point representing one assembly and symbols indicating assembly status. Boxplots of chromosome length (showing the median and interquartile range) summarize the distribution within haplogroups, highlighting extensive variation in Y-chromosome length across the human phylogeny.

Recent efforts by the Human Pangenome Reference Consortium (HPRC), Human Genome Structural Variation Consortium (HGSVC) and the Telomere-to-Telomere (T2T) consortium produced the first full and near-complete Y-chromosome assemblies^2,10^, resolving multi-megabase regions that had previously been absent or fragmented in the human reference. Our previous survey of 43 diverse Y chromosomes provided the first broad sequence-level view of variation across ampliconic, centromeric and heterochromatic regions^2,10^. However, only three of those 43 chromosomes were complete from telomere to telomere, leaving the most repetitive compartments unresolved across much of the Y phylogeny. Consequently, it remained difficult to distinguish recurrent rearrangements from lineage-specific events, resolve variation at the level of individual paralogues and repeat arrays, or determine whether apparently conserved structures reflected biological constraint or limited sampling. These studies also made clear that no single linear Y chromosome can adequately represent the full range of human Y-chromosome structural diversity.

These limitations matter because many of the Y chromosome’s most structurally complex regions are also among its most biologically and clinically relevant. The *AZF*c interval contains large, highly identical ampliconic repeats whose recurrent rearrangements are associated with spermatogenic impairment. Yet broad deletion classes do not capture the precise gene copies and structural configurations that are retained or lost. For multicopy gene families such as *DAZ*, *RBMY,* and *TSPY,* aggregate copy number alone cannot reveal which paralogues have expanded, contracted, exchanged sequence, or changed structural context. The repetitive architecture that generates this structural variation also complicates epigenetic and transcriptomic analysis because reads from near-identical copies can be collapsed or misassigned when projected onto a single reference chromosome. More broadly, human pangenome studies have demonstrated that representing multiple haplotypes can reduce reference bias and improve variant discovery^11,12^. The Y therefore presents both an acute need and an unusually difficult test case for moving beyond a single linear reference. Resolving these regions across many independent paternal lineages makes it possible to move beyond cataloguing polymorphism toward identifying recurrent mechanisms of structural change, the architectural features that remain constrained, and the consequences of reference choice for downstream analyses.

Here, we establish a phylogenetically broad, sequence-resolved reference panel of142 curated *de novo* Y chromosome assemblies spanning 17 broad haplogroup clades and more than 180,000 years of evolution. The panel includes 46 fully T2T chromosomes and substantially expands the number of lineages for which highly repetitive Y sequences can be examined directly. This allows repetitive sequences to be compared as resolved architecture rather than inferred from incomplete assemblies or aggregate measurements. We use this expanded phylogenetic framework to distinguish recurrent rearrangements from shared ancestry within the *AZFc* region, and to resolve paralogue- and array-specific evolution of the *DAZ*, *RBMY*, and *TSPY* gene families across independent Y lineages. We then examine how this structural turnover extends into the chromosome’s repetitive compartments, including palindromes, the centromeric *DYZ3* array and Yq12, and ask which features remain constrained despite extensive sequence remodeling. By integrating personalized assemblies with Oxford Nanopore methylation and long-read testis transcript data, we further examine how structural context relates to epigenetic state and transcript interpretation. Finally, we use the assembly panel as an alternative-reference and benchmarking resource to quantify how Y-chromosome structural diversity affects transcript mapping, copy-number estimation, sequence reconstruction and short-read callability. Together, this resource moves the Y chromosome from a collection of difficult-to-resolve structural polymorphisms toward a sequence-resolved framework for studying its evolution, fertility-associated variation and representation in population-scale genomic and transcriptomic analyses.

## Results

### Population-scale, near T2T Y-chromosome assemblies establish a reference resource

To establish a population-scale, sequence-resolved reference resource for the human Y chromosome, we selected samples based on availability of sequencing data at the time to maximize chances to cover broader Y haplogroups. Initially, 144 Y chromosomes from the HPRC (n=105)^12^, HGSVC (n=29)^13^, Genome in a Bottle (GIAB; n=3, 2 overlapping with HPRC), and a four-generation pedigree (CEPH1463; n=9)^14,15^ were re-assembled with Verkko v2.2.1 using Pacific Biosciences HiFi (PacBio HiFi, median coverage of 64.2x), Oxford Nanopore ultra-long (ONT-UL 155.9x total and 33.7x reads >100 kb in length), Dovetail Genomics Hi-C (Hi-C, 57x) and trio Illumina reads^16^ when samples were available (**Suppl. Table 1**). Assemblies were available for 136 of these samples (hereafter “prior-best”, **Suppl. Table 2**), of which 127 (93.4%) remained incomplete within Yq12, ampliconic, centromeric or other challenging regions. To improve resolution of these compartments, we generated additional PacBio HiFi data for 20 samples (**Suppl. Table 3-4**). After initial assessment, 61 underwent extensive manual curation of the Y-chromosome assembly graphs using graph topology and long-read support (**Suppl. Tables 4–5**). Following curation and quality control^17^, 142 Y chromosomes were retained for subsequent analyses, including 46 (32.4%) T2T, 86 (60.6%) T2T scaffold assemblies with one or more gaps, and 10 (7%) other assemblies missing one or both telomeres (**Fig. 1b**). The remaining gaps were confined predominantly to ampliconic (n=60), Yq12 (n=45), pseudoautosomal region 1 (PAR1, n=21) or the centromeric (n=18) regions (**Fig. 1c; Extended Data Fig. 1a-b; Suppl. Results (Assembly and QC); Suppl. Fig. 1; Suppl. Tables 4-6**). Structural accuracy of the curated Y assemblies were measured using PacBio HiFi and ONT-UL alignments^18^, assessed with HMM-Flagger^18^ and NucFlag^19–21^. Regions flagged as potential structural errors by both sequencing technologies and both QC tools were concentrated in PAR1 and the centromere (% affected bases in **Extended Data Fig. 1a; Suppl. Fig. 2-4**). Across the 136 prior-best matched samples, the median number of Y-chromosome sequences decreased from six to two in the present resource, demonstrating substantially increased contiguity (**Suppl. Fig. 5; Suppl. Table 2**). The resulting median error rate was 14.9 bp per Mb excluding unresolved N gaps (**Suppl. Figs. 2-4**). Base-level accuracy was also comparable or higher compared to the prior-best version, with a mean Illumina 31-mers-based^22^ Quality Value (QV) of 52.4 (S.D.±3.22, Phred scale; **Suppl. Fig. 5; Suppl. Table 2**). Residual base-level errors were most frequently observed again in PAR1, consistent with previously reported HiFi coverage dropouts in this region^2,10^ (**Fig. 1c**). In subsequent in-depth analysis, regions flagged with potential structural- or base-errors were removed accordingly to ensure the accuracy and integrity of the results (**Suppl. Table 7**).

The resulting panel in this newly established resource also substantially expands sequence-resolved representation across the human Y phylogeny. The 142 assemblies span 17 broad haplogroup clades used for population-level summaries and encompass 98 distinct terminal-marker assignments according to ISOGG v15.73, capturing substantially greater diversity at finer phylogenetic resolution than conveyed by the broad haplogroup grouping alone (**Fig. 1b,d; Suppl. Fig. 1; Suppl. Table 1**). Thirteen of the 17 broad clades are represented by at least one T2T assembly, with substantial representation of African lineages (43/142 samples). Together, this provides a phylogenetically broad, high-contiguity reference panel for population-scale analysis of Y-chromosomal regions that have remained inaccessible or poorly represented in previous assemblies. The panel was designed to represent structural and phylogenetic diversity rather than to estimate population frequencies.

Across the 142 assemblies, total Y chromosome size ranged from 37.7 to 81.7 Mb (mean 56.7 Mb), with most of this variation attributable to repeat-rich compartments rather than the single-copy regions (**Fig. 1e; Extended Data Fig. 1c-e, 2a-b; Suppl. Fig. 6; Suppl. Tables 2, 8)**. Despite this nearly twofold variation in chromosome length, every Y retained exactly one copy of each known male-specific single-copy protein-coding gene, defining a highly conserved genic core, whereas gene-content diversity was concentrated in ampliconic families and other duplicated loci (**Extended Data Fig. 1c-d; Suppl. Results (Gene content); Suppl. Figs. 7-11; Suppl. Tables 8-10**). Excluding two samples under controlled access, we additionally generated nonredundant variant callsets against both T2T-CHM13v2 and GRCh38. Compared with our similarly filtered 43-Y dataset^2,10^, the expanded GRCh38 callset contained approximately 2.6-fold as many structural variants (SVs), 1.6-fold as many indels and 2.4-fold as many SNVs (**Suppl. Results (Variant calling); Suppl. Tables 11-13**). Structural variation was strongly compartmentalised: within the non-recombining region of the Y (NRY) sequence represented in GRCh38, insertion and deletion rates were the highest in ampliconic regions (16.1 and 8.8 per Mb, respectively), compared with X-degenerate (2.4 and 1.7 per Mb) and X-transposed regions (3.2 and 1.2 per Mb) (**Suppl. Results (Variant calling); Suppl. Tables 11-13**). PAR1 also showed elevated divergence and enrichment of recombination-associated sequence features relative to PAR2 and most other Y sequence classes, consistent with its unusually high recombination rate^23^ (**Suppl. Results (PAR region); Suppl. Figs. 12-16**). This concentration of variation occurs within an exceptionally duplication-rich chromosome: assemblies contained a mean of 13.9 Mb (range 9.9 - 18.2 Mb) of interchromosomal and 12.6 Mb (range 6.2 - 21.1 Mb) of intrachromosomal segmental duplications, together covering an average of 59.2% non-redundant intervals (after excluding repetitive sequence) - substantially greater than corresponding estimates for autosomes (9.5%) or the X chromosome (6.3%) from previous assemblies^13^ (**Methods; Extended Data Fig. 1e; Supp. Results (Segmental duplications); Suppl. Figs. 17-20; Suppl. Table 14**). Mobile-element insertions were rare, with only 46 putatively polymorphic young non-LTR insertions identified in the 140/142 assemblies (**Methods; Suppl. Results (Mobile element insertions); Suppl. Fig. 21; Suppl. Table 15**).

To distinguish functional enrichment from the strong sequence-context biases of Y-chromosome SVs, we performed context-matched permutations separately within the NRY and PAR1 against both GRCh38 and T2T-CHM13v2, preserving sequence class, repeat class and segmental-duplication context. Within the NRY, insertions and deletions were consistently enriched in protein-coding gene bodies and introns, and depleted from pseudogenes across both references, whereas exon enrichment was observed only for deletions. PAR1 showed a distinct pattern, with both SV classes depleted from exons and pseudogenes, while associations with protein-coding gene bodies and introns differed between references (**Suppl. Results (Functional impact of SVs); Suppl. Figs. 22–23**). Among observed NRY gene overlaps, most involved multicopy ampliconic gene families, whereas single-copy X-degenerate genes contributed few events. Together with the concentration of SVs in ampliconic sequence and the conservation of single-copy NRY genes, these results show that Y-chromosome structural diversity is strongly compartmentalized: repeat-rich and multicopy regions account for much of the structural variation, whereas the single-copy NRY gene core remains comparatively stable.

### Base-resolved analysis of *AZFc* reveals a limited repertoire of recurrent structural haplotypes

*AZFc* is among the most structurally unstable but clinically important euchromatic intervals on the human Y chromosome, and recurrent rearrangements involving its large ampliconic repeats have been associated with spermatogenic impairment^7^. However, the complete architectures, independent origins and breakpoint diversity of these rearrangements have remained difficult to compare across the human Y phylogeny because *AZFc* is composed of large, highly identical repeat blocks spanning palindromes P1–P3 (**Fig. 2a**)^1,7^. The expanded assembly panel enabled us to resolve this architecture across diverse Y lineages. In our previous survey of 43 Y chromosomes^2,10^, 19 assemblies remained interrupted within this region, limiting population-scale comparison of complete *AZFc* structures. The expanded assembly panel now allowed us to resolve the full P1/P2/P3 interval at sequence level across 100 phylogenetically diverse Y chromosomes that passed region-specific QC for the P1/P2/P3 interval (**Methods; Suppl. Fig. 24; Suppl. Table 7**). After excluding inversions confined to palindrome spacers and balanced rearrangements that preserve repeat-block order and orientation, these chromosomes resolved into 12 major haplotypes (**Fig. 2b; Suppl. Figs. 25-26**), which among the uninterrupted assemblies ranged in size from 2.5 to 6.1 Mb (mean 4.2 Mb, median 4.4 Mb; **Fig. 2c**).

**Figure 2.**
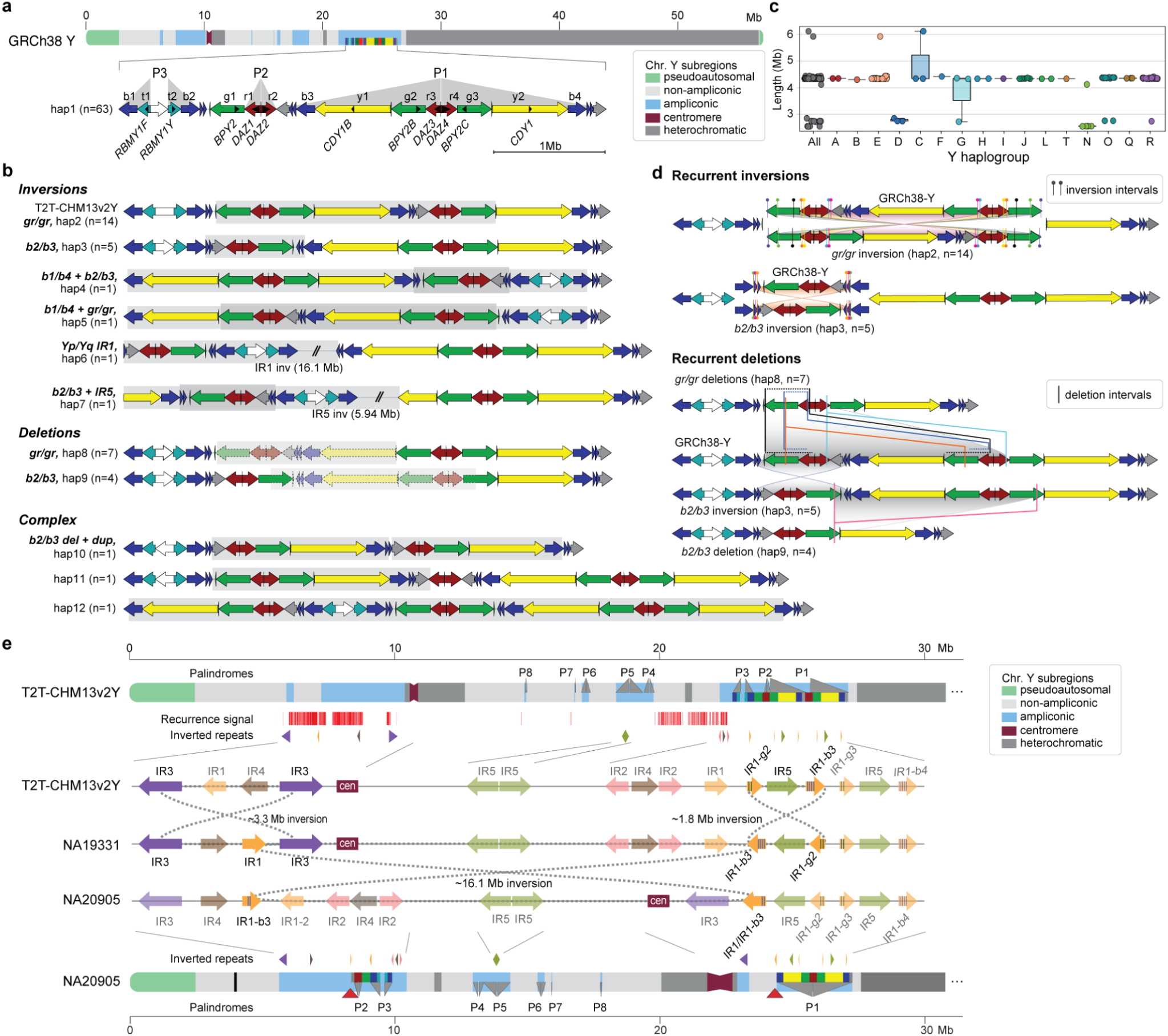
Structural diversity of the *AZFc* region and recurrent inversions across the euchromatic regions. **a**, Position and organization of the *AZFc* region in GRCh38. *AZFc* spans palindromes P1 - P3 and is subdivided into the classical amplicon families (b - blue, t - teal, g - green, r - red, y - yellow)^7^. Protein-coding genes are shown below. The GRCh38 haplotype is the most common *AZFc* structural haplotype among the 100 QC-passed Y assemblies. **b**, Additional *AZFc* structural haplotypes identified across 100 QC-passed Y assemblies, including inversions, deletions and complex rearrangements. Light grey boxes indicate approximate rearrangement intervals, and faded segments indicate the approximate location of deleted sequences. **c**, *AZFc* total length shown across all analyzed samples and across major haplogroups, with each point representing one sample. **d**, Intervals mediating recurrent *AZFc* inversions and deletions relative to the GRCh38 structure. *Top, gr/gr* and *b2/b3* inversion, *bottom*, *gr/gr* and *b2/b3* deletion; on distinct structural backgrounds. Lollipops indicate broader regions where the deletion breakpoint windows are likely located. **e,** Possible multistep history of the large inversion in NA20905. Schematic representation of the T2T-CHM13v2 structure (top) and the corresponding NA20905 haplotype (bottom), with the inversion recurrence signal detected from the pangenome graph, impacted palindromes and inverted repeats indicated. The intervening arrows show the inferred inversion events (IR3/IR3 inversion inverting the IR1 on Yp, *gr/gr* inversion inverting IR1-b3) required to transform the T2T-CHM13v2-like configuration into the NA20905 structure by permitting the final ∼16 Mb inversion.

The 12 haplotypes comprised the GRCh38-like configuration and 11 other architectures arising through six inversions, two deletions, and three complex events. The GRCh38-like configuration was present in 63/100 samples, followed by the *gr/gr* inversion (14/100; also present in the T2T-CHM13v2/HG002 Y) and the *gr/gr* deletions (7/100; **Fig. 2b**; **Suppl. Tables 16, 17).** We additionally identified recurrent *b2/b3* inversions in five chromosomes and *b2/b3* deletions in four chromosomes; the latter fixed among chromosomes in haplogroup N. The remaining seven haplotypes were each unique to a single sample. Importantly, placing these complete structures onto the Y-chromosome phylogeny revealed that several similar configurations arose independently on distinct Y backgrounds (**Suppl. Fig. 27**). Thus, the expanded panel transforms previously described rearrangement classes into a population-scale view of how often, and through which structural routes, *AZFc* is remodeled.

The phylogenetic breadth of the panel further allowed us to distinguish repeated inheritance of the same structure from independent recurrence and to compare breakpoint intervals across recurrent events. The recurrence of these haplotypes reflected repeated use of a small set of homologous substrates (**Suppl. Figs. 28-34)**. Among 14 chromosomes with *gr/gr* inversions, breakpoint intervals mapped to paired ampliconic repeat blocks and could be narrowed to a median of 5.9 kb (ranging 2.1 kb to 25.5 kb) (**Fig. 2d; Suppl. Fig. 28; Suppl. Table 16**). The estimated inversion sizes ranged from 1.34 Mb to 2.53 Mb (mean 1.87 Mb). In haplogroup J (Time to Most Recent Common Ancestor (TMRCA) estimate 37,050 years, 95% Highest Posterior

Density (HPD) 32,230 - 42,580 years), three independent *gr/gr* inversion events were identified among five chromosomes, demonstrating repeated formation of similar structures on closely related Y backgrounds. Likewise, the *b2/b3* inversions involved four independent breakpoint intervals clustered within or near a ∼30-kb segment, leading to inversions 80.9 kb to 88.9 kb in length (**Suppl. Fig. 29**), and consistent with a recurrent inversion hotspot (**Fig. 2d**). Recurrent partial deletions showed the same pattern: seven *gr/gr* deletions represented four independent deletion backgrounds, while the *b2/b3* deletion breakpoint was localised to a 1.2-kb region in three of four chromosomes (**Fig. 2d; Suppl. Figs. 32-34; Suppl. Table 17**). Together, these data show that recurrent *AZFc* diversity is repeatedly generated by non-allelic homologous recombination (NAHR) involving a restricted set of homologous repeat substrates.

To test whether the limited structural repertoire reflected a restricted accessible state space, we modeled the phylogenetic history of 98 chromosomes representing 10 of the 12 major *AZFc* architectures; two architectures whose defining inversions extended outside of the *AZFc* region were excluded (**Methods; Suppl. Results (*AZFc* architectural constraint and potential mechanisms); Suppl. Fig. 35**). Both equally parsimonious ancestral states (HG01890-like and GRCh38-like) contained 17 repeat blocks, whereas observed architectures ranged from 11 to 23 blocks. Under a permissive NAHR model allowing eligible homologous repeat blocks to undergo inversion, deletion, or reciprocal tandem duplication, simulated chromosomes showed significantly larger excursions in block copy number than observed (conservative Monte Carlo *P* = 0.003) (**Suppl. Table 18; Suppl. Fig. 36**). Thus, the observed *AZFc* architectures occupy a substantially narrower block-copy range than expected from rearrangement opportunity alone. Follow-up analyses did not identify a simple explanation for this restriction, including preferential loss of older rearrangements or preferential use of closer or longer homologous substrates (**Suppl. Results (*AZFc* architectural constraint and potential mechanisms)**).

Previous work showed that much of euchromatic Y-chromosome structural variation is driven by recurrent inversions, particularly within palindromic sequences and other inverted repeats^5,10,24^. To determine whether the recurrence resolved at base level within *AZFc* extended across the chromosome, we constructed a Y-chromosome pangenome graph and interrogated inversion-prone loci across the expanded assembly panel (**Suppl. Results (Analysis of inversion recurrence using pangenome graphs)**). We detected recurrent inversions at multiple known loci, including IR2-, IR3-, P7-, and P8-associated intervals (**Fig. 2e**), together with additional recurrent balanced inversions involving palindromic sequence (**Suppl. Table 19**). We also identified a ∼16-Mb pericentric inversion mediated by IR1 repeats in NA20905 (haplogroup J2b2a), whose structure is consistent with several sequential inversion events rather than a single rearrangement (**Fig. 2e, Suppl. Fig. 37**). Together, these analyses provide one of the largest and most phylogenetically diverse sequence-resolved surveys of *AZFc* architecture, allowing recurrent rearrangements to be distinguished from shared ancestry and their breakpoint substrates resolved across independent Y lineages. More broadly, they demonstrate that pangenome-graph approaches can identify recurrent inversions across highly repetitive and structurally polymorphic regions of the human Y chromosome that are poorly represented by a single linear reference.

### Multicopy fertility-related genes show distinct modes of paralogue and array remodeling

#### Paralog-resolved assemblies reveal asymmetric *DAZ* remodeling

The *AZFc* region typically contains four *DAZ* genes (*DAZ1-DAZ4*) embedded within palindromes P1 and P2 (**Fig. 2a**). Although *DAZ* copy-number changes and partial *AZFc* deletions have long been associated with spermatogenic failure and impairment^7^, the highly repetitive architecture of the locus has made it difficult to resolve how individual *DAZ* paralogues differ in structure and how they are remodeled across Y-chromosome lineages. The expanded assembly panel allowed us to move beyond total *DAZ* copy number and analyse individual paralogues at population scale. Across 100 QC-passed *AZFc* assemblies comprising 382 *DAZ* copies, we characterised three major features of variation: the copy number of the RNA-recognition motif (RRM), encoded by exons 2–6, the composition and copy number of exon 7 subtypes in the pre-Long Interspersed Nuclear Element (LINE) and post-LINE (L1PA2) segment, and broader gene architecture (**Fig. 3a; Methods; Suppl. Figs. 38-44; Suppl. Tables 20-21)**.

**Figure 3.**
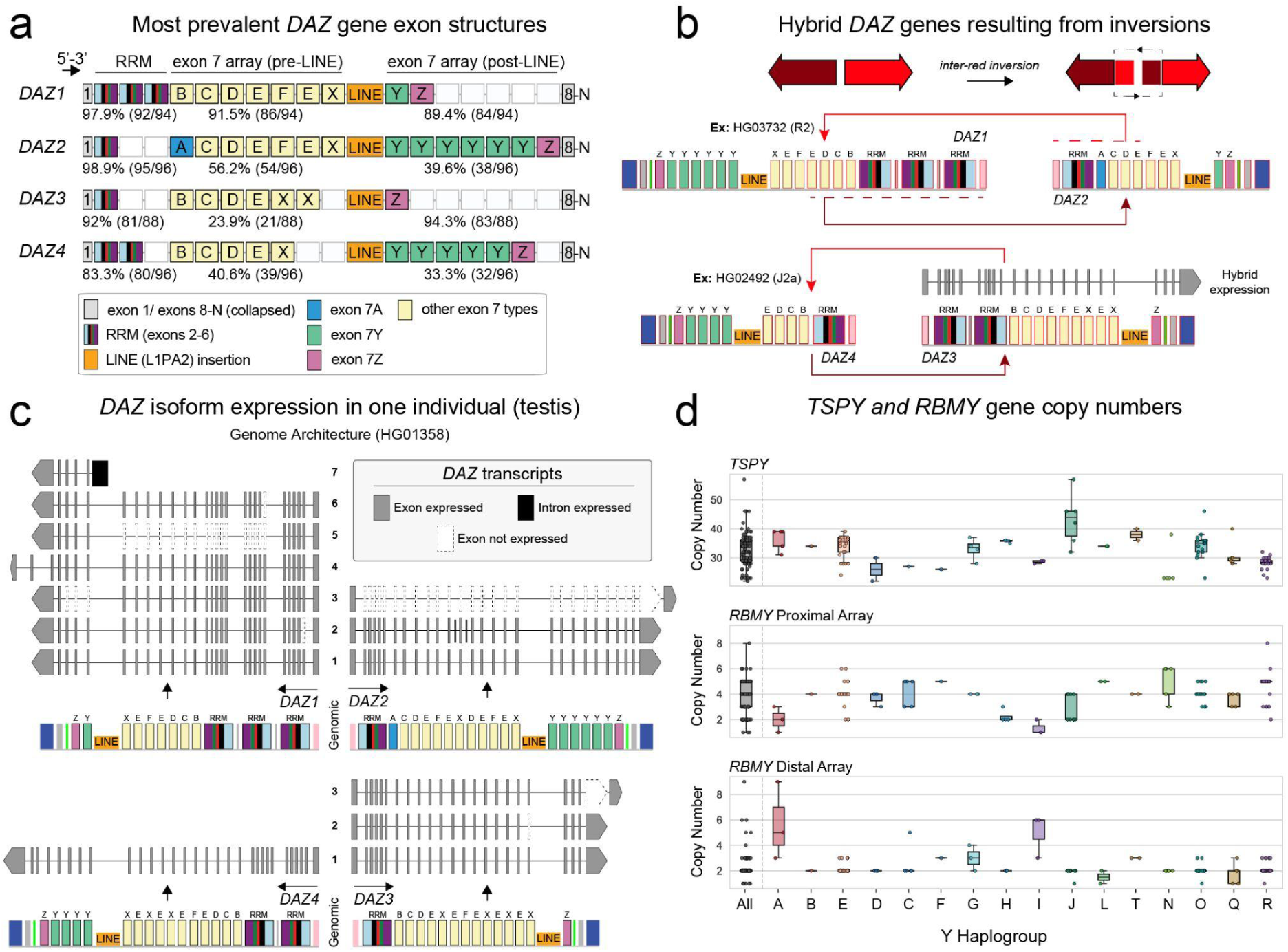
*DAZ* diversity, rearrangement and expression, and contrasting copy-number variation in *TSPY* and *RBMY*. **a**, Most prevalent *DAZ* exon structures of the four *DAZ* genes. Exons are colored by class, including RNA recognition motif (RRM) exons, exon 7 subtypes and the internal LINE (L1PA2) element. Percentages indicate the fraction of assemblies carrying the most prevalent exon block configuration. **b,** Representative hybrid *DAZ* genes in two individuals generated by reciprocal inversion events, showing preservation of 5′ RRM exons and reorganization of downstream exon 7 arrays between *DAZ1* and *DAZ2*, and exchange of RRM exons between *DAZ3* and *DAZ4*. Long-read Iso-Seq transcripts matching the hybrid gene in HG02492 were identified in the testis. **c,** Long-read kinnex *DAZ* isoforms from the testis of one individual. Reads aligned best to the HG01358 Y chromosome assembly. All four *DAZ* genes, and their respective exons, are expressed. **d,** Distribution of gene copy numbers for *TSPY* (*TSPY1* plus *TSPY2*) and *RBMY* across Y-chromosome haplogroups. *RBMY* copy numbers are shown separately for proximal and distal arrays. Boxplots indicate the median and interquartile range.

Despite extensive variation within the locus, several aspects of *DAZ* organisation were highly conserved. *DAZ1* was always linked to *DAZ2*, and *DAZ3* to *DAZ4*, with no mixing between these pairs (**Suppl. Fig. 42**). Among the 86 assemblies containing one copy of each paralogue, the *DAZ1/DAZ2* pair occupied the proximal position in 72/86 chromosomes (similarly to the GRCh38 reference) and the distal position in 14/86. At the level of individual genes, however, the four paralogues showed strikingly asymmetric structural stability (**Fig. 3a**). *DAZ1* was the most stable and almost always (92/94 gene copies) retained its canonical three-RRM structure, whereas *DAZ2* was almost uniformly (95/96 copies) a one-RRM gene, with a few changes to the RRM complement resulting from intragenic deletions and duplications (**Suppl. Figs. 40-42**). In contrast, *DAZ3* and *DAZ4* were substantially more variable. *DAZ3* usually contained one RRM (81/88 copies) and *DAZ4* usually contained two (80/96 copies). However, both genes were recurrently remodeled by reciprocal inversions, generating hybrid configurations in which *DAZ3* acquired a second RRM while *DAZ4* retained only one, as well as by intragenic deletions and duplications (**Fig. 3b; Suppl. Figs. 41-42, 44**). Because *DAZ* expression is largely restricted to male germ cells in the testis, we examined long-read testis Iso-seq data from independent datasets. These data identified transcripts from canonical *DAZ* paralogues as well as an inversion-generated hybrid *DAZ* gene, demonstrating that rearranged gene structures can contribute to the expressed DAZ repertoire (**Fig. 3b**; **Suppl. Fig. 45**). Consistent with this asymmetry, we observed only a single inversion in the *DAZ1/DAZ2* pair, but at least four independent *DAZ3/DAZ4* inversions among ten samples. Thus, population-scale paralogue resolution reveals that structurally adjacent *DAZ* copies experience markedly different evolutionary constraints.

The exon 7 arrays were substantially more polymorphic than the RRM-containing region (**Fig. 3a**), with recurrent tandem duplication and deletion generating extensive variation in copy number and subtype composition without altering overall gene order. Nevertheless, paralogue-specific patterns persisted: *DAZ1* and *DAZ3* retained relatively characteristic exon 7 architectures, whereas *DAZ2* and *DAZ4* were consistently more variable (**Fig. 3a; Suppl. Table 21**). Together with recurrent *DAZ3/DAZ4* inversions that generate mosaic paralogues (**Suppl. Figs. 41-42**), these data identify two distinct scales of *DAZ* remodeling: local expansion and contraction of tandem exon arrays and larger inversion-mediated exchange between paralogues.

Recurrent partial *AZFc* deletions further revealed non-random retention and re-establishment of *DAZ* architectures. The *b2/b3* deletion background in haplogroup N1a retained the canonical three-RRM *DAZ1* gene (**Suppl. Fig. 42**). Among four independent *gr/gr* deletion backgrounds, three retained or re-established a three-RRM *DAZ* copy: one through retention of *DAZ1* and two (in haplogroups D and O2a2a) through internal duplication within a *DAZ4*-like copy (**Suppl. Fig. 42; Suppl. Table 21**). In our dataset, such *DAZ4*-based re-establishment of a three-RRM copy was observed only in *gr/gr*-deleted chromosomes. Thus, partial *AZFc* deletions do not simply reduce *DAZ* copy number; across independent deletion backgrounds, three-RRM *DAZ* architectures were repeatedly retained or re-established through different structural routes.

### Complete arrays reveal distinct modes of *TSPY* and *RBMY* evolution

*TSPY* and *RBMY* are multicopy Y-linked gene families organized within large repetitive arrays, and substantial copy-number variation in both families has been described previously. *TSPY1* copies occur in tandem 20.4-kb repeat units, with a single *TSPY2* copy located outside the array, whereas *RBMY* genes are generally organized into proximal and distal arrays within the AMPL7 region together with two additional stand-alone copies in the P3 palindrome. Across QC-passed assemblies, *TSPY1* ranged from 22–57 copies (n=110) and *RBMY* from 5–12 copies (n=126) (**Fig. 3d; Extended Data Fig. 2b; Suppl. Fig. 46; Suppl. Tables 7-8**). The expanded set of sequence-resolved arrays allowed us to move beyond these known copy-number differences and determine how individual coding variants and array organization change across Y lineages.

Network analysis of predicted amino-acid sequences revealed contrasting evolutionary modes for the two families (**Methods; Suppl. Figs. 47–58; Suppl. Table 22**). *TSPY* copies retained similar coding sequences and relative positions within the tandem array, consistent with copy-number change occurring predominantly through local duplication or deletion of adjacent repeat units (**Suppl. Fig. 48-49**). In contrast, *RBMY* coding variants were more frequently reorganised within the proximal and distal arrays, and identical amino-acid profiles occasionally occurred in both arrays, consistent with inter-array sequence exchange (**Suppl. Fig. 54**). In both families, isolated copies outside the major arrays retained distinct coding identities. Thus, similar levels of multicopy variation can arise through fundamentally different structural processes: local tandem-array turnover in *TSPY* versus modular reorganization and sequence exchange in *RBMY*. Predicted upstream *cis*-regulatory motif profiles also distinguished array-associated from isolated copies in both families, suggesting that the structural separation of these copies is accompanied by divergence in local regulatory sequence (**Suppl. Results (Copy-number and motif variation of *TSPY* and *RBMY* genes); Suppl. Figs. 50–52, 55–58; Suppl. Table 22**).

Together, these analyses show that multicopy fertility-associated genes cannot be adequately described by aggregate copy number alone. Population-scale, copy-resolved Y assemblies reveal distinct evolutionary modes within each family: *DAZ* shows paralogue-specific constraint and recurrent exchange, *RBMY* undergoes modular array reorganization and inter-array sequence exchange, whereas *TSPY* varies primarily through local tandem-array expansion and contraction. This copy-resolved reference framework provides a basis for interpreting structural variation in Y-linked fertility genes that is obscured when these loci are represented by a single reference or by total copy number alone.

### Heterochromatic compartments drive much of Y-chromosome structural variation while preserving constrained organization

The centromere and Yq12 are the two major heterochromatic compartments of the human Y chromosome and contribute disproportionately to its structural and size variation across lineages. The centromere is typically composed of α-satellite repeats organized into higher-order repeat (HOR) arrays that assemble the kinetochore and mediate chromosome segregation^25^. On the human Y chromosome, the active centromeric array is formed by *DYZ3* α-satellite HORs and is among the smallest centromeric arrays in the human genome^13,21^. Complete centromeric α-satellite HOR arrays have historically been difficult to assemble, limiting population-scale analysis of their sequence, structural variation and evolution. Our expanded Y-chromosome resource provided 123 completely and accurately assembled centromeres, increasing Y centromere sampling by approximately 6.5 times relative to previous studies^2,10,21,26^. Across these centromeres, *DYZ3* array length varied 13.8-fold, from 121 kb to 1.67 Mb (mean 771 kb), representing, to our knowledge, the smallest and largest *DYZ3* α-satellite HOR arrays reported to date (**Fig. 4a; Methods**; **Suppl. Fig. 59**; **Suppl. Tables 7, 23**). Array length was strongly phylogenetically structured (Pagel’s λ = 1), with pronounced lineage-specific differences, including mean lengths of 1.06 Mb in haplogroup E and 401 kb in haplogroup R (**Fig. 4a; Suppl. Fig. 60a; Suppl. Table 24)**.

**Figure 4.**
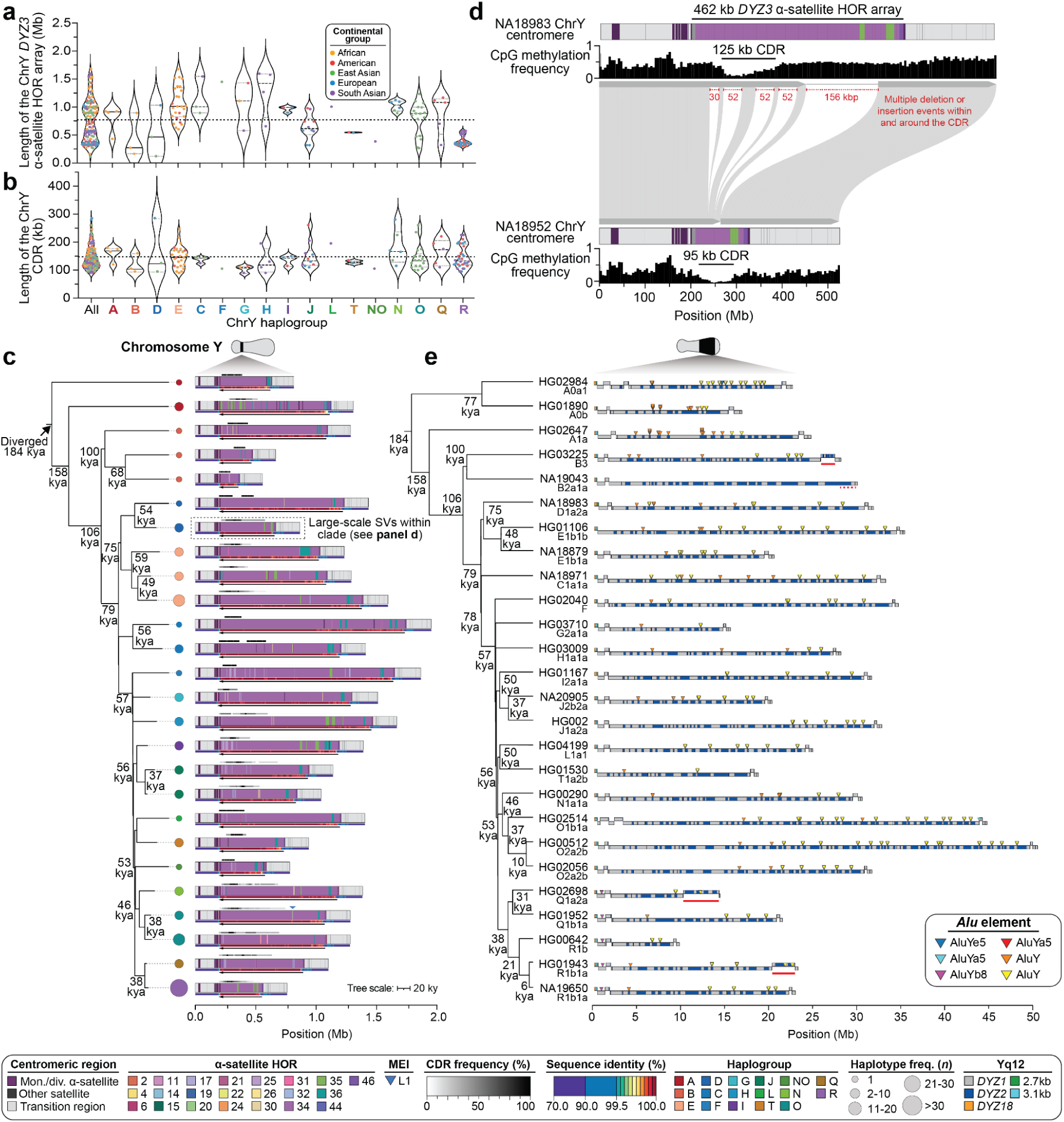
Sequence and structural diversity of Y centromeres and Yq12. **a,** Length distribution of the *DYZ3* α-satellite higher order repeat (HOR) array among Y chromosome haplogroups. Each point represents one assembly and is colored by continental group. Violin plots show the distribution within each haplogroup. Horizontal summary lines indicate the central tendency and spread, and the dashed line marks the mean across all samples. **b,** Length distribution of the putative kinetochore site, defined by the centromere dip region (CDR^28^), across Y chromosome haplogroups. Each point represents one assembly and is colored by continental group. Violin plots summarize the distribution within each haplogroup. **c,** Time-calibrated phylogeny of chromosome Y centromeres, with estimated divergence times shown on internal branches. Schematics on the *right* show the corresponding centromere structures, including α-satellite HOR composition, transition sequence, other satellite sequence, CDR frequency, and sequence identity. Dot color denotes Y haplogroup, and dot size indicates the number of assemblies sharing that centromeric haplotype. **d,** Example of extensive structural remodeling between two centromeres from the same clade (NA18983 and NA18952; clade 7 in panel **c; Suppl.** Fig. 59). A total of 342 kb of sequence is inserted or deleted within and around the CDR, altering both the local sequence composition and the span of the hypomethylated centromeric core. **e**, Yq12 structural diversity across a selected subset of samples, with a dated phylogeny shown on the left. Repeat arrays above the line are in direct orientation, whereas those below the line are in inverted orientation. Large distal inversions are indicated by solid orange lines and the deletion with a dashed line.

Despite this extensive sequence and size variation, the epigenetically defined functional centromere core was considerably constrained. Human centromeric arrays are generally highly methylated, except for a hypomethylated centromere dip region (CDR)^21,27,28^ enriched with CENP-A nucleosome, that provides a methylation-based proxy for the site of kinetochore assembly^27,29^. CDR length ranged from 90 to 285 kb across the 123 chromosome Y centromeres (3.17-fold; mean 148 kb), substantially less than the 13.8-fold variation in total *DYZ3* α-satellite HOR array length (**Fig. 4b; Suppl. Fig. 60**). Total *DYZ3* array length showed maximal phylogenetic signal (Pagel’s λ = 1), whereas CDR length showed essentially none (Pagel’s λ = 0) (**Suppl. Fig. 61a)**, and phylogenetic generalized least-squares analysis provided little evidence that CDR size increased with total *DYZ3* HOR array length (log–log slope = 0.064, 95% CI −0.024 to 0.153, *P* = 0.153) and strongly rejected proportional scaling between the two (*P* = 4.96 × 10⁻⁴² for slope = 1; **Suppl. Fig. 61b; Suppl. Table 24**). Thus, a twofold increase in *DYZ3* α-satellite HOR array length was associated with only an approximately 4.6% predicted increase in CDR length. Consistent with this decoupling, the observed 3.17-fold range in CDR size was smaller than all 10,000 simulations generated under the evolutionary size dynamics of the surrounding *DYZ3* array (median 19.80-fold, 95% interval 8.35–91.97; Monte Carlo *P* = 1 × 10⁻⁴; **Suppl. Fig. 61c; Suppl. Table 24**). This weak relationship between CDR size and total α-satellite HOR array length is consistent with centromere identity being epigenetically specified rather than determined simply by inherited α-satellite copy number, and parallels observations on other human chromosomes in which substantial differences in HOR array size are not accompanied by proportional changes in the CENP-A/kinetochore domain^30^. Thus, extensive lineage-specific expansion and contraction of *DYZ3* is largely decoupled from the size of its functional centromeric domain, although local sequence context may still influence where that domain is positioned^21^.

The depth of complete centromere sampling also allowed us to quantify sequence evolution within the *DYZ3* HOR array across Y-chromosome lineages. Using phylogenetically defined clades and one-to-one 10-kb alignments between complete centromeres, we estimated clade-specific α-satellite HOR mutation rates across 14 lineages. Rates centered around ∼5 × 10^−8^ mutations per bp per generation (median, 4.92 × 10^−8^), demonstrating substantial ongoing sequence turnover within the active Y-centromeric array (**Methods; Suppl. Fig. 62; Suppl. Table 25**), closely matching the recently reported chromosome-Y centromere estimate of 4.7 × 10^−8^ mutations per bp per generation^21^. Together with the strong phylogenetic structure in array size, these data show that the Y centromere evolves rapidly at both nucleotide and structural scales despite maintaining a comparatively stable epigenetic core. The pace of structural remodeling is apparent even among closely related Y chromosomes. Within the D1 clade, whose common ancestor is dated to ∼19,450 years ago, NA18952 carries a 121-kb *DYZ3* array whereas NA18983 carries a 462-kb array, with at least five insertion/deletion events of 33–156 kb distinguishing their centromeres (**Fig. 4c, d; Suppl. Fig. 59**). Across clades, sequence divergence also increased on transition from euchromatin into the *DYZ3* HOR array and declined again on entry into q-arm euchromatin, with the magnitude of this increase varying from 0.9-fold to 3.5-fold and averaging 1.7-fold overall (**Suppl. Fig. 62**). Thus, the Y centromere can undergo extensive nucleotide and structural remodeling over relatively short evolutionary timescales while retaining a single, comparatively size-constrained kinetochore-associated domain.

Yq12 is the largest heterochromatic compartment of the Y chromosome and the principal driver of chromosome-wide length variation. It is composed predominantly of the *DYZ1/HSat3* (single repeat: ∼3.5 kb) and *DYZ2/HSat1B* (single repeat: ∼2.4 kb) satellite families, whose large, highly repetitive arrays have historically limited sequence-level analysis. Our previous 43-Y study showed that Yq12 varies extensively in length while maintaining an approximately balanced *DYZ1*:*DYZ2* composition^2,10^. The expanded resource allowed us to resolve the internal array architecture of Yq12 at substantially greater scale and determine how this extreme length variation is generated and evolves across Y lineages. Among 84 QC-passing Yq12 assemblies, length ranged more than fivefold, from 10.3 Mb to 53.1 Mb (mean 27.1 Mb, median 25.7 Mb; **Fig. 4e**; **Ext. Data Fig. 2b**; **Suppl. Results (Yq12 analysis)**; **Suppl. Figs. 63-67**; **Suppl. Table 26**).

Despite this extreme length variation, the internal organisation of Yq12 was strikingly conserved. *DYZ1* and *DYZ2* remained near equal in abundance (mean *DYZ1:DYZ2* repeat-unit ratio = 1.004), 96.2% of repeat units shared the same orientation, and all 84 chromosomes contained matched numbers of *DYZ1* and *DYZ2* arrays arranged in alternating order (**Suppl. Figs. 63-64, 66-68)**. However, the number of arrays of each repeat family varied from 13 to 68 per chromosome (**Suppl. Table 26**). The conserved *DYZ1*:*DYZ2* balance observed previously reflects preservation of a higher-order alternating array architecture rather than simply a chromosome-wide average in repeat abundance.

Complete array resolution further revealed how Yq12 length polymorphism is generated. Variation in total *DYZ1/DYZ2* satellite-array length was driven predominantly by differences in array number rather than by changes in the mean array size of individual arrays (variance fraction = 1.144, 95% bootstrap interval 0.967–1.381) (**Suppl. Fig. 68; Suppl. Table 26**). Variance decomposition supported this distinction, with array number contributing substantially more to total length variation than mean array length, while a negative covariance between the two components indicated partial compensation between array number and size. Thus, the extreme variation in Yq12 size arises primarily through gains and losses of alternating *DYZ1/DYZ2* arrays rather than expansion or contraction of individual arrays.

Yq12 architecture was also strongly structured by Y-chromosome ancestry. Both array number and mean array length showed strong phylogenetic signals (Pagel’s λ = 1.00; **Suppl. Fig. 68; Suppl. Table 26**). Although Yq12 architecture evolved in a lineage-specific manner, changes in *DYZ1* and *DYZ2* abundance were tightly coupled (ρ = 0.911, 95% simulation interval 0.869–0.942; *P* = 3.19 × 10⁻³⁴), with 14.6-fold greater evolutionary variance in their shared abundance than in their relative proportions (**Suppl. Table 26)**. *DYZ2* nevertheless showed 1.82-fold greater evolutionary variance than *DYZ1* (*P* = 1.78x10⁻⁹). Together, these results indicate that Yq12 gains or loses large amounts of satellite DNA over evolutionary time, while predominantly preserving the coupled abundance and alternating organization of its two major repeat families.

Structural remodeling extended beyond satellite-array gain and loss. Most Yq12 assemblies (80/84, 95.2%) contained both previously described flanking inversions (*proximal*: ∼614 kb in size, *distal*: ∼472 kb in size;^10^ **Suppl. Fig. 67; Suppl. Table 26**). In three chromosomes, what initially appeared to be an expansion in a distal inverted segment was actually the result of nested inversions repositioning up to 4.5 Mb of sequence within the distal inverted interval (**Fig. 4e; Suppl. Figs. 67, 69-70; Suppl. Table 27**). These structures show that Yq12 evolution combines changes in satellite-array number with megabase-scale sequence reorganization, while preserving its broader compositional and orientational framework.

Pedigree sequencing provided direct evidence that Yq12 remains a major site of ongoing germline change. Across six father-son transmissions in the CEPH pedigree, 49 of 53 *de novo* variants mapped to Yq12, whereas only one SNV and one 4-bp insertion occurred in euchromatic sequence and two SNVs mapped to the *DYZ17* array (**Suppl. Results (*De novo* germline mutations)**; **Suppl. Figs. 71-72**; **Suppl. Tables 28-29**). Among 40 *de novo* SNVs in Yq12, grouping nearby variants yielded 35 nonredundant events, of which 26 (74.3%) were isolated SNVs most consistent with point mutation, while six (17.1%) were consistent with gene conversion events (**Suppl. Figs. 73-75**; **Suppl. Table 30)**. These transmission data independently demonstrate rapid sequence turnover within Yq12 and indicate that both point mutation and sequence exchange contribute to its ongoing evolution.

Together, the centromere and Yq12 reveal distinct forms of heterochromatic evolution within the same chromosome. The *DYZ3* centromeric array undergoes rapid sequence and size evolution while retaining a comparatively size-constrained epigenetic centromeric domain, whereas Yq12 can gain, lose and reposition megabases of satellite DNA while preserving the coupled abundance and alternating organization of *DYZ1* and *DYZ2*. Thus, extensive heterochromatic variation on the human Y chromosome occurs within higher-order architectures that are considerably more stable than the underlying sequence and copy-number variation.

### Haplotype-resolved methylomes reveal structure-associated epigenetic organization across palindromic and ampliconic sequence

To establish the structural framework for epigenetic analysis, we first characterized Y-chromosome palindromes across the expanded assembly panel. We recovered all eight canonical palindromes (P1–P8)^1^ and identified an additional conserved palindrome, P9 (median length 15.8 kbp; **Suppl. Results (Palindromes); Suppl. Tables 31-32**). P9 encompasses sequence previously annotated as the ∼12-kb direct repeat Rep1^31^ in GRCh38, but was consistently resolved as an inverted repeat across the assemblies analysed here. Outside P9 and the structurally dynamic *DAZ*-containing P1/P2 interval, palindrome architecture was highly conserved, with extremely high arm-to-arm identity across P3-P8 (median ranging from 99.787% in P8 to 99.988% in P4) and low repeat content (median RepeatMasked content from 1.7% in P4 to 4% in P7; **Suppl. Figs. 76-77**). In contrast, P9 had slightly lower arm-to-arm median identity (98.718%), and P1, P2, and P9 had relatively high repeat content (19.3%, 30.7%, and 14.1%, respectively) (**Suppl. Figs. 76-77**). The extended (∼3-fold) dataset analyzed here enabled us to study gene conversion between arms of the same palindrome in greater detail than in previous studies^10^. A phylogeny-based approach across seven palindromes (P3-P9) identified a total of 3,239 gene conversion events and a median length of gene conversion tract of 542 bp. Gene conversion had a significant GC bias (1,580 AT to GC events vs 1,088 GC to AT events, Chi-square test, *p* = 1.65×10^⁻21^) and a slightly weaker but still significant ancestral bias (905 reversals of the mutated allele vs 805 fixation events, Chi-square test, *p* = 0.0156; **Suppl. Results (Gene conversion)**; **Suppl. Figs. 78-79**; **Suppl. Table 33**). Thus, the Y palindromes undergo homogenization via gene conversion, preserving their near-perfect sequence identity and low repeat content.

The availability of matched assemblies and Oxford Nanopore methylation data allowed us to ask whether these structurally resolved repeat copies also occupy equivalent epigenetic states. We analysed 5-methylcytosine profiles from 134 EBV-transformed lymphoblastoid cell lines (LCLs), mapping reads to each individual’s own Y-chromosome assembly rather than to a single linear reference. This distinction is particularly important for palindromic and ampliconic sequences, where rearrangement and high copy identity can otherwise collapse or misassign reads. The resulting analysis therefore extends our previous reference-based Y methylation survey^2,10,21,26^ by resolving methylation in sample-specific structural and copy contexts. Because the available methylomes derive predominantly from LCLs, these patterns describe epigenetic organization in this cellular context and should not be interpreted as tissue-general Y-chromosome methylation states. Methylation analyses used the predefined canonical P1–P8 annotation framework; P9 was therefore excluded from these downstream comparisons.

Rather than exhibiting a uniform Y-chromosome methylation state, palindromes showed reproducible, locus-specific epigenetic profiles. Mean methylation and read-level methylation heterogeneity differed across Y chromosomal sequence classes and among individual palindromes (**Fig. 5a**; **Suppl. Results (Methylation of palindromes)**; **Suppl. Fig. 80**; **Suppl. Tables 7, 34**). P8 was consistently highly methylated, whereas other palindromes showed intermediate and more variable states (**Suppl. Fig. 80a**). Despite broadly conserved sequence composition (**Suppl. Fig. 80d**), palindrome arms exhibited GC and microsatellite asymmetry (**Suppl. Fig. 80e-f; Suppl. Table 35**), and reduced arm-to-arm sequence identity generally coincided with increased methylation heterogeneity (**Fig. 5b; Suppl. Table 34**), linking structural divergence to reduced epigenetic concordance. However, sequence divergence alone did not explain the observed patterns: P1, P3 and P4 showed persistent arm-specific methylation asymmetries despite high sequence similarity between their two arms (**Fig. 5c**). Our analysis shows that even palindrome arms maintained at near-identical sequence by gene conversion can occupy distinct epigenetic states, indicating that palindrome identity and local genomic context contribute information beyond primary sequence alone.

**Figure 5.**
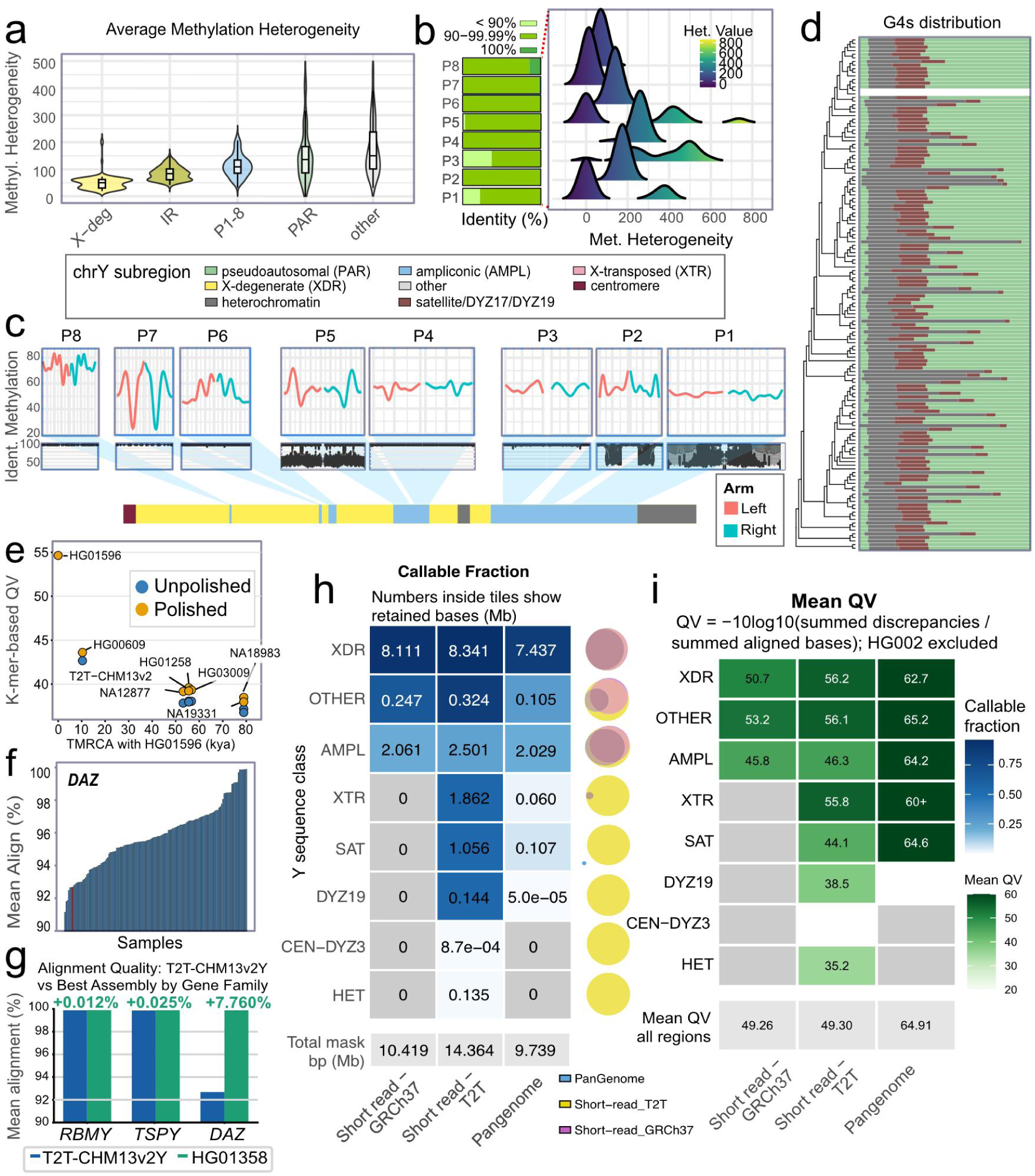
Repeat architecture, epigenetic variation, and reference-dependent analysis of the human Y chromosome. **a,** Methylation heterogeneity across Y chromosome sequence classes. Per-individual CalcHet scores were summarised within XDR, inverted repeats, palindromic, PAR, and other sequences. Violins show across-individual distributions; boxplots indicate medians and interquartile ranges. **b,** Arm-to-arm sequence identity and methylation heterogeneity across palindromes. Left: proportion of individual-specific P1–P8 palindromes in three arm-to-arm identity classes (100%, 90–99%, <90%). Right: corresponding distributions of methylation heterogeneity. **c,** CpG methylation profiles along paired palindrome arms (P1-P8). Methylation is shown across normalized arm position (0–100%) for the left (red) and right (blue) arms, with values across individuals summarized by 10 quantile trajectories per position bin. Arm-to-arm sequence identity is shown for individual assemblies. The schematic indicates palindrome positions within ampliconic (light blue) and X-degenerate-associated (yellow) regions. **d,** Distribution of predicted G-quadruplex (G4) motifs across 140 Y chromosomes, shown as local G4 density relative to major Y chromosome sequence classes **e,** Reconstruction accuracy for HG01596 using alternative Y chromosome starting haplotypes. K-mer-based QV is shown before (blue) and after (orange) polishing with HG01596 Illumina and low-coverage ONT data, plotted against the estimated TMRCA between HG01596 and each starting Y chromosome. Starting haplotypes comprised eight individual Y assemblies, including the phylogenetically closest available assembly HG00609, HG01596 itself, and T2T-CHM13v2Y. TMRCA is shown in thousands of years (kya). **f,** Distribution of *DAZ* transcript alignment quality across 140 individual Y chromosome assemblies and T2T-CHM13v2Y, measured as the average percentage of read nucleotides aligned. The red bar indicates the position of T2T-CHM13v2Y within the distribution. **g,** Improvement in mean transcript alignment quality for *RBMY*, *TSPY*, and *DAZ* using the best-performing assembly (HG01358) compared with T2T-CHM13v2Y. Alignment quality is the fraction of matched read bases among total read bases; values above the bars indicate the percentage-point improvement with HG01358. **h,** Callable fractions of T2T-CHM13v2Y sequence classes retained by three global callable-region masks. Tile values indicate retained sequence in megabases, with total callable sequence for each mask shown below. Euler diagrams show overlap in callable bases among masks within each sequence class. **i,** Phred-scaled short-read variant-calling accuracy within each callable mask, measured by discordance between variant-modified T2T-CHM13v2Y sequences and the corresponding assembled Y chromosomes. Regions where no discrepancies were observed, QV is reported as 60+, while categories without sufficient aligned sequence for QV estimation are shown as white boxes.

Ampliconic genes showed the same dependence on local structural context. Average methylation had only a modest, nonlinear relationship with exon-normalized copy number, whereas methylation heterogeneity was greater among lower-copy configurations (**Suppl. Fig. 81a-b)**. More importantly, copies of the same gene family were not epigenetically equivalent: *BPY2* and *VCY* methylation differed according to copy position, whereas *DAZ* methylation varied with ampliconic-block location, genomic copy order and inversion status (**Suppl. Fig. 81c-g)**. Across *AZFc*, overall methylation remained comparatively low across structural haplotypes (**Suppl. Fig. 80g**). Thus, structural variants in multicopy regions alter more than gene dosage: they reposition otherwise closely related copies within epigenetically distinct local contexts.

Predicted G-quadruplex-forming sequences provided an additional layer of sequence-context variation. Although G4 motifs are relatively depleted on chromosome Y compared with the autosomes^32^, their distribution varied markedly among Y sequence classes, individuals and palindromes, with reproducible local asymmetries between palindrome arms (**Fig. 5d; Suppl. Fig. 82; Suppl. Results (G4-motifs)**). These data establish a map of non-B-DNA-forming potential across structurally resolved Y chromosomes, although its relationship to methylation and structural instability remains to be tested directly.

Together, these analyses show that a highly repetitive Y-chromosome sequence is not epigenetically interchangeable. Personalized assemblies reveal a layered organization in which palindrome identity defines broad methylation states, structural divergence is associated with epigenetic heterogeneity, and local copy position further distinguishes individual ampliconic gene copies. By resolving sequence, structure and methylation on the same Y haplotypes, this resource provides a framework for determining how recurrent rearrangements alter not only gene and repeat copy number but also the chromatin contexts in which those copies reside.

### Diverse Y chromosome references reduce bias in complex locus analysis

The analyses above show that Y-chromosome diversity is strongly structured by lineage and concentrated in repetitive and ampliconic compartments that are poorly represented by any single haplotype. We therefore asked whether the expanded assembly panel could function not only as a catalogue of Y diversity, but as a reference framework for improving analyses when a sample-specific complete assembly is unavailable. We evaluated three applications: reconstruction of an individual Y chromosome from conventional sequencing data, interpretation of transcripts from multicopy ampliconic genes, and assessment of short-read copy-number and variant callability.

As a proof of principle, we first tested whether phylogenetic relatedness between the starting reference and target chromosome improves sequence reconstruction. We reconstructed the Y chromosome of HG01596 from Illumina and low-coverage ONT reads^33^ using the sample-matched HG01596 assembly and seven alternative starting sequences: the phylogenetically closest available assembly, HG00609 (TMRCA of 10,350 years, 95% HPD 8,540 - 12,330 years), five more distantly related Y assemblies and T2T-CHM13v2Y (**Methods; Suppl. Results ‘Polishing experiments’**). As expected, the sample-matched HG01596 sequence provided the highest reconstruction accuracy (QV 54.6). Among non-matched starting haplotypes, reconstruction accuracy decreased with increasing phylogenetic distance: HG00609, separated from HG01596 by only ∼10 kya, produced a polished QV of 43.6, whereas references separated by ∼53-57 kya yielded QVs of 39.2–39.6 and those separated by ∼79 kya yielded QVs of 38.0-38.5 (**Fig. 5e**). Polishing increased QV for all non-matched starting haplotypes, but did not eliminate the effect of starting-haplotype relatedness. Region-specific evaluation likewise showed that residual differences were concentrated in structurally complex compartments (**Suppl. Figs. 83-85**). Thus, even without a sample-matched long-read assembly, phylogenetic proximity of the starting Y chromosome substantially influences reconstruction accuracy, with representation of a closely related paternal lineage improving reconstruction from commonly available sequencing data.

Reference choice had an even larger effect on transcript interpretation at structurally variable ampliconic genes. We aligned long-read testis transcriptome data (SRR31360662) independently to 140 individual Y assemblies and T2T-CHM13v2Y and compared alignment performance for *DAZ*, *RBMY*, and *TSPY* ampliconic genes (**Methods; Suppl. Results (*RBMY*, *TSPY* and *DAZ* gene expression)**). Reference effects were modest for *RBMY* and *TSPY*, but pronounced for *DAZ*: for this donor, T2T-CHM13v2Y ranked in the bottom 5% of tested Y sequences for *DAZ* alignment quality, whereas HG01358 increased median aligned-read accuracy by 8.22 percentage points (adjusted P = 5.26 × 10^−22^; **Fig. 5f-g**; **Suppl. Figs. 86-91**; **Suppl. Table 36**). More importantly, reference choice altered copy-specific transcript interpretation: only 19.7% (30/152 reads) of *DAZ* read annotations agreed between T2T-CHM13v2Y and HG01358 (**Suppl. Figs. 90-91**). Mapping to the better-matching assembly resolved expression from all four *DAZ* paralogues and revealed previously unreported transcript structures, including intron retention, alternate transcription initiation in an antisense LINE element, and alternative terminal exons (**Fig. 3c; Suppl. Figs. 92-95**). Thus, at a locus where structural variation extends into gene architecture itself, a single complete Y reference can substantially distort both mapping efficiency and paralogue-specific transcript annotation.

The assembly panel also provides ground truth for extending analyses to cohorts lacking long-read assemblies. At the highly copy-variable *TSPY* array, normalized short-read depth was strongly correlated with the assembly-derived *TSPY1* copy number (R^2^ = 0.79; **Methods; Suppl. Fig. 96**), allowing copy number to be estimated in 367 GTEx samples^34^ with matched DNA and testis RNA sequencing. Despite extensive copy-number variation, TSPY1 copy number showed essentially no relationship with testis expression (R^2^ = 0.001; **Suppl. Fig. 97**), consistent with previous observations that dosage alone does not determine expression of Y ampliconic genes^35^. Nevertheless, complete assemblies can be used to calibrate scalable measurements of otherwise difficult multicopy loci while also providing the structural context needed to interpret those measurements.

Finally, we used the assembled chromosomes as independent ground truth to quantify how callable-region definitions trade genomic breadth against variant-calling accuracy. We compared three widely used callable-region masks derived from short read (GRCh37^36^), T2T-reference (T2T-CHM13v2Y^2,37^), and pangenome^38^ approaches. We compared the two newer masks to a mask originally generated for large scale Y chromosome analyses that has been used broadly in the field. The T2T-CHM13v2Y-based mask retained the greatest amount of sequence, whereas the pangenome-based mask retained fewer bases but achieved higher average variant-calling accuracy (**Fig. 5h-i; Suppl. Fig. 98; Suppl. Table 37**). Differences were modest in stable X-degenerate regions, where all three kept most callable bases and had high QV. In contrast, complex regions like satellites, heterochromatin, X-transposed sequence, and ampliconic loci showed bigger differences in both the amount of sequence kept and accuracy, and none of the masks recovered callable centromeric sequence. Thus, short-read Y-chromosome callability involves a trade-off between breadth and accuracy: the T2T-CHM13v2Y-based mask retains more sequence, whereas the pangenome mask is more restrictive but captures higher-confidence sites, with the most repetitive regions remaining dependent on assembly-based approaches. Further, the historic GRCh37 mask built using an incomplete Y chromosome and short reads still has a mean QV of 48.99, comparable to the modern short read mask which has a mean QV of 49.17, though both are lower than the pangenome mask with a mean QV of 61.9.

Together, these analyses demonstrate that the Y assembly panel has utility beyond cataloguing sequence diversity. Phylogenetically related haplotypes can improve reconstruction when only conventional sequencing data are available; diverse assemblies reduce mapping and annotation bias at structurally polymorphic genes; and complete sequences provide ground truth for calibrating copy-number estimators and evaluating short-read callability. The resource therefore provides a framework for incorporating Y-chromosome structural diversity into genomic and transcriptomic analyses that remain poorly served by a single linear reference.

## Discussion

The human Y chromosome has remained underrepresented in population-scale genomics because its highly repetitive architecture has hindered complete assembly and accurate genotyping across individuals. Our 142 curated assemblies span 17 broad haplogroup clades and more than 180,000 years of Y-chromosome evolution, including at least one T2T assembly for 13 clades and 46 fully T2T chromosomes overall: more than a threefold expansion of our previous sequence-resolved surveys^2,10^. The value of this expansion lies not only in scale, but in what increased completeness and phylogenetic sampling make resolvable at sequence resolution. Earlier studies established extensive structural polymorphism of the Y chromosome^1,2,5–9^, and the first complete sequence and our 43-Y survey resolved previously inaccessible ampliconic and heterochromatic regions^2,10,39^. The expanded panel now allows recurrent structures to be distinguished from shared ancestry, variation to be assigned to individual paralogues and repeat arrays rather than aggregate copy number, and complete centromeric and heterochromatic architectures can be compared across many paternal lineages. By integrating these assemblies with methylation and transcript data and using them as empirical benchmarks for reference-based analyses, the resource additionally provides a framework for determining how Y-chromosome structural diversity affects downstream genomic interpretation. Together, the assemblies and accompanying annotations bring the Y chromosome’s most repetitive and structurally dynamic compartments - including *AZFc*, multicopy fertility genes, palindromes, the centromere and Yq12 - into a population-scale, sequence-resolved framework.

These assemblies show that even the most structurally variable regions of the Y chromosome do not evolve without constraint. *AZFc*, where large-scale structural variation and its association with spermatogenic impairment have been studied for decades^1,5–9,35,39^, illustrates how the expanded resource moves from documenting polymorphism to resolving the processes that generate and shape it. Similar architectures repeatedly arise independently through a restricted set of homologous substrates, and the observed architectures occupy a significantly narrower block-copy-number range than expected under a permissive NAHR model. Paralogue-resolved analysis further shows that aggregate *DAZ* copy number masks substantial structural heterogeneity: the four *DAZ* genes differ in their stability, RRM complement and exon 7 architecture, with recurrent inversions, duplications and deletions generating distinct and sometimes hybrid gene structures. Independent *AZFc* deletion backgrounds nevertheless repeatedly retained or re-established a three-RRM *DAZ* architecture. Testis Iso-seq data further showed that inversion-generated hybrid *DAZ* genes can be transcriptionally expressed, demonstrating that structural remodeling can alter the expressed paralogue repertoire. Together with the presence and expression of three-RRM *DAZ* genes in great apes^40^, the recurrent preservation or re-establishment of this architecture suggests that the configuration may be functionally relevant. These findings also provide a potential structural basis for some of the variable reproductive consequences associated with partial *AZFc* deletions: chromosomes assigned to the same broad deletion class can retain different *DAZ* paralogues and exon architectures, while their effects may further depend on Y-chromosome background. Consistent with this, fixed *gr/gr* deletions in haplogroup D and *b2/b3* deletions in haplogroup N appear to be tolerated in those lineages^41,42^, whereas similar partial *AZFc* deletions in other backgrounds are associated with spermatogenic impairment. Large population studies likewise show markedly different fertility effects among recurrent *AZFc* deletion classes, emphasizing that these rearrangements are not functionally equivalent^41^. Previous population-genetic analyses inferred that purifying selection affecting repetitive or ampliconic Y sequence could help explain the chromosome’s unusually low diversity^43^, while copy-number analyses suggested that selection has countered high mutation rates to preserve ancestral amplicon dosage across deeply divergent Y lineages^31,44^. Together, these results indicate that the limited repertoire of *AZFc* architectures reflects both a mutational bias toward recurrent rearrangement pathways and differential persistence of the resulting configurations, potentially reflecting functional or selective constraint - complementing decades of clinical and population-genetic work on *AZFc* deletions and spermatogenic failure with a sequence-resolved structural framework.

The heterochromatic compartments reveal a complementary relationship between rapid sequence turnover and stable higher-order organization. The first complete Y chromosome and our 43-Y study established the alternating organization and extreme length variation of Yq12^2,10^, while recent population-scale centromere studies have demonstrated extensive structural and epigenetic diversity across human centromeres^13,21^. The expanded Y-specific panel now resolves how these features evolve across lineages. At the centromere, *DYZ3* array length is strongly phylogenetically structured and varies by more than an order of magnitude, yet the hypomethylated kinetochore-associated domain varies over a much narrower range and shows little phylogenetic signal. This decoupling is consistent with centromere identity being epigenetically specified rather than simply scaling with inherited α-satellite copy number^21,27,28,45^, while remaining compatible with an influence of local sequence context on kinetochore position. Yq12 shows a parallel but distinct form of organization, building on the previously established, approximately balanced *DYZ1:DYZ2* composition: extreme length polymorphism arises predominantly through differences in the number of whole alternating arrays rather than uniform expansion or contraction of individual arrays, and that evolutionary changes in *DYZ1* and *DYZ2* abundance are tightly coupled. Thus, both major heterochromatic compartments accommodate extensive sequence and copy-number evolution while preserving organizational features that are considerably more stable than the underlying repeats.

Haplotype-resolved methylation adds an epigenetic dimension to this architecture. Unlike our previous reference-based analysis^2,10^, methylation here was assigned to each individual’s own Y assembly, allowing highly similar palindrome arms and ampliconic copies to remain structurally distinguishable. The resulting data show that repetitive copies are not epigenetically interchangeable: palindromes retain characteristic methylation states, structural divergence is associated with increased methylation heterogeneity, and individual copies of multicopy genes differ according to position and local architecture. This more heterogeneous landscape contrasts with the comparatively uniform pattern in our earlier analysis and may reflect, at least in part, the ability of personalized assemblies to preserve copy- and context-specific signals that can be collapsed by mapping to a single reference. These patterns also raise functional questions that can now be tested. Epigenetic regulation of heterochromatin is broadly important for maintaining repetitive-DNA stability, and loss of DNA methylation at repetitive elements is associated with chromatin disruption and genome instability^46,47^; whether the Y-specific methylation states identified here similarly influence aberrant recombination or repair remains unknown. Likewise, the association between methylation and ampliconic copy position raises the possibility that rearrangements alter regulatory context as well as dosage. This is particularly relevant because previous matched genome-testis transcriptome analyses showed that copy-number variation within Y ampliconic gene families explains relatively little of the expression variation among individuals^35^. The present LCL methylomes cannot establish such regulation in germ cells, but they identify copy- and structure-specific epigenetic states that can now be tested directly using tissue-matched, haplotype-resolved germ-cell epigenomes and transcriptomes.

A second major use of the panel is as an alternative-reference and benchmarking resource. Human pangenome studies have already shown genome-wide that representing multiple haplotypes reduces mapping and variant-discovery errors relative to reliance on a single linear reference^12,13,11^. The Y presents an especially acute version of this problem because much of its biologically relevant variation involves duplicated, inverted or repetitive sequences. In the present study, reference choice substantially altered *DAZ* transcript alignment, paralogue assignment and isoform annotation, demonstrating that even a complete Y chromosome can be an inappropriate reference for a structurally divergent haplotype. Complete *TSPY* arrays provide ground truth for calibrating scalable short-read copy-number estimates, and comparison of callable-region masks against assembled chromosomes shows that short-read analysis continues to trade genomic breadth against accuracy in the most complex Y compartments. The single-target reconstruction analysis further provides proof of principle that a phylogenetically related Y assembly can improve reconstruction from conventional sequencing data, although validation across additional target chromosomes will be needed before this can be generalized. Ultimately, these assemblies also provide a foundation for Y-specific pangenome graph references, in which alternative structural haplotypes could be represented simultaneously rather than selected one at a time. Graph-based human references already improve mapping and variant discovery across much of the genome^11,12^, and multiscale pangenome graph approaches have already shown that complex Y-linked loci such as *DAZ1-DAZ4* can be represented and compared across structurally diverse haplotypes^48^. However, accurately representing the full spectrum of Y structural diversity remains particularly challenging because its high-identity palindromes, multicopy arrays and megabase-scale rearrangements generate highly complex and potentially ambiguous graph structures. The sequence-resolved haplotypes provided here therefore offer both the substrate for developing such representations and a benchmark for determining where graph-based approaches improve on, or remain limited relative to, individual Y assemblies. The practical shift is therefore from asking whether a locus is represented in the reference to asking how best to represent the structural diversity relevant to the analysis - through an appropriate Y haplotype, a panel of haplotypes or, ultimately, a validated Y pangenome graph. For male-infertility studies, population and evolutionary genetics, clinical variant interpretation and future Y-pangenome analyses, investigators can now select among phylogenetically and structurally diverse sequence-resolved Ys - or use the panel collectively - rather than assuming that one linear Y represents the species.

Several limitations define priorities for future work. The panel was assembled from samples selected to broaden available Y diversity rather than to estimate population frequencies, and deep lineages including A00 remain absent. Region-specific assembly and QC requirements also reduce the number of chromosomes available for the most repetitive compartments, raising the possibility that particularly difficult architectures remain underrepresented. Because assembly success is itself likely to depend on repeat complexity, frequencies estimated from region-specific QC-passed subsets should therefore not be interpreted as unbiased population frequencies. Functional interpretation is similarly constrained by the available datasets: methylation derives from EBV-transformed LCLs, the long-read transcript analysis uses an independent testis donor, and the reconstruction experiment currently tests one target chromosome. Larger phenotyped cohorts, tissue-matched epigenomic and transcriptomic data and direct functional studies will therefore be required to determine which recurrent structures influence spermatogenesis or other phenotypes. The contribution of this resource is to provide the structural and phylogenetic framework for those studies: regions previously represented by an unresolved gap, an aggregate copy-number estimate or a single haplotype can now be interrogated as complete, lineage-resolved architectures. More broadly, the panel provides a foundation for incorporating Y-chromosome structural diversity into reproductive, population and clinical genomics rather than treating a single Y sequence as representative of the chromosome.

## Methods

### Samples

The majority (132/144) of the samples originate from the 1000 Genomes Project (1kGP) Diversity Panel^49^, three (NA24385/HG002, NA24149/HG003 and HG005) commonly used for benchmarking by the Genome in a Bottle (GIAB) consortium^50^, and nine from the CEPH1463 family^14,15^. With the exception of the CEPH1463 pedigree samples, the sequencing data used for Y-chromosome assembly were generated from Epstein-Barr virus (EBV)-transformed lymphoblastoid cell lines (LCLs). Sequencing data for the CEPH1463 samples were obtained from the previously published pedigree studies^14,15^.

*The Human Pangenome Reference Consortium (HPRC) samples (n=105).* The HPRC samples were selected using a variation-based sampling strategy designed to maximize representation of common human genetic variation^12^. The sequencing data used for the assemblies were generated from 1KGP LCLs and are part of the HPRC release 2 dataset of the (https://humanpangenome.org/hprc-data-release-2/)^12^.

*Genome in a Bottle (GIAB) samples (n=3).* Two GIAB samples, HG002 and HG005, were included as positive controls of the assembly pipeline in the HPRC release 2 dataset. One additional sample, HG003 was assembled and the Y chromosome was curated as part of this study using sequencing data from Hansen et al.^51^.

*The Human Genome Structural Variation Consortium (HGSVC) samples (n=29).* The HGSVC samples were selected to maximize genetic diversity (see ^13^ for details), while three samples (NA18989, HG01890, HG00358) were included to represent specific Y haplogroups^10^. The sequence data used for the assemblies was derived from^13^, with two exceptions. i) Additional PacBio HiFi data was included from the HPRC efforts for 20 samples (**Suppl. Table 4**), and ii) for 27/29 samples, re-base called UL-ONT with Guppy V6.5.7 model was used (**Suppl. Table 3**).

*The CEPH1463 family (CEPH) samples (n=9)*^14,15^. Two CEPH pedigree samples are restricted access only (NA12883 and NA12884, available from dbGaP^52^ (https://dbgap.ncbi.nlm.nih.gov/home/) and with the exception of assembly QC, sequence class annotation and *de novo* mutation analysis were excluded from other downstream analysis.

### *De novo* assembly generation and evaluation

#### Verkko assembly of the genomes

The Y chromosomes were curated from Verkko assemblies that were generated in parallel to the HPRC r2^53^. In brief, a Verkko v2.2.1^17^ assembly of each genome was made using automated scripts to download data from AWS, build parental k-mer databases, and run Verkko with both Hi-C and Trio data in parallel. Standardized post-processing and QC was performed with the same pipeline which used Yak^54^ to measure QV (k=31) and switch error rate, compleasm^55^ for gene completeness, and computed T2T contig and scaffold statistics using seqtk to identify the telomeric and gap sequences in the assemblies. For each sample where a trio and Hi-C was available, the assembly with the most T2T contigs was selected. In two cases (HG02602 and HG02015) the assembly with a lower T2T contig count was selected because it had a better gene completeness score than the alternative. The pipeline was repeated with 29 HGSVC samples and HG003, all of which only had Hi-C available.

### Chromosome assignment and orientation

Mashmap v3.1.1^56,57^ was used to map the assembly to T2T-CHM13v2Y. The contigs were assigned to chromosomes if they had mappings over 99% identity covering more than 10% of the reference chromosome in a given orientation (forward or reverse-complement). If a contig matched over 10% of more than one chromosome, multiple assignments were noted and the longest one was used. The XY pseudo-autosomal regions (PAR) and distal regions of the acrocentric were excluded from the covered fraction count as they recombine between individuals and cannot be accurately assigned via reference alignment. Therefore, these regions could only be named as a chromosome if they were assembled together with a unique region of the chromosome.

In parallel, the homopolymer-compressed Verkko gfa were aligned using Mashmap v3.1.1 to a compressed version of T2T-CHM13v2Y to assign the individual nodes to a chromosome. A two pass assignment was performed. First, nodes with mappings over 99% identity and 5 Mb matches to a chromosome were assigned to that chromosome. Second, if a connected component had no nodes assigned by the first criteria, nodes over 500 kb were assigned. This identified connected components in the assembly graph belonging to each chromosome.

#### Assembly evaluation and QC

*Identifying the main Y sequence.* Initially identified Y sequences in refOriented.fasta often incorrectly originated from X or other chromosomes, or mis-oriented part or the entire Y sequence. This is primarily due to the PARs and other repeats within the ampliconic sequences, combined with alignment orientation issues due to palindromic arms that lead to the mis-orientation when the Y sequence was incompletely assembled into multiple sequences.

To address these issues, a new “main Y” sequence was identified by mapping contigs to a masked version of T2T-CHM13v2Y. This masking specifically targets PAR, heterochromatic sequences, and ampliconic sequences (as detailed in^2^). This approach ensures the proper orientation and ordering of identified sequences, and facilitates the identification of samples that require further manual curation.

The specific regions masked were taken from the following files from https://s3-us-west-2.amazonaws.com/human-pangenomics/T2T/CHM13/assemblies/annotation:

- PAR and HET: chm13v2.0_chrXY_sequence_class_v1.bed
- Inverted Repeats (IR, P1-5): Entire chm13v2.0Y_inverted_repeats_v2.bed
- Amplicons: chm13v2.0Y_amplicons_v2.bed

For data retrieval and uploading, aws cli v2.15.26 was used. For sequence indexing and manipulation, samtools v1.21^58^ and seqtk v1.4 (https://github.com/lh3/seqtk) were used. Initially, Bedtools v2.31.1^59^ was used to merge regions for masking. Masked regions are replaced with “N”s using seqtk seq -M mask.mrg.bed -n N options. Then, refOriented.fasta contig were aligned to this masked reference using minimap2 v2.28^60^ with -x asm10 --eqx -c parameters to allow multiple alignments to the Y. Post-alignment, Rustybam v0.1.33 was employed to invert reference-query coordinates. Alignments were filtered to exclude short hits (<10 kb) and those falling within other repetitive regions: AMPL, bSat and COMP MER5A1.

Alignment blocks were further excluded if the sequence was larger than 90 Mb in length to exclude X-fused Y sequences (mis-assembled) and merged for its minimum and maximum coordinate along with its mapped direction. Each sequence block was reported and evaluated if 1) it was originally identified as part of chromosome Y, or 2) it was originally not identified as any chromosome (chr prefix) and the entire Y coverage was larger than 70%, or 3) the block mappable coverage was larger than 10% and the sequence was shorter than 30 Mb, or 4) it extended mapping over PAR2, the length was shorter than 30 Mb and the entire Y coverage was larger than 0.8%. Lastly, each alignment was re-evaluated to filter out nested or duplicated sequences. Sample name, sequences and its direction was kept for final scaffolding.

Telomeres and presence of gaps were identified using the seqtk gap -l1 and seqtk telo -d 5000 command to verify the completeness of the p and q arms. A second round of alignments of the refOriented.fasta to the original T2T-CHM13v2 was employed to search for the missing PARs for genomes with missing telomeric ends. This helped find the missing PAR1 sequence in HG01106 and HG02145 and PAR2 in HG01252, which was added to the list for scaffolding. There were no sequences mapped to PAR1 in HG01358, HG01952, HG03209, NA18608, and NA20850. Likewise, nothing matched the PAR2 in HG01433, HG02145 and NA21093, hence left with missing ends. At the end, 200 kb N-gap was placed to scaffold the “main Y” sequences unless the main Y was composed of one sequence.

*Identifying unlocalized Y sequences.* Following the identification of the Y scaffolds, any node in the assembly graph connected to the nodes of the main Y scaffold that could not be ordered or oriented unambiguously was extracted as a potential unlocalized Y sequence. This step aimed to recover centromeric and ampliconic repeats that Verkko could not resolve automatically in the first place. Furthermore, components within the assembly graph that were not connected to the Y scaffolds but showed strong matches to chrY were also examined and identified as an unlocalized Y sequence. Nodes with strong matches to the Y chromosome were identified by aligning their sequences to the homopolymer-compressed T2T-CHM13v2 using Mashmap3. A node was assigned to the Y chromosome if it met three criteria: it was not already part of the Y SCAFFOLD, it was not connected to rDNA, and its best alignment hit was to the Y chromosome.

An in-house script (CollectYPathAndNodes.py and RemoveNoisyNodes.py) was developed to identify additional nodes belonging to the Y-chromosome and paths within the assembly graph, corresponding to the Y scaffolds (SCAFFOLD). This process included a step to filter out ’noisy’ nodes. In brief, a node was classified as ’noise’ if it met the following criteria, based on comparisons to the average coverage of the SCAFFOLD nodes, edge counts, and non-overlapping sequence length:

- Connectivity: The node had only one incoming and one outgoing edge, or was a singleton with fewer than two edges.
- Coverage: The node coverage was greater than 0 (where 0 indicates the node was used multiple times elsewhere in the graph) and less than half of the average node coverage.
- Size: The unique sequence length was less than the length of a HiFi read; 13 kb (derived from a 20 kb HiFi read length divided by an estimated homopolymer compression factor of 1.5). This minimum length was halved if the node had only a single edge.

The final unlocalized sequences are extracted from the original assembly.fasta file by targeting any nodes classified as UNLOC or UNLOC_Cmpnt from the script above. Note that Verkko’s own criteria may lead to the removal of isolated singleton contigs, meaning some expected ’unlocalized’ sequences might not be present in assembly.fasta. Sequences that are identified and remain are then renamed. They are given a new identifier: ${sample}_chrY_randomXXXXXXX. The XXXXXXX portion is derived from the unique identifier originally assigned to the sequence by Verkko as it appears in assembly.fasta.

*Manual graph curation.* During the identification of unlocalized contigs, we found that some T2T Y contigs contained sequences that were not integrated into the scaffold path. This was caused because of:

- Hi-C scaffolding errors: Misclassifying highly similar repeat units as distinct haplotypes, therefore avoided when walking through the path.
- Rukki errors: Choosing a sub-optimal, shorter path, which included leaving loops in tandem repeats unwalked or following a shorter path through a “noise” node instead of the correct alternate path.

These issues were frequently observed in centromeric, ampliconic, or heterochromatic regions.

The developers of Verkko investigated these problems, and subsequent improvements were incorporated into Verkko v2.3.

The manual curation involved reconstructing paths using the sample’s assembly graph, sometimes extending to the hifi graph. For unwalked hairpin-like connections, ONT-UL read support was used to determine the correct orientation. When support was equal for both possible walks, a cautious approach was taken to prevent mis-orientation: the sequence corresponding to the hairpin-like path was designated as unlocalized, resulting in a gap in the main Y sequence.

Consequently, the initial T2T contigs underwent modification. This resulted in either the addition of previously unused nodes to the sequence or the insertion of gaps where unlocalized sequences belonged. Therefore, some of these modified sequences were no longer considered complete T2T contigs after curation.

In total, 36 HPRC, 19 HGSVC and 6 CEPH Y assemblies underwent T2T-CHM13v2-guided scaffolding or manual graph curation. **Suppl. Table 5** is a summary of the resolution. Final Y-chromosomal assemblies were classified into three categories. T2T assemblies were defined as a single contig with no unlocalized contigs, no N-gaps, and the presence of both Yp and Yq telomeric sequences. HG01890 was included in this category despite two small unlocalized contigs, which are attributable to cell-line mosaicism. T2T-scaffold assemblies contain both telomeric sequences in the main Y but include one or more N-gaps and the corresponding unlocalized contigs. All remaining assemblies were classified as other.

The source code used for identifying the main Y scaffold and unlocalized Y sequences is accessible on the T2T-chrY GitHub repository (https://github.com/arangrhie/T2T-chrY). For manual curation of the graph, we recommend using Verkko-Fillet^61^, a tool developed during the analysis phase of this manuscript.

*Comparison to HPRCr2 hifiasm assemblies.* The HPRC released a set of 232 diploid hifiasm assemblies in May, 2025 as part of Data Release 2 (https://humanpangenome.org/hprc-data-release-2/^11^). To annotate suspect regions of our HPRC main Y scaffolds and unlocalized Y sequences, we first combined the Y sequences for each sample with all the non-Y verkko2 scaffolds from the source assembly used for that sample. These whole-genome, diploid verkko2 assemblies with curated Y sequences were then compared to the same sample’s HPRC release 2 hifiasm assembly using the “assemblycompare” command from GQC^51^ (https://gqc.readthedocs.io/en/latest/). Specifically, for each sample, we ran:

assemblycompare -t 2 --q1fasta $HIFIASMHAP1FASTA --q2fasta

$HIFIASMHAP2FASTA --r1fasta $VERKKOHAP1FASTA --r2fasta

$VERKKOHAP2FASTA -p $SAMPLE\_HPRC_vs_verkko4bestcurY -Q

$SAMPLE\_hifiasm -R $SAMPLE\_verkko2 --aligner minimap2 --splitdistance 500

--minalignlength 100000

GQC assemblycompare compares the first phases of the query assembly (in this case the hifiasm assembly) against the reference assembly (in this case, the Verkko assembly) using a hidden Markov model guided by reference-haplotype-specific k-mers present in the query. It then uses primary alignments of each query haplotype block to the appropriate reference haplotype to find and report regions in the reference’s coordinates that are covered by primary alignments of the query assembly with discrepancy rates lower than 0.1% (in a “Q30” BED file), 0.01% (in a “Q40” BED file), or 0.001% (in a “Q50” BED file). These “clean” regions can then be subtracted from the reference genome to determine suspect regions, i.e., regions whose consensus is not supported by the HPRC hifiasm assembly for that sample. These BED files are available https://s3-us-west-2.amazonaws.com/human-pangenomics/index.html?prefix=T2T/scratch/chrY/GQC.

To determine the set of all hifiasm assembly scaffolds for each sample which contain Y sequences, we parsed the unmerged “querycovered.bed” output file from the above GQC assemblycompare run to print all hifiasm scaffolds that had primary alignments of length 100 kb or greater to the corresponding Verkko Y sequences.

*HMM-Flagger.* HMM-Flagger detects structural errors in haplotype-resolved genome assemblies using coverage of mapped reads modelled using a Hidden Markov model augmented by a Gaussian autoregressive process. It classifies coverage anomalies as either erroneous blocks, false duplications, or collapsed blocks. To assess assembly quality, we mapped PacBio HiFi and ONT long reads, to the whole genome assemblies using Winnowmap. We also provided the censat and segdup annotation to the HMM-Flagger pipeline for detecting arrays with biased coverage and stratifying final results. We then subset the results only the Y-chromosome contigs. HMM-Flagger was run using the https://github.com/mobinasri/flagger/blob/v1.2.0/wdls/workflows/hmm_flagger_end_to_end_with_mapping.wdl wdl using the appropriate presets for each sequencing platform as mentioned here: https://github.com/mobinasri/flagger?tab=readme-ov-file.

*NucFlag.* To detect potential assembly errors in each chromosome Y sequence with NucFlag, we first aligned either PacBio HiFi or ONT reads to the relevant whole-genome assembly using pbmm2 (v1.13.1; https://github.com/PacificBiosciences/pbmm2; for PacBio HiFi reads) with the following parameters: --log-level DEBUG --preset SUBREAD --min-length 5000, or minimap2 (v2.30^60^; for ONT reads) with the following parameters: -a -x lr:hqae -y --eqx --cs -I 8g= -s 4000. PacBio HiFi and ONT alignments were filtered using SAMtools (v1.22^62^) and the filter flag (-F) 2308 to remove secondary, supplementary, and low-quality alignments. Next, we ran NucFlag (v0.3.4 for PacBio HiFi reads;^21^ or v1.0.0-a1 commit 49ea735c250cee1c23e25ebd2917bcec2282cfbc for ONT reads; Oshima et al., *in preparation*) with the following command: nucflag -i ${read_alignment.bam} -o ${output.bed} -t ${num_of_threads} -p ${num_of_processes} -c ${config.yaml}. NucFlag uses peak detection in the first- and second-highest read pileup nucleotide frequency to detect collapsed and misjoined sequences (v0.3.4) as well as insertions, deletions, softclipping, low-quality regions, mismatches, and heterozygous sites (v1.0.0) within each genome assembly. In addition, NucFlag v1.0 (https://github.com/logsdon-lab/NucFlag) annotates specific repeat-associated errors associated with long-read technologies, such as homopolymers and dinucleotide repeats, to allow downstream filtering. We called assembly errors with both PacBio HiFi and ONT reads using the following command: nucflag call -i ${read_alignments.bam} -f ${assembly.fasta} -p ${num_of_processes} -t ${num_of_threads} -x {ont_r9 or hifi} -o ${output.bed}.

*Illumina k-mers/QV*. Illumina NovaSeq reads for samples from the HPRC and HGSVC cohorts (part of the 1000 Genomes Project, “1000 Genomes 30x on GRCh38 collection,”^16^ were extracted using samtools fastq. Reads for HG002, HG003, and HG005 were sourced from GIAB, and CEPH pedigree reads were from Lin et al.^14^.

31-mers were generated with Meryl (v1.4.1) using: meryl count k=31 output $sample.k31.meryl $sample.R*.fq.gz. Base level quality value (QV) was evaluated for both the Y scaffold and unlocalized fasta sequences using Merqury^22^ (v1.4.1) via: $MERQURY/eval/qv.sh $sample.k31.meryl $sample.Y.fa.gz $sample. Using mapping results from the “Comparison to HPRC r2 version” section, comparable Y sequences from the HPRC r2 assemblies were extracted. The same command was run for HPRC r2 Y assemblies for comparison (**Suppl. Fig. 6; Suppl. Table 2**).

*Combined QC.* We derived a final label flagging potentially erroneous regions in the chromosome Y assemblies by combining the output of both Flagger and NucFlag at a resolution of 1 kb, which corresponds to Flagger’s default window size. A 1 kb window was labeled as “structural error” (label “ERRSTRUCT” in the annotation; see **Data availability**) if both tools flagged the region with both long-read technologies. This strategy was chosen to mitigate data biases such as well-known coverage dropouts in the HiFi sequencing data^63^. In addition to the structural errors, we also included “base-level errors” (label “ERRBASE” in the annotation; see **Data availability**) in the form of - typically short - sequence windows that exhibited a k-mer composition that was not supported by the sample sequence k-mers derived from highly accurate short reads (see section above “Illumina k-mers/QV”). Genome interval arithmetic such as intersections and merging of identically labeled consecutive regions was implemented using “bedtools” (v2.31)^59^ and “pyranges” (v0.1.4)^64^ (see **Code availability**).

### Y-chromosomal phylogeny construction and dating

The generation of the Y-chromosomal phylogeny for 142 samples followed the approach described earlier^10^. Briefly, variant calling was performed from high-coverage Illumina data^16,65^ across ∼10.4 Mb of Y-chromosomal sequence previously defined as accessible to short-read sequencing^36^. Variants were called with BCFtools^62,66^ (v1.16) using a minimum base quality of 20, mapping quality of 20 and haploid ploidy. Indels and SNVs located within 5 bp of an indel (SnpGap) were excluded. Genotypes were further filtered to retain sites with a minimum read depth of three and ≥85% of reads supporting the called allele. Sites with ≥5% missing data (that is, absent in more than seven of the 142 samples) were removed using VCFtools^67^ (v0.1.16). After filtering, 10,400,778 callable positions remained, including 25,426 polymorphic sites. As Illumina short-read data were not available from two samples, HG02486 and HG03471, data from their fathers (HG02484 and HG03469, respectively) was used for Y phylogeny construction and dating. To generate a dated phylogeny for samples with QC-passed centromeric regions, the filtered calls were extracted from above for 123 samples, including 23,185 polymorphic sites.

Y-chromosomal haplogroups were assigned as described previously^41^ according to the International Society of Genetic Genealogy (ISOGG v15.73) nomenclature. Because Y-haplogroup nomenclature is hierarchical, the number of represented haplogroups depends on the level of phylogenetic resolution. For summary analyses and visualization, samples were grouped into 17 broad major haplogroup clades; full haplogroup assignments and terminal markers are reported in **Suppl. Fig. 1** and **Suppl. Table 1**.

Divergence times for internal nodes were estimated using a coalescent framework implemented in BEAST (v1.10.4) both for the total dataset of 142 males and for 123 males with QC passed centromeric regions^68^. An initial maximum-likelihood tree was inferred with RAxML (v8.2.10)^69^ under a GTR plus gamma model and used as the starting tree. Markov chain Monte Carlo sampling was run for 150 million iterations with parameters logged every 1,000 steps, discarding the first 10% as burn-in. Analyses assumed a constant-size coalescent prior, a GTR substitution model with gamma-distributed rate heterogeneity, and a strict molecular clock. The clock rate was specified as a normal distribution centred on 0.76 × 10⁻⁹ substitutions per site per year with a 95% confidence interval of 0.67 × 10⁻⁹–0.86 × 10⁻⁹ ^70^. A maximum clade credibility tree was generated with TreeAnnotator (v1.10.4) and visualized in FigTree (v1.4.4).

### Variant calling

Two references were used for variant calling, T2T-CHM13v2.0 (T2T-CHM13)^63^, and GRCh38-NoALT (GRCh38)^71^ using PAV 3.0 (https://github.com/BeckLaboratory/pav3, *manuscript in preparation*).

Each variant was assigned to Y chromosome regions including pseudoautosomal (PAR), ampliconic (AMPL), X-degenerate (XDR), X-transposed (XTR), and highly repetitive regions including satellite (SAT), centromeric (CEN), *DYZ17*, *DYZ19*, heterochromatic (HET), and other (OTHER, unassigned and highly-repetitive). Because highly-repetitive tandem arrays can produce erroneous or different representations of variants, we excluded variants in SAT, CEN, *DYZ17*, *DYZ19*, OTHER, and HET loci. While PAR was not excluded from the callset, many of the analyses exclude this region and focus on the non-recombining regions of the Y chromosome.

A nonredundant callset was generated with Agglovar (https://github.com/BeckLaboratory/agglovar; *manuscript in preparation*) using the same algorithm and parameters as was previously used^13^, but with a much faster implementation. The same parameters and approach was used for intersecting variants between this callset and our previous callset on 43 Y chromosomes^10^.

#### Functional enrichment of SVs

We performed compartment-stratified permutation analyses of chromosome Y SVs (length >50 bp) separately for GRCh38 and T2T-CHM13v2. Analyses were restricted to PAR1 and the NRY; PAR2 was excluded because of the small number of SVs (50 in GRCh38 and 47 in T2T-CHM13v2). Observed overlaps were defined as the number of unique allele-level SV IDs intersecting five genomic elements—protein-coding genes, lncRNA genes, pseudogenes, protein-coding exons, and protein-coding introns—and were calculated separately for insertions and deletions. GRCh38 genomic elements were derived from GENCODE v47, whereas T2T-CHM13v2 elements were derived from the RefSeq Liftoff v5.2 annotation. Repeat and segmental-duplication annotations were assembly-specific: GRCh38 annotations were obtained from the UCSC hg38 rmsk and genomicSuperDups tables, whereas T2T-CHM13v2 annotations were obtained from the T2T-CHM13v2.0 RepeatMasker and segmental-duplication tracks. For each of 1,000 permutations, each SV was independently relocated within its observed compartment (PAR1 or NRY), while preserving its BED interval length. Candidate locations were required to match the corresponding observed interval endpoints in sequence class, broad RepeatMasker class, and whether each endpoint fell within a segmental duplication. Known GRCh38 assembly gaps were excluded from both the observed and permuted datasets. Enrichment was calculated as log_2_[(observed + 0.5)/(mean permuted + 0.5)]. Two-sided empirical *P*-values were calculated from the permutation tails using a plus-one correction and adjusted using the Benjamini–Hochberg method across the ten tests defined by two compartments and five genomic-elements, separately within each assembly and SV type.

### Y Chromosome Annotations

#### Draft annotations for assembly curation

*Y subregion draft annotations.* To ensure the integrity of the Y chromosome assemblies before and after curation, a draft sequence class annotation was created. This annotation marked sequence compositions, as detailed in^2^, in BED format to facilitate manual inspection on IGV. Sequences were annotated primarily for:

- Telomere
- N-gap
- HSat3 (from Y)
- HSat1A and HSat1B
- Centromere (from Y) and alpha-satellite
- Ampliconic sequences (P5-AZFb, P3-AZFc, blue, teal, red, green, yellow, gray as in the ampliconic palindrome designation)

Annotation targets were first identified from the T2T-CHM13v2 CenSat annotation^2^. To maximize sensitivity, the full refOriented.fasta was queried against these target sequences using minimap2. Coordinates were then inverted back to the assembly space using Rustybam, a process similar to that described in the “Identifying the main Y sequence” section. Throughout and after the manual curation process, sequences identified as Y components were annotated.

Resulting sequences annotated as HSat, centromere, or ampliconic were merged if the same annotation occurred within 500 base pairs. Any remaining regions without a specific annotation were labeled as “SEQ.” Color codes and directions (+ or -) were applied according to the specifications in src/annotate_regions.txt.

The annotation code is accessible on the T2T-chrY GitHub repository (https://github.com/arangrhie/T2T-chrY) under src/annotate.sh.

#### Final sequence class annotation

*Alignment-based transfer of sequence class labels*. The final annotation of the chromosome Y sequence class labels was realized in a similar process as described in^10^. Briefly, the labels were transferred from both reference annotations (GRCh38^31^, T2T-CHM13v2^2^ and **Suppl. Table 6**) via sequence alignment using minimap2 v2.28^60,72^. The labeled sequence windows in the chromosome Y assemblies were then extracted and aligned back to the respective reference and only those labels were retained that could reliably be re-aligned in this way. Next, the annotation was extended by adding previously curated sets of AZFc repeats (“color blocks”) using coordinates from Teitz et al.^31^ (**Suppl. Fig. 24)**; motif hits for *TSPY*, *DYZ19*, and Yq12 repeats identified with HMMER v3.4^73^ and post-processed with custom Python scripts (see **Code Availability**); telomere repeats determined with the seqtk v1.4 “telo” command; and the centromere annotation as a single “CEN” window (see Section “Centromere analysis” below). Error annotations for both structural and base-level errors were added as overlapping annotations. Regions of unresolved sequence (“N-gaps”) were included by subtracting the respective interval coordinates from the annotation and then adding the N-gap regions into the annotation file. All regions that were not labeled by a sequence class or N-gap in that way were labeled as “unassigned”. We note that this process of transferring the sequence class labels can lead to overlapping labels due to an imprecise alignment process that may be caused by factors such as biological meaningful sequence variation or structural errors remaining in the assemblies.

*Simplification of sequence class labels*. We reduced the sequence class label annotation to just the major annotation labels as defined in^1^, to which we refer to as “umbrella” labels (**Suppl. Table 6**). For example, the AMPL7 umbrella encompasses the AZFc region with all color block repeats and the P1-P3 palindromes. Overlapping umbrella intervals were split in half and evenly assigned to the neighbouring labels. Sequences of identically labeled regions were either joined or extended up to the next N-gap within the larger region. A further step of reducing the label complexity was applied for certain analyses by clustering labels by their prefix, e.g., all ampliconic regions were reduced to “AMPL” and so on (see, e.g., **Fig. 1c**). The post-processing steps were implemented in custom Python scripts (see **Code Availability**).

#### Segmental duplications

Repetitive elements in each genome assembly were first annotated and masked using a combination of three independent approaches. Tandem repeats were identified with TRF (v4.1.0)^74^ using the parameters trf [asm.fa] 2 7 7 80 10 50 2000 -l 30 -h -ngs. Interspersed repeats were detected with RepeatMasker (v4.1.5)^75^ using the NCBI engine and human repeat library (RepeatMasker -s -e ncbi -xsmall -species human [asm.fa]). Low-complexity regions were additionally annotated using WindowMasker (v2.2.22)^76^ in a two-stage procedure consisting of count generation (windowmasker -mk_counts -mem 16384 -smem 2048 -infmt fasta -sformat obinary -in [asm.fa] -out [asm.count]) followed by interval masking (windowmasker -infmt fasta -ustat [asm.count] -dust T -outfmt interval -in [asm.fa] -out [asm.interval]). The resulting repeat annotations from all three tools were merged, and the genome sequences were soft-masked accordingly. Segmental duplications were subsequently detected on the repeat-masked assemblies using SEDEF (v1.1)^77^ . Duplication pairs exceeding 1 kb in length, sharing greater than 90% sequence identity, and containing less than 70% satellite sequence were annotated as segmental duplication. The duplication sequences overlapping with Flagger and nucFlag regions were further filtered out. In addition, duplications orthogonally validated by FastCN^78^ computed on the respective *de novo* assemblies were retained.

#### Gene annotation

*Graph-based gene annotations.* minigraph-Cactus^79^ was used to build a Y-chr pangenome using this -cactus-pangenome ./js hprc-hgsvc-ceph-chrY.seqfile --outDir hprc-hgsvc-ceph-chrY --outName hprc-hgsvc-ceph-chrY --logFile hprc-hgsvc-ceph-chrY.log --reference HG002_chrY --lastTrain --snarlStats --filter 46 --giraffe --viz --haplo --vcf --vcfwave --chrom-vg --chrom-og --gfa clip full --gbz clip full filter --mgCores 32 --mapCores 8 --consCores 32 --indexCores 64 --doubleMem true --batchSystem slurm --batchLogsDir batch-logs-hprc-hgsvc-ceph-chrY --slurmTime 168:0:0

CAT2 (improving upon CAT^80^) was then used to annotate genes on all of the chrY assembly using this pangenome alignment with HG002_chrY as the reference gene set. CAT2 uses multiple methods to transfer genes from the reference to each of the targets- transMap module lifts genes over pairwise chains derived from the pangenome and pairwise chains derived from minimap2 alignments, Liftoff^81^ is integrated to annotate any additional genes missed by the transMap methods and AUGUSTUS gene prediction fixes gene and CDS boundaries.

*LIFTOFF Annotation.* Liftoff^81^ (v1.6.3) was run using minimap2^60^ (v2.24) as the alignment engine, with 8 parallel threads. A minimum sequence identity score threshold of 0.80 was applied (-sc 0.8) to ensure only high-confidence annotation mappings were retained. The -copies flag was enabled to allow detection and mapping of extra gene copies present in the target assembly beyond those in the reference, and the -cds flag was used to perform additional polishing to ensure valid open reading frames in the transferred coding sequences. Unmapped features that failed to meet the similarity threshold were written to a separate unmapped output file for downstream quality assessment. All analyses were performed on a high-performance computing cluster using SLURM. Following the liftover, transferred annotations were further filtered to retain only high-quality mappings. Specifically, features with a sequence identity and coverage of less than 0.95 were excluded, ensuring that only annotations with at least 95% sequence identity and 95% coverage relative to the reference were carried forward for downstream analyses.

*Ampliconic gene annotation.* A custom RepeatMasker (v4.1.2)^82^ library containing the canonical exonic sequences from a representative from each of the nine ampliconic genes (*DAZ* (ENST00000405239.6), *PRY* (ENST00000303804.5), *BPY2* (ENST00000331070.8), *CDY* (ENST00000306609.5), *TSPY* (ENST00000451548.6), *RBMY* (ENST00000383020.7), *HSFY* (ENST00000307393.3), *VCY* (ENST00000250825.5), and *XKRY* (ENST00000510392.1)) was used to annotate ampliconic gene exon locations in assemblies. Canonical ampliconic gene exon sequences were obtained from Ensembl (release 113)^83^. Truncated (containing at least half of all canonical exons) and full exon ampliconic gene copies were identified by detecting exons present in their canonical order and in the same orientation.

#### Polymorphic MEI annotations

Mobile element insertions (MEIs) specifically non-long terminal repeats (non-LTRs: SINE/Alu elements, LINE/L1, and Retroposon/SVA) were first annotated within assemblies (140/142 open access assemblies only) with RepeatMasker (v4.1.7)^82^ and the Dfam (v3.8) library^84^. Candidate MEIs ±30 bp of flanking sequence were directly extracted from assemblies using Pysam (v.0.23.3)^58,85^. These candidate sequences were re-Repeatmasked (with the same library) and then provided to L1ME-AID (v1.4.1-beta)^13^. Putative MEIs were grouped by family (e.g., AluYb8, AluYa5, etc.), and their sequences aligned with MUSCLE (v3.8.31)^86^. Family groups were consolidated into unique element insertions based on manual curation of the alignments. Finally, unique candidate MEIs ±500 bp flanking sequence were retrieved from assemblies using Pysam (v.0.23.3)^58,85^ and then aligned against all Y assemblies, using Minimap2 (v2.30-r1287). Minimap2 high-confidence matches to candidate MEI sequences (sequence identity ≥99%, and coverage ≥99%) were used to remove duplicated elements (not unique insertions) and confirm/correct sample genotyping (**Suppl. Table 15**). In parallel, we generated a Yq12-specific MEI callset in which duplicated elements were retained, allowing us to evaluate both unique and repeated MEI copies within this highly repetitive region (**Suppl. Table 26**). Polymorphic element insertion ages were estimated based on the most recent common ancestor (MRCA) of samples sharing the insertion and the SNV-based Y male phylogeny times (**Suppl. Fig. 1**).

#### Ampliconic regions and palindromes

Palindrome identification was performed *de novo* for each repeat-masked Y chromosome assembly. Individual contigs were extracted and self-aligned using LASTZ^87^. Self-alignment output was processed with PALINDROVER^88^ with the following parameters: minimum arm length of 8 kb, a minimum inter-arm sequence identity of 98%, and a maximum spacer length of 500 kb. Repetitive elements annotated by RepeatMasker (classes: Low_complexity, Simple_repeat, and Satellite) were used as a blacklist; palindromes overlapping blacklisted regions by ≥80% were excluded. Arm coordinates from all samples were collected and formatted as BED files.

To cluster palindromes across samples, left arm sequences from palindromes were merged into a single combined FASTA, and each sample’s left arms were aligned to this combined reference using LASTZ^87^ (minimum identity 90%, minimum query coverage 70%, HSP threshold 5,000). Alignment results were used to construct an undirected graph in which nodes represent individual palindrome arm sequences and edges connect pairs of arms with bidirectional alignment coverage exceeding 80%. Connected components of this graph represent sets of homologous palindromes across samples. Components were then annotated by mapping to known Y chromosome palindrome designations and re-named according to established “P” nomenclature^1^ whenever possible. One additional palindrome, initially designated Q6 following the PALINDROVER convention (based on its T2T-CHM13v2Y/HG002 homolog), was renamed P9 here for consistency with the existing Y-chromosome palindrome nomenclature. A short palindrome containing the first copy of the RNA recognition motif of *DAZ* was reported separately. “P1” designates the standard palindrome as reported in ^1^ and “P2_inversion” designates the recurring *gr/gr* inversion where one arm of P1 is inverted and part of the gene content creates a larger palindromic formation at the P2 palindrome (as observed in T2T-CHM13v2Y/HG002 - ^2^). Table of palindrome occurrences visualised as a heatmap (**Suppl. Fig. 76**).

For each of the ten reported palindrome loci (P1-P8, *DAZ*, and P9), arm length, spacer length, inter-arm sequence identity, and repeat content were extracted from the corresponding self-alignment files and summarised as boxplots with per-locus median annotations (**Suppl. Fig. 77, Suppl. Tables 31-32**).

#### Inversion recurrence analysis

*Pangenome graph construction.* A pangenome graph of 142 Y-chromosome assemblies (105 HPRC, 29 HGSVC, and 6 CEPH samples, plus T2T-CHM13v2Y and GRCh38) was constructed using the PGGB pipeline v0.7.4-26-g498c5d7^89^ with wfmash v0.23.0-41-gb5f0ff1c, seqwish v0.7.11-4-g9a7131d^90^, smoothxg v0.8.2-2-g2a6f17f, odgi v0.9.2-0-gbe6a0202^91^, and GFAffix v0.2.1 (https://github.com/codialab/GFAffix). Assembly sequences were renamed to PanSN format (sample#haplotype#chrY) before graph construction. All-versus-all alignments were computed with wfmash at a minimum percent identity of 98% (-p 98) and the default segment length of 1,000 bp (-s 1000). Graph induction was performed with seqwish using a minimum exact match length filter of 311 bp (-k 311), chosen to reduce spurious graph complexity arising from the highly repetitive ampliconic and palindromic sequences on chrY. Graph normalization and diagnostic output were run with PGGB defaults.

We analyzed the recurrence status of 22 potential homology-mediated inversion loci in the region spanning inverted repeats IR1 to IR5 and palindromes P1 to P8. For this purpose, we used PIVOT, a tool for detecting recurrent inversions within a graph-based pangenomic framework^92^ (Chapter 5). It builds upon the theoretical framework of the tiSNPs-based approach we proposed in Porubsky et al.^24^, while extending it by analyzing all within-inversion variant types–not just SNPs–to identify variants inconsistent with a single-inversion origin. Additionally, it operates directly on pangenome graphs, thereby mitigating reference bias. Locating an inverted region across all haplotypes represented in a pangenome is nontrivial due to the absence of haplotype-specific breakpoints. PIVOT addresses this by using the inverted repeats flanking and potentially mediating the inversion. In the graphical framework, the nodes representing these flanking repeats serve as entry and exit paths, allowing haplotype paths to traverse the inversion locus in both orientations. These nodes act as “anchors” to locate the region of interest across the entire haplotype panel. We used a chrY pangenome graph generated by PGGB for this analysis (see details in section ‘Pangenome graph construction’). The coordinates of the regions analyzed for recurrence, provided as input to PIVOT, were derived from repeat and palindrome annotations for T2T-CHM13v2. Since the input coordinates already covered the flanking repeats, PIVOT was run with the parameters -flank 0 and -limit 0 . In addition, we used -safe_len 15 and -safe_len_limit 5.

#### *AZFc* region analysis

AZFc Colorblock Region Annotation and clustering. Each assembly was scanned for 100 base pair (bp) kmers derived from the GRCh38 AZFc colorblock coordinates from Teitz et al.^31^. Regions of high kmer density (defined as ≥8,000 kmer matches within a 10,000 bp sliding window, step size = 1,000 bp) were identified across each Y chromosome assembly. The start and end of each color block (blue, teal, green, red, yellow, and gray) and their spacer regions were defined based on the boundaries of the kmer signal.

*Clustering of various structures of the AZFc region.* We set to cluster 100 structural haplotypes at chromosome Y *AZFc* region. There were 10 repeat units (Color blocks: Blue, Teal, Red, Green, Yellow, Grey, Green-IR1, Blue-IR1, Blue-Plus, and Spacer) defined in each haplotype. Then in every haplotype we assigned each repeat unit a unique numeric identifier. This identifier was either a positive or a negative number for direct and reverse oriented repeat units, respectively. Then for each possible pair of haplotypes we calculated the distance between these numeric encodings of each haplotype structure as described before^93^. All pairwise distances were organized into the distance matrix, which was then used to construct the UPGMA tree using the function ‘upgma’ from the R package phangorn (v2.11.1)^94^. Visually we defined 20 nonredundant haplotype groups (which included spacer orientations) that were separated by the ‘cuttree’ function (set parameter: k=20) from the base R package.

*AZFc Colorblock Region Recurrent Inversions.* Assemblies carrying recurrent AZFc *b2/b3* inversions (HG01109, HG01074, NA20870, NA21093, HG02074) and gr/gr inversions (HG002, HG01255, HG01258, HG01952, HG02027, HG00706, HG02083, HG02492, HG02514, HG03471, HG03688, HG04199, NA18620, HG00642) were each compared to the phylogenetically closest sample with a GRCh38-like *AZFc* architecture to localize inversion breakpoint interval (**Suppl. Table 16**). Colorblock sequences were extracted from each assembly using Pysam (v.0.23.3)^58,85^, inspected in AliView (v1.3.0)^95^, and manually aligned. Alignments were parsed with custom Python scripts (**see Code Availability**), and only SNVs were called; regions with homopolymer variation, low-complexity sequence, or indels were excluded. Breakpoint intervals were defined at positions where the shared SNV allele pattern shifted from being located on the same (both on either 5’ or 3’) colorblock copy to different (one on 5’ and the other on 3’ or vice versa - a sign of inversion) copies between the two samples or where the change in SNV density was greatest (HG02514, HG02492, HG02083, HG02027, HG002) (**Suppl. Figs. 28, 29, 32**). Single SNVs consistent with the inverted pattern but flanked by SNVs showing non-inverted patterns were ignored as those could result from double crossover events between the colorblocks. Inversion intervals were considered distinct based on their relative distances from the start and/or end of the colorblock containing the breakpoint.

*AZFc inversion window percent identity.* Sequences corresponding to the AZFc proximal and distal inversion breakpoint window for each sample were extracted from the genome assembly using BEDTools (v2.31.1)^59^ (*bedtools getfasta -fi assembly.fasta -bed interval_coordinates*). Pairwise sequence alignments between the proximal sequence and the reverse-complemented distal sequence were performed using Biopython (v1.85)^96^ with scoring parameters derived from EMBOSS Water^97^ (*Bio.Align.PairwiseAligner(mode="global", match_score=5, mismatch_score=-4, open_gap_score=-10, extend_gap_score=-0.5)*)). Percent identity for the highest scoring alignment was reported (**Suppl. Table 16**).

*AZFc inversion and deletion window repeat content*. The repeat content within the breakpoint windows was extracted from the RepeatMasker annotation track (generated in this study) using BEDTools (v2.31.1)^59^ (*bedtools window -w 0 -a interval_coordinates -b repeatmasker_annotation.bed*) (**Suppl. Figs. 30, 31, 33 and 34**).

*AZFc architectural constraint analysis.* To test whether observed AZFc architectures occupy a more restricted structural state space than expected from rearrangement opportunity alone, each architecture was represented as an ordered series of ampliconic repeat-family blocks and their orientations. Of the 100 AZFc QC-passed chromosomes, two architectures whose defining inversions extended outside the modeled AZFc block system (HG02666 and NA20905) were excluded, leaving 98 chromosomes representing 10 internally modeled architectures. The Y-chromosome phylogeny (**see ‘Y-chromosomal phylogeny construction and dating’**) was pruned to these 98 samples while retaining branch lengths. Potential NAHR substrates were defined as pairs of blocks belonging to the same repeat family: oppositely oriented pairs could generate inversions, whereas directly oriented pairs could generate deletions or reciprocal tandem duplications. Structurally silent outcomes were excluded. At each simulated event, an eligible homologous pair was selected uniformly from the current architecture, followed by one of its permitted state-changing products. Substrate availability was recalculated after each event, allowing architectures to evolve dynamically along the empirical phylogeny. No explicit cap was imposed on simulated architecture size. Parsimony reconstruction identified two equally parsimonious ancestral states, corresponding to the HG01890-like and GRCh38-like internal *AZFc* configurations, each containing 17 modeled blocks. Minimum-event reconstruction indicated that 30–32 state-changing internal NAHR events were required across the phylogeny; simulations were therefore conditioned on exactly 30 events for the primary analysis and repeated at 31 and 32 events as sensitivity analyses. Architectural restriction was quantified as the maximum absolute terminal deviation from the ancestral block count, D_max_ = max_i_|B_i_ - B_root_|, where B_i_ is the block count of terminal architecture i and B_root_ is the block count of the inferred ancestral architecture. Observed architectures contained 11–23 blocks, giving D_max_= 6. For each possible root and event count, 1,000 accepted simulations were generated, and one-sided Monte Carlo *P* values were calculated using the +1 correction, P = (X+1)/(N+1), where x is the number of simulated replicates with D_max_ ≤ 6 and N is the total number of accepted replicates. The null represents a permissive NAHR opportunity model and does not incorporate pair-specific differences in sequence identity.

*Potential mechanisms underlying AZFc architectural restriction.* First, to test whether older rearrangements were preferentially absent, simulated event histories were reduced to terminal architectures and architecture changes were reconstructed using the same parsimony procedure applied to the observed data; the mean phylogenetic age of reconstructed changes was then compared with ascertainment-matched simulations conditioned on 30–32 state-changing events. Second, to test whether NAHR preferentially involved physically closer homologous blocks, symbolic *AZFc* architectures were mapped to assembly coordinates and analyses were restricted to reconstructed transitions explainable by a single internal NAHR event. For each transition, all compatible substrate pairs producing the observed destination were compared with available state-changing opportunities in the source architecture using center-to-center distance and one-sided permutation tests against opportunity sets matched by source architecture and event type, with or without additional matching by repeat family. Third, to test whether NAHR preferentially involved longer homologous blocks, the same one-step transitions and matched opportunity sets were analyzed using the length of the shorter homologous block as the primary measure, with one-sided permutation tests for preferential use of longer substrates.

*DAZ gene copy classification.* To distinguish individual *DAZ* gene copies (e.g., *DAZ1* from *DAZ2*), 15 previously published paralogous sequence variants (PSVs) were used (**Suppl. Table 20**)^41,98,99^. Each PSV plus 50 bp of flanking sequence (101-bp PSV-centered windows) were extracted from the GRCh38 Y reference sequence using PySam (v0.23.3)^58,85^. *DAZ* gene sequences (plus 50 bp of flanking sequence) were retrieved from each assembly using PySam (v0.23.3)^58,85^. The 101-bp PSV-centered windows were then aligned to the gene sequences using BLAT (v35)^100^. Only full-length alignments with 100% sequence identity were retained. BLAT output was parsed with custom Python scripts (**see Code Availability**), and *DAZ* copies were assigned as *DAZ1*, *DAZ2*, *DAZ3*, or *DAZ4* based on the highest total number of PSVs detected at unique positions within each *DAZ* sequence (multiple PSVs mapping to the same position were counted only once). In instances where the highest total count was the same, both gene labels were used (e.g., *DAZ3/DAZ4*; **Suppl. Fig. 40-42**).

*DAZ gene exon composition. DAZ* exons were extracted from assemblies using PySam (v0.23.3)^58,85^ and aligned to reference exon sequences with previously assigned *DAZ* exon identifiers using MUSCLE (v3.8.31)^86,101^. *DAZ1* exons from the canonical Ensembl sequence (**Methods: Ampliconic gene annotation**) corresponded as follows (’EXON_17’:’B’, ’EXON_18’:’C’, ’EXON_19’:’D’, ’EXON_20-22’:’E’, ’EXON_21’:’F’, ’EXON_23’:’X’, ’EXON_24’:’Y’, ’EXON_25’:’Z’). *DAZ1*‘EXON_17’ copies that diverged from the canonical Ensembl exon sequence were assigned the *DAZ2* specific exon “A” annotation.

#### *de novo* mutation analysis

*Detection and validation of de novo mutations.* Eight samples from the CEPH Platinum pedigree^14,15^ were used to identify *de novo* mutations (DNMs) in two independent Y-chromosome pedigrees belonging to distinct lineages within haplogroup R1b (**Fig. 1b; Suppl. Fig. 71; Suppl. Table 1**). In the first pedigree, individual 200080 (R1b-Z326) was analyzed together with his two sons, 200084 and 200085. In the second, NA12877 (R1b-Z302) was analyzed together with his four sons, NA12882, NA12883, NA12884 and NA12886.

For each family, the paternal Y-chromosome assembly was used as the reference to identify *de novo* variants in the sons across the non-recombining regions of the Y chromosome, excluding the pseudoautosomal regions. The paternal NRY spanned approximately 51.4 Mb in individual 200080 and 48.8 Mb in NA12877. Variants were called from each son’s Y assembly relative to the corresponding paternal assembly using Dipcall (v0.3)^102^ with default parameters recommended for male samples.

Only variants on the main chrY contig were retained. Variants overlapping regions flagged by HMM-Flagger or NucFlag, based on either HiFi or ONT support in either the reference or query assembly, were removed. Indels in homopolymer tracts or annotated by RepeatMasker as Simple_repeat or Low_complexity were also excluded.

Read-based validation of DNMs was performed using both PacBio HiFi and ONT-UL reads. Support for each candidate site was evaluated in four alignment contexts: paternal (i.e., reference) reads mapped to the paternal assembly, son’s reads mapped to the son’s assembly, son’s reads mapped to the paternal assembly, and paternal reads mapped to the son’s assembly using minimap2 (v2.28)^60^ with default parameters. Only primary alignments were retained (samtools view -F 2308), and per-base read counts were extracted using bam-readcount (v1.0.1) with minimum mapping and base qualities both set to 20. In addition, all alignments covering the DNMs were visually checked in IGV (v2.16.0)^103^.

For validation of *de novo* SNV, no paternal reads were allowed to support the son’s *de novo* allele and no son’s reads were allowed to support the paternal allele in the HiFi read alignments. For ONT-UL data, a small amount of discordant support was tolerated because of the higher error rate. Among the 43 *de novo* SNVs, three sites showed low-level support for the son’s allele in the paternal ONT-UL reads (≤1.5% of paternal reads), while a different set of three sites showed low-level support for the paternal allele in the son’s ONT-UL reads (≤5% of son’s reads). At all remaining SNV sites, the ONT-UL reads were fully concordant with the expected paternal and son genotypes. As an additional control, sibling read data mapped to the paternal/reference assembly were examined and in all cases supported the paternal allele. All read alignments were visually checked for support for the *de novo* indels and SVs.

Additional validation was performed for DNMs identified in NA12886 using pedigree data from Porubsky et al. 2025^15^. All candidate variants were confirmed to be in the reference state in the grandfather’s Y assembly (NA12889, father of NA12877), and transmission was assessed in the three sons of NA12886 (200101, 200102 and 200105) using available PacBio HiFi and ONT-UL data. HiFi data were available for all three sons, whereas ONT-UL data were available for individuals 200101 and 200102. Reads were mapped with minimap2 to the NA12877 and NA12886 Y assemblies, and support was evaluated as above. All DNMs identified in NA12886 were confirmed in his three sons.

Classification of 40 Yq12 DNMs. Single nucleotide variants identified in six father–son pairs were restricted to the DYZ1 and DYZ2 repeat arrays within Yq12 (see Methods ‘Detection and validation of de novo mutations’). Any intervals flagged by HMM-Flagger and/or NucFlag were excluded. For each SNV, we tested whether the son’s alternate allele could be explained by intra-array gene conversion (GC) by searching the father’s Yq12 for homologous donor copies using a seed-and-extend strategy. Using Pysam (v.0.23.3)^58,85^ and custom python code (see Code availability), exact 4-bp seeds immediately flanking, but not including, the focal site was required, and matching sequence was extended outward until the first mismatch to quantify total flanking homology. Hits were discarded if they overlapped masked error regions, occurred within 200 bp of the focal site, or had <200 bp of total flanking homology. For each retained donor, the allele at the homologous site was recorded, and redundant donors were collapsed by unique combinations of flanking homology length and carried allele before statistical testing. Using Scipy (v1.16.1; scipy.stats.binom.sf)^104^ one-sided binomial test was then used to assess whether the observed number of concordant donor copies exceeded expectation under a trinucleotide-context-dependent mutational null. The null model was estimated in two passes: first, all events were screened using a simple uniform alternate-allele model to identify low-support events provisionally classified as de novo mutations (DNMs), excluding clustered SNVs. Second, those provisional DNMs were used to fit an empirical trinucleotide-context mutation model that was applied in the final per-site tests. SNVs separated by ≤300 bp were grouped into clusters and additionally evaluated by a joint donor search that required preservation of inter-SNV spacing across the cluster. Clusters with at least one spanning donor fully matching the son’s allelic state at every site were classified as concordant clusters, whereas clusters with spanning donors but no fully concordant donor were classified as partial clusters. Benjamini–Hochberg false-discovery-rate correction (α = 0.05)^105^ was then applied only to singleton ‘GC_candidate’ sites; fully concordant cluster representatives were not included in this FDR step and were instead directly classified as ‘Likely_GC.’ Singleton ‘GC_candidates’ with a significant FDR q-value were re-classified as ‘Likely_GC’ events. Final calls were therefore assigned as ‘DNM’, ‘GC_candidate’, or ‘Likely_GC’.

#### *TSPY* and *RBMY* arrays

For analyzing the *TSPY* and *RBMY* gene arrays, exon sequences for each gene copy were extracted from assemblies using Pysam (v.0.23.3)^58,85^ and concatenated (in exon order) to generate a per-copy coding sequence. Concatenated exon sequences were translated in AliView (v1.3.0) ^95^, and amino-acid sequences were aligned with MUSCLE (v3.8.31)^86^. Multiple-sequence alignments were parsed with custom Python scripts (see **Code Availability**), and pairwise Hamming distances among aligned amino-acid sequences were computed with SciPy (v1.16.1)^104^. Gene copies with identical amino-acid sequences (100% identity) were collapsed into a single group and represented as one node in an undirected NetworkX (v3.5) graph^106^. Edges were placed between nodes that contained the lowest hamming distance from one another compared to all other nodes (ties were allowed). Iterative graph simplification was performed in NetworkX by repeatedly merging removable nodes into adjacent “representative” nodes. In each round, leaf nodes (node degree = 1) whose neighbor had degree > 1 were collapsed into their neighbor, and any node with a group size of 1 gene was collapsed into its largest-size neighbor. The nine highest-degree nodes were designated hubs and assigned unique colors (blue, green, etc.); colors were then propagated through the network to assign a color to all remaining nodes based on proximity to the hub nodes. For *TSPY* and *RBMY* separately, the sequence upstream of exon 1 (2 kb for *TSPY* and 1 kb for *RBMY*, reflecting differences in array length) was extracted using PySam (v0.23.3)^58,85^. Redundant upstream sequences showing 100% sequence identity were collapsed to unique representatives before motif analysis. These unique upstream sequences were then screened for exact identity matches to transcription factor binding motifs using the JASPAR 2024 CORE vertebrate database^107^. A representative-by-motif count matrix was constructed from these motif calls and analyzed by principal component analysis (PCA) using sklearn.decomposition^108^. K-means clustering (sklearn.cluster)^108^ was evaluated across multiple values of *k*, and silhouette analysis was used to assess cluster performance, with *k* = 2 selected for downstream analyses^109^. To identify motifs most strongly associated with each cluster, motif enrichment was calculated relative to the global mean across all promoters, and the top enriched motifs were ranked within each cluster. To aid biological interpretation of the motif-defined clusters, the ten motifs with the greatest enrichment relative to the global mean motif count across upstream sequences were selected for each cluster. For composite motifs containing two transcription factors, the motif name was separated into its component gene symbols, and Gene Ontology Biological Process (GO-BP) annotations for the corresponding genes were retrieved from MyGene.info using the mygene Python package (v3.2.2)^110–112^. GO-BP terms were summarized descriptively for each cluster; no statistical GO over-representation test was performed.

### Yq12 array architecture and evolutionary analyses

Analyses of Yq12 architecture were restricted to the 84 assemblies that passed Yq12 quality control. Total *DYZ* array number was calculated as the sum of *DYZ1* and *DYZ2* arrays, and total satellite-array length as the summed physical length of all *DYZ1* and *DYZ2* arrays. Mean array length was calculated as total satellite-array length divided by total array number. Because total array length is exactly the product of array number and mean array length, we decomposed the variance of log-transformed total satellite-array length as Var[log(*N*)] + Var[log(*L̄*)] + 2Cov[log(*N*), log(*L̄*)], where *N* is array number and *L̄* is mean array length. Each component was expressed as a fraction of the total log-length variance. Uncertainty was estimated by resampling chromosomes with replacement 10,000 times and recalculating the decomposition; 95% bootstrap intervals were taken from the 2.5th and 97.5th percentiles.

Phylogenetic analyses used the Y-chromosome phylogeny (see **‘Y-chromosomal phylogeny construction and dating’**) and the 84 QC-passing chromosomes. Phylogenetic signal in log2-transformed total array number and mean physical array length was quantified using Pagel’s λ^113^, with λ constrained between 0 and 1. The fitted-λ model was compared by AIC^114^ with fixed λ = 1, representing Brownian phylogenetic covariance, and λ = 0, representing no shared phylogenetic covariance.

Evolutionary coupling between *DYZ1* and *DYZ2* was assessed using a bivariate Brownian-motion model fitted to log2-transformed *DYZ1* and *DYZ2* repeat-unit counts^115^. The fitted evolutionary covariance matrix was used to calculate the evolutionary correlation (ρ), variance along a shared-size axis and variance along a *DYZ1:DYZ2* imbalance axis. Coupling was tested by likelihood-ratio comparison with a model in which evolutionary covariance was constrained to zero, and rate asymmetry was tested against a model with equal marginal evolutionary variances. Simulation-based 95% intervals for evolutionary correlation, the shared-to-imbalance variance ratio and the *DYZ1/DYZ2* evolutionary-variance ratio were obtained from 10,000 parametric simulations under the fitted Brownian model.

### Y callable regions

*Callable regions and prior masking*. The sequence annotation for T2T-CHM13v2 was restricted to chromosome Y^2^. Global masks BED files define intervals considered callable using different methodology: pangenome structural variation for pm151b.easy (strict) regions^38^, reference based illumina callability mask^37^, short read and depth statistics^36^. Using Bedtools intersect (v2.31.1), each mask was intersected with the Y sequence annotation to define a callable fraction across sequence classes (**Fig. 5h**). Callable fraction is defined as the fraction of total base pairs within a class that overlap the mask (base pairs in the mask / total base pairs in sequence class). We further partitioned the class specific callable intervals using bedtools multiinter to report shared base pairs and unique intervals to each combination of the three masks. XTR1 and XTR2 were combined into one XTR category, and SAT and *DYZ17* were combined into SAT. Summed overlap counts were visualized using Euler diagrams to show qualitative relationships between the global masks within each sequence class; specific basepair overlap counts between masks can be found in **Suppl. Table 37**.

*Short read variant calling.* Short-read whole-genome data were aligned with BWA-MEM (v0.7.19-r1273)^116^, and the resulting alignments were sorted using Samtools (v1.22.1)^58^. Per-sample Y chromosome variants were called from the processed alignment files using GATK (v4.6.2.0)^117^ HaplotypeCaller in gVCF mode (–ERC GVCF) against the CHM13v2.0 human reference genome. To reflect sex-chromosome ploidy, HaplotypeCaller was run twice with region-restricted intervals: (i) a *diploid* call over both pseudoautosomal regions (PAR1: chrY:1-2458320 and PAR2: chrY:62122809-62460029) and (ii) a *haploid* call over non-pseudoautosomal X/Y regions (chrY:2458321-62122808). The resulting diploid and haploid gVCFs were then merged into a single per-sample gVCF using GATK CombineGVCFs.

For each partition (diploid and haploid calls), inputs were imported into separate GenomicsDB workspaces with GenomicsDBImport. Joint genotyping was then performed with GenotypeGVCFs on each GenomicsDB to produce diploid/haploid-specific cohort VCFs. Finally, the resulting diploid and haploid-specific VCFs were concatenated into a single cohort VCF using GatherVcfs.

*Per sample level consensus and QV within callable regions*. To estimate the accuracy of variant calls within different annotation classes and different callability masks, we used the variant genotypes from the short read variant calls described above to infer a Y chromosome “pseudo reference” for each sample, and then we measured the agreement between this constructed pseudo reference and the sample’s Verkko2 long read assembly.

Briefly, for each sample, bcftools was first used to create a sample-specific VCF file of variant calls and a sample-specific BED file of high confidence regions (both against the T2T-CHM13v2 reference) from the cohort VCF file produced in the short read variant calling step. To limit an analysis to regions of T2T-CHM13v2 within one of the three callability masks, the sample-specific VCF and high confidence BED file were intersected with the mask’s BED file using bedtools. The resulting set of called Y chromosome variants was then applied to the reference using “bcftools consensus” to generate a sample-specific pseudo reference sequence in FASTA format, as well as a chain file for lifting reference coordinates from T2T-CHM13v2 to the new sample-specific pseudo reference. Finally, the high confidence regions within the callable mask were lifted to the new sample-specific Y coordinates and used to mask any regions of the pseudo reference FASTA that were outside the callability mask or were not deemed high confidence in the original variant calling (see script “sh.make_sample_gvcf_genome”, https://github.com/nhansen/chrY_callable_analysis).

Then, using minimap2^60^ (v2.28) and GQC’s “compare” command, each masked pseudo reference FASTA file was aligned to its corresponding Verkko Y assembly for the same individual, first masking sequences in the Verkko assembly that were annotated as base errors (“ERRBASE”). To assess base level accuracy, differences between the pseudo references and the assemblies were counted as errors within each sequence class as defined by annotation along the T2T-CHM13v2 reference (see script “sh.run_gqc_compare_norandoms_minalign1000”, https://github.com/nhansen/chrY_callable_analysis). Error rates were combined within these sequence classes across individuals. For each sample and sequence class, the error rate was calculated as the number of errors in class / length of class and then converted to phred quality values using QV (−10*log_10_(error rate)). QV values were visualized across all individuals (**Suppl. Fig. 98;** https://github.com/sifordrebecca/functional_chrY) to view sample specific variation by sequence class and QV values were also averaged across samples for each class by totaling errors and class lengths across samples, then converting the overall error rate to phred quality values as above. Similarly, we produced QV estimates within the intersection of each sequence class and each of the global masks to compare sequence accuracy within the callable regions of each mask to see which mask called the highest QV sequence (**Fig. 5i)**. Regions with zero discrepancies were reported as 60+ to indicate no observed errors. QV was not calculated for categories with fewer than 1000 aligned bases and are represented as white boxes.

### Centromere analyses

*Chromosome Y centromere mapping and annotation.* To identify completely and accurately assembled chromosome Y centromeric regions in whole-genome assemblies, we applied a Centromere Mapping and Annotation Pipeline (CenMAP; https://github.com/logsdon-lab/CenMAP; v1.1.0, commit b1aebc0308b4e05fa42466793f15e8ceef08fdb5) that we recently developed^21^. Briefly, CenMAP aligns each whole-genome assembly to the centromeric regions of the T2T-CHM13v2 reference genome^63^ (v2.0) using minimap2^60^ (v2.28) with the following parameters: -x asm20 --secondary=no -s 25000 -K 15G. The resulting BAM files are then converted into BED files using rustybam (v0.1.34)^118^. Then, contigs containing centromeric regions are reoriented from p to q arm relative to the T2T-CHM13v2 reference genome (v2.0). Contigs are renamed to indicate the chromosome they most accurately align to based on the percent of sequence aligned to the CHM13 reference genome (v2.0), with a minimum of 5% required. To detect centromeric contigs *de novo*, CenMAP (v1.0 and greater) runs three tools: K-Mer Counting (KMC)^119^, Satellite Repeat Finder (SRF)^120^, and Tandem Repeat Finder (TRF)^74^. First, CenMAP identifies repetitive sequences in a whole-genome assembly by identifying *k*-mers (*k* = 171 bp; the average length of an α-satellite repeat) that occur more than 10 times using KMC 3 (v3.2.4)^119^. Then, satellite motifs are constructed using SRF (commit e54ca8c8eccf6b1f19428b0f862f2c90575290a0)^120^, and tandem repeats are annotated in each motif with a fork of TRF (trf-mod; commit 3e891db310124f7e5f7a630a1c006650be9d1f3a; github.com/lh3/TRF-mod). Using a custom script we developed to work with both outputs, srf-n-trf (v0.1.1; https://github.com/koisland/srf-n-trf), and the subcommand *motifs*, we remove any SRF motif without a TRF motif and with a bp length of 170 n, where n <= 5 for α-satellite monomers or <=42 for HSat1A monomers^121^ within 2% difference in length with the following command: srf-n-trf motifs -f ${srf_motifs} -m ${trf_monomers} -s 170 340 680 850 1020 42 -d 0.02. With a candidate set of motifs, CenMAP maps each assembly to the motifs using minimap2 (v2.29)^60^ and recommended parameters by SRF with the following command: minimap2 -c --eqx -N 1000000 -f 1000 -r 100,100 <(srfutils.js enlong ${srf_motifs}) ${assembly.fasta}. The output PAF is then filtered to find monomers from aligned regions or CIGAR elements that contain an α-satellite or HSat1A monomer with the command: srf-n-trf monomers -p ${paf} -m ${trf_monomers} -s 170 340 680 850 1020 42 -d 0.02 --max-seq-div 0.3. All α-satellite and HSat1A regions are iteratively merged with srf-n-trf *regions* if they are within 100 kb of each other and contain at least one α-satellite monomer using the following command: srf-n-trf regions -b ${monomers.bed} -m 30000 -d 100000 -s 170 340 680 850 1020 --diff 0.02. This yields the coordinates of the centromeric α-satellite HOR array(s) greater than 30 kb. The coordinates of the α-satellite HOR array(s) are subsequently extended on either side by 1 Mb using BEDtools *slop*, producing the coordinates of the centromeric regions within each contig. Finally, we intersect the centromeric regions with the combined QC BED files using BEDtools (v2.31.1)^59^ and exclude any centromeric region with an assembly error.

#### Annotation of centromeric regions

To determine the sequence composition and organization of each chromosome Y centromeric region, CenMAP (v1.1.0) uses Snakemake-HumAS-SD (v0.1.0; https://github.com/logsdon-lab/Snakemake-HumAS-SD), which combines the α-satellite monomer HMM libraries from HumAS-HMMER (https://github.com/fedorrik/HumAS-HMMER_for_AnVIL) with the rapid annotation of StringDecomposer^122^ to efficiently annotate centromeric regions. Snakemake-HumAS-SD first constructs a comprehensive α-satellite monomer library based on an existing HumAS-HMMER HMM profile (https://github.com/fedorrik/HumAS-HMMER_for_AnVIL/blob/main/AS-HORs-hmmer3.4-071024.hmm). To detect α-satellite HOR structural variants, CenMAP uses an α-satellite HOR variation detection method that generates BED-format annotations of α-satellite HOR variants for each centromere based on the previous StV approach (https://github.com/fedorrik/stv). The method takes α-satellite monomer sequences as input and segments them based on three criteria: (i) adjacent monomers are separated by gaps >160 bp, (ii) adjacent monomers are on opposite strands, or (iii) monomer IDs are not monotonically increasing (on the forward strand) or decreasing (on the reverse strand).

#### Estimation of the length of active α-satellite HOR arrays within centromeric regions

To estimate the length of the chromosome Y active α-satellite HOR arrays, we ran CenStats (v0.0.10) with the `length` subcommand, which leverages the annotations generated by Snakemake-HumAS-SD (described above) to output length statistics of each active α-satellite HOR array. CenStats first uses Snakemake-HumAS-SD α-satellite HOR StV BED files to identify the coordinates of active α-satellite HOR monomers (those with names containing ‘L’). Adjacent active α-satellite HOR monomers are then merged into one block if they are within ∼1.5 α-satellite monomer lengths (256 bp) of each other, and blocks containing fewer than two α-satellite monomers are removed. To account for interruptions of stretches of α-satellite HORs caused by mobile element insertions (MEIs), adjacent blocks of active α-satellite HORs separated by less than 8 kbp are iteratively merged into a single active α-satellite HOR array. Finally, the coordinates of α-satellite HOR arrays that consist of at least five α-satellite HOR monomers and have at least 90% of their total length composed of active α-satellite HORs is outputted as a BED file. To determine the orientation of active α-satellite HOR arrays, we followed a similar procedure, but incorporated both strand orientation and distance to group active α-satellite HOR monomers into arrays.

#### Detection of CDRs within centromeric regions

In addition to annotating centromeric sequences, CenMAP (v1.1.0) uses methylation information in native long-read sequencing data to annotate the CpG methylation profile across each centromeric region. CenMAP first aligns native ONT long-read sequencing data from each genome to the relevant whole-genome assembly using minimap2^60^ (v2.28) with the following command: `minimap2 -x lr:hqae -s 4000 -y -I 8G {ref.fa} {ont_reads.bam}`. CenMAP filters the alignments using SAMtools^58^ (v1.9) -F 2308 to retain only primary alignments and remove partial and secondary alignments. Finally, CenMAP uses CDR-Finder^123^ (v1.0.4), a tool to detect hypomethylated regions termed “centromere dip regions” (CDRs^28^), spanning from the first hypomethylated base to the last hypomethylated base and inclusive of the sequence in between. We manually curated the CDR calls by checking each region outputted in the BED file using IGV^124^, removing false positives, recovering missed calls, and merging calls that were within 150 kbp of each other.

*Estimation of the chromosome Y centromere mutation rate.* To estimate the mutation rate of active α-satellite HOR arrays and flanking sequences for each chromosome Y centromere haplogroup, we identified clades within the phylogenetic tree (see Mathods section ‘Y-chromosomal phylogeny construction and dating’) that were separated by 35,000 years of evolution and contain at least four centromeres, from which one was selected as a reference and the remaining as queries. We then aligned the reference’s active α-satellite HOR array, along with 1 Mb of the flanking pericentromeric sequence on either side, to each query in the same clade using minimap2 (v2.28)^60^ with the following parameters: -x asm20 -K 8G --eqx -Y -r 500000,500000. Unmapped, supplementary, and partial alignments were removed using SAMtools (v1.22)^58^ flag 260. We then generated a BED file of 10-kb windows located within each reference centromere. We used *subseq* (https://github.com/EichlerLab/subseq) to extract query sequences. For each window, the corresponding query sequence was required to be between 8,000 and 12,000 bp in length and exhibit a one-to-one, unambiguous mapping. For sequences in each window, a multiple sequence alignment was performed using MAFFT (v.7.526)^125^ with the following parameters: --maxiterate 1000 --globalpair. The mutation rate per segment was estimated based on Kimura’s model of neutral evolution^126^. In brief, we modeled the estimated divergence (D) as a result of between-species substitutions and within-species polymorphisms, i.e.,

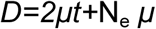

where μ is the mutation rate, t is the divergence time estimated from our phylogenetic tree, and N_e_ is the ancestral human effective population size. We assume a generation time of 20 years. To convert the genetic unit to a physical unit, our computation also assumes N_e_ =10,000 and uniformly drawn values for the generation and divergence times. The estimated divergence (D) is calculated from the Tamura-Nei genetic distance using the libtn93 library (https://github.com/sdwfrost/libtn93?tab=readme-ov-file#api) with a match mode of RESOLVE and a minimum overlap of 100 characters.

*Phylogenetic analysis of centromeric size variation.* Analyses were performed using 123 QC-passing Y chromosomes centromeres with measurements of total active *DYZ3* α-satellite HOR array length and CDR length. All length measurements were natural log transformed prior to phylogenetic analyses so that model coefficients represented proportional changes in size. The amount of HOR sequence outside the CDR was calculated as total *DYZ3* HOR length minus CDR length and was used as a sensitivity analysis. The 142 male Y-chromosome phylogeny (**see Y-chromosomal phylogeny construction and dating**) was pruned to only the 123 chromosomes with complete centromere measurements, and branch lengths were rescaled by the median root-to-tip distance for numerical stability. Phylogenetic covariance matrices were constructed from the shared branch length between each pair of chromosomes from the root to their most recent common ancestor, corresponding to the expected covariance under a Brownian-motion model^115,127^. Phylogenetic signal in CDR, total *DYZ3* HOR, and non-CDR HOR length was quantified using Pagel’s λ^113^. λ was estimated by maximum likelihood over the interval 0–1. Approximate 95% profile-likelihood support intervals were defined using the standard likelihood-ratio criterion, retaining values of λ whose log likelihood was within 1.92 units of the maximum. To test whether CDR length scaled with total *DYZ3* HOR array length, we fitted a phylogenetic generalized least-squares model of ln(CDR) against ln(*DYZ3* HOR), with residual Pagel’s λ estimated by maximum likelihood. The fitted slope was tested against both β=0, representing no relationship with array size, and β=1, representing proportional scaling. To determine whether the observed dispersion of CDR lengths was narrower than expected under the phylogenetic size dynamics of the underlying array, we generated 10,000 multivariate-normal datasets using the covariance structure estimated from total *DYZ3* HOR length, centered on the observed mean log CDR length. The observed CDR fold range was compared with the simulated distribution, with one-sided Monte Carlo P values calculated as (r+1)/(N+1), where (r) is the number of simulations with dispersion equal to or smaller than observed, across all simulations (N). The analysis was repeated using non-CDR *DYZ3* HOR length as a sensitivity reference. Analyses were implemented in Python v3.12.13 using NumPy v2.5.2^128^, pandas v3.0.5^129^, SciPy v1.18.1^104^, and Biopython v1.88 for phylogenetic-tree parsing and pruning^97^; figures were generated using Matplotlib v3.11.1^130^.

### Gene conversion in palindromes

Gene conversion and mutation events were analyzed across seven Y chromosome palindromes (P3–P9; identified *de novo* for each assembly) in 142 male assemblies from the HGSVC, HPRC, and CEPH cohorts. For each palindrome, arm A and arm B sequences were extracted from RepeatMasked^75^ assemblies using coordinates from a reference BED file using samtools faidx^62^, and each arm was aligned to the HG002 arm A sequence using minimap2^60^. A per-sample genotype was then assigned at each variable site by comparing the two arms: positions where both arms carried the same allele were called pseudohomozygous (0/0 or 1/1), and positions where the arms carried different alleles were called pseudoheterozygous (0/1). Sites genotyped in fewer than 80% of samples or exhibiting more than two alleles in a population sample were excluded.

A maximum-parsimony approach (Fitch state reconstruction with parsimony method^131^, was applied to a pruned clade of the HGSVC/HPRC/CEPH Y-chromosome phylogeny to reconstruct the most likely ancestral state at each variable site and assign events to internal nodes. A transition from pseudohomozygous to pseudoheterozygous state was classified as a point mutation; a transition from pseudoheterozygous to pseudohomozygous state was classified as a gene conversion event, provided the converted genotype was supported by the parsimony tree. Sites requiring two or more independent mutations to explain the observed pattern were excluded. Putative gene conversion tracts were identified by clustering conversion events assigned to the same phylogenetic node within a 1 kb window, retaining clusters of at least two sites. For each palindrome, we report the total number of gene conversion events, events per kilobase of arm sequence, the proportion of conversions that restored the ancestral allele (reversions) versus fixed the derived allele, nucleotide conversion bias (A/T to G/C versus G/C to A/T), and the distribution of estimated tract lengths (**Suppl. Fig. 78**).

### Gene expression analysis

*Testis Expression Data and Preprocessing*. The identification of *DAZ* hybrid transcript shown on **Figure 3b** used data from^132^. The other analyses used long-read testis expression data retrieved from the NCBI Sequence Read Archive (Accession: SRR12544672, SRR12544673, SRR31360662, PRJNA911852). The polyA tails of the acquired IsoSeq data were trimmed from the reads using Cutadapt (v1.18)^133^ then aligned to T2T-CHM13v2Y (GCF_009914755.1) using pbmm2 (v1.13.1) with –isoseq presets.

*Extraction of Gene Copies*. The combined annotation of assembly Y chromosome gene loci for 140 individuals was parsed for the coordinates of all copies of *TSPY*, *RBMY*, and *DAZ* using Python. The gene copies were extracted from the personal Y chromosome assemblies using PySam (v0.23.3)^58,85^ and compiled into fasta files by gene name. Finally, the exonic sequence of *RBMY1A1, RBMY1B, RBMY1D, RBMY1E, RBMY1F, RBMY1J, TSPY1, TSPY2, DAZ1, DAZ2, DAZ3*, and *DAZ4* were sequestered from the UCSC Genome Browser^134^ and sorted into Fasta files by gene family.

*Identification of Target Reads*. Trimmed IsoSeq reads were aligned with pbmm2 (v1.13.1) to a Fasta file containing all *RBMY*, *TSPY*, and *DAZ* gene copies from 140 individuals (combinedFiles) and T2T-CHM13v2Y (GCF_009914755.1). As a preliminary filter, reads aligned to the target gene copies with a minimum of >35% accuracy were then mapped with pbmm2 (v1.13.1) to the Y chromosomes of 140 individuals and T2T-CHM13v2. This initial filter removed poor alignments while preserving reads containing novel sequences absent from gene copy annotations. The genome producing the best alignment per read was identified and read alignment coordinates were intersected with the corresponding genome’s annotation using python. Reads were assigned gene IDs according to the annotation overlap (combinedFiles or GCF_009914755.1). *DAZ* exon sequences were then annotated in all target reads using Repeatmasker (v4.1.5) with a custom repeat library. Finally, considering the best alignment for each read, mono-exon reads and reads with an accuracy of < 99% were filtered.

*Genome Alignment Comparisons*. Read alignments extracted from the 141 alignment files and categorized by gene family (*RBMY*, *TSPY*, and *DAZ*) and genome. Read alignment quality was calculated by dividing the number of matching nucleotides by the total read nucleotides. We found the mean read alignment quality per gene family and took the average across families for each sample to identify the genome producing the overall best alignments. The significance of the improvement in read alignment percentages per family afforded by the best overall assembly, HG01358, and T2T-CHM13v2Y were tested by Mann-Whitney U and Bonferroni-corrected. Confusion matrices comparing the annotation of *DAZ* reads using T2T-CHM13v2Y Refseq annotation (GCF_009914755.1) and assembly annotations (combinedFiles) were generated using Matplotlib (v.3.9.4)^130^.

*Visualization of DAZ Expression.* Repeatmasker (v4.1.5)^75,82^ with the Dfam library^84^ (v3.8) was utilized to identify repeats present in the assemblies and T2T-CHM13v2Y. The repeat locations and Encode read alignments to the best matching assembly, HG01358, and T2T-CHM13v2Y were then visualized using gviz (v.1.46.1)^135^ in R. Further, a custom Repeatmasker (v4.1.5) library of *DAZ* exons was implemented to identify *DAZ* exon order and repetition counts in expression data. Then, a Python script was created to illustrate the exon-intron structure of *DAZ* transcripts plotted alongside the genomic architecture of HG01358. Additionally, the target gene read alignments to the assembly producing the best alignments, HG01358, were further scrutinized using the Integrative Genome Viewer (IGV; v2.17.4)^103^ to predict the likelihood of nonsense-mediated decay (NMD) of transcripts containing the unreported *DAZ* exon 29. NMD was predicted for transcripts with a premature stop codon > 50 bp upstream of a splice junction.

### *TSPY* gene expression and copy number variation

*Validate read depth as an approximation for copy number variation.* We used short read 30x whole genome alignments from 1kGP samples (n=130)^16^. The *TSPY1* locus (chrY: 9466954-9469749) was taken from the basic gene level annotation for GRCh38 from GENCODE (https://www.gencodegenes.org/human/). For each 1kGP sample^49^, the mean read depth over the *TSPY1* interval was computed from the CRAM alignments using samtools depth, with zero coverage positions included. Multiple normalized depth metrics to control for sequencing differences were calculated, including normalization by autosomal mean depth (chr 1-22), a single copy Y control (*DDX3Y*), and average depth across XDR windows (1kb regions, MAPQ > 20). 1kGP samples were subset from the same/overlapping available HPRC samples, and a linear model was fit to relate normalized read depth to the assembly HPRC truth *TSPY1* copy number (CN_hat = m*X +b), where X is normalized read depth over *TSPY1*.

*RNA-seq expression and copy number correlation analysis.* The same read depth extractions and normalizations were applied to GTEx^34^ DNA alignments (n=367) over *TSPY1*. The linear model was applied to these GTEx samples to generate sample specific estimated *TSPY* copy numbers converted from normalized read depth. Copy number estimates were then compared to RNA-seq expression (TPM) in the testis to test whether copy number variation is related to gene dosage (https://github.com/sifordrebecca/functional_chrY).

### PAR regions

*Selection of high confident PAR sequences.* To mitigate the effect of assembly errors for downstream analyses, we selected PAR1 and PAR2 sequences for each sample based on assembly status and QC tracks. First, we defined PAR1 as the region ranging from the last coordinate of 5’ telomeric region (“TELO”) plus one to the first coordinate of XDR minus one, and PAR2 as the region ranging from the last coordinate of Yq12 (“HET”) region plus one to the first coordinate of 3’ telomeric region (“TELO”) minus one. We then selected T2T or T2T-scaffold samples that had no structural error (“ERRSTRUCT”) or gap (“NGAP”) within these PAR1 and PAR2, respectively, and no unlocalized sequences annotated with “PAR1” or “PAR2”. The base errors (“ERRBASE”) were masked on the resulting sequences (n=106 for PAR1 and n=126 for PAR2).

*Alignment and visualization of PAR sequences.* PAR1 and PAR2 sequences were all-vs-all aligned by using minimap2 (v2.30 r1287)^60^ with the following command: minimap2 -t ${cores} -x asm10 -c --eqx -D -P --dual=no ${input.fasta} ${input.fasta} > ${output.paf}. The alignments were then visualized using SVbyEye (v0.99.0)^136^ with “plotAVA” function in the phylogenetic order generated from “*Y-chromosomal phylogeny construction and dating*”.

*G-quadruplex (G4) prediction.* Potential G-quadruplex (G4) structures within PARs were identified using Quadron^137^, a machine-learning framework that predicts G4 folding stability based on sequence context and loop architecture. To ensure the inclusion of physiologically relevant structures, we retained only predictions with a Quadron score more than or equal to 19.

*PRDM9 binding motif scan.* Candidate PRDM9 binding sites were identified by scanning sequences with the FIMO (Find Individual Motif Occurrences) tool from the MEME Suite (v5.5.9)^138^. We utilized a Position Weight Matrix (PWM) from JASPAR database^139^, corresponding to the PRDM9 binding motif (https://jaspar.elixir.no/matrix/MA1723.2). Motifs were considered significant at a q-value threshold of < 0.05.

*Statistical test for comparing sequence features across sequence classes.* Statistical comparisons of sequence feature coverages (GC content, G4, *Alu*, L1, SVA, and PRDM9 binding motif) across different sequence classes were performed in R v4.5.1. To account for phylogenetic relationships and the non-independence among different sequence classes, we employed Phylogenetic Generalized Least Squares (PGLS) models using the “gls” function in the nlme R package^140^. Phylogenetic relationships were modeled using the phylogenetic tree generated from “*Y-chromosomal phylogeny construction and dating*”, with the correlation structure defined by Pagel’s λ (“corPagel” function in ape package^141^). The λ parameter was estimated simultaneously with the model coefficients using Restricted Maximum Likelihood (REML). The sample identifier was included as a grouping factor (“form” parameter) to control for intra-assembly correlation. For post-hoc analysis, we utilized the emmeans package^142^ to calculate estimated marginal means and perform pairwise comparisons between groups (PAR1 vs. other classes or PAR2 vs. other classes). Raw p-values were adjusted for multiple testing using the Dunnett’s method.

*LD-based recombination map generation.* To estimate fine-scale recombination rates in the PARs, we utilized pyrho (v0.2.1)^143^. To ensure the independence of the genetic data, related samples were excluded, retaining a single representative per family for each PAR. Using minimap2 (v2.30-r1287) ^60^, each PAR sequence was aligned to the HG002 PAR sequence that was soft-masked for repeats using RepeatMasker (v4.1.7-p1)^82^ (http://www.repeatmasker.org) with -species “human” option. Variants were called using bcftools^62^ mpileup and bcftools call, assuming a ploidy of 1. To generate high-confidence variant sets for LD inference, we applied stringent filtering: only biallelic SNPs with a quality score (QUAL) >= 30, mapping quality (MQ) >= 30, and a minor allele count (MAC) > 1 were retained. We further excluded sites with > 5% missing data, required a depth (DP) of 1 per sample, and filtered for a minimum SNP gap (-SnpGap) of 10 bp to minimize mapping artifacts. Recombination rates were then inferred after generating a custom lookup table assuming a constant effective population size N_e_ of 10,000 and a mutation rate of 1.25 x 10^-8^ per base pair per generation. The Moran sample size was defined as 1.25 times the actual sample size as recommended by the developer. Optimal hyperparameters were determined by minimizing the log(L2), resulting in a window size of 60 and a block penalty of 15 for both PAR1 and PAR2. Final recombination maps were estimated using these optimized parameters and region-specific lookup tables.

*Tandem repeat identification.* Tandem repeats in the PAR1 recombination hotspot were identified using Tandem Repeats Finder^74^. The program was executed with the following alignment parameters: Match = 2, Mismatch = 7, Delta = 7, PM = 80, PI = 10, Minscore = 50, and MaxPeriod = 500.

### Reconstruction of chromosome Y sequences

We adapted our previously developed pipeline, originally designed for polishing consensus haplotype sequences derived from phased short-read genotypes (https://github.com/eblerjana/polishing-pipeline). The pipeline leverages short reads and low-coverage long-read data to correct errors in haplotype sequences. Specifically, it identifies small variants from alignments of short reads to the haplotype sequence and structural variants from long alignments of long reads. The detected variants are then incorporated into the haplotype sequence to correct errors. The original pipeline was developed for polishing haplotypes of a diploid genome and therefore included steps for phasing called variants to be able to distinguish true differences from variants corresponding to the opposite haplotype. For the experiments presented here, we used v2.0 of the pipeline which omits these steps for haploid chromosomes.

We conducted multiple polishing experiments for sample HG01596 with the goal to reconstruct its chromosome Y sequence based on Illumina reads^16^ and low-coverage ONT reads ^33^, starting from the chromosome Y sequences of different samples. As starting points, we selected the phylogenetically closest sample (HG00609), as well as five samples from different continental groups: HG03009 (SAS), HG01258 (AMR), NA12877 (EUR), NA18983 (EAS) and NA19331 (AFR). Additionally, we used the T2T-CHM13 Y chromosome (HG002) as another starting point.

For evaluation, we compared the polished sequences to the chromosome Y assembly of HG01596. We applied our previously developed QV estimation pipeline which calculates local QVs across genomic regions based on variant calls obtained from aligning the polished sequence to the ground truth assembly^13^. Instead of using fixed 1 Mb windows for QV calculation, we modified the pipeline to compute QVs over annotation intervals defined relative to the ground truth assembly (**Suppl. Fig. 83**). Additionally, we computed chromosome-wide QV estimates with Merqury^22^ (k=31) based on Illumina data of the target sample HG01596. Furthermore, we analyzed the distribution of variants identified between the predicted and assembly sequences, capturing differences before and after polishing (**Suppl. Figs. 84-85**).

### Methylation analysis

#### *TSPY* methylation analysis

CpG methylation across *TSPY* array regions was quantified from Oxford Nanopore sequencing data aligned to per-sample verkko assemblies (n = 140; HPRC 106, CEPH 6, HGSVC 28). Methylation calling was performed using modkit (v0.6.0) (https://github.com/nanoporetech/modkit)^144^, reporting only 5-methylcytosine (5mC) CpG dinucleotides with both strands combined per site. Per-sample verkko references were provided for CpG context identification. CpG sites with fewer than 5 covering reads were excluded.

*TSPY* gene bodies were defined from per-sample Liftoff annotations, and each CpG site was classified as genic or intergenic based on overlapping with gene-level features. Per-sample mean methylation was computed separately for genic and intergenic regions.

To assess the effect of the *TSPY* array inversion on methylation, samples were stratified by inversion status and compared using the Mann-Whitney U test. Main array genes (*TSPY*1/3/4/8/9/10) and *TSPY2* were analyzed separately. Methylation was also compared across Y chromosome haplogroups, with groups ordered by mean methylation. All statistical tests were two-sided. Visualizations were generated using matplotlib.

#### Methylation analysis across Y subregions

*Quality control of BAM files.* Quality control of aligned files was performed using the mapq_softclip module of raaqa^145^ (v0.1.1) (https://github.com/kuzsam/raaqa). Mapping quality and soft-clipped base percentages were computed in sliding-window framework along the Y-chromosome, with a window size of 5 kb and a step size of 2.5 kb, producing per-window and per-contig quality profiles used to assess the overall alignment confidence and coverage across examined regions.

*Methylation calling.* DNA methylation calling was performed on Oxford Nanopore sequencing data using modkit^144^. Prior to methylation analysis, BAM files were filtered to retain only high-confidence alignments (MAPQ > 20). For each sample, 5-methylcytosine (5mC) methylation was quantified at CpG sites using the modkit pileup function, with strand information combined. A per-sample Y-chromosome reference sequence was provided to ensure accurate methylation assignment. Methylation calls were filtered using a minimum modified base probability threshold of 0.8, and output was generated in bedGraph format for downstream analyses.

*Methylation on palindromic regions.* DNA methylation patterns across Y-chromosome palindromic regions were analyzed by computing average methylation levels and spatial methylation profiles along each palindrome. For each palindromic copy, methylation values were aggregated across CpG sites and summarized both as overall averages and within fixed genomic bins spanning the length of the palindrome, enabling comparison of methylation distribution along palindromic structures. To facilitate comparison among palindromes of different sizes and across individuals, methylation values were projected onto normalized coordinates corresponding to 0–100% of palindrome arm length. Methylation profiles were summarized within equally spaced position bins and aggregated across individuals using decile (10-quantile) summaries. Palindromic information, including color block assignment and color block order, was retained to facilitate direct comparison among structural contexts. Methylation profiles were further stratified by the palindromic arm to assess epigenetic symmetry between homologous arms. The number of palindromic copies contributing to each analysis was recorded and incorporated into the visualization and downstream interpretation to account for copy-specific representation and structural variability.

*Methylation on ampliconic genes.* Methylation analyses at the level of ampliconic genes were performed by integrating gene annotations with palindrome structure information using a genomic overlap–based approach. Gene annotations and palindromic interval data were converted into GRanges objects (GenomicRanges 1.50.2)^146^ to enable sample-specific overlap analysis. Overlapping gene names and exon-level metadata were aggregated using dplyr^147^ and summarized into a consolidated data frame, which was merged back into the main dataset to annotate each gene-associated interval. Structural metadata, including palindromic arm type, color block assignment (when available), and sequence class order, were retained throughout the analysis.

To distinguish gene-associated from intergenic regions, overlap logic was refined to handle empty datasets robustly and to explicitly define genomic coordinate fields during GRanges construction, minimizing metadata inconsistencies. Multi-gene overlaps were collapsed into single, interval-level annotations using a purrr^147^–based mapping workflow (purrr 1.2.1), and resulting categories were encoded as factor variables for downstream visualization. Gene copy numbers were estimated by normalizing exon occurrences across the Y chromosome and were integrated with methylation levels. A data.table-based workflow was used to pre-collapse gene metadata into a unique reference table prior to merging, with manual garbage collection applied to maintain computational efficiency during large data joins.

*Methylation on rearranged/inverted regions.* DNA methylation data were integrated with curated AZFc structural annotations and architecture classifications by intersecting per-sample CpG methylation calls with annotated genomic regions using a custom Python workflow. Each interval was assigned to a palindromic arm (P#A/P#B) when overlapping palindrome coordinates, while preserving chromosome and strand information; non-overlapping regions were labeled accordingly. For each sample–region combination, the number of CpG sites, mean methylation, and methylation variance were calculated, and *AZFc* architecture information was incorporated by matching sample identifiers. The resulting integrated dataset was used for downstream comparative analyses and visualization in R.

*Methylation data visualization.* All data visualizations were generated using the R statistical environment. Plots were primarily produced with ggplot2 (version 4.0.2)^148^, with additional visualization layers and genomic track representations created using Gviz (version 1.42.1)^135^. Multi-panel figures and composite layouts were assembled using gridExtra (version 2.3)^149^, while selected plots were generated using the lattice package. Consistent color schemes, ordering of genomic features, and factor levels were maintained across visualizations to enable direct comparison between samples, genomic regions, and analytical layers.

*Methylation heterogeneity quantification.* Methylation heterogeneity was quantified using modbamtools^150^ (version 1.0) to assess variability in CpG methylation within defined genomic regions. Filtered BAM files containing Nanopore methylation information were analyzed against BED files defining annotated ampliconic genes, palindromic regions, and major Y chromosome sequence classes (X-degenerate, inverted repeat, pseudoautosomal, and other regions). For each sample, regional heterogeneity scores were computed using the **calcHet** function implemented in modbamtools with default parameters. Unlike average methylation, which summarizes CpG methylation levels across a region, calcHet quantifies methylation heterogeneity within the specified region, thereby capturing variation in methylation states. Higher values indicate greater methylation heterogeneity. Heterogeneity scores are unitless and are not directly comparable to methylation percentages. The resulting output contained one heterogeneity score for each analyzed genomic interval and was used for downstream comparative analyses. For analyses comparing broad Y chromosome sequence classes, individual intervals were first assigned to their corresponding sequence category, and calcHet scores were then summarized within each category for each individual by calculating the arithmetic mean across all intervals belonging to that category. Thus, each individual was represented by a single mean heterogeneity value for each sequence category. For analyses of individual palindromic regions and other defined genomic intervals, the regional calcHet scores were retained without this aggregation. Heterogeneity quantification was performed using a batch-processing workflow to ensure consistent processing across samples.

*Arm-to-arm sequence identity analysis.* For each palindrome, arm-to-arm sequence identity was computed using EMBOSS^151^ stretcher after generating reverse-complement of the second arm. Percent identity was normalized to the length of the shorter arm to account for structural differences. Results were compiled into per-sample and global summary tables. Additionally, regional sequence identity was assessed using a custom Python sliding-window approach (see **Code availability**: sliding_window_identity_batch.py, 200 bp window, 50 bp step), considering only aligned, non-gap positions. All downstream visualizations and summary statistics were generated from the consolidated identity tables. For visualization alongside methylation profiles, sliding-window sequence identity values were projected onto normalized arm coordinates (0–100% of arm length), enabling direct comparison between local sequence conservation and methylation patterns across palindrome arms.

*Analysis of palindromic structure.* Palindrome sequences were analyzed from FASTA files containing both arms. GC content was computed using *seqrequester* (https://github.com/marbl/seqrequester) and refined per arm with *SeqKit*^152^. Low-complexity regions were identified with *dustmasker*^153^, and homopolymers (≥10 bp) were detected using custom *awk* scripts. Microsatellites were annotated using Tandem Repeats Finder (TRF)^74^. All metrics (length, count, and density) were calculated per arm following sequence separation. Results were aggregated into a summary table and visualized in Python using *seaborn*^154^.

*Annotation of non-canonical DNA structures.* Non-B DNA motifs in all accessions were predicted using non-B DNA Motif Search Tool (nBMST)^155^ with default parameters, and G4Hunter^156^ with parameters -w 25 -s 1.2. G4Hunter output was parsed and converted to GFF format using a custom script. G4 motifs were then filtered for stability scores > 1.5, a commonly used threshold corresponding to sequences with high predicted G-quadruplex-forming potential and reduced false-positive predictions. For chromosome-wide visualization, predicted G4 motifs were intersected with Y chromosome sequence class annotations. Local G4 density was calculated as the number of predicted G4 motifs within fixed genomic windows (250kb) along the Y chromosome and normalized by window length. Density profiles were visualized together with sequence class annotations to assess the distribution of G4 motifs across major genomic compartments. Comprehensive code is available in the project code repository. All analyses related to this section were performed using the latest stable versions of Python and R, with package versions and dependencies documented in the accompanying code repository (see **Code Availability** section).

## Data availability

The curated Y-chromosome assemblies generated in this study are available on Genbank with accession numbers listed in **Suppl. Table 1**. The HGSVC assemblies are available under bioproject PRJNA1435860. Raw sequencing data for HPRC, HGSVC, GIAB and CEPH samples are available through the accessions listed in **Suppl. Tables 1,3** and from^13–15^. Restricted-access CEPH samples are available through dbGaP under accession phs003793.v1.p1. Released resources, including quality-control annotations, sequence class annotations, segmental duplication and ampliconic gene annotations, methylation tracks and Y pangenome graph can be found in the IGSR release directory hosted publicly via HTTP and/or FTP (https://ftp.1000genomes.ebi.ac.uk/vol1/ftp/data_collections/HGSVC3/release/chrY_analysis/1.0) and on the Globus end point ‘EMBL-EBI Public Data’ in the directory ‘/1000g/ftp/data_collections/HGSVC3/release’.

## Code availability

All custom code used for generating the HPRC and HGSVC assemblies is available at https://github.com/marbl/gis-bulk-assembly. Codes for manual curation of the graph, GQC annotation and initial sequence class annotation is available at https://github.com/arangrhie/T2T-chrY. Code for sequence class annotation and merging of QC annotations is available at https://github.com/core-unit-bioinformatics/project-chrom-y-extended. Methylation profiling, palindrome architecture and G4 characterization and scripts for visualization are available at https://github.com/bioinf-fi/Y-meth-pangenome.

The polishing pipeline used for reconstructing chrY sequences is available at: https://github.com/eblerjana/polishing-pipeline (v2.0), all scripts and pipelines used to produce the polishing results described in this manuscript are available at: https://github.com/eblerjana/hprc2-companion/tree/chrY-paper.

The PIVOT pipeline used for pangenome graph based inversion recurrence analysis is available at https://github.com/Hufsah-Ashraf/PIVOT.

Custom scripts used for analyses of Yq12, *DAZ*, *AZFc*, gene annotation, mobile element insertions, and de novo mutations are publicly available at: https://github.com/Markloftus/PanYChromosome.

Scripts for calculating normalised read depth and correlations for the *TSPY1* gene and Y callable analyses are available at https://github.com/sifordrebecca/functional_chrY. Code used for *de novo* palindrome assembly and palindrome aided gene conversion is available at https://github.com/makovalab-psu/142_human_Y-chr_palindromes_gene_conversion. Scripts and code used for methylation analysis and visualization of the TSPY arrays are available at https://github.com/shilab/ChrY_Methylation. Code for NucFlag QC is available at https://github.com/koisland/NucFlag-chrY. The CenMAP pipeline used in this analysis is publicly available at https://github.com/logsdon-lab/CenMAP/tree/b1aebc0308b4e05fa42466793f15e8ceef08fdb5. Code used for the analysis of *DAZ*, *TSPY*, and *RBMY* testis expression and visualizations available at https://github.com/gmartinoresearch/testisExpression. Code used for the estimation of variant call QV in callability-masked chrY regions is publicly available at https://github.com/nhansen/chrY_callable_analysis.

## Supporting information

Supplementary results and figures

## Acknowledgements

We thank Marie Kratka for annotation of non-canonical DNA structures (nBMST and G4Hunter). We are grateful to the people who contributed samples through the 1000 Genomes Project. This work used computational resources provided by the National Institutes of Health (NIH) HPC Biowulf cluster; the e-INFRA CZ project (ID:90254), supported by the Ministry of Education, Youth and Sports of the Czech Republic; the ELIXIR-CZ project (ID:90255), part of the international ELIXIR infrastructure (MetaCentrum); and the Penn State Institute for Computational and Data Sciences (RRID:SCR_025154), including access to the Roar Core Facility (RRID:SCR_026424). We also thank the Centre for Information and Media Technology and the Research IT Department of the Medical Faculty at Heinrich Heine University Düsseldorf, the staff at Clemson University for access to the Palmetto HPC, and the Scientific Services at The Jackson Laboratory, including Research IT, for computational infrastructure and support.

We thank the Intramural Research Program of the National Human Genome Research Institute, NHGRI ZIA HG200398 (A.R., S.K., N.H. and A.M.P.); NIH grants U24HG007497 (to E.E.E., T.M., J.O.K. and C.L.), R00GM147352 (to G.A.L.), R01HG002385 and R01HG010169 (to E.E.E.), R35GM151945 and R03HG014804 (to K.D.M.), R01HG014490, U01HG010961, U24HG010262, OT2OD026682 and U24HG011853 (to B.P.), and U01HG013748 (to B.P. and T.M.); NIH National Institute of General Medical Sciences R35GM133600 (to P.A.A., P.B. and C.R.B.); 1P20GM139769 (to M.K.K. and M.L.); and National Institutes of Health (NIH) National Center for Advancing Translational Sciences (NCATS) training award TL1TR001880 (to C.M.). This work was also supported by the MUNI Award in Science and Humanities StG/CoG (MUNI/SC/1916/2024) (to M.C.) and the Jack Baskin and Peggy Downes-Baskin Fellowship (to P.He.). E.E.E. is an Investigator of the Howard Hughes Medical Institute. The contributions of the NIH authors are considered Works of the United States Government.

## Author contributions

Assembly generation: A.R., S.K., J.Li., E.E.E., and A.M.P. Assembly quality control: A.R., S.K., P.E., K.K.O., P.He., N.F.H., B.P. and A.M.P. Y Sequence class annotation: P.H. A.R. M.L. and P.E. Y phylogeny: P.H. and C.L. Segmental duplication annotation: D.Y., and E.E.E. Gene annotation: M.L., F.Y., P.He., B.P., C.L., and M.K.K. Assembly-based variant calling: P.A., and C.R.B. Short-read mapping and variant calling: M.L., R.S., B.J.P., and M.A.W. Callable regions analysis: R.S., N.F.H., M.A.W., and A.M.P. *AZFc* structure: P.H., M.L., D.P., J.O.K., E.E.E. and C.L. *AZFc* deletion and inversion repeats: P.H., M.L., P.B., C.R.B. and C.L. *DAZ* gene analysis: P.H., M.L., and C.L. Gene isoform analyses: G.V.M., L.R., E.E.E. and M.K.K. Functional impact of structural variants: Y.J., J.L., M.J., T.K., X.S., L.N. and M.G. Methylation analysis: M.M., S.Ku., O.P., T.K., X.S. and M.C. Non-B DNA analyses: M.M. and M.C. Palindrome identity: K.P., O.P., M.M., M.C. and K.D.M. Gene conversion in palindromes: K.P. and K.D.M. Yq12 analyses: P.H., M.L. and C.L. *De novo* mutation (DNM) analyses: P.H., M.L., P.A., C.R.B. and C.L. Mobile element insertions: M.L. and M.K.K. Inversion recurrence from graphs: H.A. and T.M. Polishing experiments: J.E. and T.M. Centromere analyses: S.G., C.M. and G.A.L. PAR analyses: K.K. and C.L. Pangenome graph construction: A.G. and P.He. Conceptualisation: P.H., A.R., A.M.P. and C.L. Supervision of the analysis: M.A.W., M.K.K., T.M., M.C., G.A.L., A.M.P. and C.L. All of the authors contributed to writing the manuscript and to the final interpretation of data.

## Declaration of generative AI and AI-assisted technologies in the manuscript preparation process

During the preparation of this work, the authors used ChatGPT (OpenAI) and Claude (Anthropic) to improve language and readability, assist with editorial consistency checks, including figure and table cross-references, and support code development and debugging for computational analyses. After using these tools, the authors reviewed and edited the content as needed and take full responsibility for the content of the published article.

