## Supplementary results and figures for "A population-scale Y chromosome reference resource resolves structural diversity and reduces reference bias"

### Supplementary Results, Extended Data Figures and Supplementary Figures

### Table of content:


**Supplementary Results, Extended Data Figures and Supplementary Figures** 1
**Table of content:** 1
**Supplementary results** 3
Assembly and QC 3
Gene content 3
Variant calling 5
PAR regions 6
Copy-number and motif variation of TSPY and RBMY genes 7
Segmental duplications 8
Mobile element insertions 8
Functional impact of SVs 9
AZFc architectural constraint and potential mechanisms 9
Analysis of inversion recurrence using pangenome graphs 11
*De novo germline mutations* *11*
Palindromes 13
Gene conversion in palindromes 13
Methylation of palindromes 14
G4-motifs 14
Polishing experiments 15
*RBMY, TSPY and DAZ gene expression* *16*
**Extended Data Figures** 18
**Supplementary Figures** 22
Y phylogeny 22
Assembly and QC 24
Gene content 30
PAR regions 37
Segmental duplications and ampliconic regions (incl. palindromes) 46
Mobile Element Insertions 49
Functional impact of SVs 50
*AZFc Region* *52*
*gr/gr recurrent inversions* *59*
*b2/b3 recurrent inversions* *65*
*AZFc region deletion breakpoints* *70*
AZFc constraint modeling 76
*DAZ genes* *82*

*TSPY and RBMY gene families* 99
*TSPY array and genes* 99
*RBMY genes* 109
Centromere 116
Yq12 and DYZ19 123
De novo germline mutations 134
Repeat architecture shapes epigenetic organization across palindromic and ampliconic sequence 139
Palindromes 139
Gene conversion 142
Methylation on palindromes and ampliconic genes 144
Non-canonical structures and G4-motifs 148
Polishing Experiments 150
*RBMY, TSPY and DAZ gene expression* 153
*TSPY gene expression and copy number variation* 163
Y callable regions 166
**References** 168

### Supplementary results

#### Assembly and QC

**Contributing authors:** Arang Rhie, Sergey Koren, and Peter Ebert

Recent HPRC[1], HGSVC[2] and CEPH[3] assembly releases marked improved Y chromosome contiguity and accuracy relative to earlier versions[2–4] (https://humanpangenome.org/hprc-data-release-2/) resolving many segmental duplications and centromeric structures. Nevertheless, 127 out of 136 (93.4%) remained incomplete within the most challenging regions of the chromosome, including the Yq12, ampliconic, and the centromeric region (**Extended Data Fig. 1b, Suppl. Table 2**). To improve recovery of these difficult regions, we generated additional HiFi sequencing data for 20 samples (**Suppl. Table 4**) and constructed alternative assemblies using Verkko v2.2.1. Because the large palindromes and highly identical repeats of the ampliconic region are particularly prone to misassembly, we manually curated 61 Y chromosomes using assembly-graph topology together with long read support (**Suppl. Table 5**). After quality control, 142 assemblies were retained for downstream analyses **(Fig. 1b** and **Suppl. Table 1**).

T2T assemblies were defined as a single contig with no unresolved region (N-gaps), and the presence of both Yp and Yq telomeric sequences (n=46, 32.4%). HG01890 was included in this category despite having two small unlocalized contigs, which are attributable to cell-line mosaicism. T2T-scaffold assemblies have both telomeric sequences present in the main Y but include unresolved gaps (N-gaps) and the corresponding unlocalized contigs (n=86, 60.6%). The unlocalized contigs retained in the gaps could not be unambiguously assigned their order or orientation. This problem was especially pronounced in the Yq12 (54.9%, 439 out of 799) and the ampliconic region (31.7%, 253 out of 799), where highly similar repeat arrays and partially nested palindromic arms complicate unique placement of the unlocalized sequences within this region. All remaining assemblies were classified as other (n=10, 7.0%).

#### Gene content

**Contributing authors:** Mark Loftus, Prajna Hebbar, Feyza Yilmaz, Miriam K. Konkel

We annotated gene content across the 142 Y chromosomes using three complementary approaches: 1) Liftoff, 2) a pangenome graph-based pipeline, and 3) a RepeatMasker exon stitching method (**Methods**). The Liftoff and pangenome graph pipeline were used to annotate all gene classes, whereas the Repeatmasker-based approach was applied specifically to the nine high-identity, copy-number variable ampliconic gene families (*VCY*, *PRY*, *HSFY*, *XKRY*, *CDY*, *BPY2*, *DAZ*, *RBMY*, and *TSPY*) for which we have previously demonstrated the utility of this approach[5].

In the 140 open-access Y assemblies, the number of annotated protein-coding gene copies on the Y chromosome varied depending on the annotation method, from 77-90 copies (Liftoff mean: 90.1, median: 90.0; Pangenome graph mean: 76.5, median: 77.0) (**Suppl. Fig.**

**7**). Most of this difference originated from the ampliconic gene families. By comparison, the number of non-ampliconic protein coding genes was much more consistent, averaging 35.55 copies with Liftoff and 32.91 with the pangenome graph pipeline. Every assembly had exactly one copy of each male specific single copy protein coding gene, showing that this set of genes is highly stable across human Y chromosomes (**Suppl. Fig. 8**).

As the Repeatmasker exon stitching pipeline is not affected by mapping or graph alignments, and has previously proven to accurately annotate ampliconic genes[5], we utilized these annotation results as the reference framework for ampliconic gene presence and absence. Compared with this benchmark, the ampliconic gene content was generally similar to that reported in earlier studies, but there were clear differences among gene families (**Suppl. Fig. 9**). *VCY* showed the least variation, with a copy number of two in 138 out of 140 assemblies (98.57%). In contrast, *TSPY* was the most variable, ranging from 22 to 57 copies per chromosome (mean: 32.77; median: 33.0). Median copy numbers for the remaining families were concordant with previous Y chromosome assemblies[5]: *PRY*: 2 copies; *HSFY*: 2 copies; *XKRY*: 2 copies; *CDY*: 4 copies; *BPY2*: 3 copies; *DAZ*: 4 copies; *RBMY*: 8 copies; and *TSPY*: 33 copies (**Suppl. Table 8)**. Interestingly, ampliconic gene pseudogene counts are strongly gene family–dependent, with the greatest abundance and variability observed in *CDY*, *TSPY*, *RBMY* (**Extended Data Fig. 1d, Suppl. Table 10**). Compared to Repeatmasker baseline, the pangenome graph approach recovered a larger fraction of ampliconic genes versus Liftoff (mean recall of 98.52% versus 93.68%) (**Suppl. Fig. 10)**. The errors of the two methods were also systematic rather than random. Liftoff frequently missed *DAZ* copies, whose copy number is relatively stable but exon structures are highly variable, whereas the pangenome graph-based approach more often struggled with *RBMY* and *TSPY*, which have relatively stable exon structures but occur in long, highly polymorphic tandem arrays (**Suppl. Fig. 11)**. These results highlight that different classes of ampliconic genes pose distinct annotation challenges depending on whether complexity arises primarily from exon diversification or array expansion.

We next asked whether Y chromosome assemblies contained sequences homologous to GENCODE (v49)[6] gene annotations whose reference loci occur elsewhere in the genome. Because highly similar duplicated sequences can inherit the annotation of their closest reference paralog, we refer to these as homology-based annotations rather than assuming that they represent intact or functional gene copies. We identified these loci using the Liftoff whole-genome pipeline and retained annotations with high sequence identity and gene coverage (**Methods**).

Across 140 assemblies, we identified 797 such loci from 21 genes, averaging 5.69 loci per assembly. Most loci (88.6%, 707/797) were homologous to sequences on chromosomes X, 15, 3, and 16 (chrX: 280; chr15: 181; chr3: 140; chr16: 105), with fewer originating from other chromosomes (chr1: 29; chr4: 24; chr2: 12; chr19: 9; chr20: 6; chr7: 4; chr9: 3; chr10: 3; chr14: 1) (**Suppl. Table 9**). These annotations were strongly concentrated in regions of known sequence sharing or duplication, including PAR2 (n = 275), PAR1 (n = 2), X-degenerate regions XDR2 (n = 140), XDR7 (n = 138), XDR6 (n = 2), and the *AZFc* yellow blocks (n = 240). The PAR annotations reflect established regions of X–Y sequence homology, whereas many of the remaining loci likely reflect the extensive history of interchromosomal and intrachromosomal duplication that has shaped the Y chromosome[7,8]. Consistent with this interpretation, 43% of these annotations (343/797) overlapped segmental duplications, including 239 within the *AZFc* yellow blocks of the P1 region.

The 21 annotations belonged to 12 gene families (*CICP, DDX11L, GYG2, MIR4448, MIR6859, RNU6-689P, RPL41P1, SEPTIN14P, SPRY3, UBE2Q2P, WASIR2,* and *PRKX-AS1*), with several represented by multiple pseudogene copies. Importantly, the presence of sequence sufficiently similar to receive a gene annotation does not imply that the Y-linked copy is functionally equivalent to its X-linked or autosomal counterpart. Most annotations were non-protein-coding (82.4%, 657/797), comprising pseudogenes (n = 240), lncRNAs (n = 208), and miRNAs (n = 205), and some protein-coding annotations may similarly represent partial or otherwise nonfunctional paralogous sequence. Previous studies have documented both truncated gene remnants and transcriptionally silenced Y-linked homologues, indicating that functional activity must be evaluated independently of sequence homology. We therefore consider these loci primarily as evidence of duplicated or ancestrally shared sequence rather than as evidence for additional functional Y genes.

Six annotations were restricted to a single haplogroup (*CICP9*_chr10 in haplogroup A, *RNU6-689P*_chr14 in B, *SEPTIN14P17*_chr1 in C, *CICP19*_chr19 and *SEPTIN14P2*_chr2 in E, and *GYG2*_chrX in N), although after multiple-testing correction only enrichment of the chr10-derived pseudogene *CICP9* in haplogroup A remained significant (FDR q = 0.0046). Such lineage-restricted loci could reflect comparatively recent duplication or insertion events, although rare loci are also the most susceptible to assembly or paralog-assignment errors and therefore require independent structural validation. Overall, these results indicate that the single-copy genic core of the Y chromosome is highly conserved, whereas ampliconic genes and duplicated gene-derived sequences record the extensive structural and evolutionary remodeling of the chromosome.

#### Variant calling

**Contributing authors:** Peter Audano
We created a nonredundant callset across our 140 Y chromosomes for two references, GRCh38 and T2T-CHM13, and we apply strict filtering on VNTR loci to reduce bias in alignment-based variant calling (**Methods**). In T2T-CHM13, we identify 535 SVs, 24,664 indels, and 88,736 SNVs. GRCh38 produces similar but lower yields with 425 SVs, 21,911 indels, and 70,591 SNVs (**Suppl. Table 11**). While no T2T-CHM13 variants were fixed in all samples with 100% allele frequency, we note one fixed SV (an insertion), 72 fixed indels, and 119 fixed SNVs in GRCh38 (**Suppl. Table 11**). While SV insertion bias in GRCh38 is well known[9,10], a similar bias for small variants on the GRCh38 Y chromosome has not yet been described.

We compared this callset with our previous results across Y chromosomes[5] after applying the same strict VNTR filtering strategy and identified 1.6X more SVs, 0.6X more indels, and 1.42X more SNVs (**Suppl. Table 13**). In ampliconic loci enriched with high-identity segmental duplications (SDs), new SV insertions found at a rate of 16.1 per Mbp (160 total), which decreases to 2.4 per Mbp (21 total) in X-degenerate regions and 3.2 per Mbp (11 total) in X-transposed regions (**Suppl. Table 12**). As expected, these rates are higher than SV deletions after combining samples (8.8 ampliconic, 1.7 X-degenerate, 1.2 X-transposed per Mbp).

#### PAR regions

**Contributing authors:** Kwondo Kim

The human pseudoautosomal regions (PARs) are short homologous segments shared by the X and Y chromosomes that enable pairing and recombination during male meiosis. PAR1, located at the distal ends of Xp and Yp, spans ~2.7 Mb and undergoes exceptionally high recombination rates (4.33–20.48 cM/Mb), typically ensuring at least one crossover per male meiosis[11]. In contrast, PAR2, located at the distal ends of Xq and Yq, spans ~0.3 Mb and recombines far less frequently (~6.06 cM/Mb)[11]. Their telomeric locations and recombination dynamics give rise to distinctive patterns of sequence diversity and evolution compared with other regions of the Y chromosome. Leveraging fully assembled sequences, we investigated the structural and sequence features of PARs associated with elevated recombination activity.

Analysis of aligned sequences from PAR1 (n = 106) and PAR2 (n = 126) confirmed that PAR1 is substantially more divergent than PAR2 (**Suppl. Figs. 12-13**), consistent with its higher recombination rate. In PAR1, overall sequence identity decreases toward the 5′ telomeric end, forming a positional gradient that mirrors the concentration of recombination activity near the telomere[12]. In contrast, PAR2 shows no comparable gradient, with most regions exhibiting very high sequence identity between the closest phylogenetic pairs (> 99.5%), except at the 3′ distal end. Despite the pronounced diversity in PAR1, within-family divergence among NA12882, NA12883, and NA12886 was markedly lower than divergence among unrelated individuals, indicating that most observed variation reflects long-term evolutionary accumulation rather than *de novo* changes within pedigrees.

We next examined recombination-associated sequence features in PAR1 and PAR2 and compared them with those in 21 other Y-chromosomal regions (**Suppl. Fig. 14**). Overall, PAR1 exhibited a distinct sequence composition, whereas PAR2 shared broadly similar features with other regions. PAR1 showed consistently higher GC content (median = 47.86%; adjusted p-value $\leq 6.17 \times 10^{-11}$) and increased G-quadruplex (G4) coverage (median = 1.73%; adjusted p-value $\leq 6.02 \times 10^{-11}$) compared with all other regions. *Alu* element coverage (median = 1.31%; adjusted p-value $\leq 6.22 \times 10^{-11}$) and PRDM9 binding motif coverage (median = 0.06%; adjusted p-value $\leq 5.30 \times 10^{-3}$) were also consistently elevated relative to most Y-chromosomal regions, with the exception of ampliconic regions 3 (AMPL3) and 4 (AMPL4). In contrast, PAR2 displayed more modest enrichment, with only G4 coverage (median = 0.64%; adjusted p-value $\leq 6.05 \times 10^{-11}$) consistently higher than that of other regions, except for PAR1 and AMPL3.

To further characterize the regional recombination landscape, we generated linkage disequilibrium (LD)-based recombination maps for each region. Consistent with differences in sequence features, fine-scale recombination maps revealed contrasting recombination landscapes across the two PARs. PAR1 exhibited substantial local variation in recombination rate with a median of 9.76 x $10^{-8}$ per bp per generation, whereas PAR2 showed a comparatively uniform profile with a median of 8.71 x $10^{-9}$ per bp per generation (**Suppl. Figs. 15 a, 16a**). Despite these differences, both regions contained localized recombination signals. In PAR1, the localized hotspot overlapped a tandem repeat locus (repeat unit: CCCTCTCCCTGCATCCCCTTTCCCTGTGGGAG[C/-]TGAGAGCTCCTT, unit length: 44 or 45 bp and copy number: 26-186), while the strongest signal in PAR2 coincided with a GA-rich tract (median GA content = 94.59%). Consistent with their high recombination rate estimates, both hotspots exhibited significantly higher GC content than their respective regional backgrounds and extensive haplotype diversity in the flanking regions adjacent to the loci (**Suppl. Figs. 15 , 16**). Although both PARs contained putative G4-forming sequences and PRDM9 binding motifs, their relative contributions differed between regions: G4 motifs predominated at the PAR1 hotspot, whereas PRDM9 binding motifs were more prominent at the PAR2 hotspot, suggesting distinct sequence determinants underlying recombination initiation in these two hotspots (**Suppl. Figs. 15c , 16c**).

Together, these results show distinct sequence features that may be associated with different recombination landscapes in the two PARs. In addition, the discrete recombination hotspots in both PAR1 and PAR2 provide focal points for future mechanistic studies. While both G4-forming sequences and PRDM9 binding motifs are associated with elevated recombination in the PARs, their differing enrichment patterns suggest that their relative contributions may vary between PAR1 and PAR2 hotspots.

#### Copy-number and motif variation of *TSPY* and *RBMY* genes

**Contributing authors:** Mark Loftus and Pille Hallast

Among 110 assemblies that passed region-specific QC, *TSPY1* copy number ranged from 22 to 57 copies per chromosome (mean 32.6, median 32), corresponding to tandem-array lengths of 445 kb to 1.16 Mb (mean 661 kb, median 648 kb) (**Extended Data Fig. 2b; Suppl. Fig. 46; Suppl. Tables 7-8**). *RBMY* copy number ranged from 5 to 12 among 126 QC-passed assemblies (mean 8.26, median 8). The proximal *RBMY* array contained 1–8 copies (mean 3.96, median 4), whereas the distal array contained 1–9 copies (mean 2.28, median 2) and was generally shorter and less variable in size (**Suppl. Tables 7-8**). Variation in predicted transcription-factor motif composition was examined by principal component analysis (**Methods; Suppl. Figs. 50, 55; Suppl. Table 22**). For *TSPY*, PC1 and PC2 explained 55.0% and 16.9% of the variance, respectively, whereas for *RBMY* they explained 93.9% and 3.3%.

Predicted transcription-factor motif composition in the upstream cis-regulatory sequence distinguished array-associated from isolated copies in both gene families (**Suppl. Figs. 50-52, 55-58; Suppl. Table 22**). Array-associated *TSPY1* copies were enriched for motifs corresponding to transcription factors associated with spermatogenesis, morphogenesis, and Sertoli and immune signaling, whereas the isolated *TSPY2* copy showed a distinct profile linked to neural and sensory development. *RBMY* groups were more similar, although array-associated copies contained additional motifs corresponding to transcription factors associated with stress responses, angiogenic/hypoxic signaling, and tissue remodeling.

#### Segmental duplications

**Contributing authors:** DongAhn Yoo
Examining segmental duplication (SD) of the nearly complete Y chromosomes, we identify 13.9 (9.9-18.2) and 12.6 (6.2-21.1) Mb/sample of inter- and intrachromosomal SDs, respectively (**Extended Data Fig. 1e, Suppl. Fig. 17, Suppl. Table 14**). Among different haplogroups, we find reduced inter- and intrachromosomal SD content for haplogroup D and N, largely restricted to individuals from East Asian and Eurasian ancestry; however only haplogroup N passes Bonferroni multiple test correction showing significantly lower intrachromosomal SD content. On the other hand, greater duplication content was observed for haplogroup E, found predominantly in individuals with African ancestry. This is consistent with the genome-wide trend observed for higher SD content among individuals of African ancestry[10,13]. The pairwise SD sequence identity is in general similar to that observed for T2T-CHM13v2 (**Suppl. Fig. 18**). With one exception, we observe a distinct pattern of more divergent intrachromosomal SDs (93-94.5%) among Y haplogroups Q and R. These SDs are specifically enriched for the *TSPY* and *FAM197* gene families. Assessing paired chromosomes for interchromosomal SDs, we find that the largest SDs paired with acrocentric chromosomes (**Suppl. Fig. 19**), consistently in all haplogroups, in exception for D and J showing less acrocentric SDs than the other haplogroups, and G with more SDs (**Suppl. Fig. 20**).

#### Mobile element insertions

**Contributing authors:** Mark Loftus and Miriam K. Konkel
Mobile element insertions (MEIs) are important to study because they can disrupt or alter gene function[10]. They also serve as substrates for *Alu*-mediated recombination[14]. As absence of an insertion represents the ancestral state, Y chromosomal insertions can be distinguished from deletions containing MEI sequence based on haplogroup information. Here, we limited our analyses to non-long terminal repeat (non-LTR) elements (*Alu*, L1, and SVA), as these are the only transposable elements mobilizing in the human population[15], and to 140 of 142 male samples with open-access data (excluding dbGaP-restricted males). Across these 140 Y chromosome assemblies, we identified only 46 putatively polymorphic young MEIs (36 *Alu* elements and 10 L1s), highlighting the overall rarity of MEIs on the Y chromosome (**Suppl. Fig. 21a, Suppl. Table 15)**. Nearly half of these insertions (47.82%, 22/46) were specific to an individual, while the remaining elements were dated to insertion

times ranging from 84 to 157,461 years ago based on the most recent common ancestor (TMRCA) of carriers (**Suppl. Fig. 21b**).

Analysis of insertion density across Y chromosome sequence classes revealed distinct regional patterns (**Suppl. Fig. 21c**). The pseudoautosomal region 1 (PAR1) was enriched for *Alu* element insertions (10/36, 27.78% of all *Alu* insertions), while the centromeric region had the highest density of L1 insertions (per Mbp) and was relatively enriched for L1s compared to *Alu* elements, consistent with a hotspot for L1 activity. Notably within the centromere, we identified two unique L1 insertions in O lineages located within higher-order repeat (HOR) arrays, including one full-length L1 element shared across seven males in the O2 lineage that retains intact open reading frames (ORFs). Despite the large size of the Yq12 region, comprising nearly half of the Y chromosome, we identified few *Alu* insertions (3 - only counting elements which have not been duplicated within the region yet) and zero L1 or SVA elements, indicating a striking depletion of MEIs in this heterochromatic domain.

#### Functional impact of SVs

**Contributing authors:** Yunzhe Jiang and Matthew Jensen
To account for the heterogeneous sequence composition of chromosome Y, we evaluated SV enrichment separately in PAR1 and the MSY using a compartment- and context-matched permutation framework. In the MSY, insertions and deletions were enriched in protein-coding gene bodies in both GRCh38 ($log_2FC$ = 1.48 and 1.46, respectively) and hs1 ($log_2FC$ = 1.60 and 0.79), with similar enrichment in protein-coding introns (GRCh38: $log_2FC$ = 1.58 and 1.52; hs1: $log_2FC$ = 1.76 and 0.79). Pseudogene overlaps were depleted for both SV types in GRCh38 ($log_2FC$ = −1.06 and −0.66) and hs1 ($log_2FC$ = −1.14 and −1.13). Protein-coding exons were enriched for MSY deletions in GRCh38 and hs1 ($log_2FC$ = 2.24 and 0.90), but not for insertions. In PAR1, insertions were depleted from protein-coding gene bodies in both assemblies (GRCh38: $log_2FC$ = −0.20; hs1: $log_2FC$ = −0.19), and both SV types were depleted from exons and pseudogenes. PAR1 deletion patterns differed between references: protein-coding gene bodies and introns were enriched in GRCh38 ($log_2FC$ = 0.30 and 0.36), whereas hs1 showed weak protein-coding depletion ($log_2FC$ = −0.11) and no significant intronic enrichment. lncRNA associations also varied by compartment, SV type, and reference assembly. These results indicate that the distribution of SVs relative to genomic elements differs between PAR1 and the MSY and cannot be represented adequately by a single chromosome-wide enrichment estimate (**Suppl. Figs. 22–23**).

#### AZFc architectural constraint and potential mechanisms

**Contributing authors:** Mark Loftus, Pille Hallast

We tested whether the limited range of observed *AZFc* architectures could arise under a permissive model of repeat-mediated structural change. The primary analysis included 98 chromosomes representing 10 internally modeled architectures. Both equally parsimonious ancestral states contained 17 modeled repeat blocks, whereas observed architectures ranged from 11 to 23 blocks, giving $D_{max}$= 6 (**Supplementary Figure 35**). Under the dynamic NAHR model, simulated chromosomes generally showed substantially larger departures from the ancestral block count (**Supplementary Figure 36)**. At 30 state-changing events, the median null $D_{max}$ was 14 with Architecture:1 (HG01890-like) as the root and 12 with Architecture:3 (GRCh38-like) as the root; the 2.5th percentile was 12 for both. Only 2/1,000 and 1/1,000 simulations, respectively, had $D_{max}$ ≤ 6, corresponding to Monte Carlo *P*=0.002997 and *P*=0.001998 (**Suppl. Table 18**). Using the larger root-specific value as a conservative summary gave *P*=0.002997. The result remained significant when conditioning on 31 events (*P*=0.001998) and 32 events (*P*=0.000999) (**Suppl. Table 18**).

Other measures of terminal diversity were less unusual. At 30 events, the observed number of terminal architectures was not significantly reduced relative to the null (*P*=0.295 and 0.293 for the two roots), nor was Shannon entropy (*P*=0.093 and 0.111). Dominant-state frequency and total block-count span also showed only modest departures from expectation. The strongest departure from the null was therefore the limited excursion in block copy number rather than a general reduction in the number or diversity of architectural states.

We then tested whether the restricted state space could reflect preferential loss of older rearrangements. Simulated event histories were reduced to terminal architectures, and ancestral changes were reconstructed using the same parsimony procedure applied to the observed chromosomes. Reconstructed changes in the observed data were somewhat younger than expected, but the difference was not significant: conservative *P* values across the two possible roots were 0.157, 0.181 and 0.199 when conditioning on 30, 31 and 32 events, respectively. The observed 23 reconstructed architecture changes were also within the range expected under the ascertainment-matched simulations. We therefore found no clear evidence that preferential removal of older rearrangements accounts for the restricted state space.

Finally, we tested whether one-step rearrangements preferentially involved physically closer or longer homologous repeat blocks. Physical block coordinates were reconciled with the symbolic architecture model for all 98 chromosomes, yielding 1,504 sample-specific state-changing NAHR opportunities. Eighteen reconstructed branches, representing five distinct transitions, could be explained by a single modeled NAHR event. Observed substrates showed a modest tendency toward shorter physical separation in the broad comparison (mean distance percentile=0.433, *P*=0.085), but this was weaker after matching by repeat block family (0.456, *P*=0.225) and was absent when each distinct transition was counted once. Substrate length likewise did not differ significantly from matched opportunities when measured using the shorter homologous block (broad *P*=0.458; family-matched *P*=0.134) or alternative length summaries. These analyses did not identify a general effect of physical proximity or homologous-block length on substrate use.

#### Analysis of inversion recurrence using pangenome graphs

**Contributing authors:** Hufsah Ashraf, Andrea Guarracino

PIVOT was able to locate and analyze all 22 potential inverted loci in the region spanning inverted repeats IR1 to IR5 and palindromes P1 to P8, directly from the chrY pangenome graph generated using PGGB (**Methods, Fig. 2e**). This is a substantial improvement over the HPRC r1 chromosome Y PGGB graphs[4], where none of the Y chromosome inversions could be assessed for recurrence, either due to the absence of anchor nodes or due to low-quality, fragmented assemblies containing contigs not long enough to span the inverted loci together with a flanking non-inverted region required to synchronize the orientation of extracted subpaths across haplotypes[16] **(**Chapter 5**) (https://github.com/Hufsah-Ashraf/PIVOT)**. PIVOT detected evidence of inversion recurrence for IR2-, IR3-, P7-, and P8-mediated inversions (**Fig. 2e**). Additionally, we observed a recurrence signal in the region between palindrome P4 and the first IR2 copy, indicating recurrent rearrangements in this region. In our cohort, only one sample, NA20905, carries the IR1 inversion. Structural comparison of this haplotype with T2T-CHM13v2 showed that the region between the first copy of IR1 and the second copy of IR3 is inverted in place, whereas, in the case of an IR1 inversion relative to T2T-CHM13v2, it would be expected to shift to the distal end. Additionally, we observed that the region between the first copy of IR3 and the first copy of IR1 is shifted to the distal end in the NA20905 haplotype, while maintaining the same orientation as T2T-CHM13v2 (**Fig. 2e**). These observations suggest the occurrence of intermediate inversion events leading to the NA20905 haplotype. One possible sequence of events (as carried by NA19331) involves an IR3-mediated inversion flipping the first IR1 repeat copy, and another, independent inversion toward the distal end flipping the region between IR1-g2 and IR1-b3 (**Fig. 2e**). A subsequent ~16Mb inversion between the inversely oriented first IR1 copy and the IR1-b3 results in the haplotype structure observed in NA20905 (**Fig. 2e**). None of the samples in our cohort carry an IR4-mediated inversion. Although at least two haplotypes carrying an inversion are required to determine recurrence, considering the haplotype architecture and the observed recurrence signal across the region by PIVOT particularly close to both repeat copies, we hypothesize that both of these loci have a potential to recurrently invert in different haplotype backgrounds. Note that PIVOT analyzes only non-repetitive regions to detect recurrence evidence. Therefore, in cases where repeat copies are too close or in repeat-rich regions, such as the region between IR2 and the last IR5 copy, it lacks sufficient power to detect recurrence. Consequently, the recurrence signal in this region is observed primarily between P4 and the IR2 repeats.

#### *De novo* germline mutations

**Contributing authors:** Mark Loftus and Pille Hallast

The Y-chromosome assemblies included eight samples from the CEPH Platinum pedigree[3,17], enabling the identification of *de novo* mutations (DNMs) across generations. Both pedigrees belonged to haplogroup R1b: individual 200080 (R1b-Z326) was analyzed together with his two sons, and NA12877 (R1b-Z302) together with his four sons (**Fig. 1b,**

**Suppl. Fig. 71**). For each family, the paternal Y-chromosome assembly was used as the reference to identify *de novo* variants in the corresponding sons across the non-recombining region of the Y chromosome (approximately 51.4 Mb for 200080 and 48.8 Mb for NA12877). Candidate variants were then subjected to extensive filtering and validation (**Methods, Suppl. Table 28**).

Across the six father-son transmissions, we detected a total of 53 DNMs, corresponding to 6 - 13 variants per transmission and averaging 7.2 SNVs, 0.8 indels and 0.8 structural variant per transmission (**Supp. Tables 28-29**). Based on the callable male-specific Y sequence in these pedigrees, this corresponds to a mean per-transmission SNV rate of $1.46 \times 10^{-7}$ (95% CI: $9.14 \times 10^{-8}$ - $2.02 \times 10^{-7}$) per bp per generation across the male-specific Y overall, with rates of $7.62 \times 10^{-9}$ (range: 0 - $4.57 \times 10^{-8}$) in euchromatic sequence and $2.82 \times 10^{-7}$ (95% CI: 1.78 - $3.87 \times 10^{-7}$) in Yq12 (**Supp. Table 28**). Four of the 53 DNMs were located in euchromatic or centromeric/pericentromeric sequence, whereas the remainder mapped to Yq12 (**Suppl. Table 29)**. Considering SNVs alone, only one of 43 occurred in euchromatic sequence across six transmissions, consistent with previous reports[3,18,19], while two mapped to the pericentromeric region and 40 to Yq12. These results show that both the count and rate of *de novo* SNVs are strongly elevated in the distal Yq12 heterochromatin relative to euchromatic Y sequence.

We next examined the sequence context of the 40 *de novo* SNVs identified in Yq12. Because some SNVs occurred in tightly clustered groups, we first collapsed fully concordant multi-SNV clusters into single nonredundant events. Using a maximum distance threshold of 300 bp, this reduced the dataset from 40 SNVs to 35 unique events, reflecting the collapse of 9 clustered SNVs into 4 representative multi-SNV events. Each of these 35 events was then analyzed using a donor-discovery and post-classification framework to identify homologous donor tracks (≥200bp) and assign events to one of three categories: i) *de novo* mutation, ii) gene conversion (GC) candidate, iii) likely gene conversion (**Methods, Suppl. Fig. 73, Suppl. Table 30)**. Briefly, the donor-discovery and post-classification framework evaluated whether *de novo* SNVs in the Yq12 *DYZ1/DYZ2* repeat arrays were more consistent with isolated mutation or gene conversion by identifying homologous donor copies in the paternal assembly and comparing their alleles to the son's alternate allele. Candidate donors were filtered for assembly quality, distance, and flanking homology, then singleton and clustered SNVs were statistically assessed and classified as DNM, GC_candidate, or Likely_GC (**Methods**).

This analysis classified 26 of the 35 nonredundant events (74.3%) as DNMs, 3 (8.6%) GC candidates and 6 (17.1%) as likely GC events. The six likely GC events included four multi-SNV clusters, including three clusters containing two SNVs and one cluster with three SNVs, as well as two single-SNV events that were significant in the donor-discovery pipeline (one-sided binomial test and FDR correction = $1.595 \times 10^{-203}$ and $1.4 \times 10^{-4}$). The nearest perfectly homologous donor (≥200bp) in the paternal assembly for the nine GC candidate and likely GC events were located 1,657 bp - 2.26 Mb away from the mutation loci (mean distance ~715 kb, median distance 40,763 bp, with 7/9 events having the closest donor <80-kb away) and the number of homologous donor sequences ranged from 1-2727 (mean

520, median 18) (**Suppl. Figs. 74-75**, **Suppl. Table 30**). Donor tracts spanned 217 - 681 bp of perfect flanking homology, with a mean length of 426 bp.

Thus, most Yq12 substitutions were most consistent with *de novo* mutation, but a substantial minority were more consistent with interlocus templated copying than with independent substitution events. Although sequence comparison alone cannot resolve the precise mechanism, these results indicate that a subset of Yq12 *de novo* SNVs arise in the context of highly similar homologous tracts elsewhere in the paternal assembly.

#### Palindromes

**Contributing authors:** Karol Pal and Kateryna D. Makova
The most complex palindrome architecture involved the *DAZ* gene regions. Here, the palindrome-calling algorithm[20] detected separate short *DAZ*-containing inverted repeats. Most assemblies contain two "*DAZ* palindromes" (n=118), but single (n=19) and three (n=5) copies were also observed. Across assemblies, P1 and P2 showed a striking mutually exclusive structural configuration (**Suppl. Fig. 76**). In most Y chromosomes, P1 retained its canonical form (89 occurrences) while P2 was represented solely by the smaller *DAZ* palindrome, consistent with the structure described in Skaletsky et al.[8]. In the remaining assemblies, the *gr/gr* inversion expanded P2 substantially (35 occurrences, **Fig. 2c**) while simultaneously disrupting the inverted repeat structure of P1, matching the alternative architecture observed in T2T-CHM13v2Y/HG002[21].

Overall, these results show that human Y chromosome palindromes are far more conserved than their surrounding structural volatility might predict. Even within the highly dynamic P1/P2 interval, short *DAZ*-containing palindromes remain consistently preserved, forming local islands of architectural stability within one of the most rearrangement-prone regions of the chromosome.

#### Gene conversion in palindromes

**Contributing authors:** Karol Pal and Kateryna D. Makova
To investigate gene conversion dynamics within Y chromosome palindromes, we applied a maximum-parsimony approach to 142 male Y chromosome assemblies spanning seven palindromes (P3–P9), reconstructing mutation and gene conversion events across the Y chromosome phylogeny. Briefly, pseudohomozygous-to-pseudoheterozygous transitions at variable sites were classified as mutations, while the reverse transitions were classified as gene conversion events (**Methods, Suppl. Table 33**). We found a total of 3239 gene conversion events. Excluding 571 transversions (A-T or G-C swapped), we found that gene conversion events had a significant GC-bias with 1,580 AT to GC events compared to 1,088 GC to AT events (Chi-square test, $p = 1.65\times10^{-21}$). Additionally, we found a significant trend toward the ancestral state. Out of 1710 events in which the root node is reconstructed as homozygous, we found with 905 reversals of the mutated allele compared to 805 fixation events ($p = 0.0156$). Median conversion tract length, estimated from co-occurring events, was 542 bp, with a maximum of 8.5 kbp. GC bias has been consistently reported across

previous studies[5,22–24]. Regarding directionality, while Hallast et al. (2013)[22] and Skov & Schierup (2017)[23] reported a significant excess of conversions toward the ancestral state, a subsequent study found a non-significant trend in the opposite direction[24]. Our observed ancestral-to-derived ratio (905:805) shows a stronger skew toward the ancestral state than Hallast et al. (2023) (374:357), which now becomes  statistically significant. The previous results are more likely underestimated as reversals “fixing” a mutation fast enough within a branch of the phylogenetic tree would become undetectable. The larger dataset presents a more realistic picture.

#### Methylation of palindromes

**Contributing authors:** Mariateresa Mazzetto and Monika Cechova

We analyzed 5-methylcytosine (5mC) profiles in 134 individuals, focusing on repetitive elements and palindromic structures (**Fig. 5a**). Palindromes also showed elevated methylation heterogeneity relative to most other sequence classes (**Suppl. Fig. 80, Suppl. Tables 7, 34**). Measuring how similar the DNA sequences are within each arm showed that structural changes are a major driver of methylation differences. In the same way, when the DNA sequence was less similar between the two arms, there was greater methylation variation, linking structural changes to less stable methylation.

But looking at where these changes occur showed that structure alone does not fully explain the methylation patterns. Each palindrome had its own unique methylation pattern: P7 and P8 had typical patterns, while P4, P3, and P1 showed clear differences between the left and right arms that could not be explained by how similar the sequences were. Across AZFc, methylation was lower overall, regardless of structure. These results demonstrate that uneven methylation across Y chromosome palindromes arises from both structural differences and their locations, not just from the type of rearrangement (**Suppl. Fig. 80d**).

We then examined how the gene arrangement in these repeated DNA regions affects methylation. Beyond simple gene identity, the complex spatial organization of ampliconic regions serves as a primary determinant of local epigenetic landscapes. Analysis reveals that while methylation levels follow a modest, non-linear relationship with exon-normalized copy number (**Suppl. Fig. 81a**), methylation heterogeneity is more strictly copy-dependent, with fewer copies showing greater differences (**Suppl. Fig. 81b**). Where the genes were located also made a difference in methylation across ampliconic blocks and among paralogs. BPY2 and VCY genes showed differences in methylation depending on the order of the copies (**Suppl. Fig. 81c-d**), while DAZ methylation changed based on block location, copy order, and inversion status (**Suppl. Fig. 81e-g**). These data demonstrate that methylation patterns of Y-linked genes with multiple copies depend not just on which gene it is, but also on how the genes are arranged and where they are in the repeated regions.

#### G4-motifs

**Contributing authors:** Mariateresa Mazzetto and Monika Cechova

G-quadruplexes (G4s) are non-canonical DNA secondary structures formed by guanine-rich sequences and have been implicated in the regulation of replication, transcription, and genome stability. While G4s are widespread across the genome, their distribution and potential functional roles within highly repetitive regions, such as those of the human Y chromosome, remain poorly characterized. To address this, we systematically mapped predicted G4-forming sequences across telomere-to-telomere Y chromosome assemblies.
Analysis of non-B DNA motif composition revealed substantial variability across sequence classes, with multiple motif types contributing to G4 signal **(Suppl. Fig. 82a**). Across telomere-to-telomere Y chromosome assemblies, G4 density exhibits pronounced inter-individual variability, while maintaining conserved large-scale patterns along the chromosome, as shown by clustering of haplotypes based on G4 distribution (**Fig. 5d, Suppl. Fig. 82**). Within pseudoautosomal regions, G4 density is not uniform but varies along the telomere–centromere axis, with distinct local enrichment patterns in PAR1 and PAR2 (**Suppl. Fig. 82d**). Notably, palindromic regions show highly heterogeneous G4 content, with substantial differences in both total G4 counts and sequence diversity across palindromes (**Suppl. Fig. 82e-f**). For example, P1 and P5 harbor the highest number of G4 motifs and unique sequences, whereas other palindromes exhibit more limited G4 representation. Fine-scale analysis of palindrome arms reveals distinct structural profiles of G4 density, with reproducible patterns across individuals and asymmetries between arms in specific regions (**Suppl. Fig. 82g**). Together, these results indicate that G4 motifs are preferentially associated with structurally complex and repetitive regions of the Y chromosome and display both conserved and variable features across individuals, suggesting a potential link between local sequence architecture and non-canonical DNA formation.

### Polishing experiments

**Contributing authors:** Jana Ebler and Tobias Marschall
Reconstruction of Chromosome Y sequences was done using a modified version of our previously developed polishing pipeline (https://github.com/eblerjana/polishing-pipeline). The pipeline starts from an initial Chromosome Y reference sequence and uses sequencing data of the target sample to reconstruct its Y chromosome. In our experiments, the target sample was HG01596 (haplogroup O2a1b1a1a1a1c, **Suppl. Table 1**) and we used Y chromosome assemblies of HG01596 itself, and six different samples as well as the T2T-CHM13v2Y (HG002, J1a2a1a2c1a1) as references to initialize the pipeline. These samples included the phylogenetically closest one, HG00609 (O2a1b1a1a1a), as well as five samples from different continental groups: HG03009 (SAS, H1a1a4b2c1a1a), HG01258 (AMR, J1), NA12877 (EUR, R1b1a1b1a1a2e2), NA18983 (EAS, D1a2a1a2b1) and NA19331 (AFR, E1b1b1b2b2a1) (**Suppl. Fig. 1**).
In the first step, we align Illumina data[25] of the target sample to the initial Y chromosome reference with strobealign (v0.16.1)[26]. We then call SNPs and indels with DeepVariant (v1.8.0)[27]. Next, we align low coverage ONT reads[28] of the target sample with minimap2 (v2.3)[29] and call structural variants with Sniffles2 (v2.6.1)[30]. Finally, variant calls are incorporated into the initial Y chromosome sequence with bcftools[31] consensus. The original pipeline (v1.0) was developed for polishing diploid genomes and thus included additional

steps to phase variants in order to distinguish true differences from variant calls originating from the opposite haplotype. For our experiments presented here, we used v2.0 of the pipeline which omits these steps for haploid chromosomes.

For evaluation, we applied Merqury (v1.3)[32] with k-mer size of 31 to compute chromosome-wide QVs by comparing the polished Y chromosome sequence to Illumina reads of the target sample. We additionally applied our previously introduced variant-based QV estimation pipeline[10] which compares the reconstructed sequence to the Y chromosome assembly of HG01596 to compute QVs for different regions of the chromosome based on variant calls generated by PAV[33]. The original pipeline computed variant-based QVs across windows of 1 Mb in length. We modified it to compute QVs for annotation intervals defined relative to the ground truth assembly (**Suppl. Fig. 83**). We additionally analyzed the distribution of variants identified between the predicted and assembly sequences, capturing differences before and after polishing (**Fig. 5e, Suppl. Figs. 84-85**).

#### *RBMY*, *TSPY* and *DAZ* gene expression

**Contributing authors:** Gianni Martino, Luyao Ren, and Miriam K. Konkel

We implemented a read-alignment quality scoring pipeline (**Methods**) to determine the best genome for alignment of ampliconic gene expression in testis belonging to a single individual (SRR31360662) outside of the 142. Alignment of the transcripts to the respective *DAZ* gene copies revealed expression of all four *DAZ* genes in our testis samples. Furthermore, we identified transcripts with presence of all possible exons for each gene, including up to three RNA recognition motifs (RRMs). Following alignment of reads for three ampliconic genes (*RBMY*, *TSPY*, and *DAZ*) to 140 Y chromosomes and T2T-CHM13v2Y, HG01358 was determined to provide the highest quality alignments with five others producing similar quality results (HG01934, NA12877, NA20509, NA12882, NA12886). Alignment to HG01358 improved median read alignment accuracy in all three genes (*RBMY*: +0.002%, adjusted p=0.132; *TSPY*: +0.015%, adjusted p=2.30e-16; *DAZ*:+8.222%, adjusted p=5.26e-22; **Fig. 5g, Suppl. Fig. 86, Suppl. Table 36**) when compared to T2T-CHM13v2Y. Thus, in particular the highly variable *DAZ* region exemplifies the need for mapping to the assembly of a genetically similar individual for transcriptomic analysis of structurally highly variable regions.

The ability to unambiguously map *DAZ* transcripts across the four copies revealed unreported splicing events including, intron retention between exon 7 subtypes (**Suppl. Figs. 92-95**), LINE-containing alternate transcription start sites (**Suppl. Figs. 92**), and two alternate last exons. The first and more prominent unreported 3’ exon, exon 29 (**Fig. 3c,** purple), was present in *DAZ1, DAZ2,* and *DAZ3* while the other, exon 30 (orange) was observed only following exon 29 in *DAZ1*. Of the three transcripts containing exon 29 (**Fig. 3c,** *DAZ1* transcript 4, *DAZ2* transcript 3, and *DAZ3* transcript 3): 1) *DAZ1* transcript 4 contains a truncated exon 28 splicing to exon 29 then again to exon 30. 2) *DAZ2* transcript 3 contains only exon 1 with start codon splicing directly to exon 29. 3) *DAZ3* transcript 3 lacks exon 28, splicing from exon 27 to 29. Only the second transcript produces an altered putative protein, while the other two transcripts contain the typical stop codon in exon 27. Additionally, *DAZ1* transcript 4 is predicted to undergo nonsense-mediated decay based upon the presence of a stop codon upstream of two splice junctions, while *DAZ2* transcript 3 and *DAZ3* transcript 3 are not predicted to undergo NMD.

To better understand the improvements afforded by the 140 Y chromosome as a transcriptomic analysis resource, we performed a series of comparisons between transcriptomic pipelines using the ENCODE individual testis data (SRR31360662), assemblies, and the reference genome assembly, T2T-CHM13v2Y. We first visualized the position of T2T-CHM13v2Y within the distribution of overall read alignment quality for the three ampliconic genes among the 140 individual assemblies (**Fig. 5f**). Surprisingly, while T2T-CHM13v2Y placed in top 50% of assemblies for *RBMY* and *TSPY*, it placed in the bottom 5% (7th/141 genomes) for *DAZ* alignment accuracy. While we do not know the background of the ENCODE individual beyond Asian ancestry, several other assemblies provided better matches than T2T-CHM13v2Y. In the context of the *DAZ* alignment quality distribution curve this explains the relatively poor performance of the assemblies most divergent from the R1 group, and strengthens the importance of using a genome from the same haplogroup for transcriptomic analysis of highly variable loci. We next explored the impact of these improved alignments on annotation of reads to the *DAZ* gene copies by comparing pipelines using T2T-CHM13v2Y versus one of the best performing assemblies, HG01358 (**Suppl. Fig. 90**). The comparison revealed only 19.7% of read annotations were in agreement (30/152 reads) between the two reference annotations. Interestingly, the concordance of HG01358 and HG002, the same individual as T2T-CHM13v2Y, was notably higher, 66.4% (101/152 reads; **Suppl. Figs. 90-91**).

### Extended Data Figures

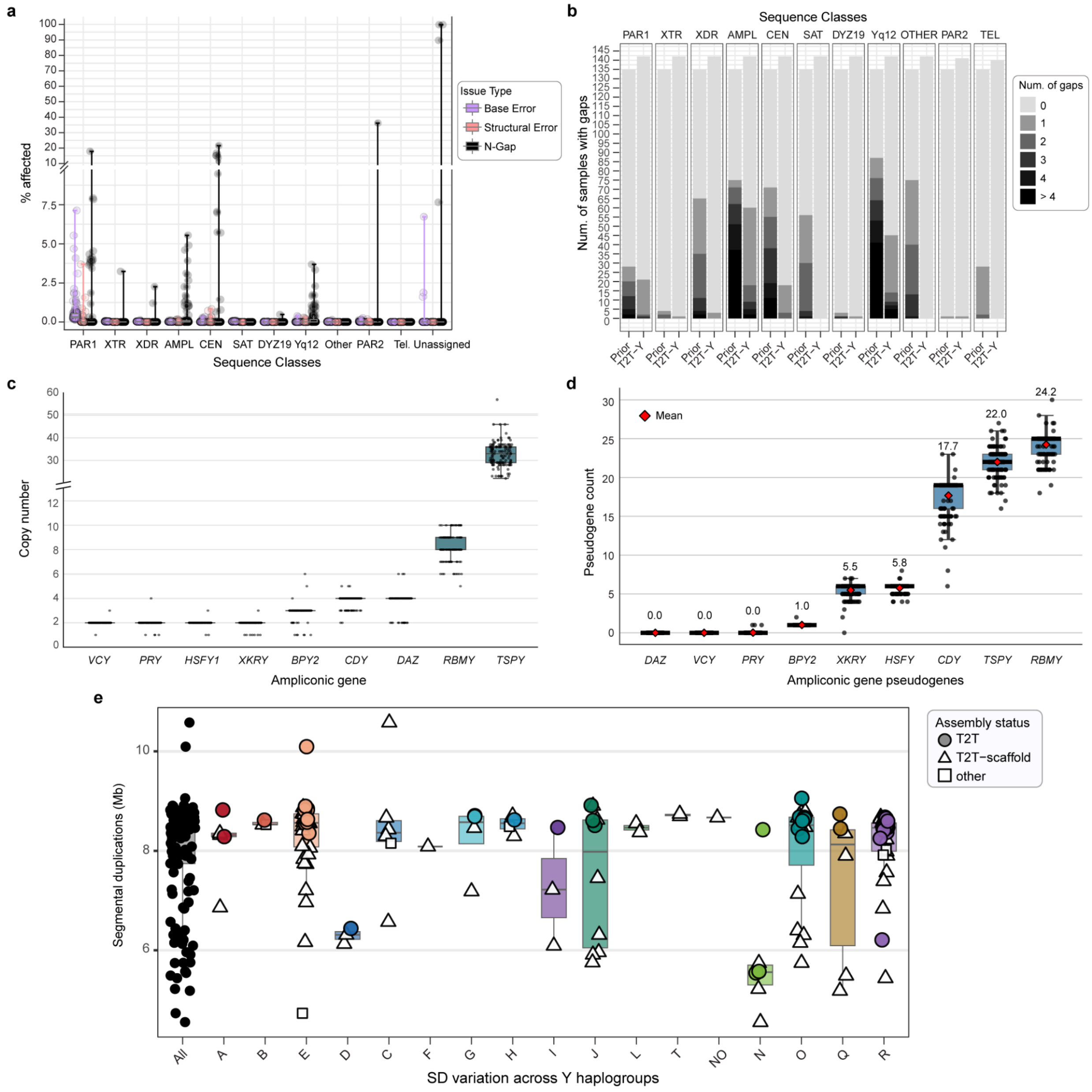


**Extended Data Figure 1. Assembly completeness and repeat-associated features across human Y chromosome assemblies. a, Percent annotated sequence classes affected by assembly issues**. Each circle represents a Y chromosome, with the % bases affected in each sequence class by an error type. Black lines indicate minimum and maximum values, whereas gray lines indicate median with interquartile range boxes (IQR, Q1-Q3). Whiskers indicate 1.5 x IQR. Sequences unable to assign a sequence class based on CHM13v2Y or GRCh38Y are labeled as "Unassigned". Locations of the error type on each Y is shown in **Supplementary Fig. 2**. **b, Number of samples found with the number of gaps in each sequence class**. Prior best assemblies (left) are compared to the Y chromosomes presented in this study (right). Current assemblies show less gaps, therefore more complete representation of the sequence classes, especially in the heterochromatin and its surrounding sequences (Other), ampliconic region (AMPL), centromere (CEN), and PAR1. **c-d, Liftoff- and pangenome graph-based gene and pseudogene counts for**

**Y ampliconic genes.** Boxplots showing the distribution of matched pseudogenes identified across Y chromosome assemblies (main contig only) for each ampliconic gene family. Only pseudogenes identified and classified (as a pseudogene) by both the Liftoff- and pangenome graph-based approaches were counted. Points indicate individual Y assemblies, boxes represent the interquartile range, the center line marks the median, and whiskers extend to 1.5x the interquartile range. Red diamonds indicate the mean matched pseudogene count for each gene family. Gene names are shown on the x-axis, with the mean value listed under each label. Pseudogene counts are strongly gene family-dependent, with the largest counts/variation in *TSPY*, *RBMY*, and *CDY*. **e, Intrachromosomal segmental duplication content.** Non-redundant intrachromosomal segmental duplication content across 140 open-access Y chromosomes, shown for all samples and grouped by major haplogroup. Boxplots indicate the median and interquartile range. Each point represents one assembly, and symbols indicate assembly status.

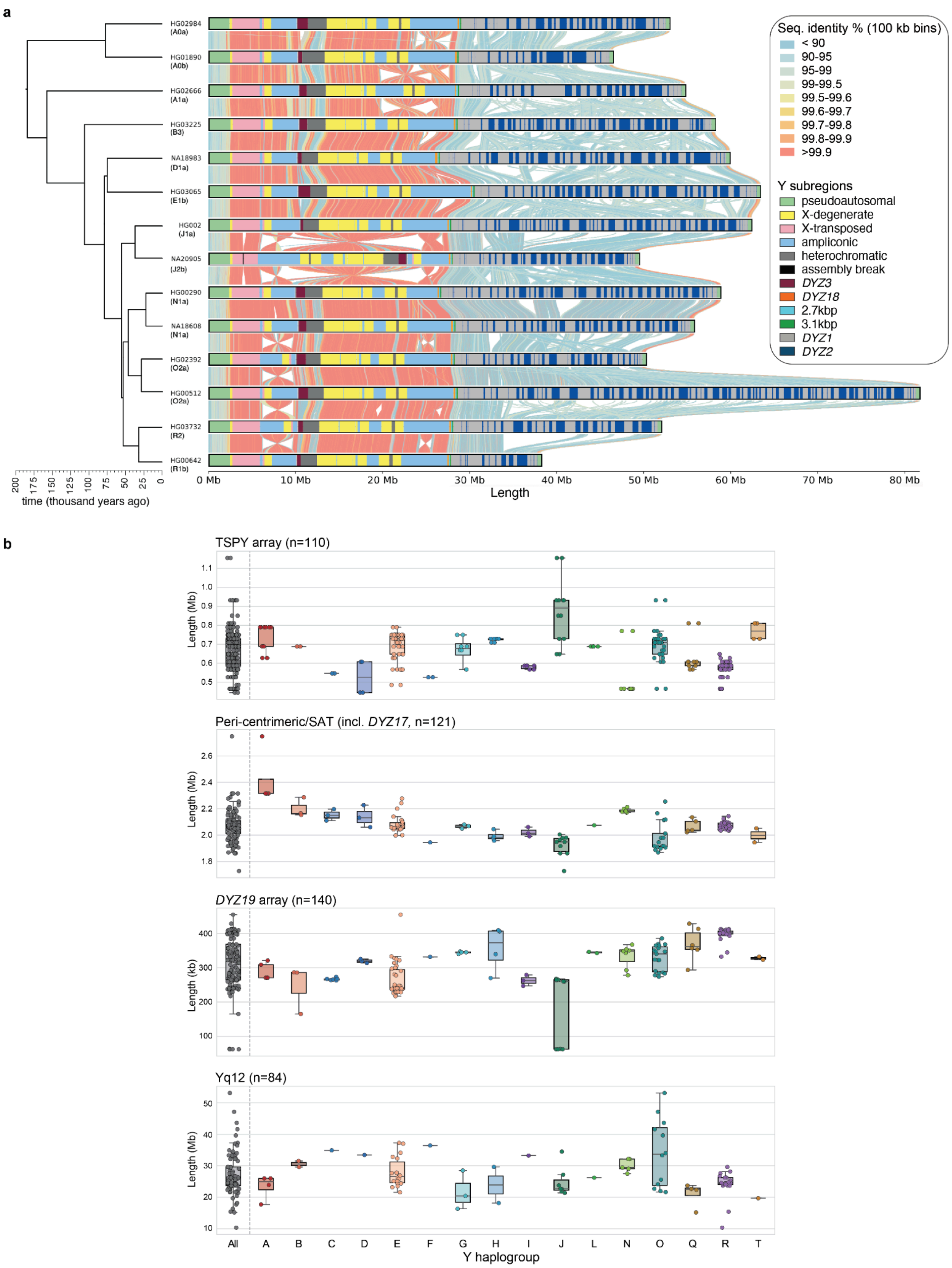


**Extended Data Figure 2. Structural and repeat-array variation across assembled human Y chromosomes. a,** Pairwise comparison of 14 assembled Y chromosomes illustrating extensive structural variation. Colored bars indicate Y-chromosomal sequence classes for each sample, and vertical lines between assemblies indicate sequence identity; a dated phylogeny is shown at left. **b,**

Length variation across four Y-chromosomal repeat arrays, shown for open-access and QC-passed samples (samples with N-gaps and structural error annotations in the analysed subregion were excluded) and grouped by major Y haplogroup. Boxplots indicate the median and interquartile range.

### Supplementary Figures

#### Y phylogeny

Contributing authors: Pille Hallast

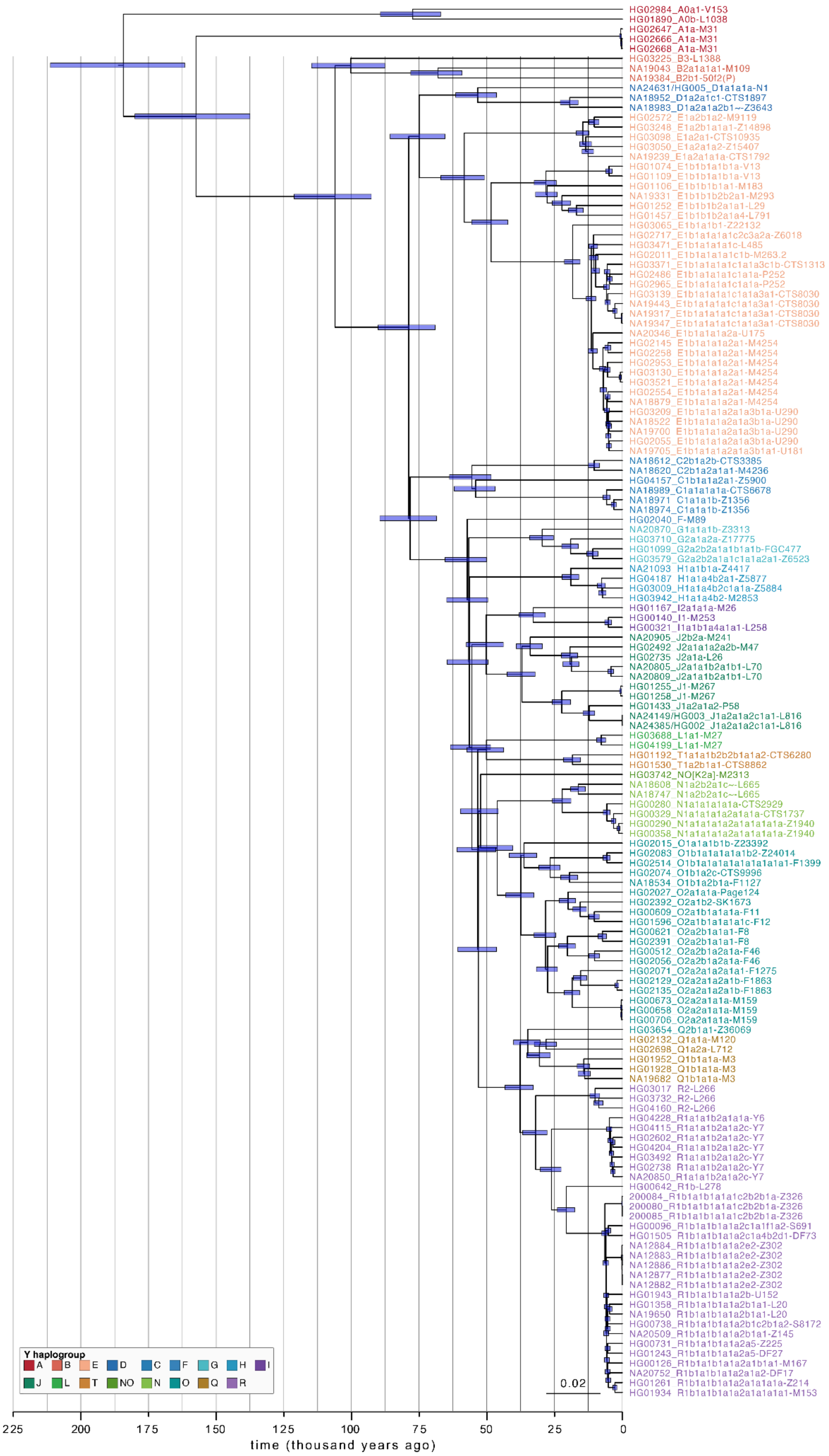


**Supplementary Figure 1. Phylogenetic relationships among the 142 assembled Y chromosomes.** The 95% HPD intervals for split-time estimates inferred by BEAST are shown as blue horizontal bars. Each sample ID is followed by the full Y-haplogroup label (according to ISOGG v15.73), and the terminal marker ID.

#### Assembly and QC

Contributing authors: Arang Rhie, Peter Ebert, Sergey Koren, Nancy Hansen, Prajna Hebbar, Jiadong Lin, Keisuke Oshima, Glennis Logsdon, Benedict Paten, Adam Phillippy

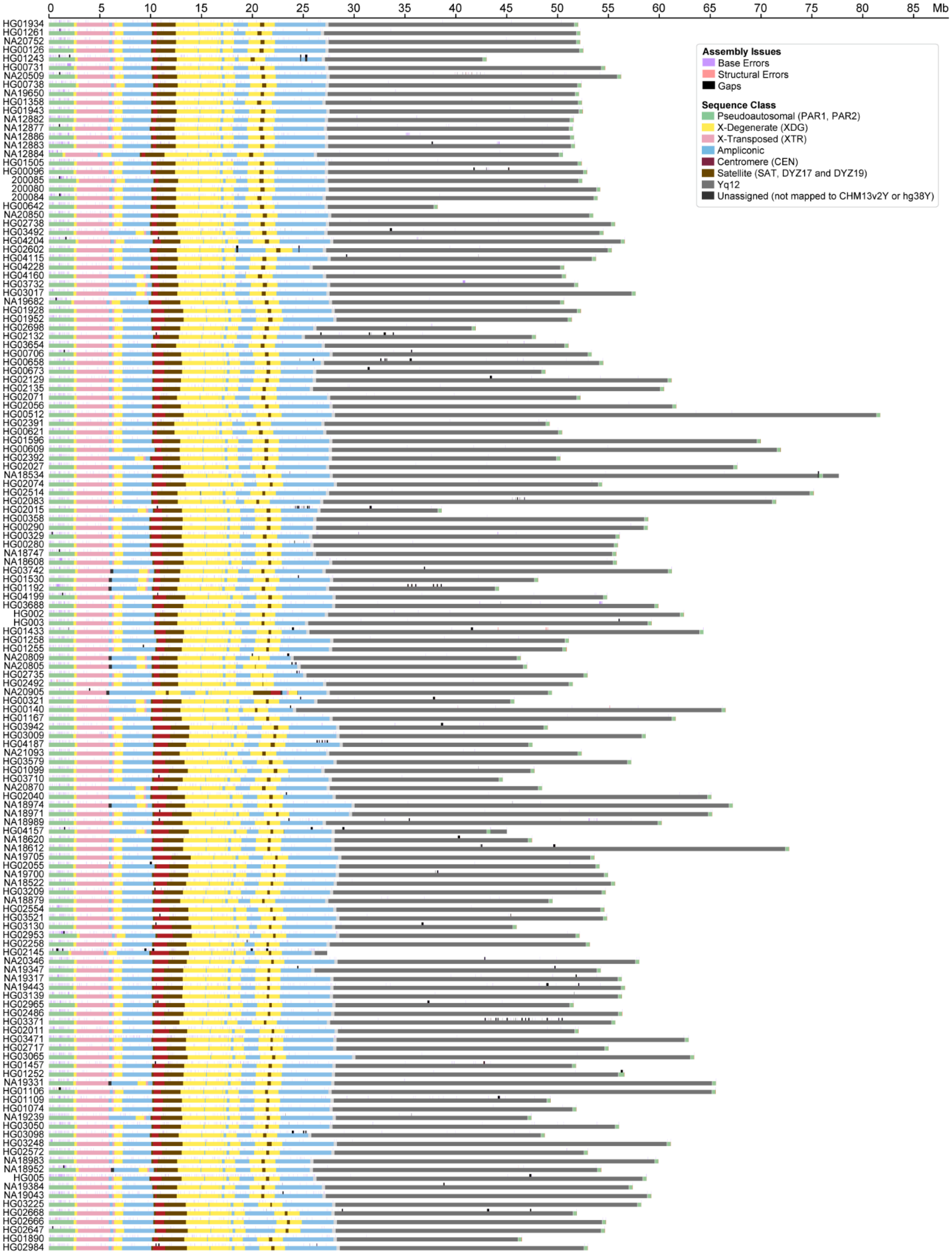


**Supplementary Figure 2. Sequence class annotation with assembly issues**. Assembly issues are shown on top of the ideogram. Percent of each assembly issue type is shown in Extended Data Fig. 1a.

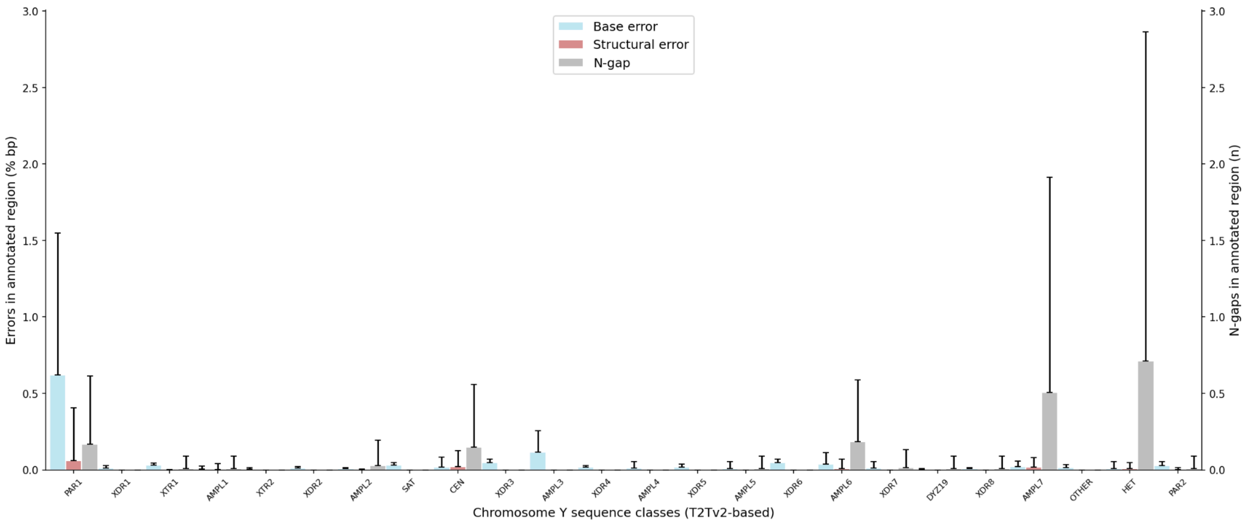


**Supplementary Figure 3. Percent annotated sequence classes affected by assembly issues.** The x-axis shows sequence classes, the left y-axis shows the mean fraction of bases in each annotated class affected by base-level errors (blue) or structural errors (red), and the right y-axis shows the mean number of unresolved gaps (N-gaps; grey) within each class. Error bars indicate one standard deviation across assemblies. Base-level and structural errors are low across most sequence classes, whereas unresolved gaps are concentrated in a subset of repetitive classes, including PAR1, centromere (CEN), ampliconic region with P1-P3 palindromes (AMPL7) and Yq12 (HET). Overall, these data show that assembly errors are generally rare across the Y chromosome but remain enriched in a small number of highly repetitive sequence classes.

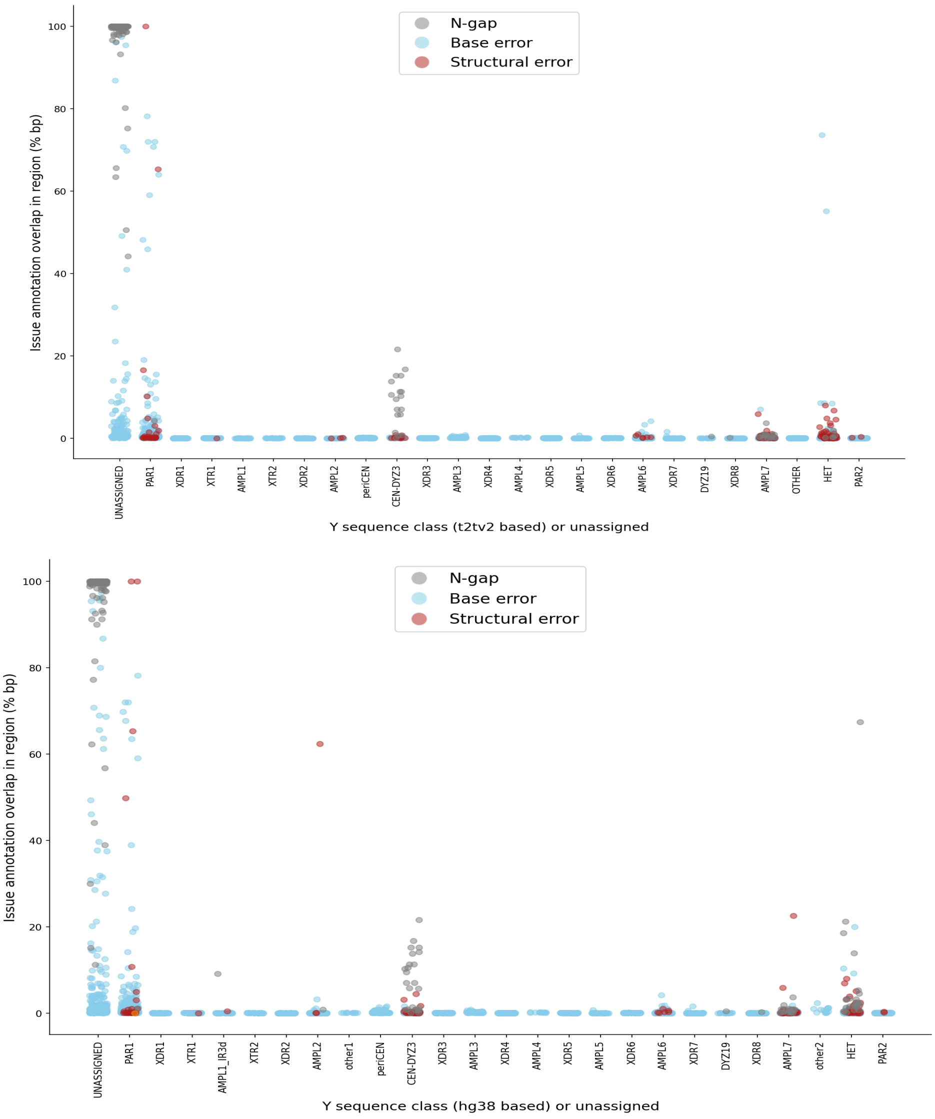


**Supplementary Figure 4. Percent annotated sequence classes affected by assembly issues by T2T-CHM13v2Y or GRCh38Y.** Scatter plot showing the proportion of bases in each Y-chromosome sequence class that overlap flagged assembly issues. The x-axis denotes sequence classes defined from the T2T-CHM13v2Y-based (top) or GRCh38Y-based (bottom) annotation, together with unassigned sequences. The y-axis shows the percentage of bases in each class affected by a given issue. Each point represents one assembly. Colors indicate unresolved gaps or N-bases (grey), base errors (blue), and structural errors (red). Most sequence classes show few affected sequences in most assemblies. Elevated issue rates are concentrated in certain classes, such as PAR1, centromere (CEN), ampliconic region with P1-P3 palindromes (AMPL7) and Yq12 (HET). This analysis was used to evaluate assembly quality and guide sample inclusion. Original data is available as **issues-pct-cov-by-region.pe.t2tv2** and **issues-pct-cov-by-region.pe.hg38**.

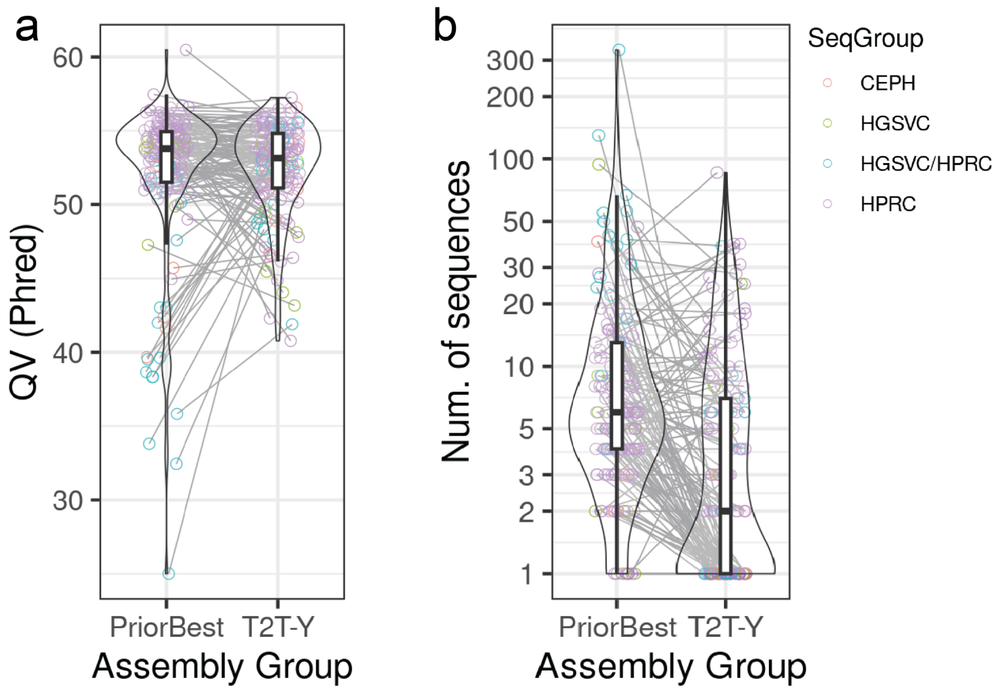


**Supplementary Figure 5. QV estimates and assembly fragmentation in current versus previous best Y-chromosome assemblies.** Comparison of **(a)** QV estimates and **(b)** numbers of assembled Y sequences in 136 matching samples for which previous Y assemblies were available (see **Suppl. Table 2** for details). Samples are colored by the data group, as defined in **Suppl. Table 1**. Lower numbers of assembled Y sequences indicate improved assembly contiguity.

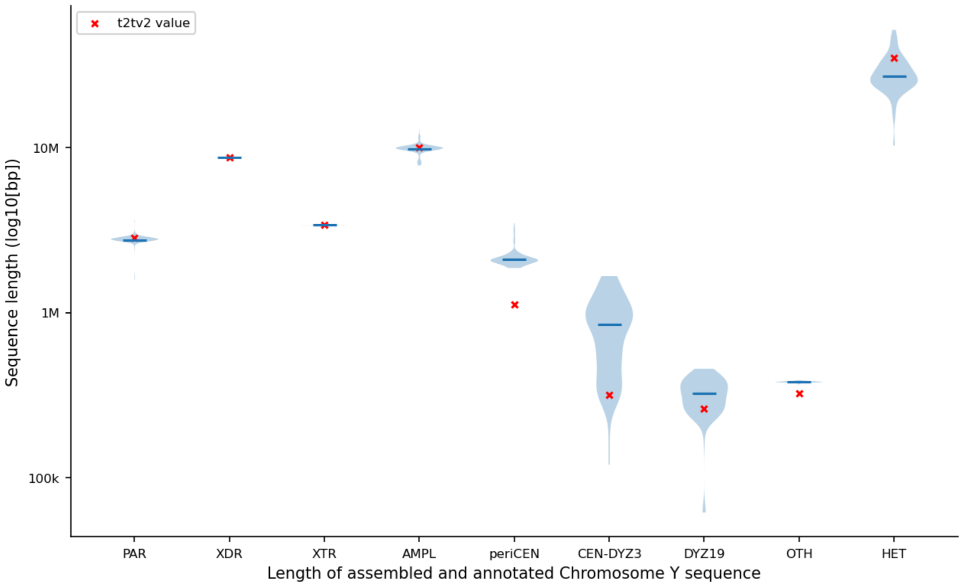


**Supplementary Figure 6. Sequence-length distributions across major T2T-CHM13v2 sequence class groups.** Violin plots showing the distribution of assembled and annotated sequence length across the 142 chromosome Y assemblies for each major sequence class group defined using the T2T-CHM13v2 annotation. The x-axis shows broad sequence class groups, and the y-axis shows the assembled sequence length on a log10 scale (bp). The blue violin plots show the observed distribution across assemblies, and the horizontal blue bars highlight the mediancentral value for each class. Red crosses mark the class length in the T2T-CHM13v2 reference assembly. Sequence lengths are tightly constrained for PAR, XDR, XTR and AMPL sequence, but show substantially greater variability in periCEN, CEN-DYZ3, DYZ19 and Yq12 (HET), indicating that the largest differences among assembled Y chromosomes are concentrated in centromeric and heterochromatic compartments. Overall, these data show that some Y-chromosome sequence classes are highly stable in size, whereas centromeric and heterochromatic classes contribute disproportionately to inter-assembly length variation.

### Gene content

Contributing authors: Mark Loftus, Feyza Yilmaz, Prajna Hebbar, Benedict Paten, Charles Lee

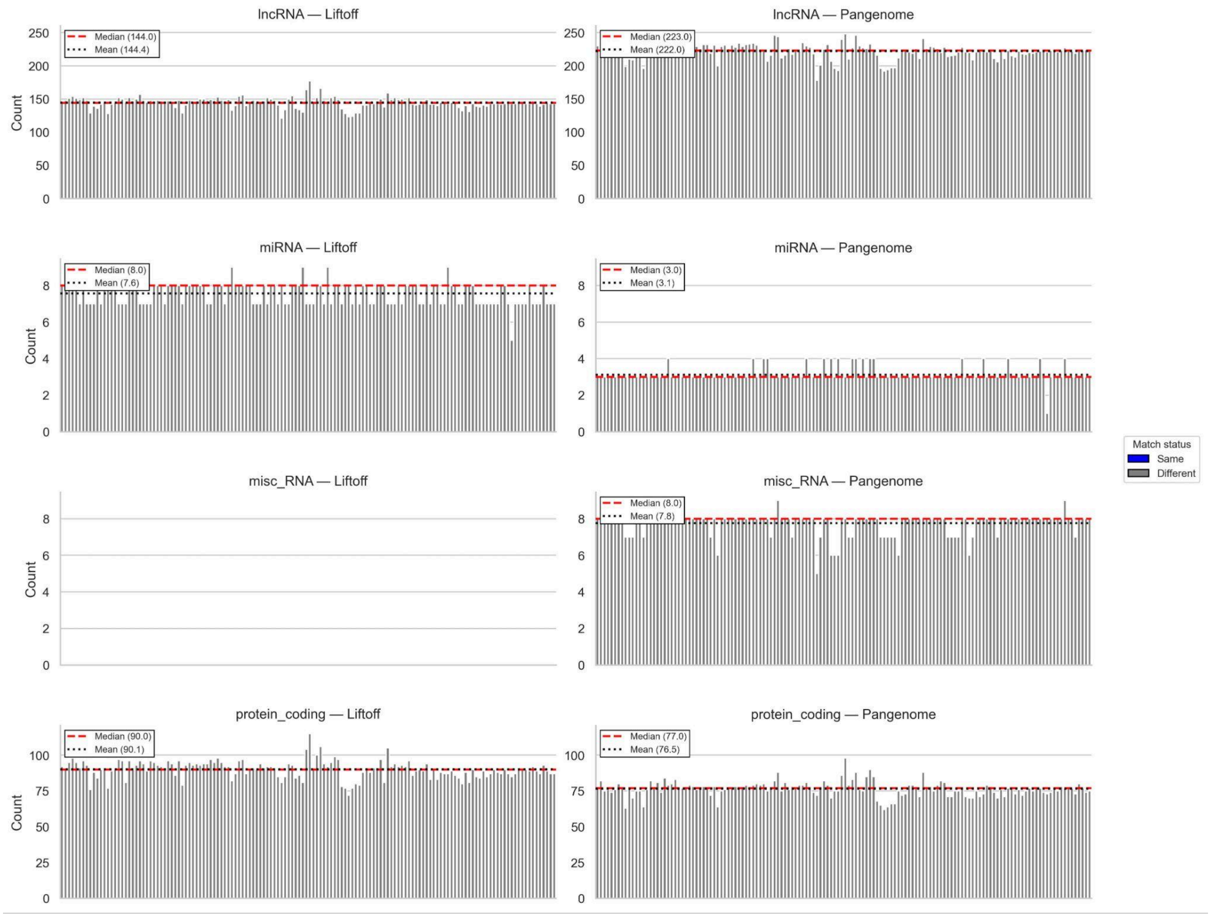

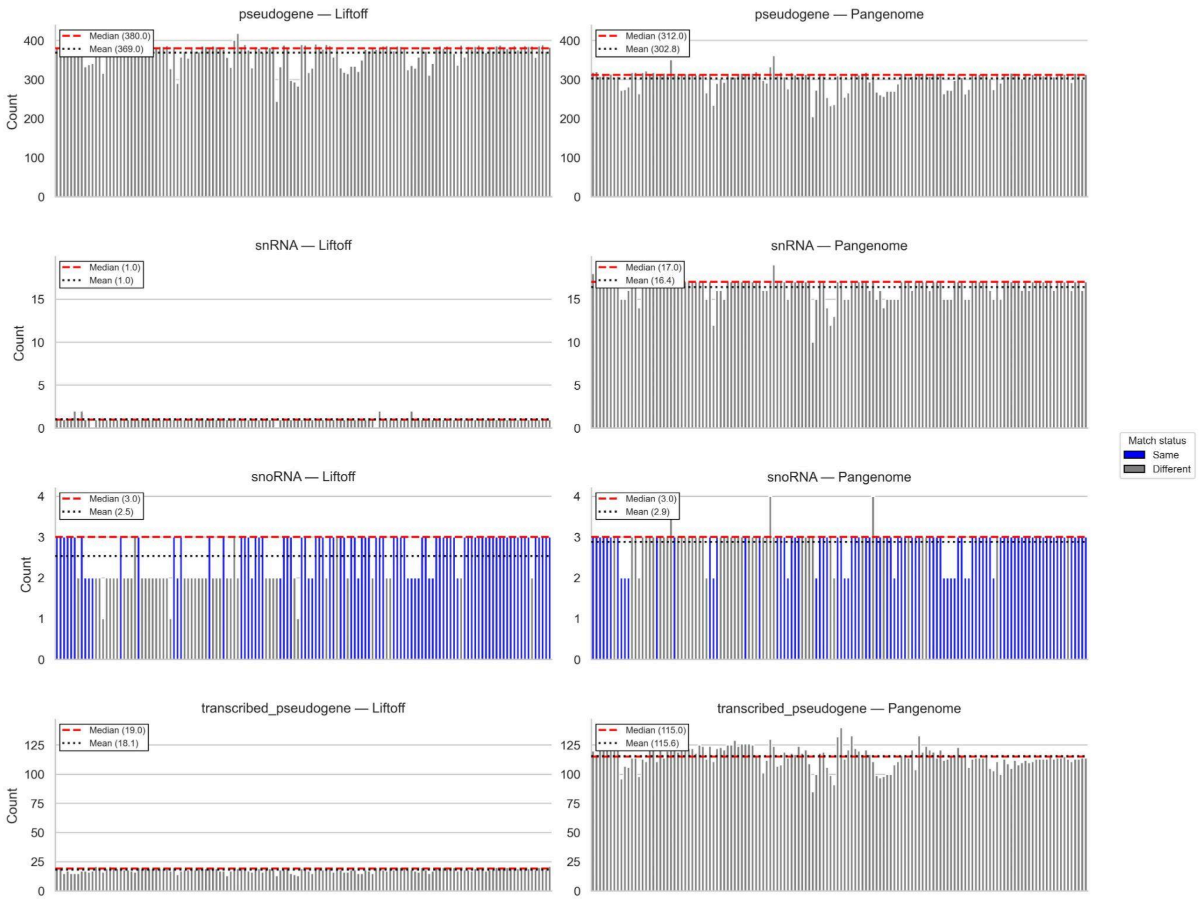


**Supplementary Figure 7. Gene biotype counts are largely consistent between Liftoff and pangenome-graph annotation, with discrepancies concentrated in select biotypes.** Bar plots show per sample gene counts for major gene biotypes inferred using two annotation approaches: Liftoff-based lift-over (***left***) and pangenome graph (***right***) annotation. Panels are stratified by biotype (protein-coding, lncRNA, miRNA, misc_RNA, pseudogene, transcribed pseudogene, snoRNA, snRNA), and samples are arranged by phylogeny from left to right (starting with haplogroup A lineages). Bar color indicates whether the two pipelines report the same biotype-specific count for a given sample (***blue***) or different counts (***gray***). Red dashed and black dotted lines indicate the median and mean counts, respectively, with values annotated in each panel.

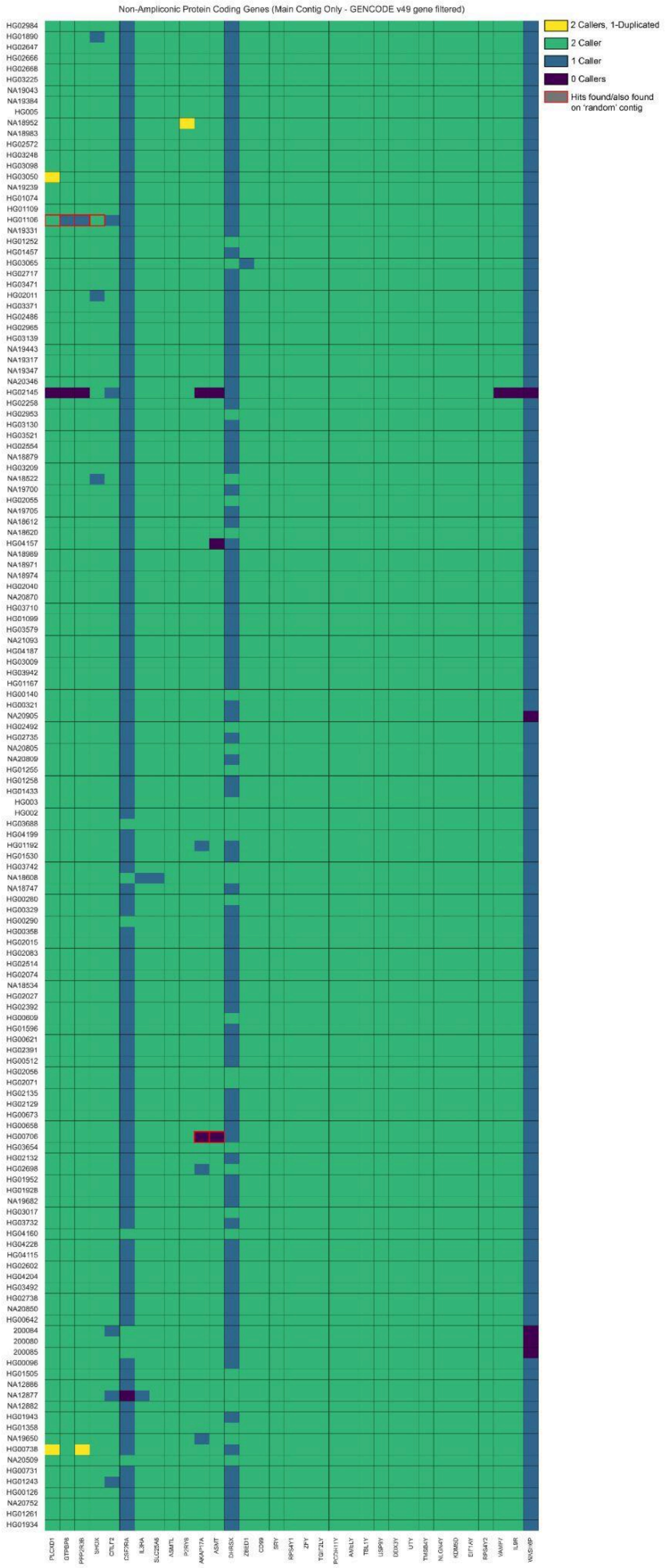
Non-Ampliconic Protein Coding Genes (Main Contig Only - GENCODE v49 gene filtered)
2 Callers, 1-Duplicated
2 Caller
1 Caller
0 Callers
Hits found/also found on 'random' contig

**Supplementary Figure 8. Concordance of non-ampliconic, protein coding gene annotations across Y chromosome assemblies.** Heatmap summarizing gene calls from Liftoff and pangenome graph-based annotation pipelines across non-ampliconic Y-chromosome protein coding genes. Rows correspond to samples ordered by phylogeny with haplogroup A lineages at top. Columns correspond to genes, ordered by genomic position, from PAR1 on the left to PAR2 on the right. Cell color indicates concordance between pipelines: ***yellow*** denotes genes identified by both pipelines where one caller annotated the gene as duplicated; ***green*** indicates genes identified by both pipelines as single copy; ***blue*** indicates genes identified by only one pipeline; and ***purple*** indicates genes not identified by either pipeline. A ***red outline*** highlights genes also detected in non-primary ("random") assembly contigs. All single-copy male-specific Y-chromosome genes (*SRY* → *RPS4Y2*) were identified exactly once in the assemblies by both annotation pipelines.

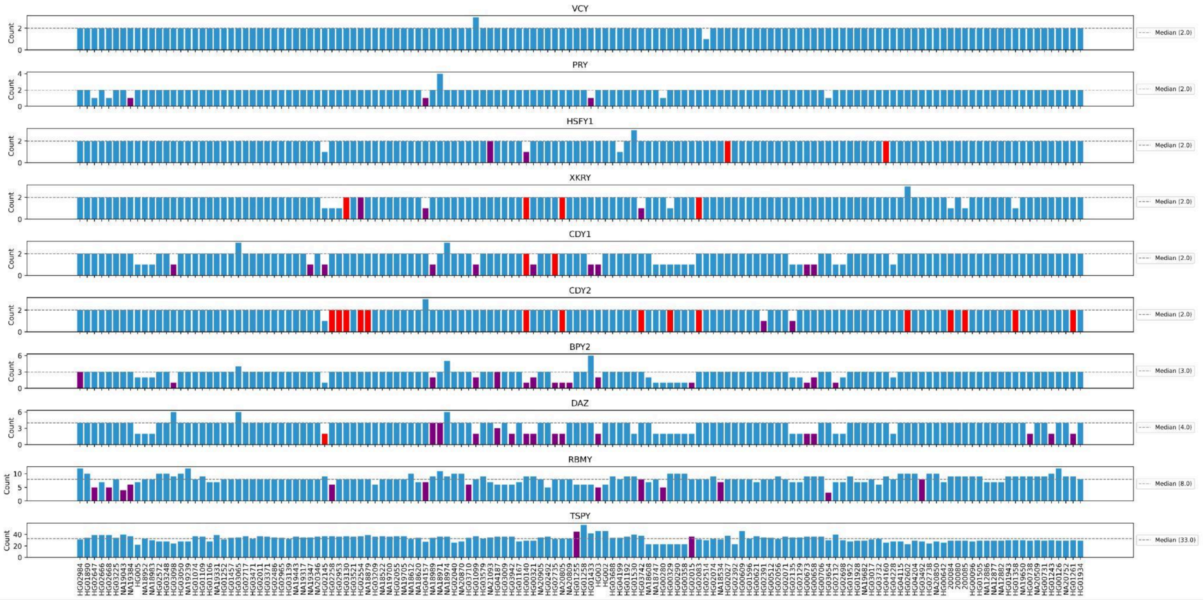


**Supplementary Figure 9. Sample specific copy number variation in Y chromosome ampliconic gene families.** Bar plots show total copy number for nine Y-chromosome ampliconic gene families (with *CDY* reported separately as *CDY1* and *CDY2*) across 140 open-access samples. Gene families are ordered from top to bottom by increasing copy-number variability (standard deviation), from *VCY* (lowest variability) to *TSPY* (highest variability). For each gene family, the median copy number across samples is indicated by a dashed reference line (median value annotated). Bar colors denote gene assembly localization: ***blue***, all copies on the main contig; ***red***, copies detected only on non-main contigs; ***purple***, copies detected on both main and non-main contigs. Hence, ampliconic gene families vary widely in how many copies they have across individuals - some are fairly stable (e.g., *VCY*), while others (e.g., *TSPY*) show large, sample-to-sample copy number changes.

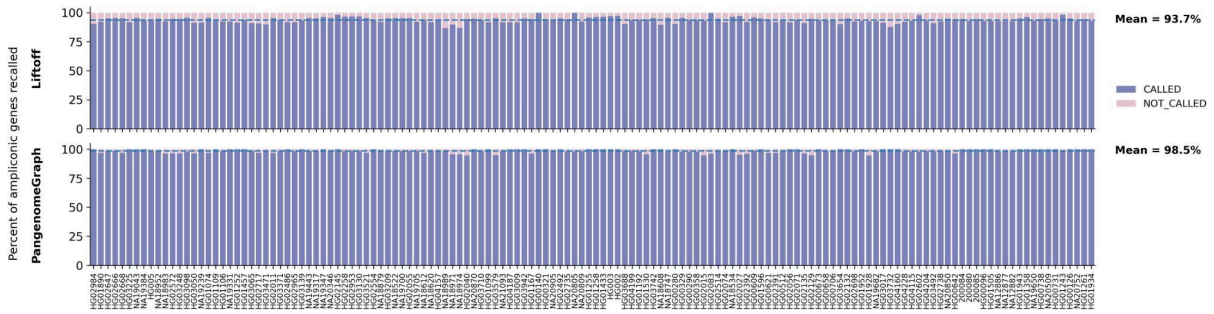


**Supplementary Figure 10. Recovery of RepeatMasker-stitched ampliconic genes by Liftoff and pangenome graph annotation.** Stacked bar plots show per-sample recovery (recall) of Y-chromosome ampliconic genes (full exon copy genes only) recalled by Liftoff (***top panel***) or the pangenome graph pipeline (***bottom panel***), measured relative to a RepeatMasker-based ampliconic gene stitching annotation used as the baseline reference. For each sample, the colored fraction indicates the proportion of baseline ampliconic genes on the primary Y contig that are also identified by the evaluated pipeline ("called", purple), with the remainder not covered ("not called", pink). Samples are ordered phylogenetically, beginning with haplogroup A on the left. Mean recall across samples is higher for the pangenome graph-based pipeline than for Liftoff (Liftoff mean: 93.7%; pangenome graph mean: 98.5%).

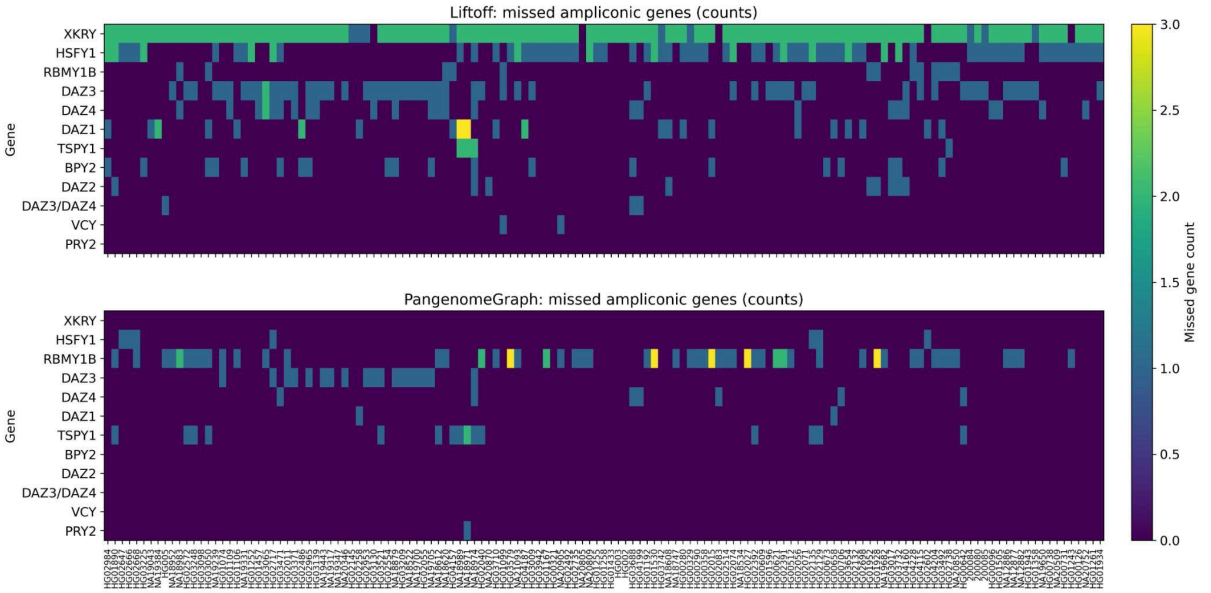


**Supplementary Figure 11. Pipeline specific patterns of missed ampliconic gene copies relative to a RepeatMasker baseline.** Heatmaps show, for each sample (columns; ordered phylogenetically) and ampliconic gene family (rows), the number of baseline ampliconic gene copies on the primary Y contig that were not recovered ("missed") by Liftoff (***top panel***) or by the pangenome graph pipeline (***bottom panel***) where the baseline is a RepeatMasker-based ampliconic gene stitching annotation. Color indicates the missed-copy gene count per sample. The results are only for the genes missed on the main contig, and reveal strong caller-specific biases in the genes that were missed. Liftoff failed to recover a broader range of ampliconic genes overall, with most omissions occurring within the *DAZ* and *XKRY* gene families. In contrast, the pangenome graph–based pipeline mostly failed to identify *TSPY* and *RBMY* genes.

### PAR regions

Contributing authors: Kwondo Kim, Charles Lee

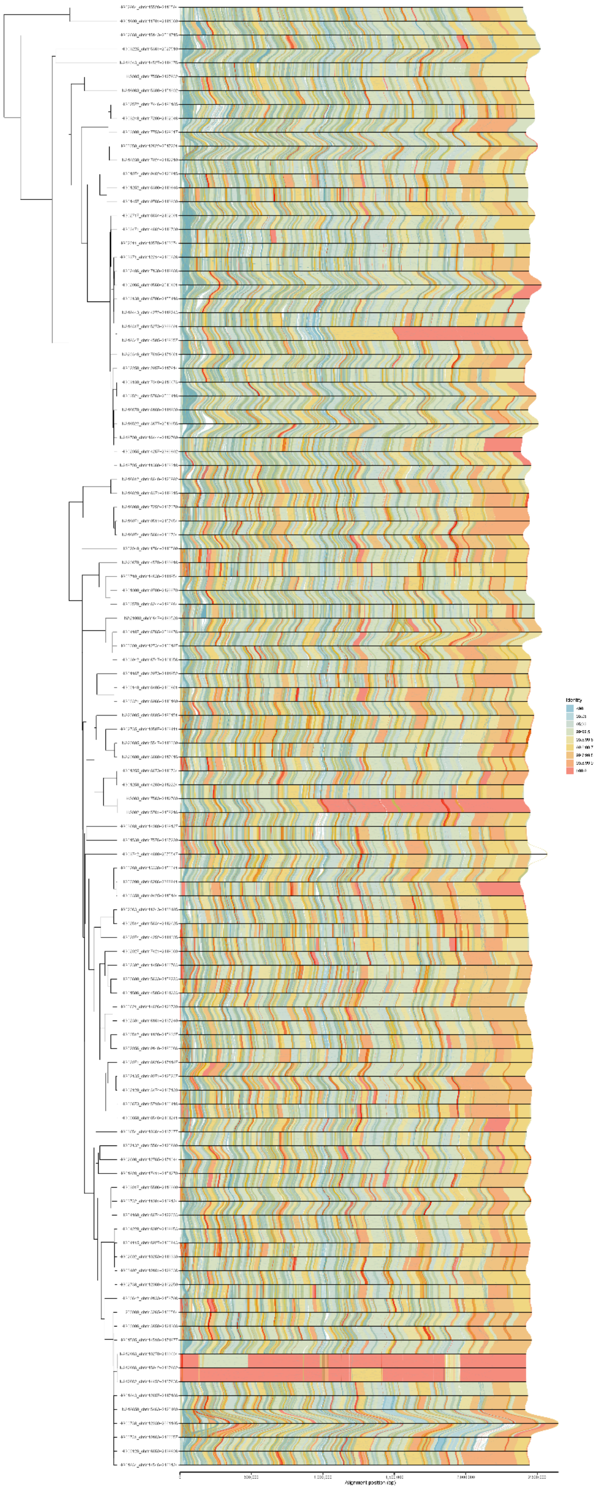

**Supplementary Figure 12. Sequence alignments of PAR1.** The alignments between the sequences, without structural errors ("ERRSTRCT") or gaps ("NGAP"), were shown in phylogenetic order. The color code represents sequence identity in each alignment block. The coordinates on the sample label indicate the genomic position of the PAR1 sequence shown here, relative to the main Y contig.

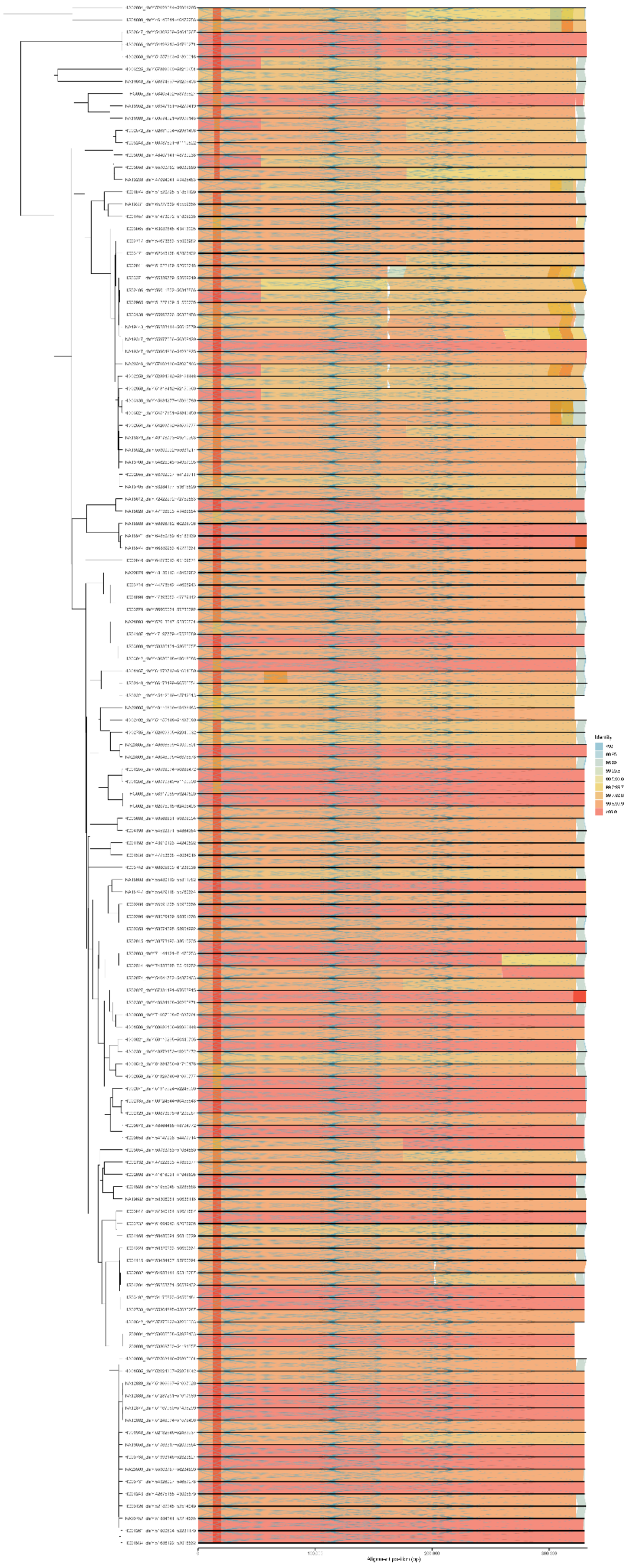

**Supplementary Figure 13. Sequence alignments of PAR2.** The alignments between the sequences, without structural errors (“ERRSTRCT”) or gaps (“NGAP”), were shown in phylogenetic order. The color code represents sequence identity in each alignment block. The coordinates on the sample label indicate the genomic position of the PAR2 sequence shown here, relative to the main Y contig.

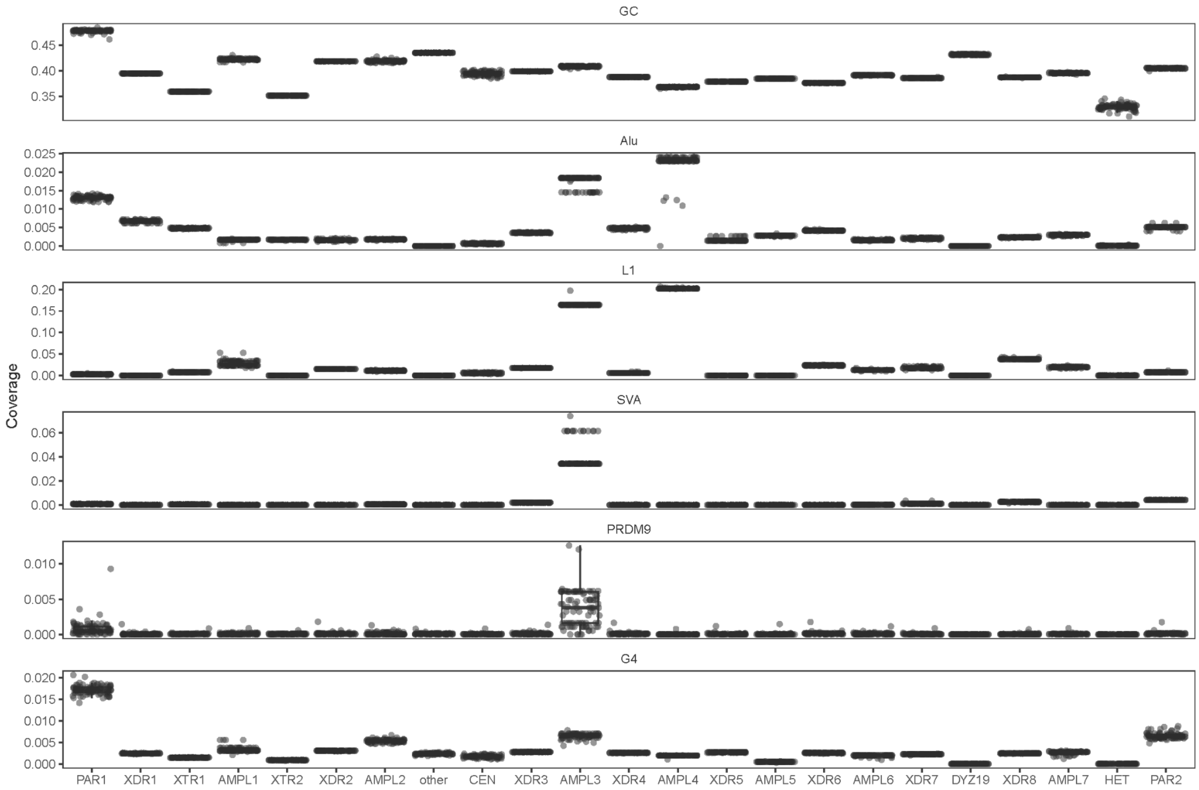


**Supplementary Figure 14. Recombination-associated sequence features of pseudoautosomal regions compared with other regions of the Y chromosome.** Each dot represents the coverage of the indicated sequence feature within a given genomic region for an individual.

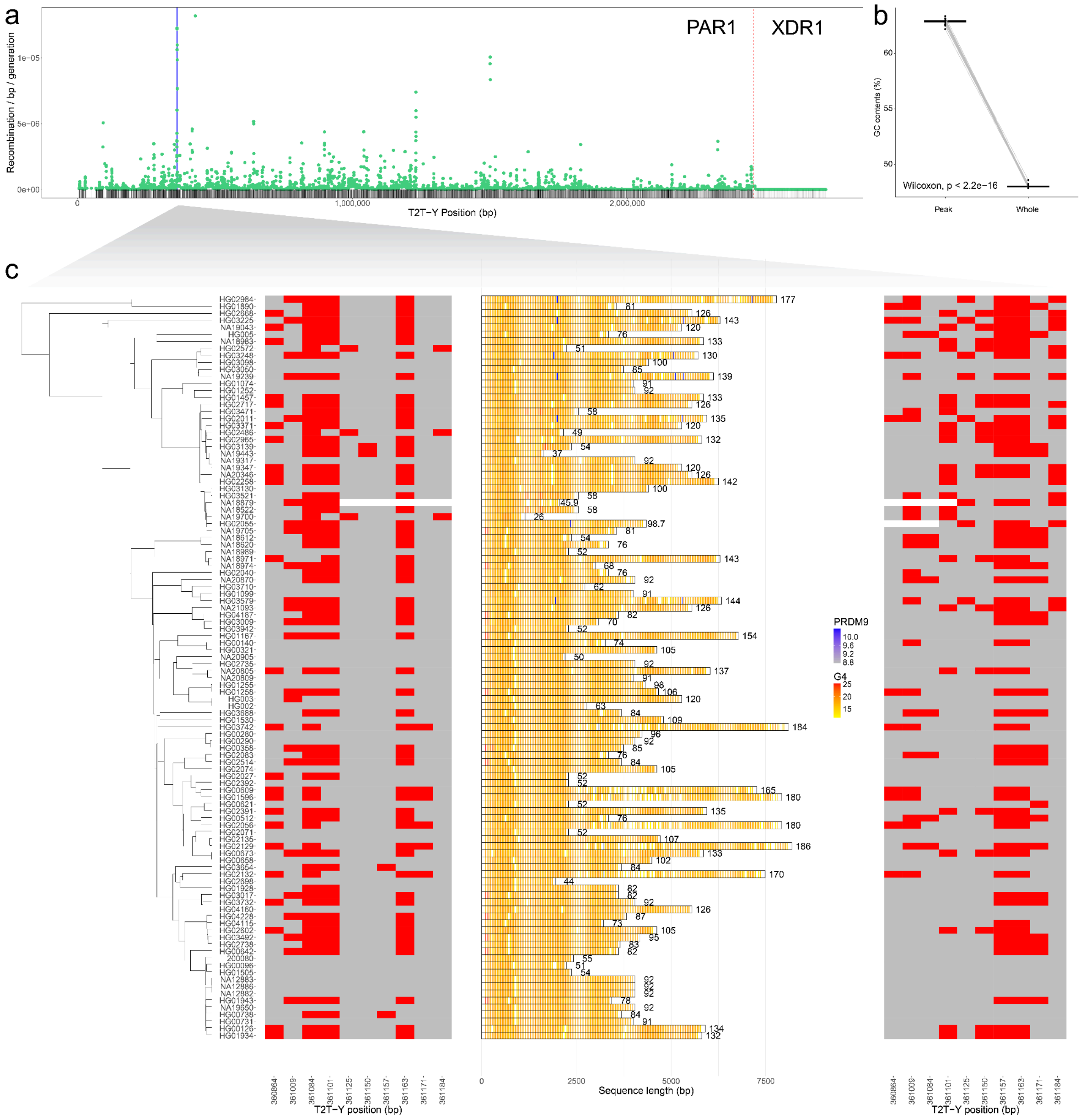


**Supplementary Figure 15. Tandem repeats at the PAR1 recombination hotspot. (a)** Recombination map of pseudoautosomal region 1 (PAR1) inferred from linkage disequilibrium. Related individuals (NA12883 and NA12886) were excluded from the analysis. The solid blue bar denotes the tandem repeat locus shown in panel (c), and the dashed red line marks the PAR1 boundary. **(b)** Comparison of GC content between the recombination hotspot ("Peak") and the entire PAR1 ("Whole"). GC content is significantly higher in "Peak" than in "Whole" with Wilcoxon signed-rank test p-value < 2.20 x 10-16. Related individuals (NA12883 and NA12886) were excluded from the analysis. **(c)** Tandem repeats identified at the recombination hotspot (HG002_chrY:361,257-364,030). The center panel shows the length of tandem repeats across samples with the location of possible G-quadroduplex (G4) and PRDM9 binding motifs. PRDM9 binding motif scores are scaled from grey (low) to blue (high), and G4 scores are scaled from yellow (low) to red (high). Numbers at the tips of the bar plots indicate the copy number of the repeat unit. Left and right panels show the ten 5′ and 3′ flanking SNVs, respectively. Reference and alternative alleles based on HG002 PAR1 sequence are shown in grey and red, respectively. Missing allele is shown in white.

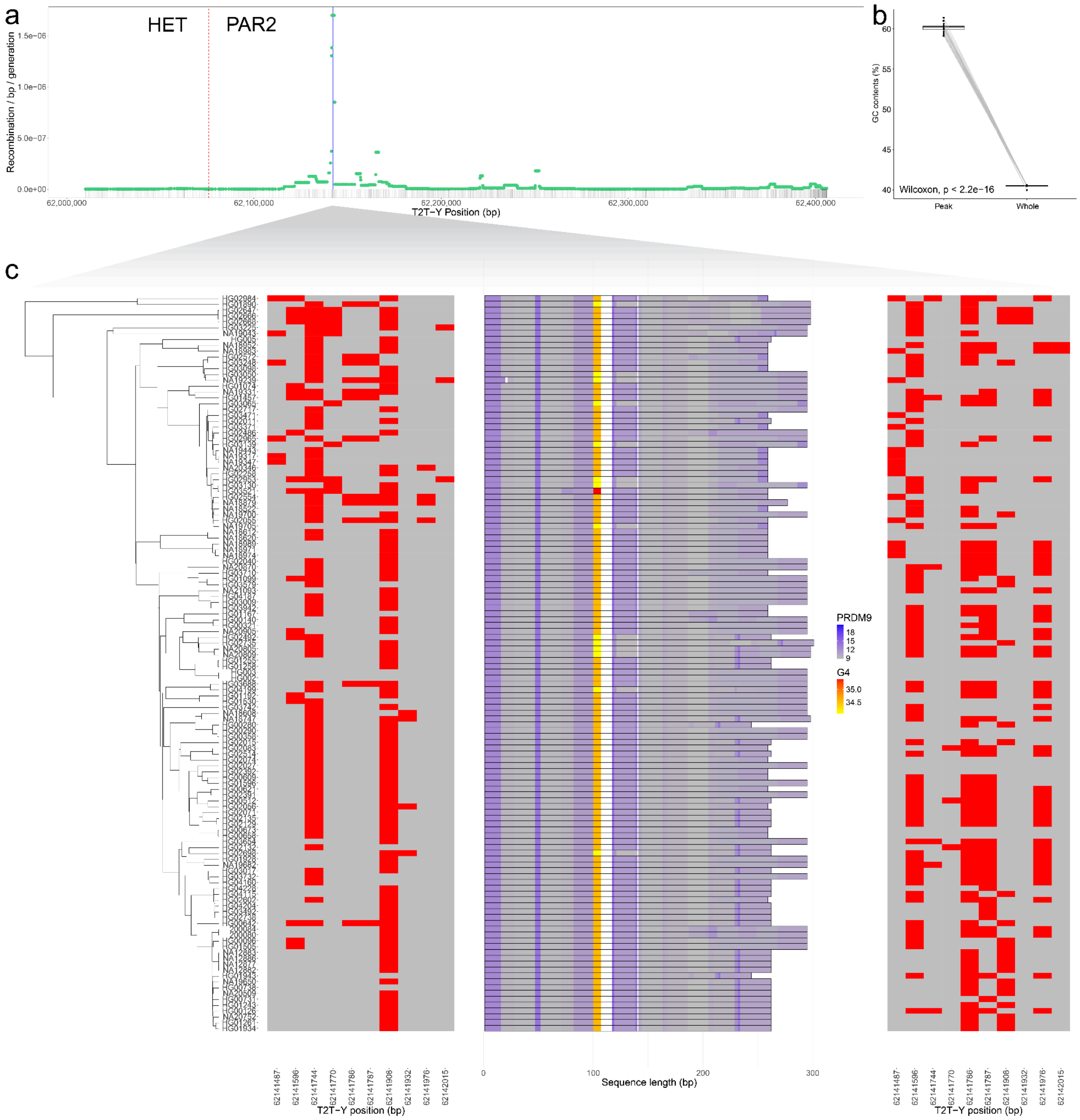


**Supplementary Figure 16. GA-rich tract at the PAR2 recombination hotspot. (a)** Recombination map of pseudoautosomal region 2 (PAR2) inferred from linkage disequilibrium. Related individuals (200084, NA12877, NA12882 and NA12883) were excluded from the analysis. The solid blue bar denotes the GA-rich tract locus shown in panel (c), and the dashed red line marks the PAR2 boundary. **(b)** Comparison of GC content between the recombination hotspot ("Peak") and the entire PAR2 ("Whole"). GC content is significantly higher in "Peak" than in "Whole" with Wilcoxon signed-rank test p-value < 2.20 x 10-16. Related individuals (200084, NA12877, NA12882 and NA12883) were excluded from the analysis. **(c)** GA-rich tract identified at the recombination hotspot (HG002_chrY:62,142,043-62,142,337). The center panel shows the length of GA-rich tracts across samples with the location of possible G-quadroduplex (G4) and PRDM9 binding motifs. PRDM9 binding motif scores are scaled from grey (low) to blue (high), and G4 scores are scaled from yellow (low) to red (high). Left and right panels show the ten 5′ and 3′ flanking SNVs, respectively.

Reference and alternative alleles based on HG002 PAR2 sequence are shown in grey and red, respectively. Missing allele is shown in white.

### Segmental duplications and ampliconic regions (incl. palindromes)

Contributing authors: DongAhn Yoo, Evan Eichler

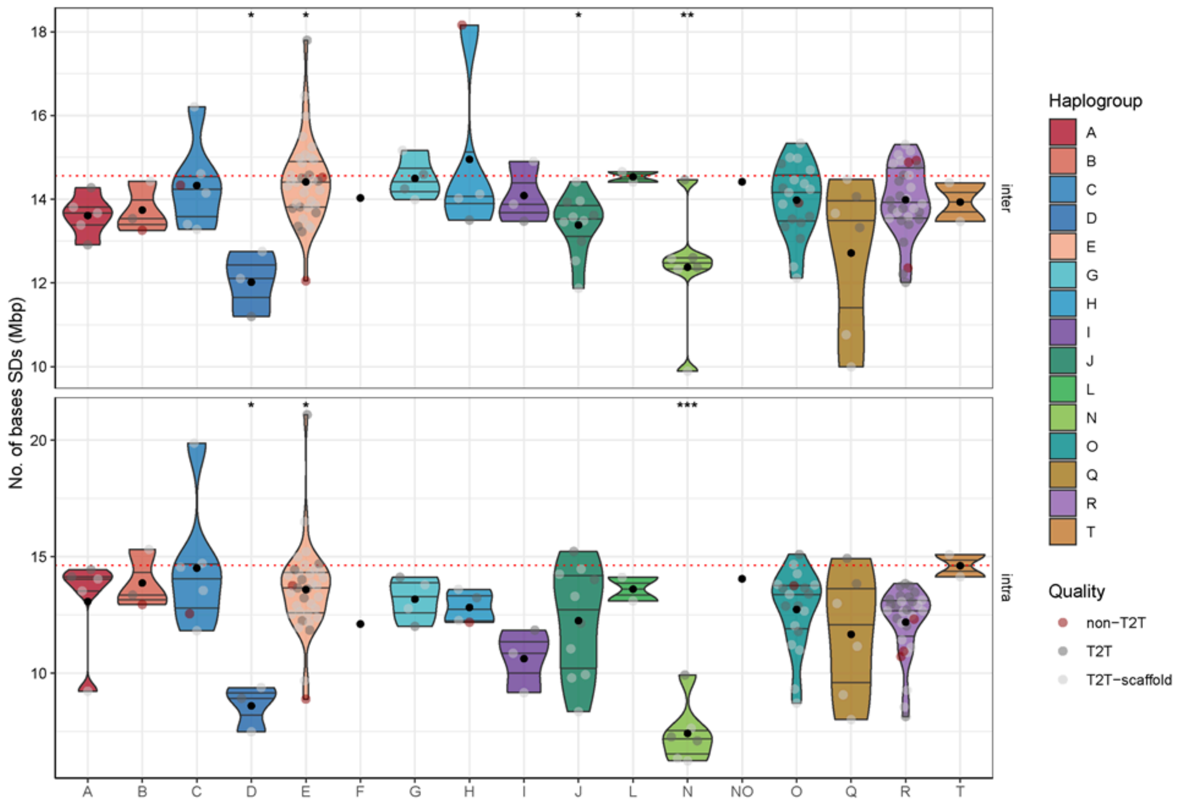


**Supplementary Figure 17. Quantification of SDs per haplogroup.** The number of redundant bases of duplicated sequences in chromosome Y are summarized for inter- and intrachromosomal SDs on the top and bottom, respectively. For each haplogroup, the two-tailed Wilcox test against the rest of the groups are indicated on the top (*: p<0.05, **: <0.01, and *** p<0.001 & adj.p<0.05). Individual dots indicate samples, and color of the dots represents the completeness of chromosome Y assembly. The horizontal dotted line in red indicates the amount of SD in T2T-CHM13v2.

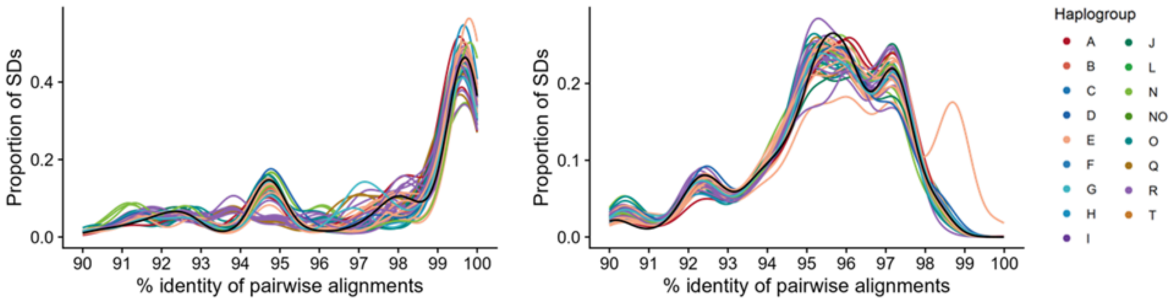


**Supplementary Figure 18. Distribution of sequence identity of duplicated sequences.** Sequence identity distributions are only shown for 46 T2T-level Y chromosome assemblies. Individual line indicates a sample with colors representing the haplogroup. Indicated by black is the distribution observed for T2T-CHM13v2.

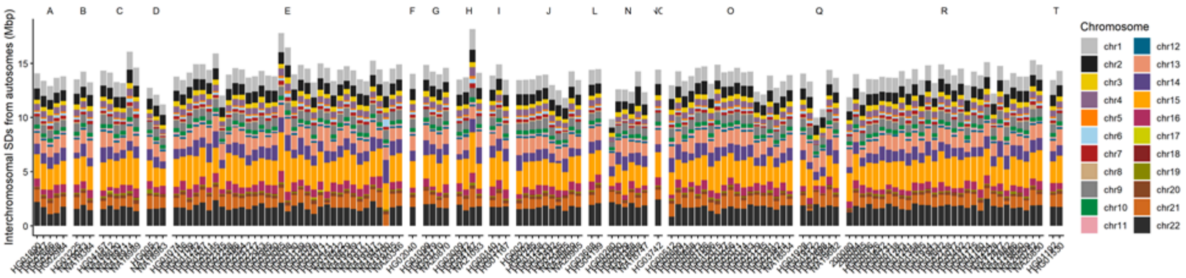


**Supplementary Figure 19. Interchromosomal SD content.** Chromosome Y SDs pairing with other chromosomes are quantified with colors of bars representing different chromosomes. On the top, a haplogroup of each population is indicated.

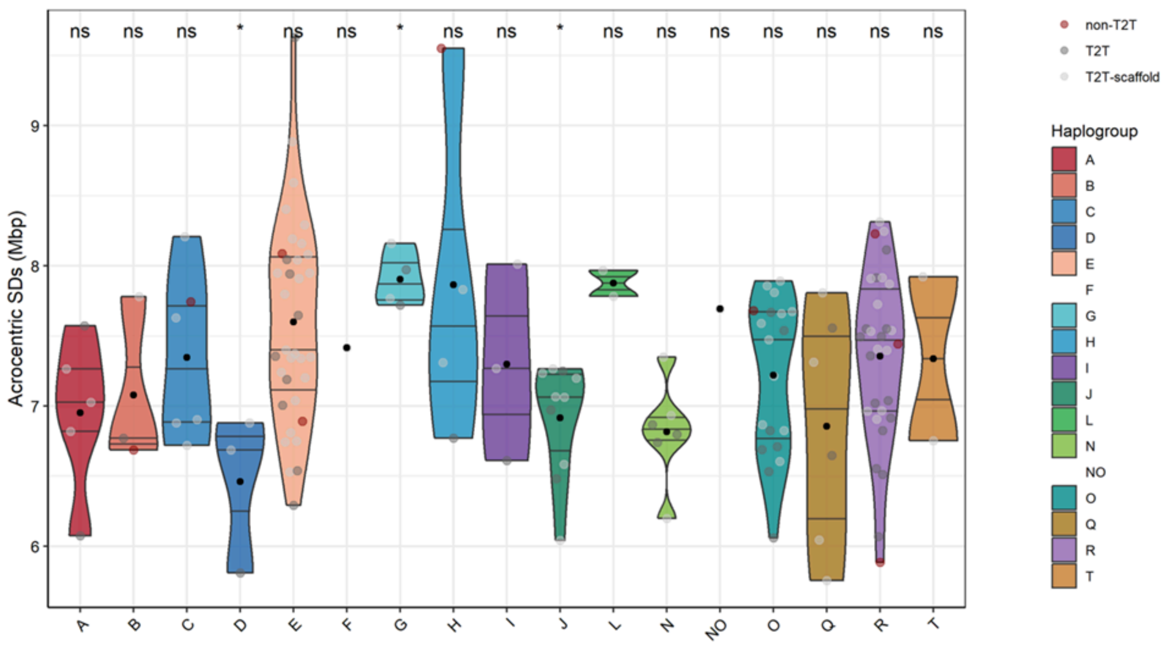


**Supplementary Figure 20. Quantification of SDs paired with acrocentric chromosomes.** Per each haplogroup, acrocentric SDs are quantified. Each point indicates a sample, with the color indicating the completeness of the chromosome Y assembly. Indicated on the top of the distribution, is the significance of Wilcox two-sided test; *: p<0.05 and ns: non-significant.

### Mobile Element Insertions

Contributing authors: Mark Loftus and Miriam K. Konkel

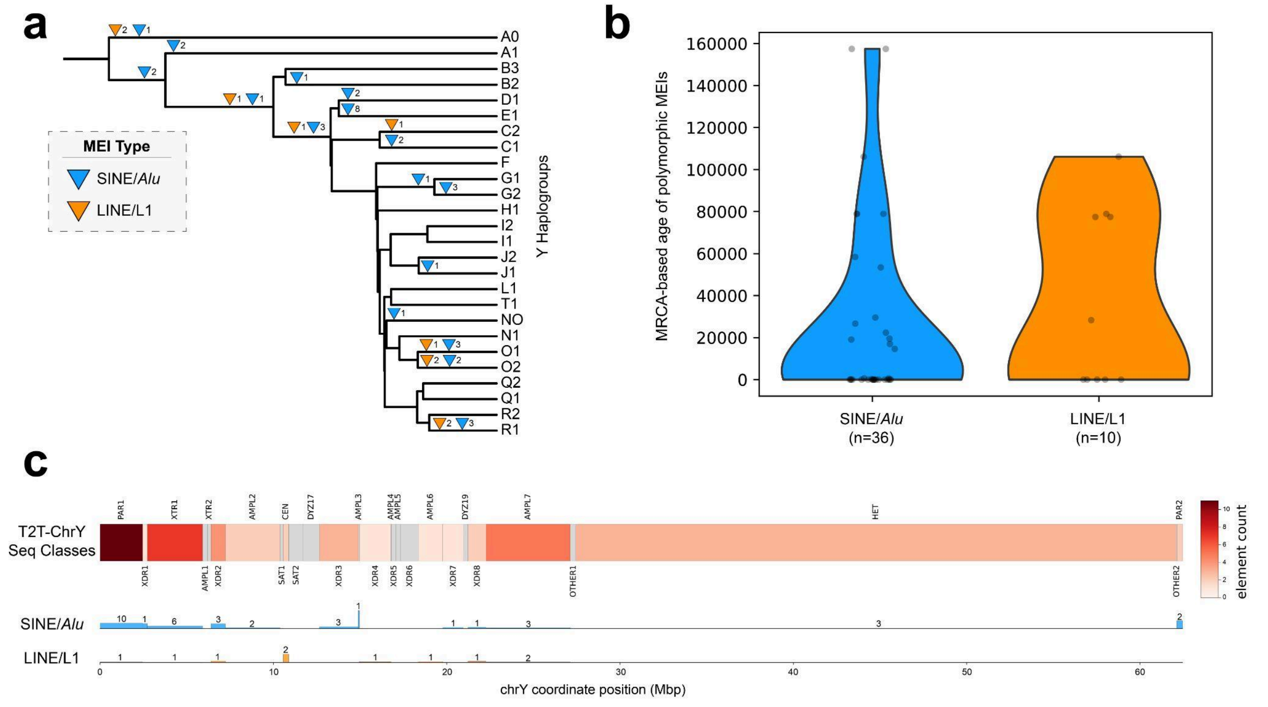


**Supplementary Figure 21. Polymorphic mobile element insertions across 140 Y chromosomes. (a)** Diagram of a phylogenetic tree of broad Y haplogroups showing the inferred/general placement of polymorphic mobile element insertions (MEIs). ***Triangles*** indicate insertion events, with blue representing SINE/*Alu* elements and orange representing LINE/L1 elements. Numbers adjacent to triangles denote the number of independent insertions assigned to each branch. Young polymorphic MEIs are sparsely distributed across the tree. **(b)** Violin plots showing the distribution of MRCA-based ages for polymorphic MEIs by element type. SINE/*Alu* insertions (n=36) are generally more numerous. ***Points*** represent individual insertions. **(c)** Relative genomic distribution of MEIs across the T2T-CHM13v2 Y chromosome, shown alongside sequence classes. SINE/*Alu* and LINE/L1 insertions are plotted by sequence class bin. The size of element density bar plots (beneath the T2T-CHM13v2 Y) represent counts per million base pairs with total counts shown above each bar. Polymorphic MEIs are rare across the Y chromosome overall, with limited accumulation even within large repetitive regions such as the Yq12 heterochromatin, while LINE/L1 elements show relative enrichment in specific regions (i.e., centromere) compared to *Alu* elements.

### Functional impact of SVs

Contributing authors: Yunzhe Jiang, Matthew Jensen, Mark Gerstein

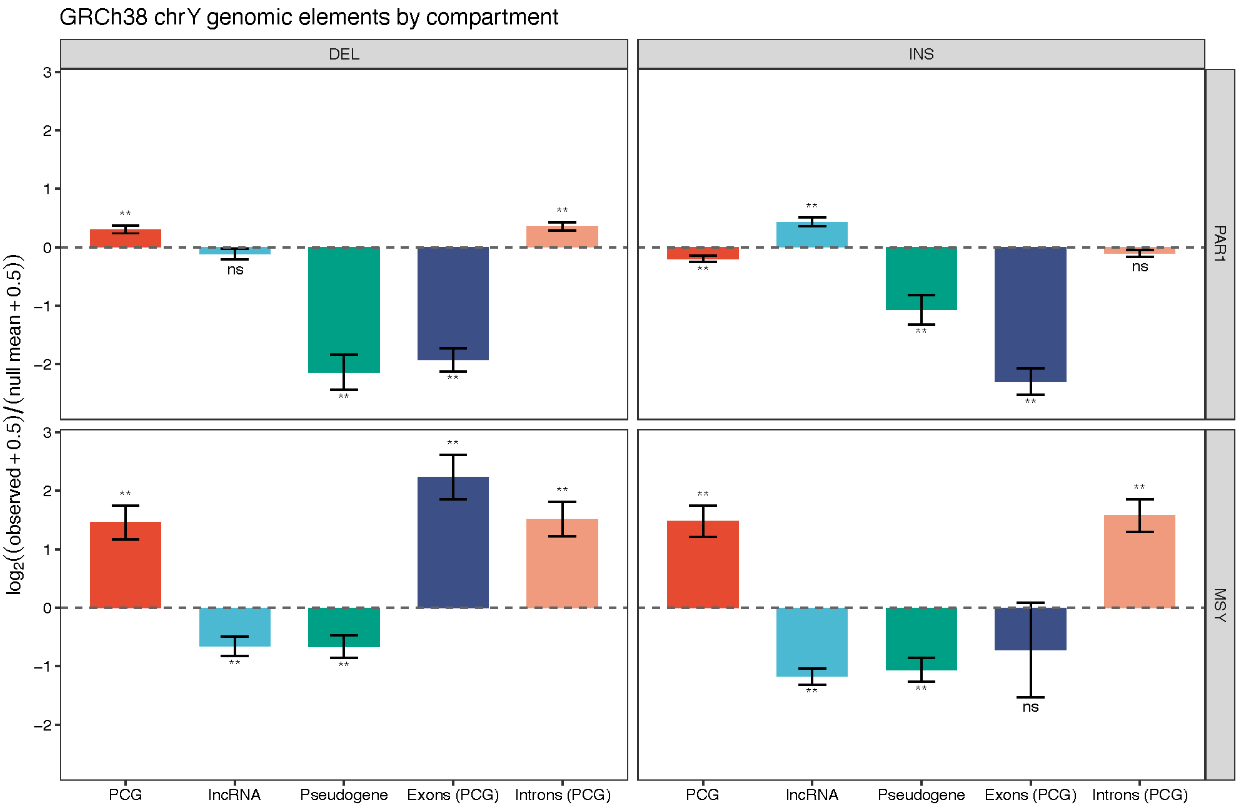


**Supplementary Figure 22. Compartment-specific enrichment of chromosome Y SVs across genomic elements in GRCh38.** Enrichment and depletion of SVs across GENCODE v47 protein-coding genes (PCGs), long noncoding RNAs (lncRNAs), and pseudogenes are shown separately for insertions and deletions in PAR1 and the male-specific region of the Y (MSY). Exonic and intronic regions of PCGs were analyzed separately. Error bars represent one standard deviation of the permutation null distribution transformed to the plotted log2 scale. * $q < 0.05$; ** $q < 0.01$; ns, not significant.

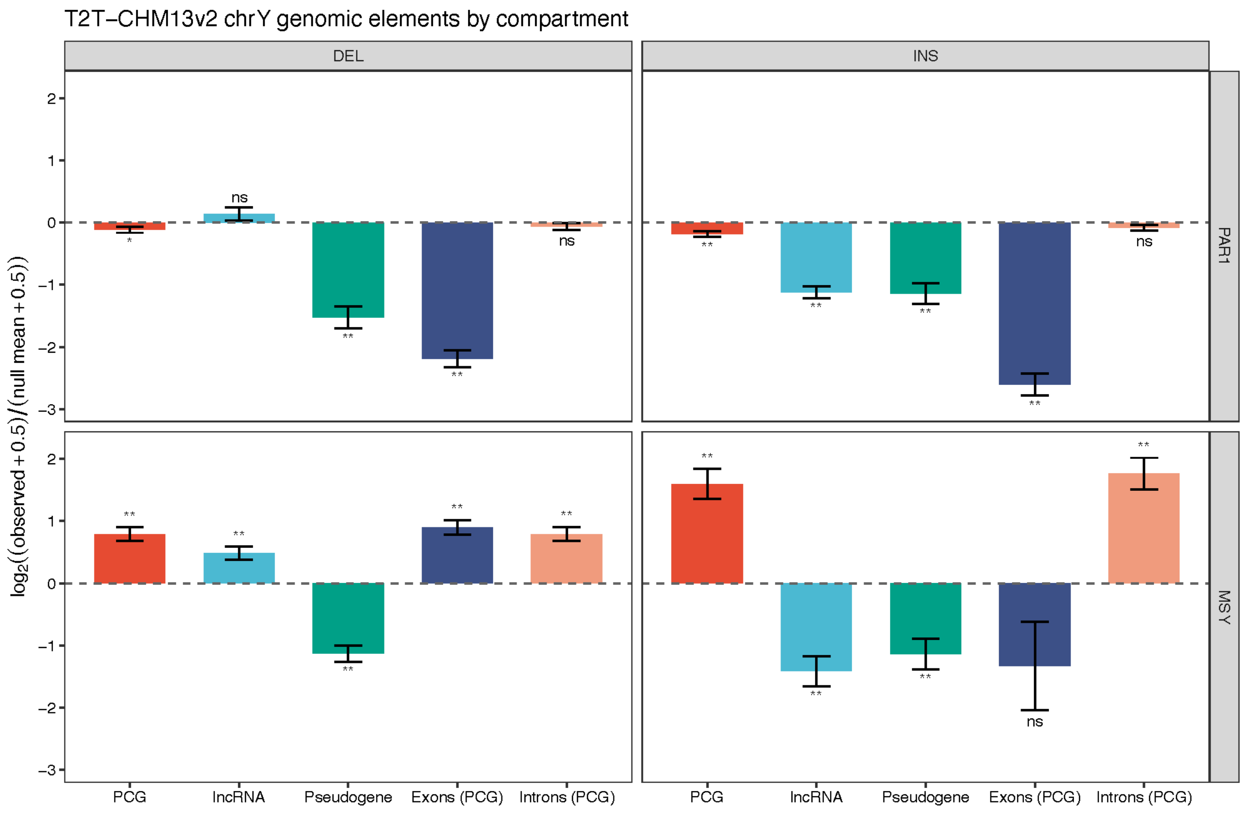


**Supplementary Figure 23. Compartment-specific enrichment of chromosome Y SVs across genomic elements in T2T-CHM13v2.** Enrichment and depletion of SVs across PCGs, lIncRNAs, and pseudogenes are shown separately for insertions and deletions in PAR1 and the MSY. Exonic and intronic regions of PCGs were analyzed separately. Error bars represent one standard deviation of the permutation null distribution transformed to the plotted log2 scale. * $q < 0.05$; ** $q < 0.01$; ns, not significant.

### *AZFc* Region

Contributing authors: Mark Loftus, Pille Hallast, David Porubsky

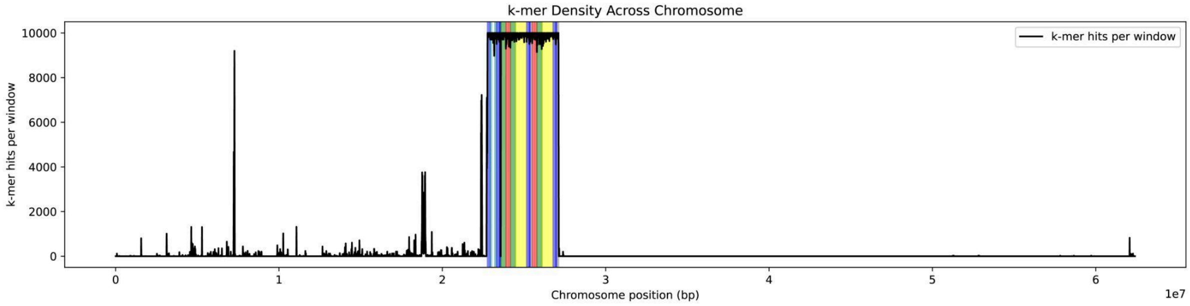


**Supplementary Figure 24. Example of *AZFc* colorblock localization using k-mer density in HG002.** The black trace line shows the number of matches per window for a set of *AZFc* "colorblock" k-mers derived from GRCh38, plotted across the HG002 Y-chromosome assembly (x-axis; assembly coordinates). Colored segments mark the inferred locations of each colorblock within the assembly based on local enrichment of their corresponding k-mer sets. K-mer match counts were computed in 10 kb sliding windows with a 1 kb step. This method can be used to find and label *AZFc* (and its internal blocks) directly from the sequence to allow the comparison of the *AZFc* structure and copy number organization across assemblies and individuals.

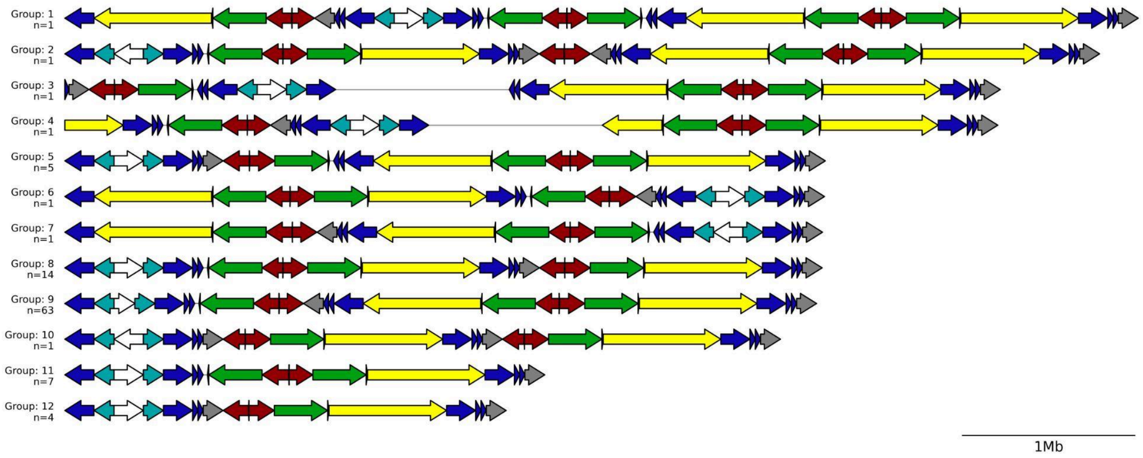


**Supplementary Figure 25. Distinct *AZFc* haplotypes.** One representative example is shown for each distinct AZFc haplotype architecture observed among 100 accurately and completely assembled AZFc regions. Twelve haplotype architectures were identified (frequency shown as *n*). ***Colored arrows*** denote the ordered sequence of *AZFc* colorblocks (arrow direction indicates orientation) and ***white arrows*** denote spacers. Group architectures were defined based on the presence and order of the major *AZFc* colorblocks. Spacer orientation and minor structural differences (e.g., small partial block fragments) were not used to distinguish groups. ***Black line breaks*** indicate intervening sequence of variable length between adjacent colorblocks (not to scale). Overall, across 100 high-quality assemblies, *AZFc* falls into 12 distinct haplotypes, with the two most common haplotypes accounting for 77% of samples (63% and 14%), while the remaining haplotype are much rarer (most seen only once).

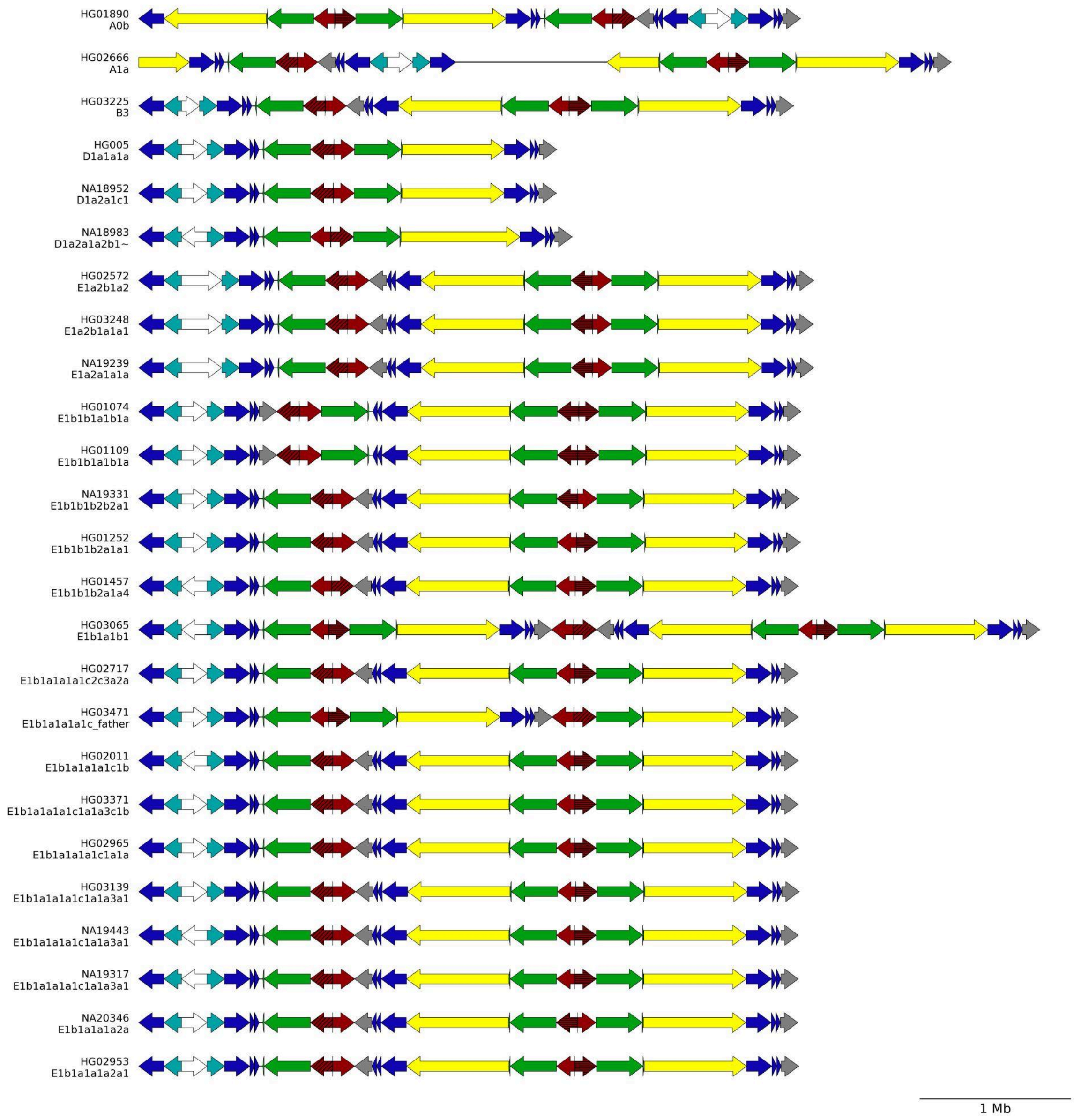
HG01890
A0b
HG02666
A1a
HG03225
B3
HG005
D1a1a1a
NA18952
D1a2a1c1
NA18983
D1a2a1a2b1~
HG02572
E1a2b1a2
HG03248
E1a2b1a1a1
NA19239
E1a2a1a1a
HG01074
E1b1b1a1b1a
HG01109
E1b1b1a1b1a
NA19331
E1b1b1b2b2a1
HG01252
E1b1b1b2a1a1
HG01457
E1b1b1b2a1a4
HG03065
E1b1a1b1
HG02717
E1b1a1a1a1c2c3a2a
HG03471
E1b1a1a1a1c_father
HG02011
E1b1a1a1a1c1b
HG03371
E1b1a1a1a1c1a1a3c1b
HG02965
E1b1a1a1a1c1a1a
HG03139
E1b1a1a1a1c1a1a3a1
NA19443
E1b1a1a1a1c1a1a3a1
NA19317
E1b1a1a1a1c1a1a3a1
NA20346
E1b1a1a1a2a
HG02953
E1b1a1a1a2a1
1 Mb

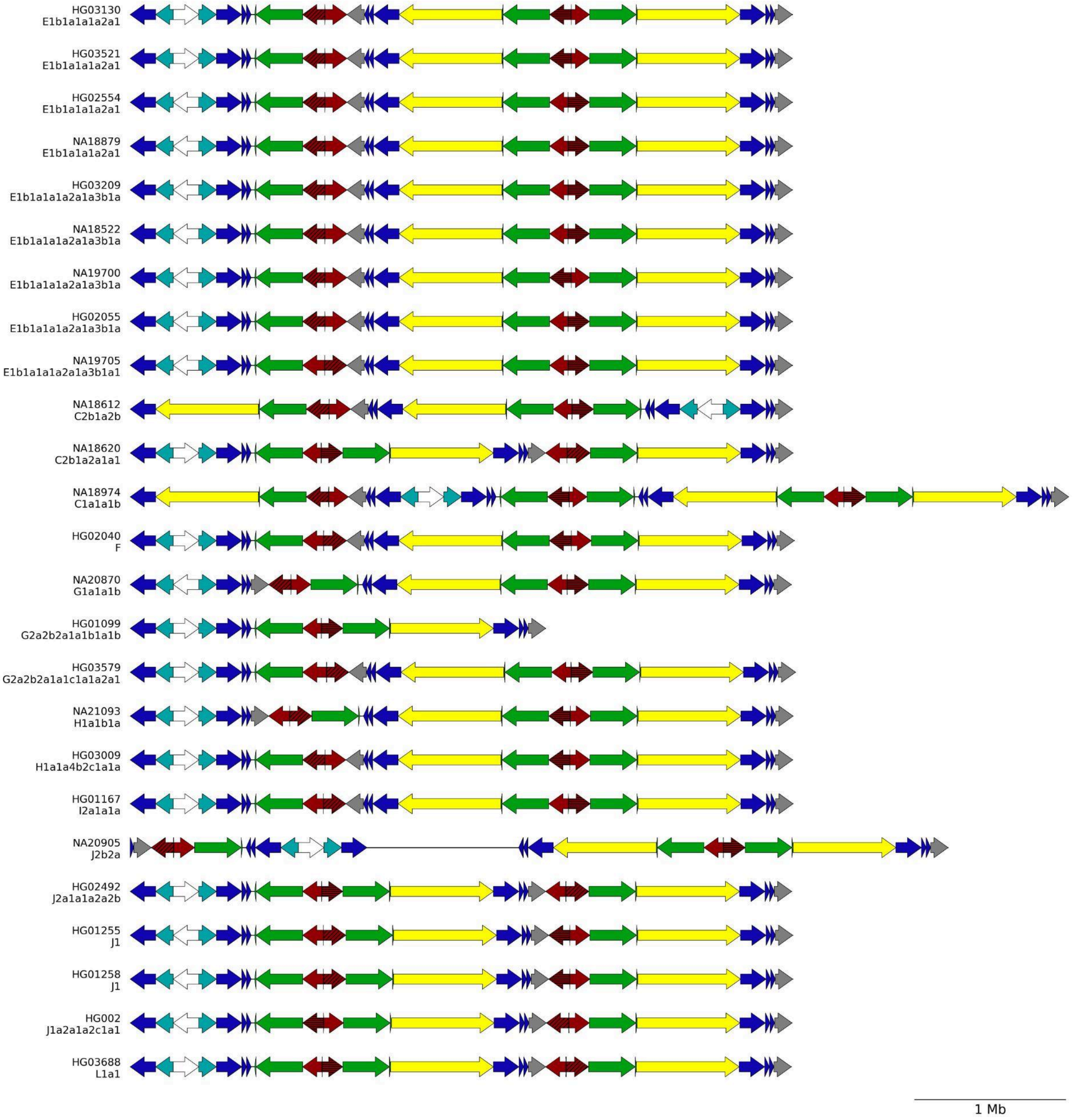
HG03130
E1b1a1a1a2a1
HG03521
E1b1a1a1a2a1
HG02554
E1b1a1a1a2a1
NA18879
E1b1a1a1a2a1
HG03209
E1b1a1a1a2a1a3b1a
NA18522
E1b1a1a1a2a1a3b1a
NA19700
E1b1a1a1a2a1a3b1a
HG02055
E1b1a1a1a2a1a3b1a
NA19705
E1b1a1a1a2a1a3b1a1
NA18612
C2b1a2b
NA18620
C2b1a2a1a1
NA18974
C1a1a1b
HG02040
F
NA20870
G1a1a1b
HG01099
G2a2b2a1a1b1a1b
HG03579
G2a2b2a1a1c1a1a2a1
NA21093
H1a1b1a
HG03009
H1a1a4b2c1a1a
HG01167
I2a1a1a
NA20905
J2b2a
HG02492
J2a1a1a2a2b
HG01255
J1
HG01258
J1
HG002
J1a2a1a2c1a1
HG03688
L1a1
1 Mb

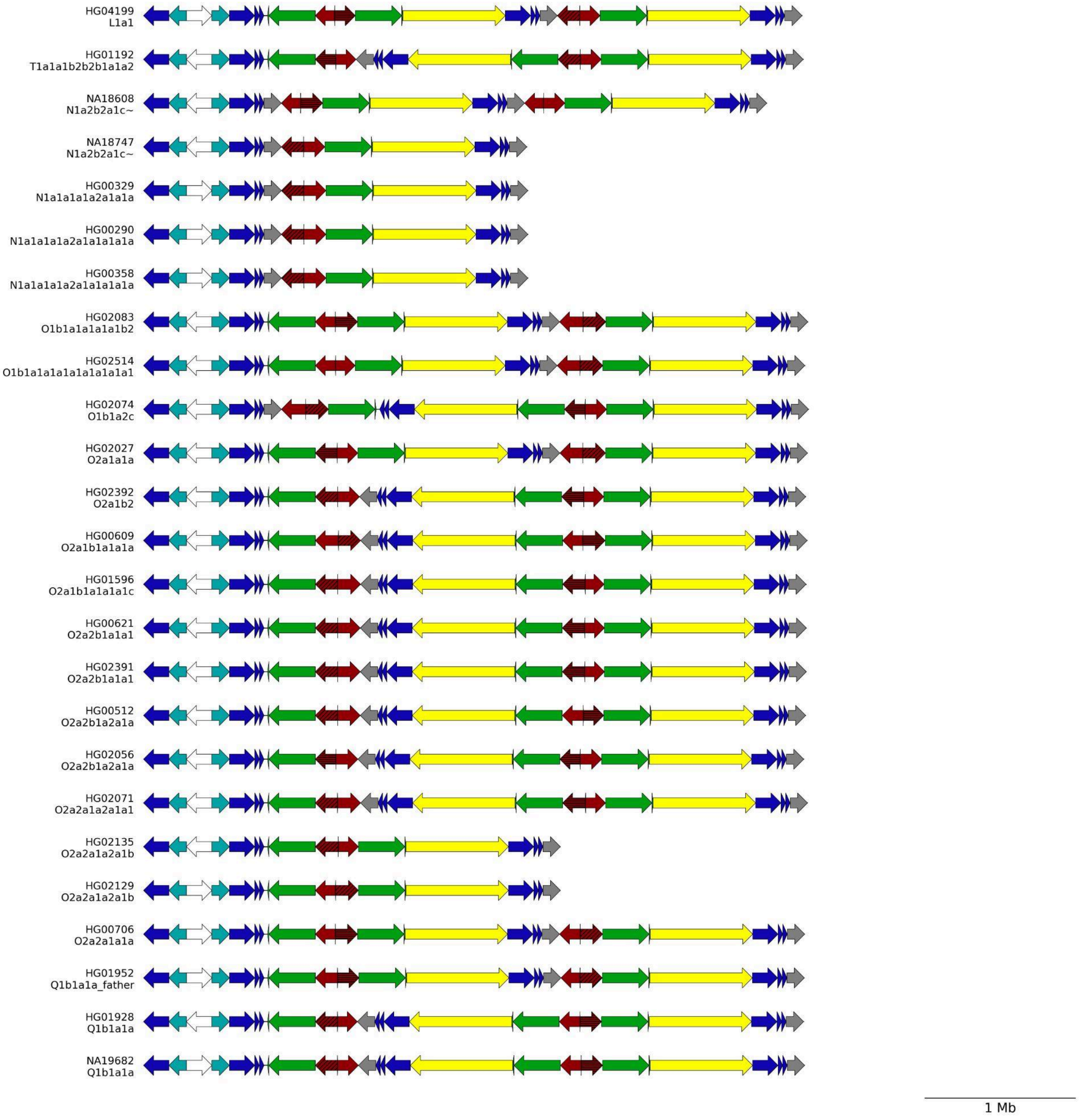
HG04199
L1a1
HG01192
T1a1a1b2b2b1a1a2
NA18608
N1a2b2a1c~
NA18747
N1a2b2a1c~
HG00329
N1a1a1a1a2a1a1a
HG00290
N1a1a1a1a2a1a1a1a1a
HG00358
N1a1a1a1a2a1a1a1a1a
HG02083
O1b1a1a1a1a1b2
HG02514
O1b1a1a1a1a1a1a1a1a1
HG02074
O1b1a2c
HG02027
O2a1a1a
HG02392
O2a1b2
HG00609
O2a1b1a1a1a
HG01596
O2a1b1a1a1a1c
HG00621
O2a2b1a1a1
HG02391
O2a2b1a1a1
HG00512
O2a2b1a2a1a
HG02056
O2a2b1a2a1a
HG02071
O2a2a1a2a1a1
HG02135
O2a2a1a2a1b
HG02129
O2a2a1a2a1b
HG00706
O2a2a1a1a
HG01952
Q1b1a1a_father
HG01928
Q1b1a1a
NA19682
Q1b1a1a
1 Mb

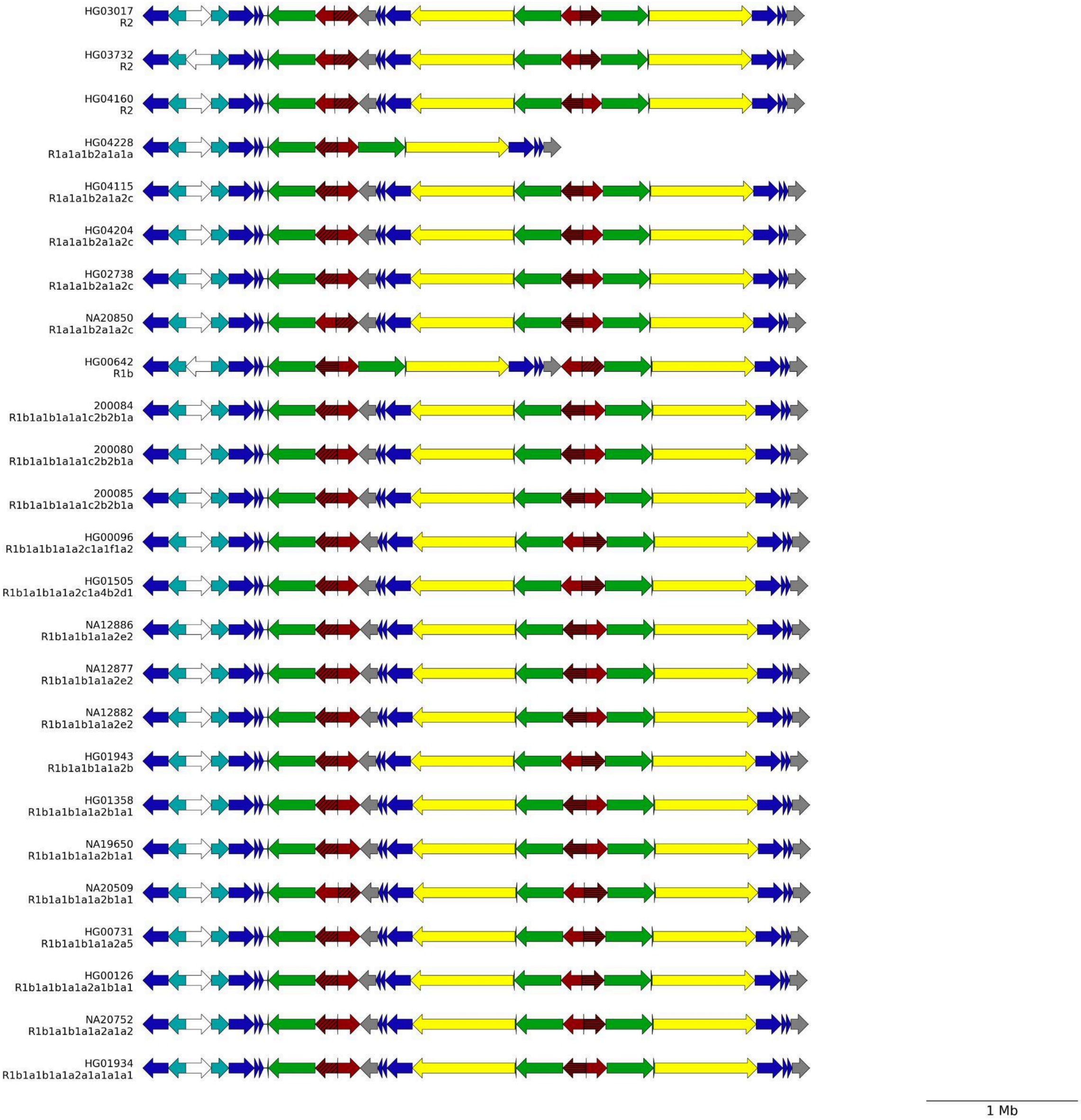


**Supplementary Figure 26. *AZFc* colorblock architectures across 100 QC-passing assemblies, with DAZ gene RRM copy number annotation.** *AZFc* colorblocks spanning palindromes P1-P3 are shown for 100 QC-passing samples. Samples are ordered phylogenetically with A haplogroup samples at the top, and Y-haplogroups are listed beneath each sample name. Colored arrows denote *AZFc* colorblocks. Arrow direction indicates colorblock orientation (forward sense/+ to the ***right***; reverse antisense/- to the ***left***). *DAZ*-containing blocks (***red***) are further annotated by hatching to indicate the number of RNA-recognition motifs (RRMs) in the associated *DAZ* gene copy: 1 RRM (***no hatching***), 2 RRMs (***horizontal hatching***), and 3 RRMs (***diagonal hatching***).

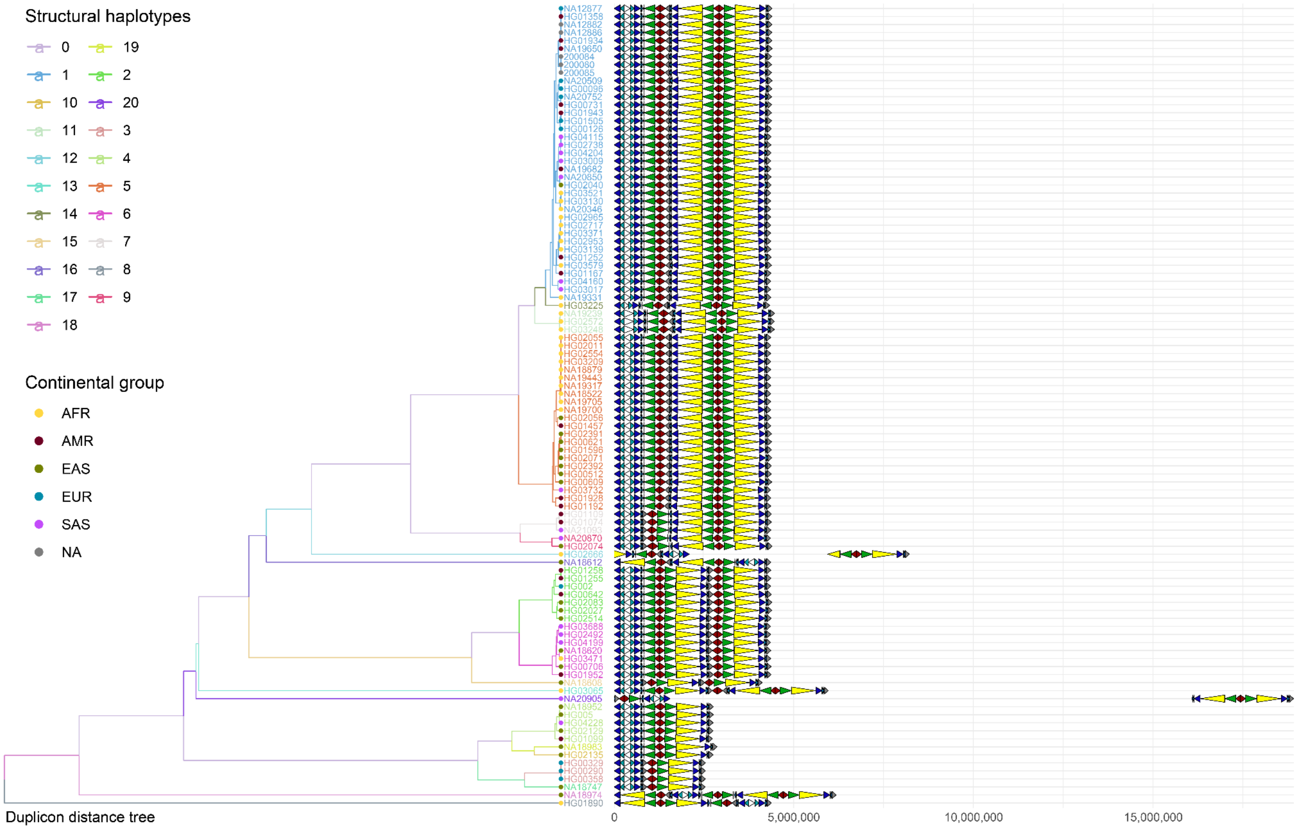


**Supplementary Figure 27. Clustering of *AZFc* region based on structural similarity.** A hierarchical clustering ("UPGMA tree"; ***left***) groups 100 QC-passing assemblies by similarity of *AZFc* duplicon/colorblock architecture. Each row corresponds to one sample, with the sample label indicating structural haplotypes (***top legend***) and continental groups (***bottom legend***; AFR, AMR, EAS, EUR, SAS, NA). To the right of the tree, colored arrows depict the ordered *AZFc* colorblocks (palindromes P1–P3) for each sample; arrow direction indicates block orientation (+/forward to the ***right***; −/reverse to the ***left***). The x-axis shows position along the plotted *AZFc* architecture (bp), with large gaps indicating variable-length intervening sequence.

### *gr/gr* recurrent inversions

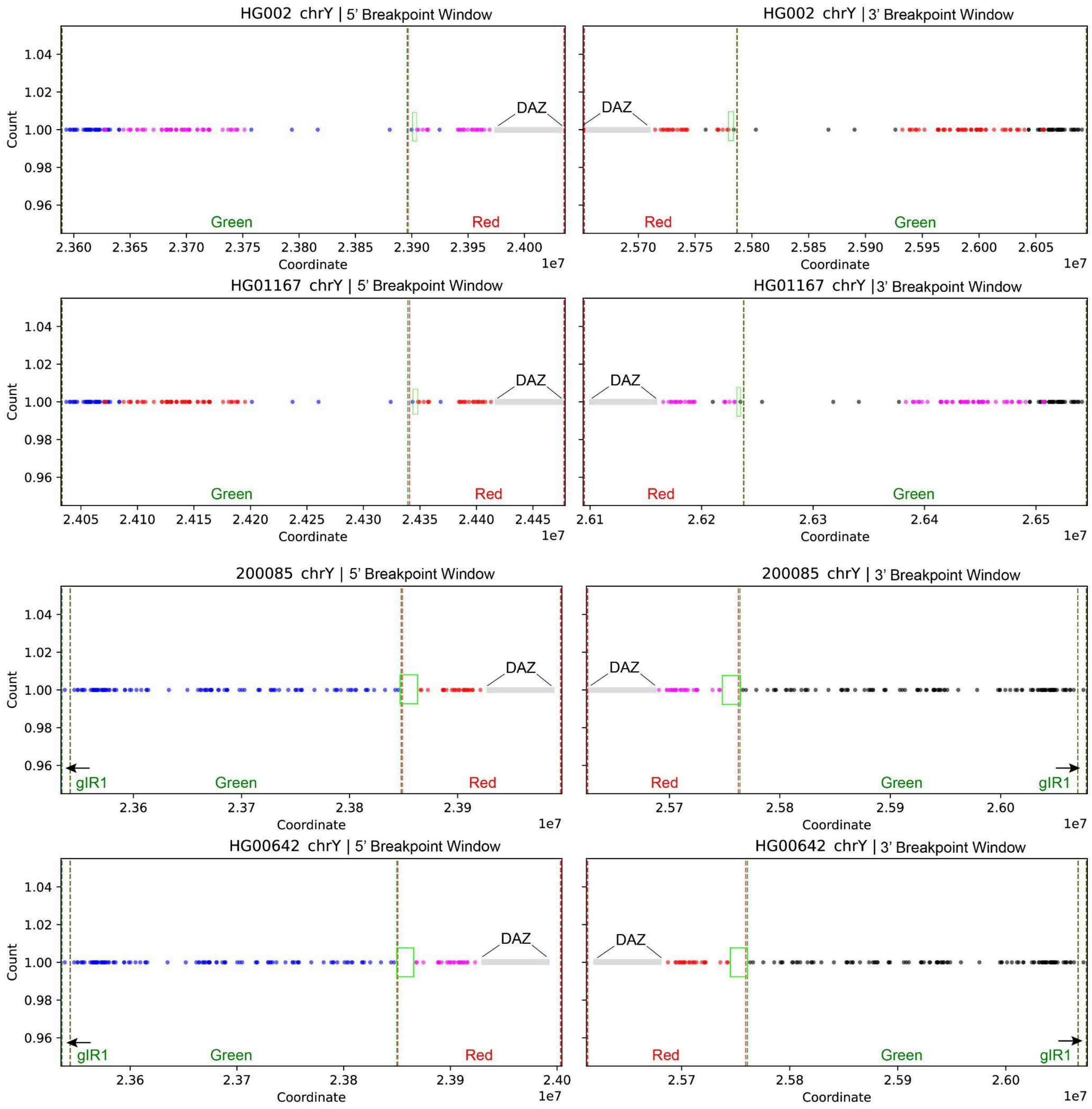

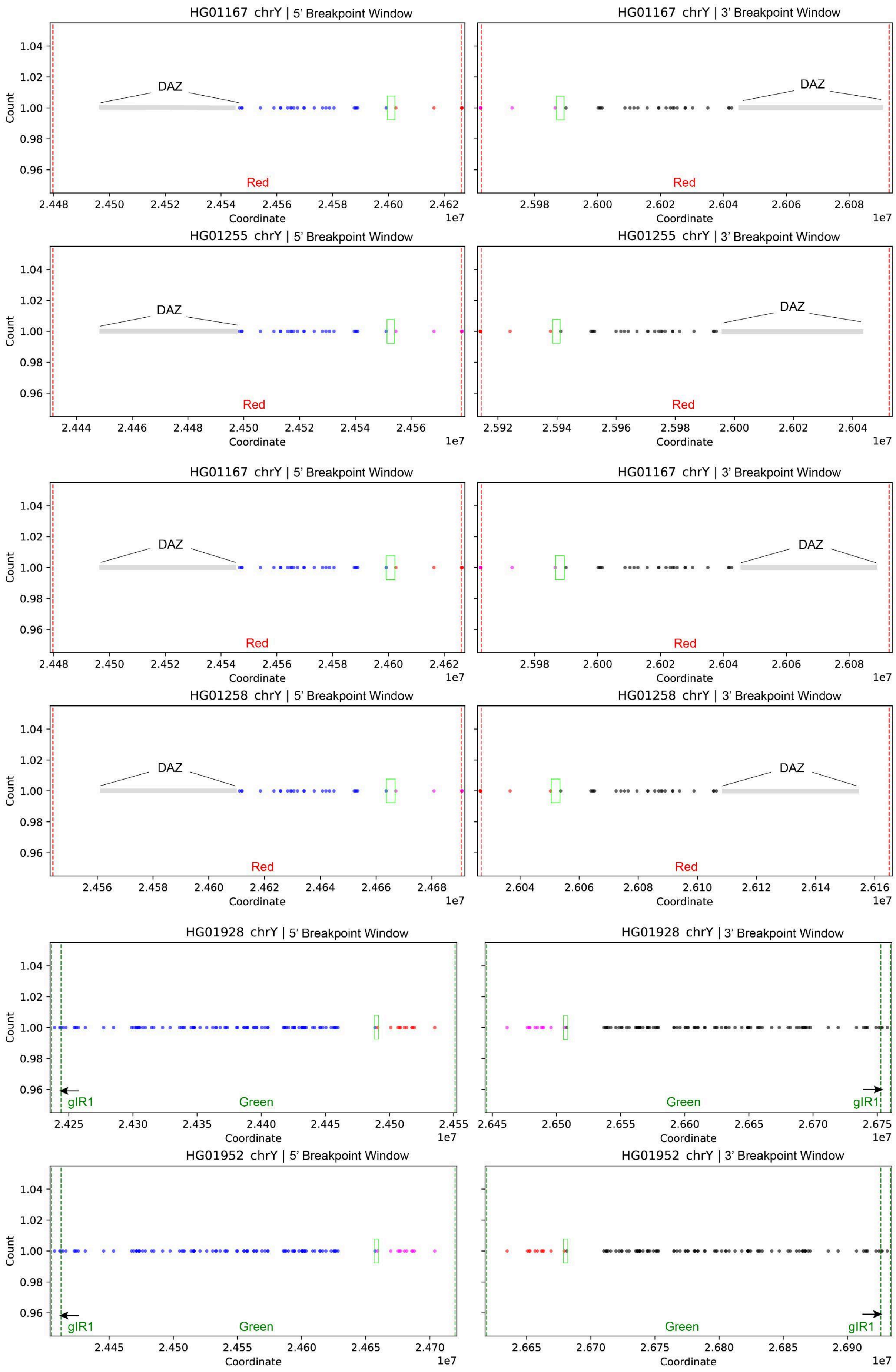
HG01167 chrY | 5' Breakpoint Window
HG01167 chrY | 3' Breakpoint Window
HG01255 chrY | 5' Breakpoint Window
HG01255 chrY | 3' Breakpoint Window
HG01167 chrY | 5' Breakpoint Window
HG01167 chrY | 3' Breakpoint Window
HG01258 chrY | 5' Breakpoint Window
HG01258 chrY | 3' Breakpoint Window
HG01928 chrY | 5' Breakpoint Window
HG01928 chrY | 3' Breakpoint Window
HG01952 chrY | 5' Breakpoint Window
HG01952 chrY | 3' Breakpoint Window
DAZ
Red
Green
gIR1
Count
Coordinate
1e7

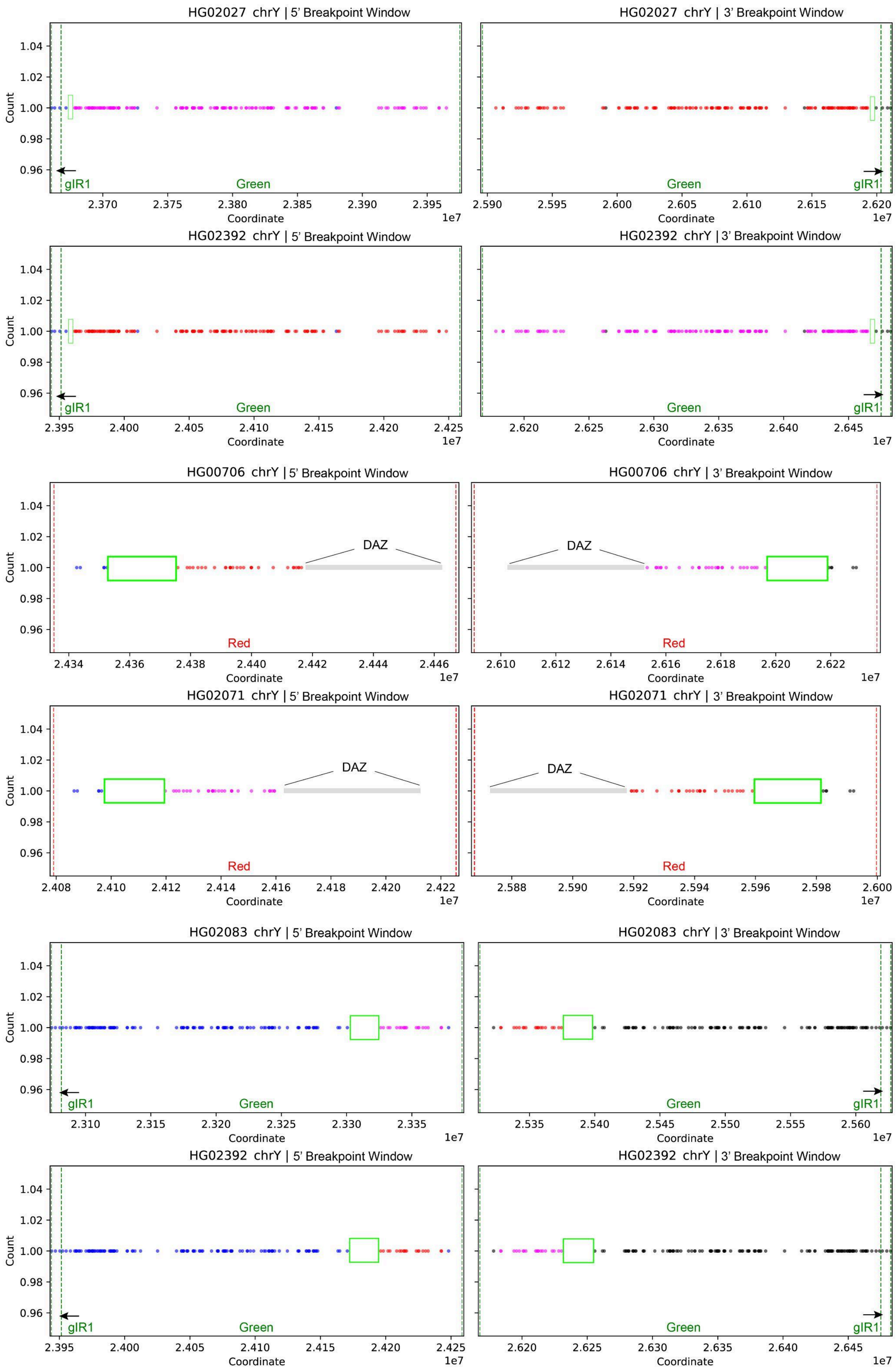
HG02027 chrY | 5' Breakpoint Window
HG02027 chrY | 3' Breakpoint Window
HG02392 chrY | 5' Breakpoint Window
HG02392 chrY | 3' Breakpoint Window
HG00706 chrY | 5' Breakpoint Window
HG00706 chrY | 3' Breakpoint Window
HG02071 chrY | 5' Breakpoint Window
HG02071 chrY | 3' Breakpoint Window
HG02083 chrY | 5' Breakpoint Window
HG02083 chrY | 3' Breakpoint Window
HG02392 chrY | 5' Breakpoint Window
HG02392 chrY | 3' Breakpoint Window
Count
Coordinate
1e7
gIR1
Green
Red
DAZ

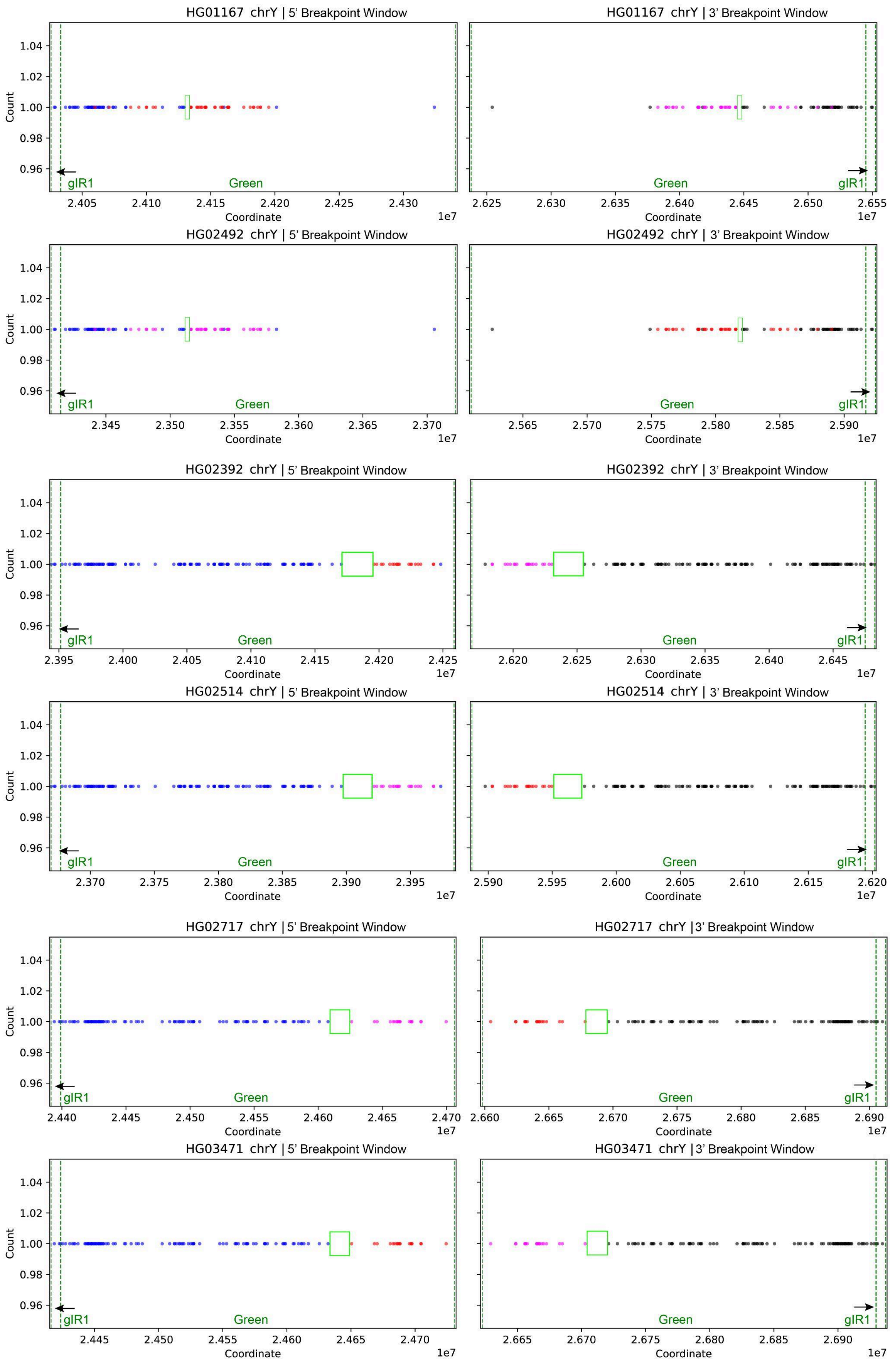
HG01167 chrY | 5' Breakpoint Window
HG01167 chrY | 3' Breakpoint Window
HG02492 chrY | 5' Breakpoint Window
HG02492 chrY | 3' Breakpoint Window
HG02392 chrY | 5' Breakpoint Window
HG02392 chrY | 3' Breakpoint Window
HG02514 chrY | 5' Breakpoint Window
HG02514 chrY | 3' Breakpoint Window
HG02717 chrY | 5' Breakpoint Window
HG02717 chrY | 3' Breakpoint Window
HG03471 chrY | 5' Breakpoint Window
HG03471 chrY | 3' Breakpoint Window
Count
Coordinate
1e7
gIR1
Green

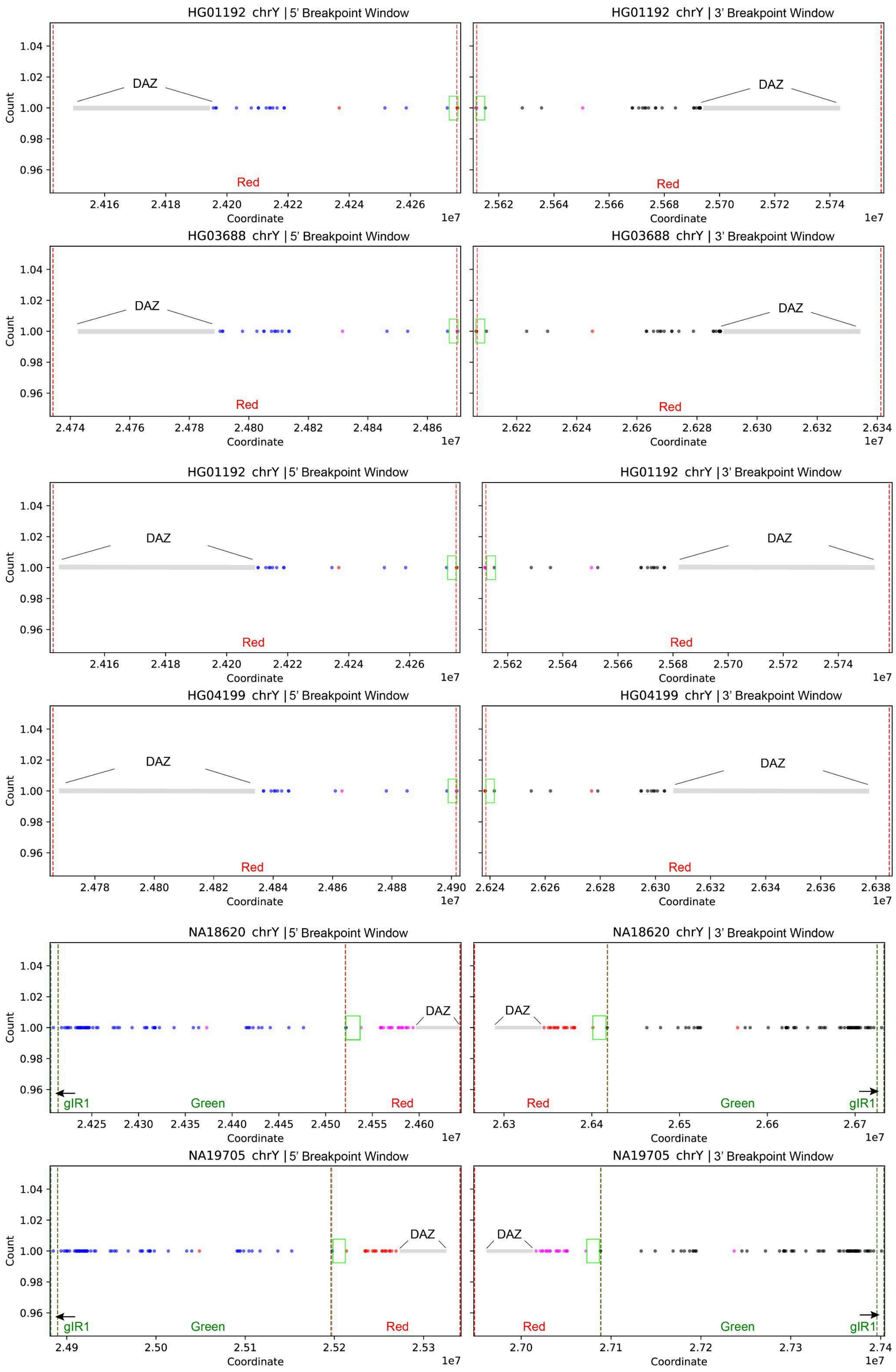

HG01192 chrY | 5' Breakpoint Window
HG01192 chrY | 3' Breakpoint Window
HG03688 chrY | 5' Breakpoint Window
HG03688 chrY | 3' Breakpoint Window
HG01192 chrY | 5' Breakpoint Window
HG01192 chrY | 3' Breakpoint Window
HG04199 chrY | 5' Breakpoint Window
HG04199 chrY | 3' Breakpoint Window
NA18620 chrY | 5' Breakpoint Window
NA18620 chrY | 3' Breakpoint Window
NA19705 chrY | 5' Breakpoint Window
NA19705 chrY | 3' Breakpoint Window
DAZ
Red
Green
gIR1
Count
Coordinate
1e7

**Supplementary Figure 28. *gr/gr* recurrent inversion breakpoint windows.** Each panel shows *AZFc* colorblocks from a sample carrying the *gr/gr* inversion compared with the nearest phylogenetic neighbor that has a GRCh38-like architecture. Points represent individual SNPs positioned by assembly coordinates (x-axis). Text annotation across the bottom indicates the colorblock region boundaries (Green-IR1, ***Green***, ***Red***). ***Blue*** and ***black*** SNPs fall outside the inverted interval, whereas ***red*** and ***pink*** SNPs lie within the inversion. ***Lime-green squares*** mark the inferred inversion breakpoint window(s).

### *b2/b3* recurrent inversions

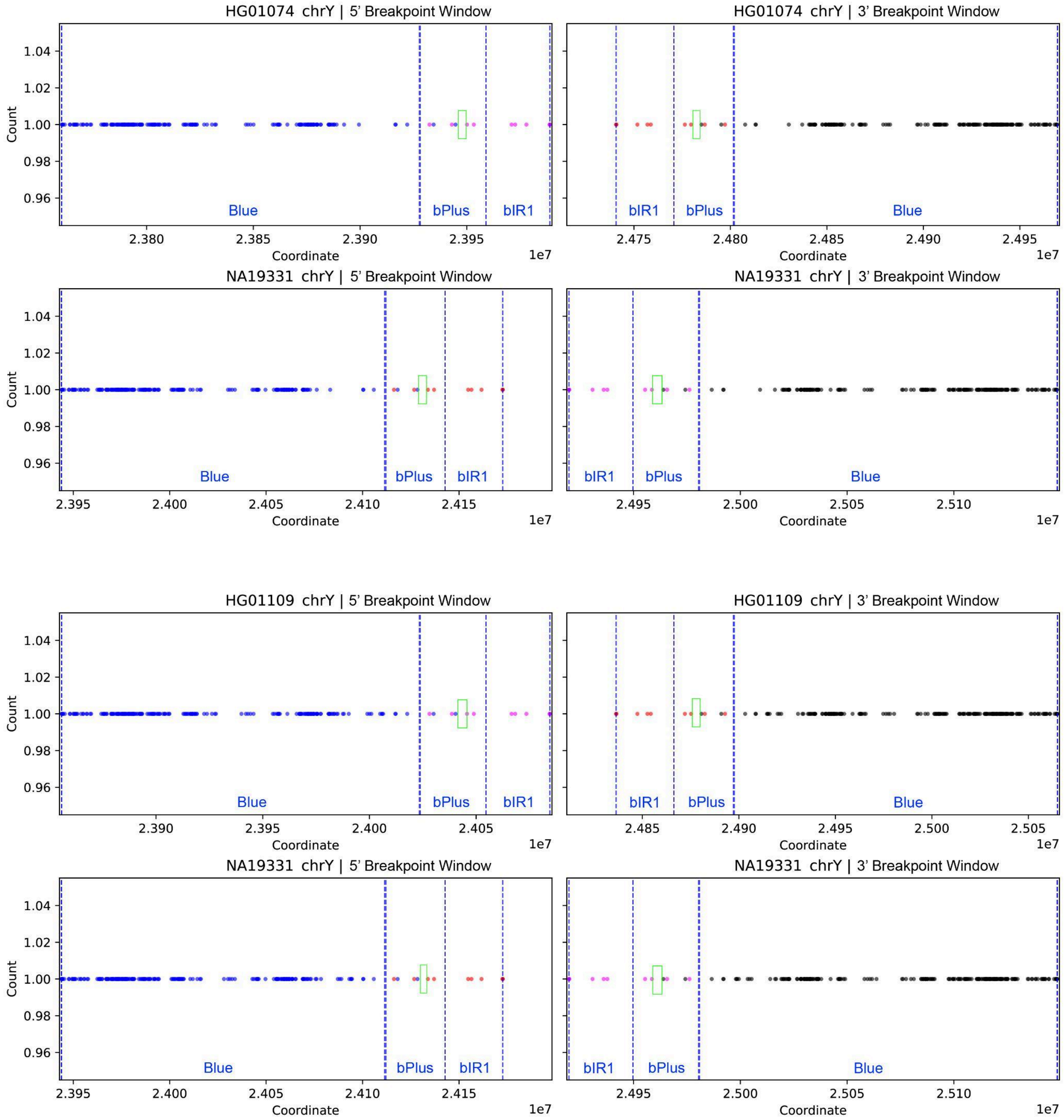

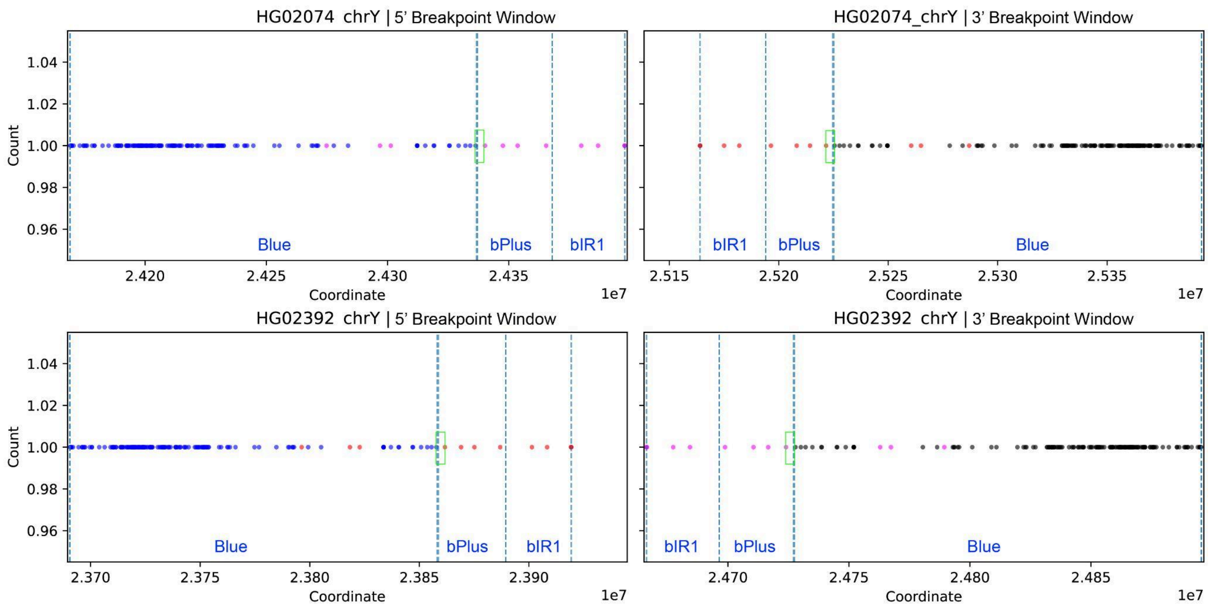
HG02074 chrY | 5' Breakpoint Window
HG02074_chrY | 3' Breakpoint Window
HG02392 chrY | 5' Breakpoint Window
HG02392 chrY | 3' Breakpoint Window
Count
Coordinate
1e7
Blue
bPlus
bIR1

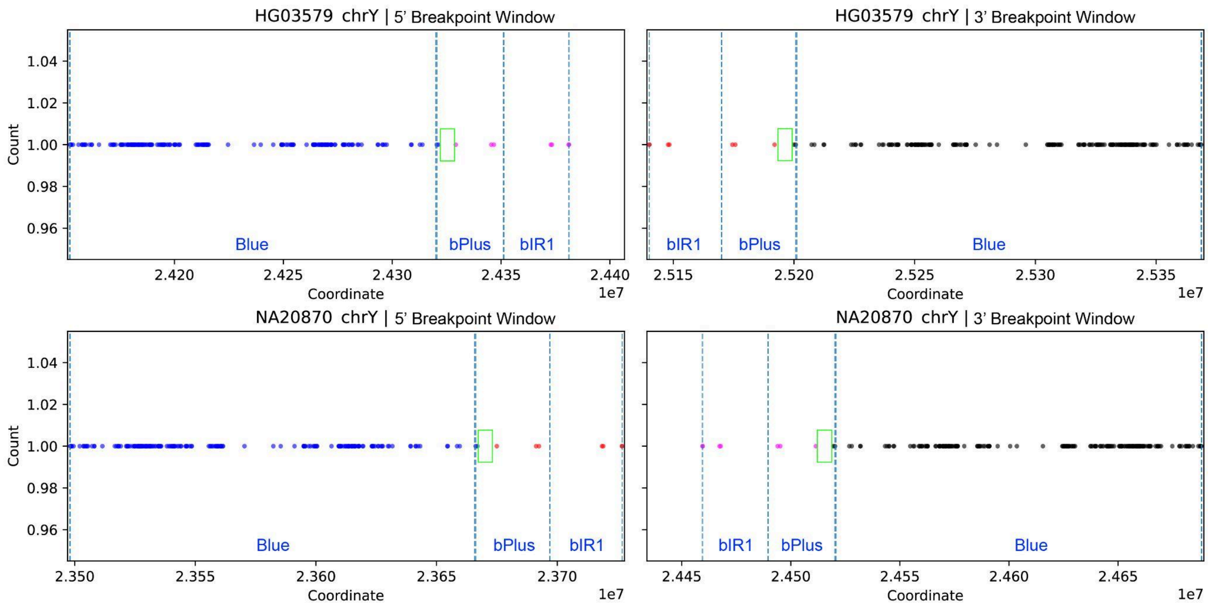
HG03579 chrY | 5' Breakpoint Window
HG03579 chrY | 3' Breakpoint Window
NA20870 chrY | 5' Breakpoint Window
NA20870 chrY | 3' Breakpoint Window
Count
Coordinate
1e7
Blue
bPlus
bIR1

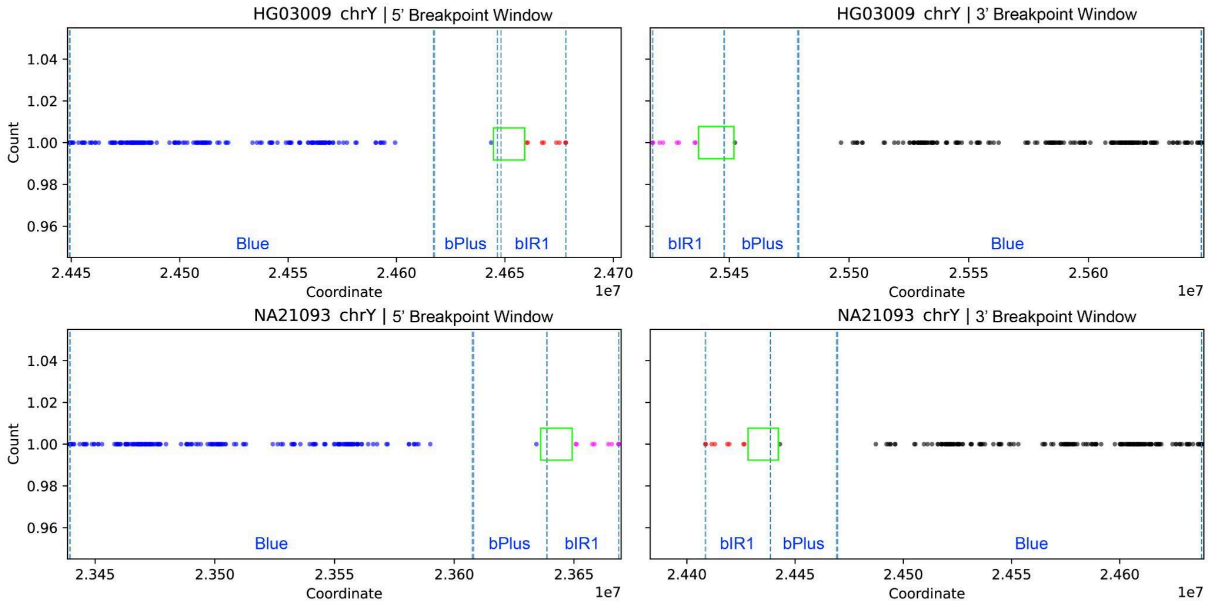


**Supplementary Figure 29. *b2/b3* recurrent inversion breakpoint windows inferred from SNP patterns.** Each panel shows *AZFc* colorblocks from a sample carrying the *b2/b3* inversion compared with the nearest phylogenetic neighbor that has a GRCh38-like architecture. Points represent individual SNPs positioned by assembly coordinates (x-axis). Text annotation across the bottom indicates the colorblock region boundaries (***Blue block***, Blue-Plus, and Blue-IR1). Blue and black SNPs fall outside the inverted interval, whereas ***red*** and ***pink*** SNPs lie within the inversion. Lime-green squares mark the inferred inversion breakpoint window(s).

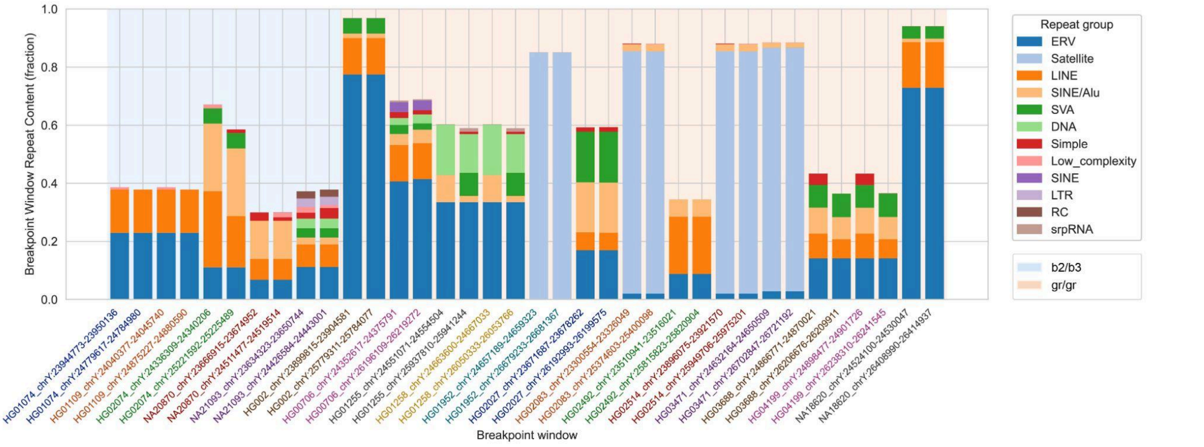


**Supplementary Figure 30. Repeat composition of *AZFc* inversion breakpoint windows.** Stacked bar plots show the repeat content of each inversion breakpoint window, where each bar represents a single breakpoint window and stacked segments indicate the fraction of the window annotated as each repeat group. Repeat classes were consolidated into major categories (e.g., ERV, LINE, SINE, SINE/Alu, LTR, DNA (transposon), Satellite, Simple, and low-complexity). Breakpoint windows are ordered by the corresponding comparison sample. Shaded background bands denote groups of windows associated with the same inversion type (*b2/b3* or *gr/gr*), and x-axis tick label colors indicate the sample from which each breakpoint window was derived.

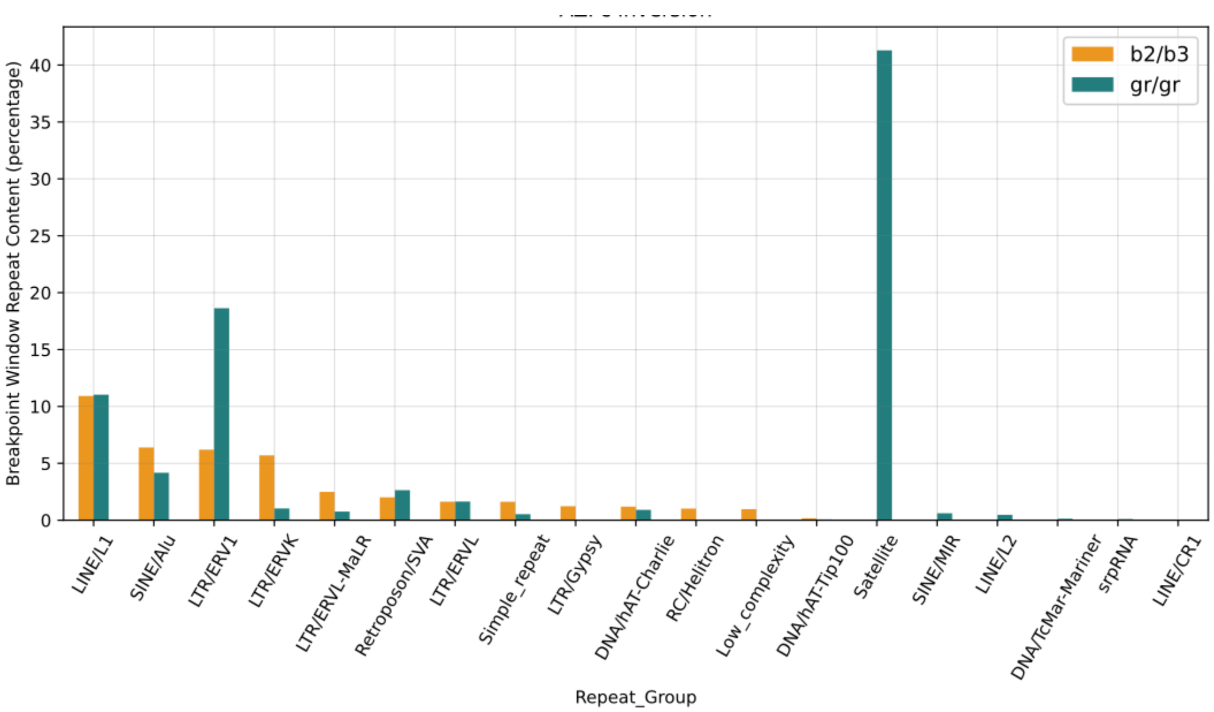


**Supplementary Figure 31. Percentage of repeats within *AZFc* inversion breakpoint windows.** Bar plot showing the total percentage of each Repeat Group present within the breakpoint window across all samples, by inversion type.

### *AZFc* region deletion breakpoints

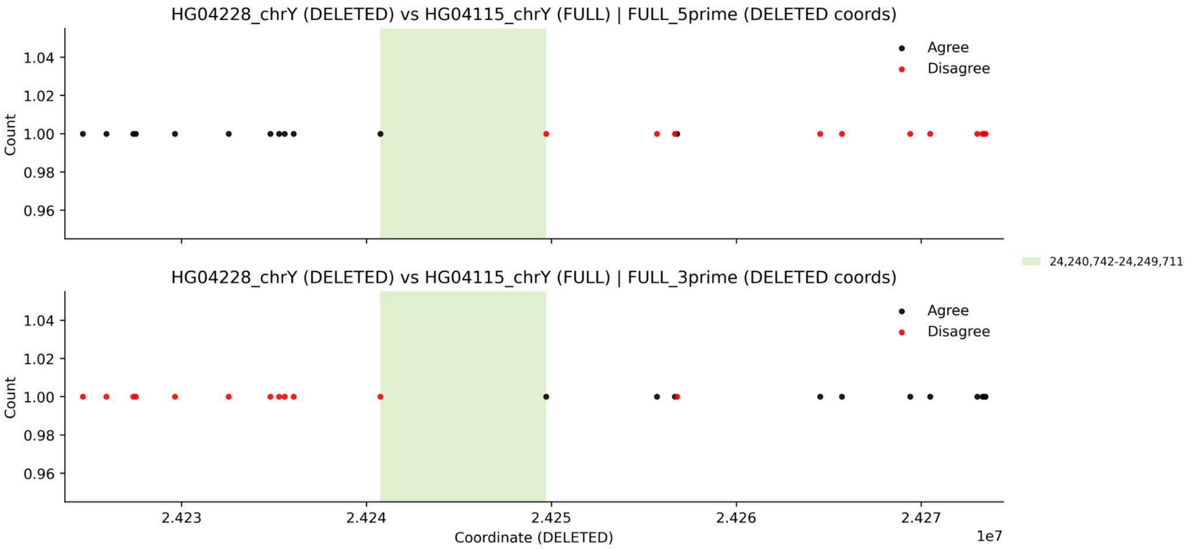


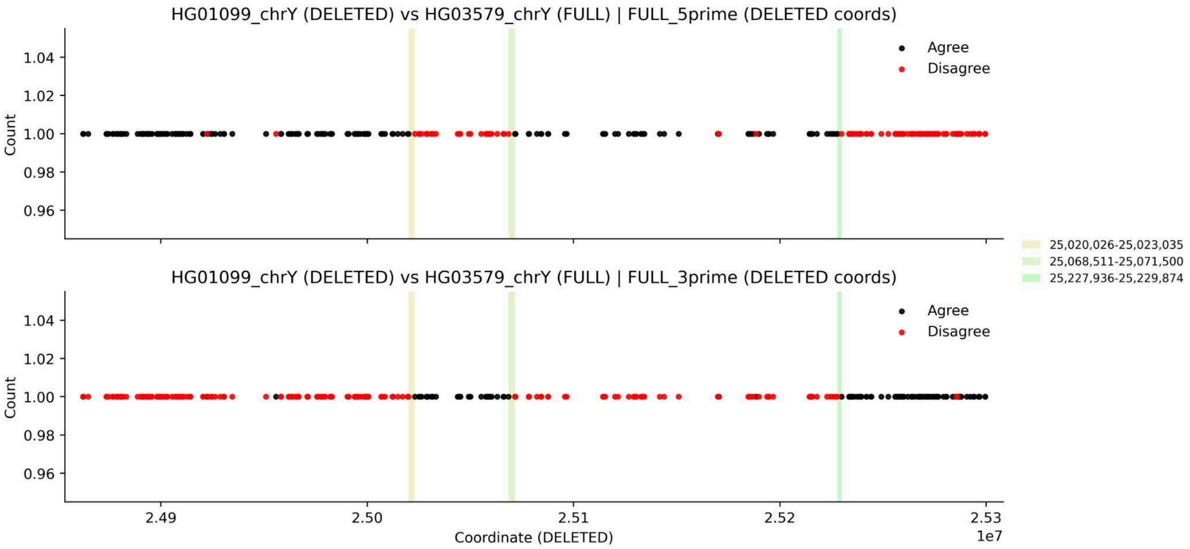


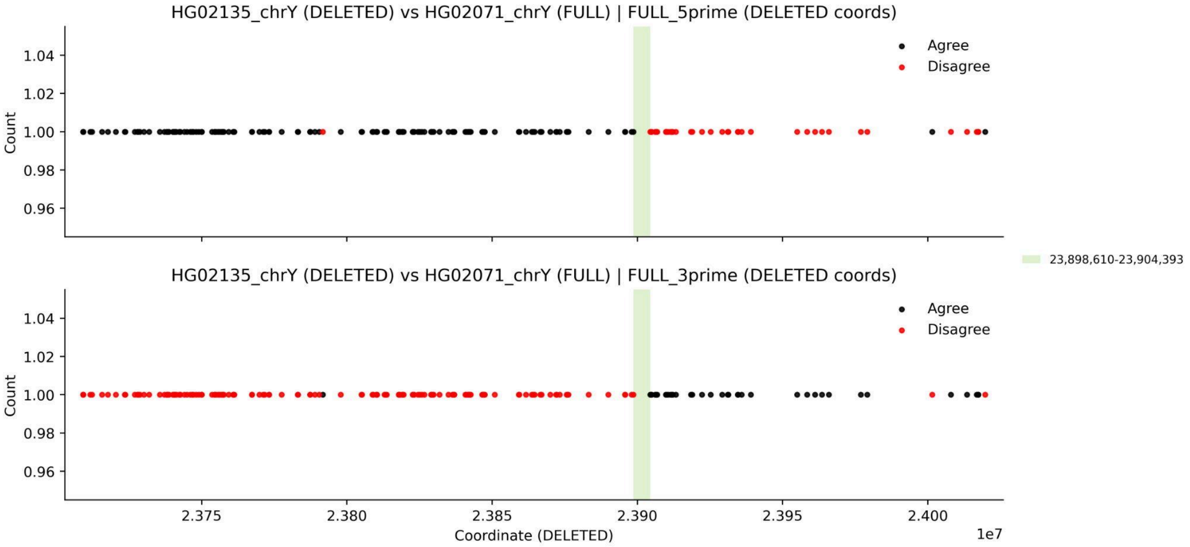

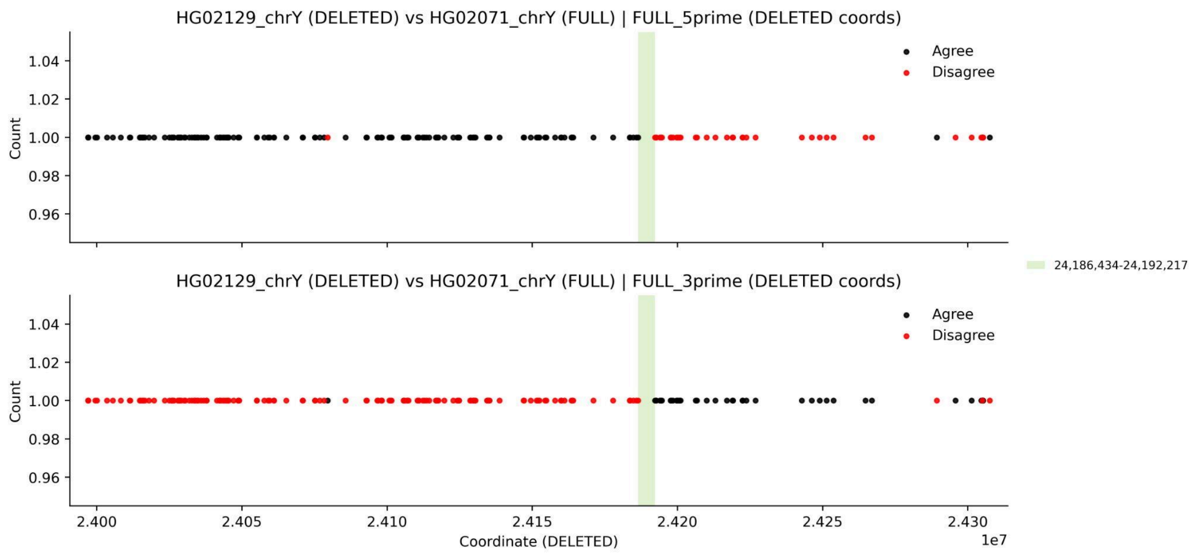
HG02129_chrY (DELETED) vs HG02071_chrY (FULL) | FULL_5prime (DELETED coords)
HG02129_chrY (DELETED) vs HG02071_chrY (FULL) | FULL_3prime (DELETED coords)
Agree
Disagree
Count
Coordinate (DELETED)
1e7
24,186,434-24,192,217

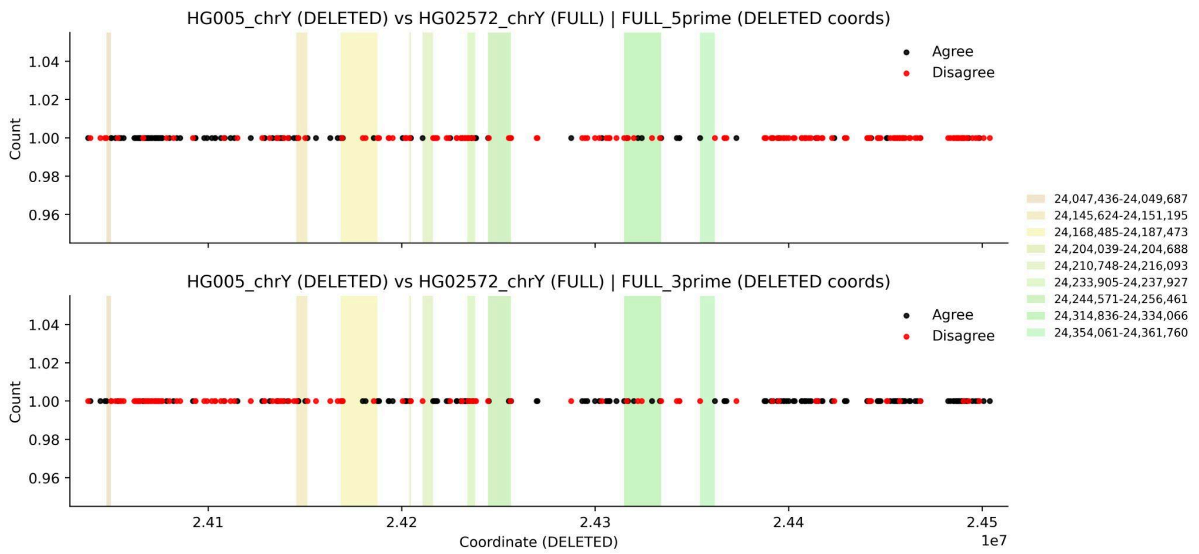
HG005_chrY (DELETED) vs HG02572_chrY (FULL) | FULL_5prime (DELETED coords)
HG005_chrY (DELETED) vs HG02572_chrY (FULL) | FULL_3prime (DELETED coords)
Agree
Disagree
Count
Coordinate (DELETED)
1e7
24,047,436-24,049,687
24,145,624-24,151,195
24,168,485-24,187,473
24,204,039-24,204,688
24,210,748-24,216,093
24,233,905-24,237,927
24,244,571-24,256,461
24,314,836-24,334,066
24,354,061-24,361,760

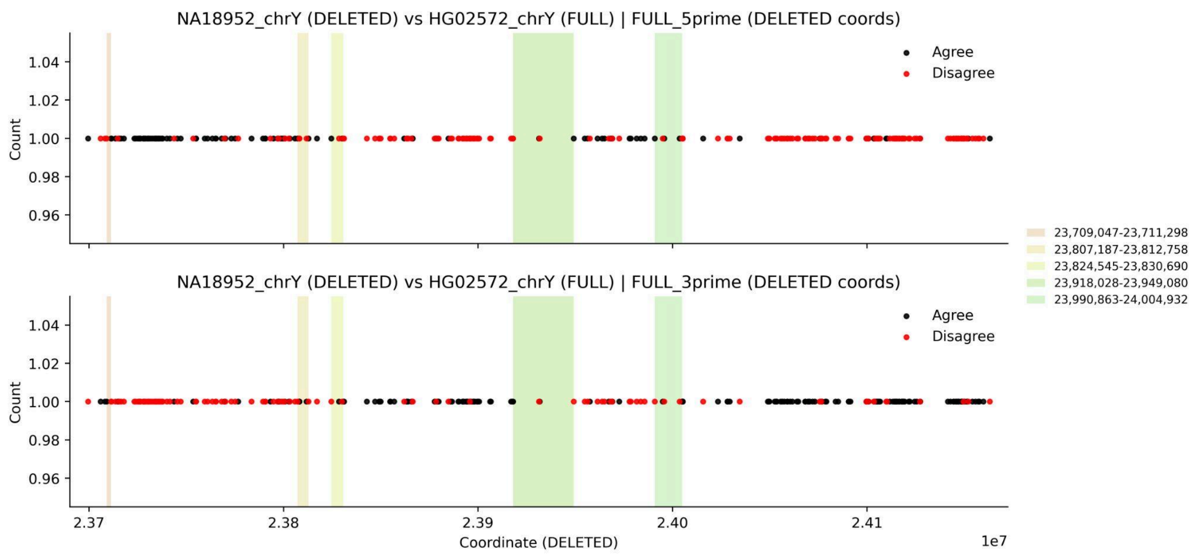
NA18952_chrY (DELETED) vs HG02572_chrY (FULL) | FULL_5prime (DELETED coords)
NA18952_chrY (DELETED) vs HG02572_chrY (FULL) | FULL_3prime (DELETED coords)
Agree
Disagree
Count
Coordinate (DELETED)
1e7
23,709,047-23,711,298
23,807,187-23,812,758
23,824,545-23,830,690
23,918,028-23,949,080
23,990,863-24,004,932

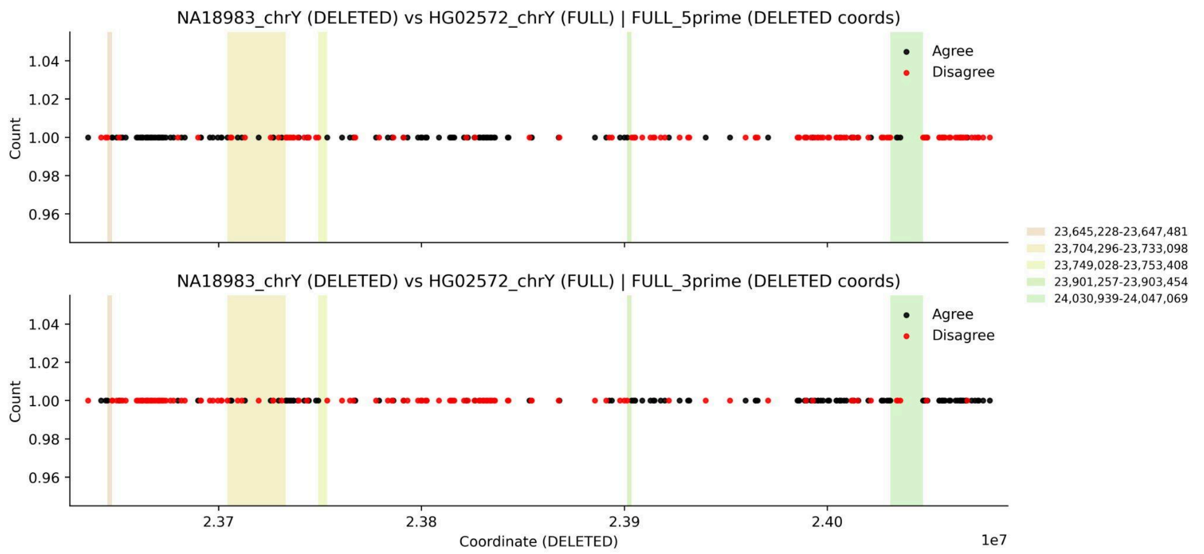
NA18983_chrY (DELETED) vs HG02572_chrY (FULL) | FULL_5prime (DELETED coords)
NA18983_chrY (DELETED) vs HG02572_chrY (FULL) | FULL_3prime (DELETED coords)
Count
Agree
Disagree
23,645,228-23,647,481
23,704,296-23,733,098
23,749,028-23,753,408
23,901,257-23,903,454
24,030,939-24,047,069
Coordinate (DELETED)
1e7

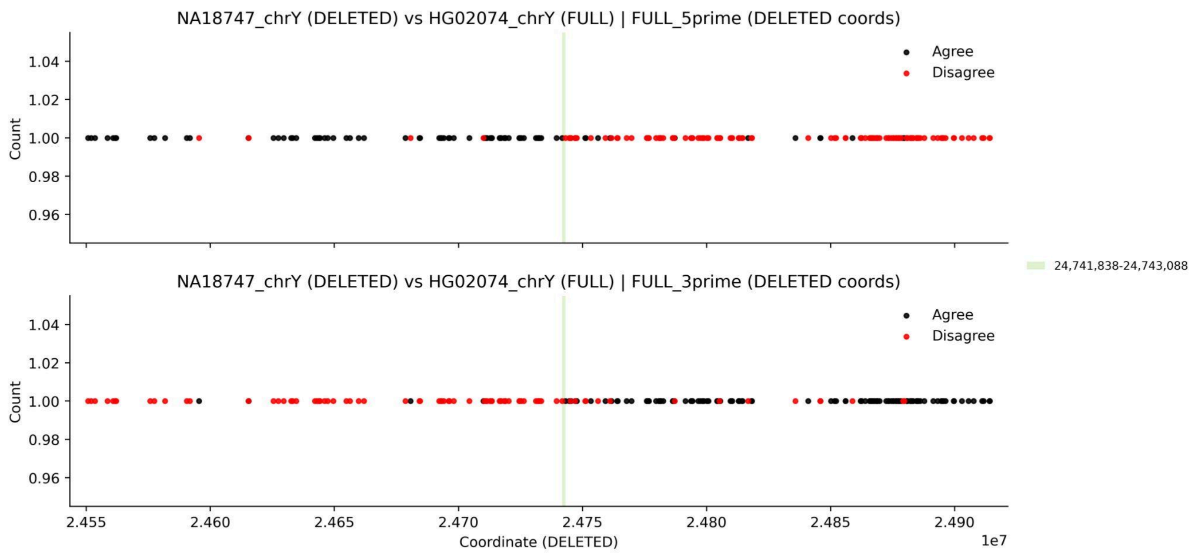
NA18747_chrY (DELETED) vs HG02074_chrY (FULL) | FULL_5prime (DELETED coords)
NA18747_chrY (DELETED) vs HG02074_chrY (FULL) | FULL_3prime (DELETED coords)
Count
Agree
Disagree
24,741,838-24,743,088
Coordinate (DELETED)
1e7

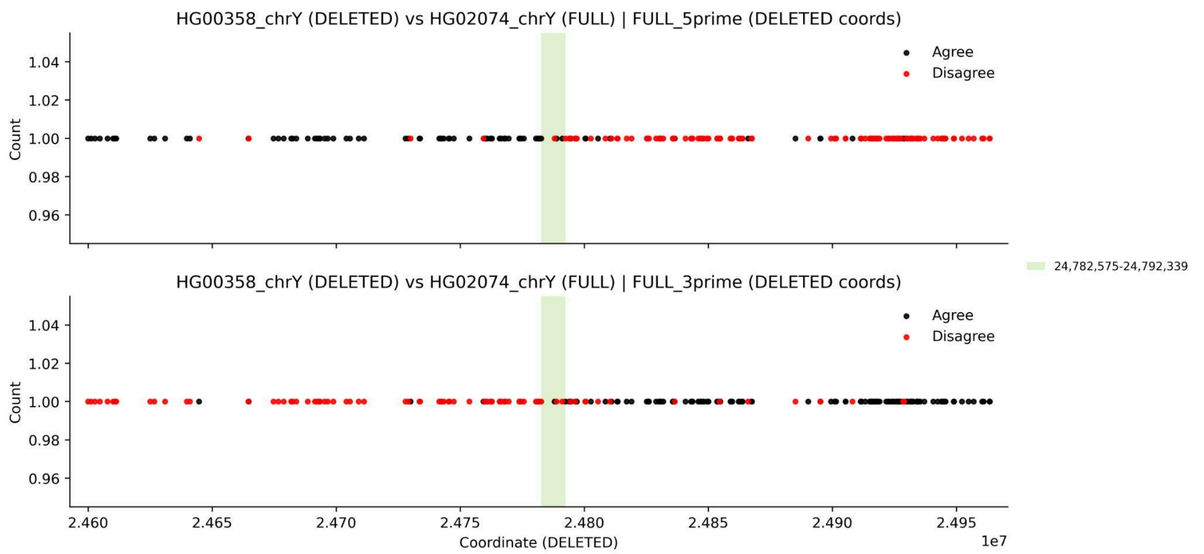
HG00358_chrY (DELETED) vs HG02074_chrY (FULL) | FULL_5prime (DELETED coords)
HG00358_chrY (DELETED) vs HG02074_chrY (FULL) | FULL_3prime (DELETED coords)
Count
Agree
Disagree
24,782,575-24,792,339
Coordinate (DELETED)
1e7

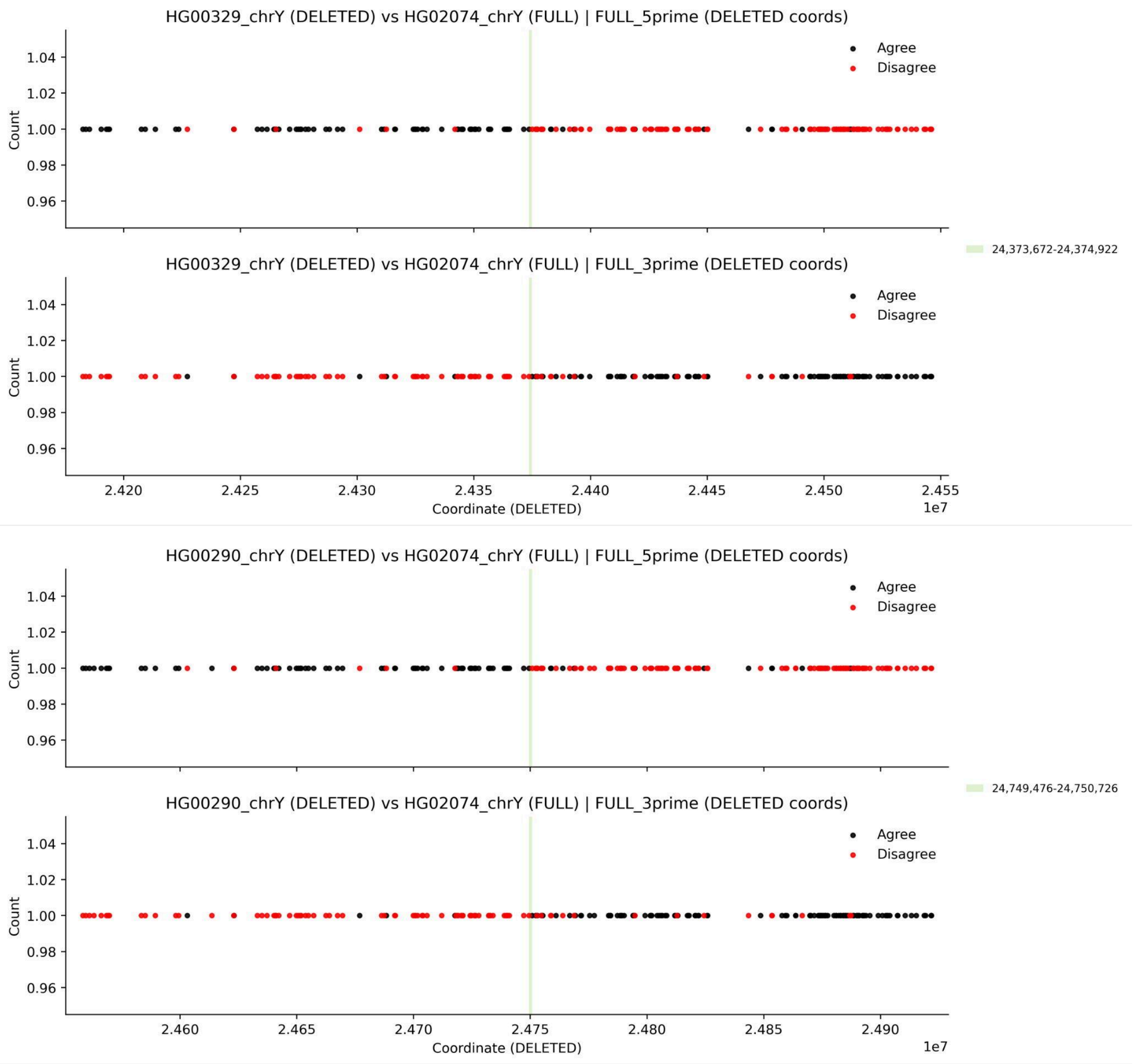


**Supplementary Figure 32. Inferred breakpoint windows for large *AZFc* deletions using SNP concordance with a phylogenetically closest non-deleted neighbor.** For each sample carrying an *AZFc* deletion (*b2/b3* or *gr/gr*; "***DELETED***"), SNP concordance is shown relative to the closest phylogenetic neighbor lacking the deletion ("***FULL***"). Each DELETED-FULL pair is displayed in two panels corresponding to the two deletion breakpoints, using alignments anchored on the 5′ side of the FULL sample (***top***; "***FULL_5prime***") and the 3′ side (***bottom***; "***FULL_3prime***"). SNPs are plotted by coordinates in the DELETED assembly (x-axis) and colored by whether the alleles match between samples (***black***, agree; ***red***, disagree). Shaded bands indicate the inferred deletion breakpoint window(s), with the corresponding breakpoint window coordinates in the DELETED assembly are listed at right.

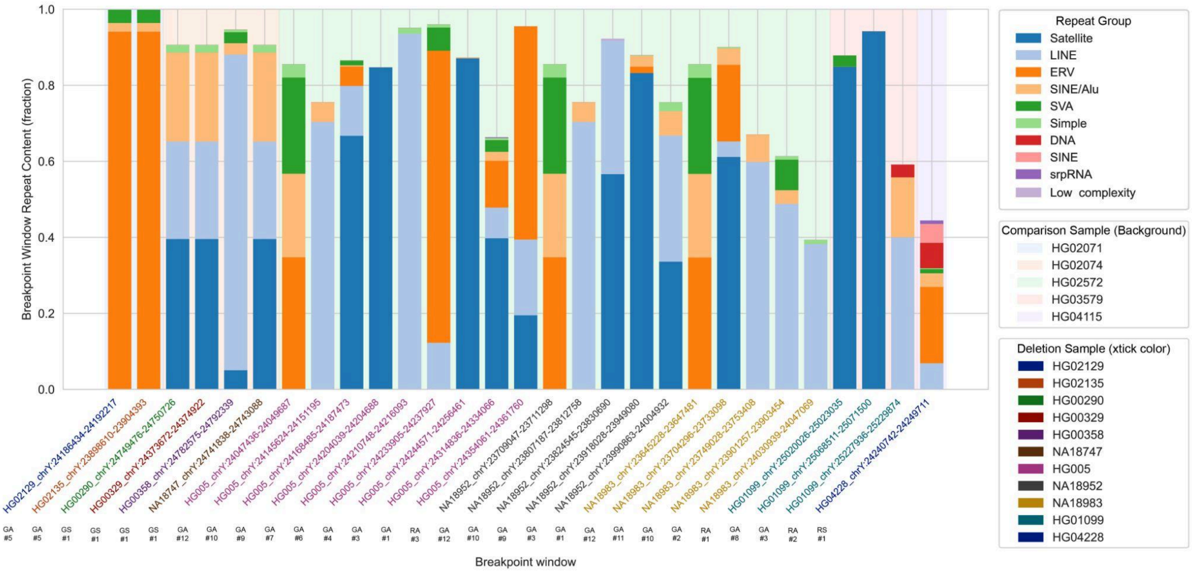


**Supplementary Figure 33. Repeat composition of *AZFc* deletion breakpoint windows.** Stacked bar plots show the repeat content of each inferred deletion breakpoint window. Each bar represents a single breakpoint window, and colored segments indicate the fraction of bases in that window annotated to major repeat groups (RepeatMasker-derived categories). Repeat classes were consolidated into major categories (e.g., ERV, LINE, SINE, SINE/Alu, LTR, DNA (transposon), Satellite, Simple, and low-complexity). Breakpoint windows are ordered by the FULL comparison sample used to localize the breakpoint (background shading), and x-axis label color denotes the DELETED sample from which the breakpoint window was derived. Windows with similar positions relative to their enclosing AZFc colorblock were further binned into four breakpoint classes shown beneath each bar: ***GA***, green antisense; ***GS***, green sense; ***RA***, red antisense; ***RS***, red sense. Overall, breakpoint windows are frequently enriched for satellite and TE sequence.

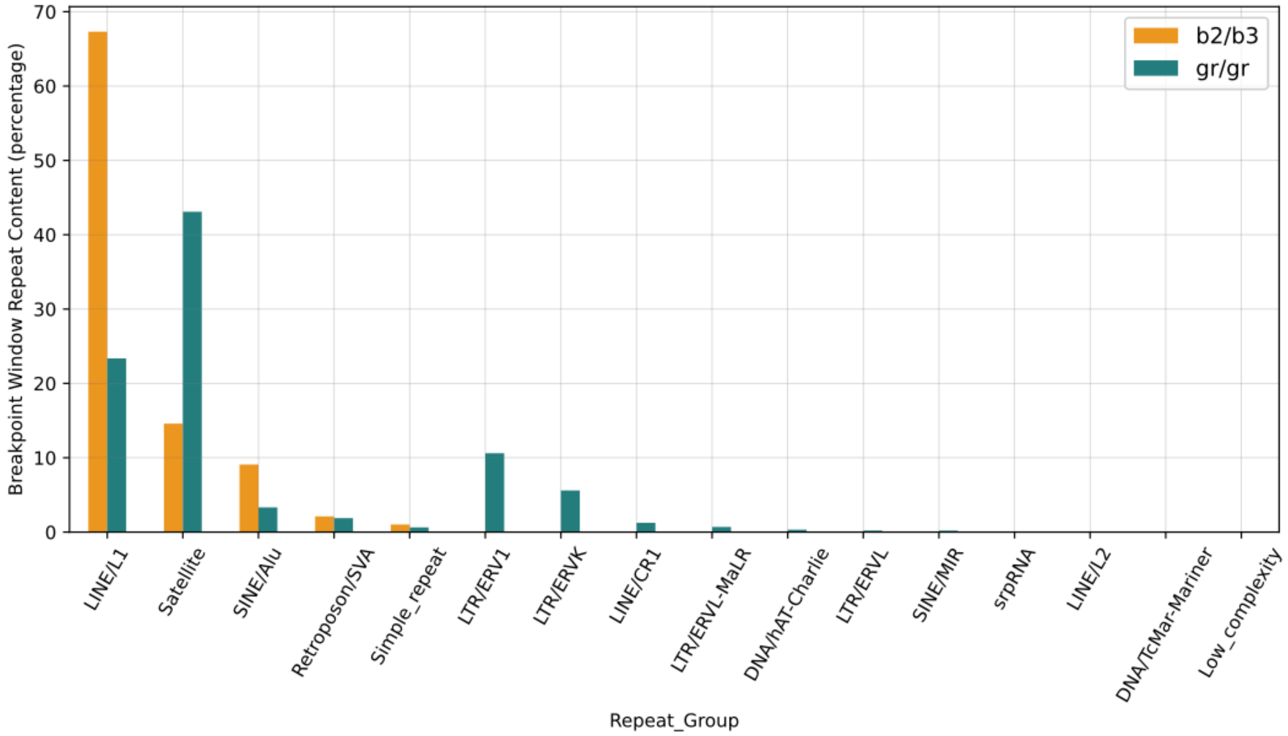


**Supplementary Figure 34. Percentage of repeats within *AZFc* deletion breakpoint windows.** Bar plot showing the total percentage of each Repeat Group present within the breakpoint window across all samples.

*AZFc* constraint modeling

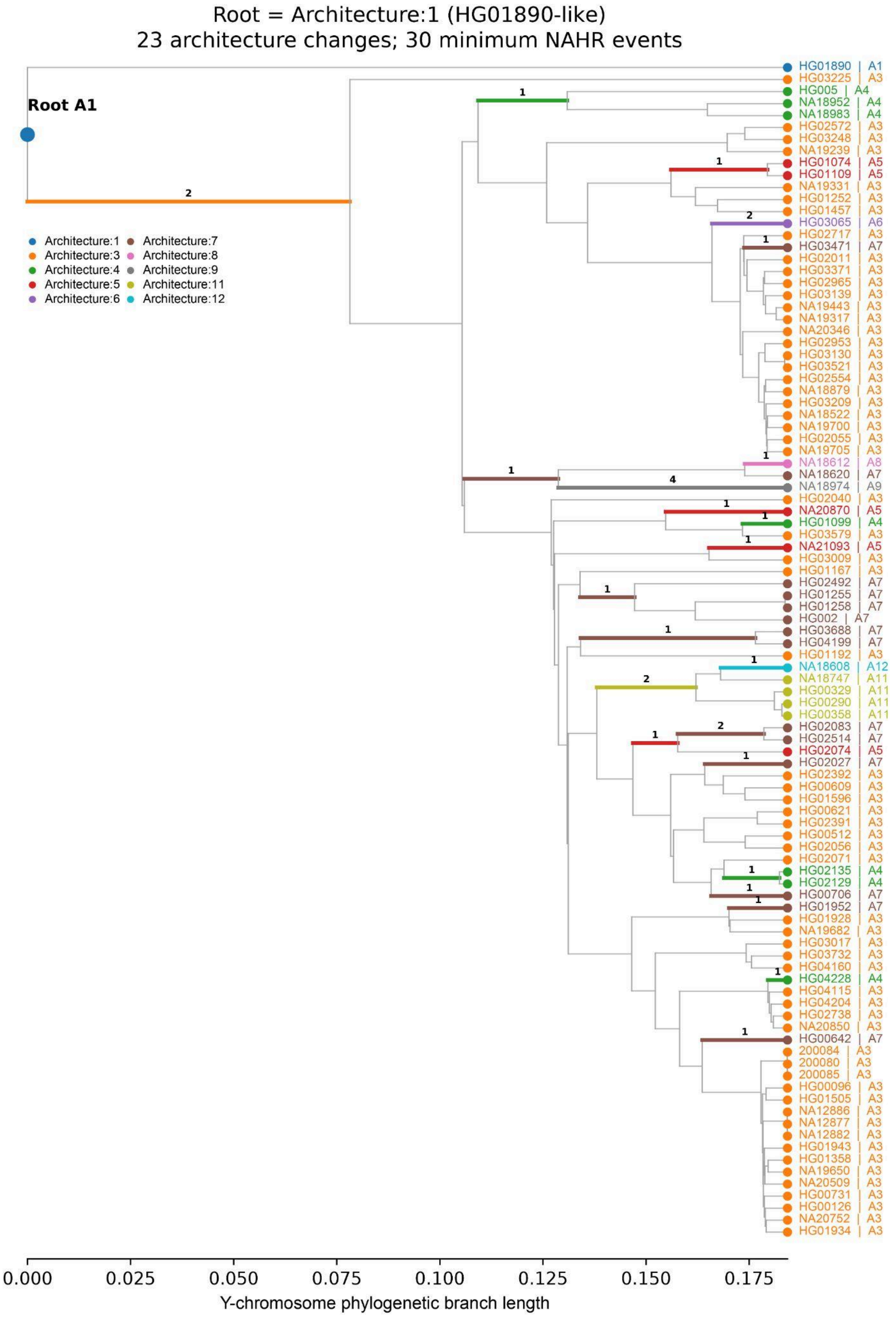
Root = Architecture:1 (HG01890-like)
23 architecture changes; 30 minimum NAHR events
Root A1
Architecture:1
Architecture:3
Architecture:4
Architecture:5
Architecture:6
Architecture:7
Architecture:8
Architecture:9
Architecture:11
Architecture:12
HG01890 | A1
HG03225 | A3
HG005 | A4
NA18952 | A4
NA18983 | A4
HG02572 | A3
HG03248 | A3
NA19239 | A3
HG01074 | A5
HG01109 | A5
NA19331 | A3
HG01252 | A3
HG01457 | A3
HG03065 | A6
HG02717 | A3
HG03471 | A7
HG02011 | A3
HG03371 | A3
HG02965 | A3
HG03139 | A3
NA19443 | A3
NA19317 | A3
NA20346 | A3
HG02953 | A3
HG03130 | A3
HG03521 | A3
HG02554 | A3
NA18879 | A3
HG03209 | A3
NA18522 | A3
NA19700 | A3
HG02055 | A3
NA19705 | A3
NA18612 | A8
NA18620 | A7
NA18974 | A9
HG02040 | A3
NA20870 | A5
HG01099 | A4
HG03579 | A3
NA21093 | A5
HG03009 | A3
HG01167 | A3
HG02492 | A7
HG01255 | A7
HG01258 | A7
HG002 | A7
HG03688 | A7
HG04199 | A7
HG01192 | A3
NA18608 | A12
NA18747 | A11
HG00329 | A11
HG00290 | A11
HG00358 | A11
HG02083 | A7
HG02514 | A7
HG02074 | A5
HG02027 | A7
HG02392 | A3
HG00609 | A3
HG01596 | A3
HG00621 | A3
HG02391 | A3
HG00512 | A3
HG02056 | A3
HG02071 | A3
HG02135 | A4
HG02129 | A4
HG00706 | A7
HG01952 | A7
HG01928 | A3
NA19682 | A3
HG03017 | A3
HG03732 | A3
HG04160 | A3
HG04228 | A4
HG04115 | A3
HG04204 | A3
HG02738 | A3
NA20850 | A3
HG00642 | A7
200084 | A3
200080 | A3
200085 | A3
HG00096 | A3
HG01505 | A3
NA12886 | A3
NA12877 | A3
NA12882 | A3
HG01943 | A3
HG01358 | A3
NA19650 | A3
NA20509 | A3
HG00731 | A3
HG00126 | A3
NA20752 | A3
HG01934 | A3
0.000
0.025
0.050
0.075
0.100
0.125
0.150
0.175
Y-chromosome phylogenetic branch length

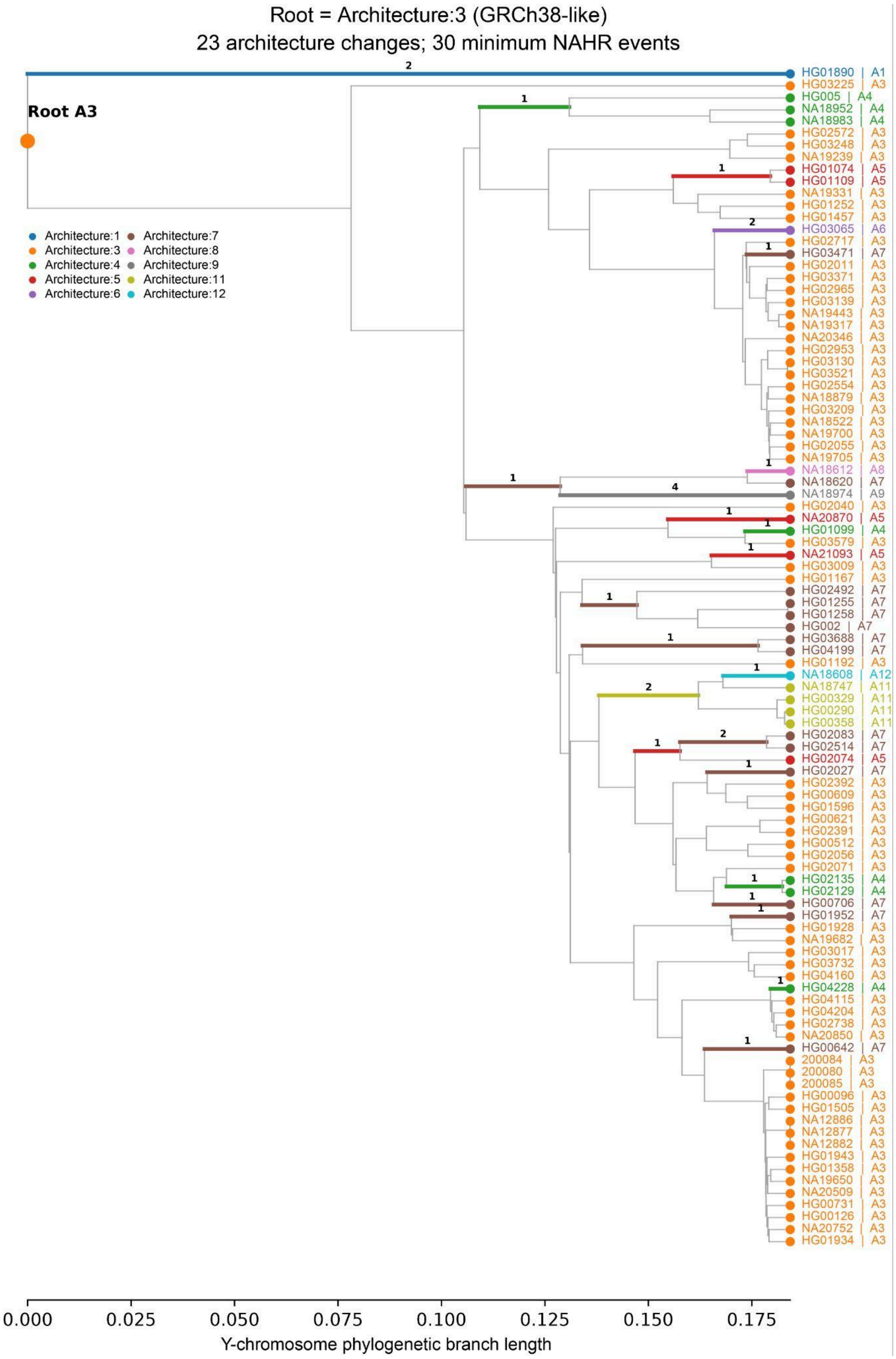
Root = Architecture:3 (GRCh38-like)
23 architecture changes; 30 minimum NAHR events
Root A3
Architecture:1
Architecture:3
Architecture:4
Architecture:5
Architecture:6
Architecture:7
Architecture:8
Architecture:9
Architecture:11
Architecture:12
HG01890 | A1
HG03225 | A3
HG005 | A4
NA18952 | A4
NA18983 | A4
HG02572 | A3
HG03248 | A3
NA19239 | A3
HG01074 | A5
HG01109 | A5
NA19331 | A3
HG01252 | A3
HG01457 | A3
HG03065 | A6
HG02717 | A3
HG03471 | A7
HG02011 | A3
HG03371 | A3
HG02965 | A3
HG03139 | A3
NA19443 | A3
NA19317 | A3
NA20346 | A3
HG02953 | A3
HG03130 | A3
HG03521 | A3
HG02554 | A3
NA18879 | A3
HG03209 | A3
NA18522 | A3
NA19700 | A3
HG02055 | A3
NA19705 | A3
NA18612 | A8
NA18620 | A7
NA18974 | A9
HG02040 | A3
NA20870 | A5
HG01099 | A4
HG03579 | A3
NA21093 | A5
HG03009 | A3
HG01167 | A3
HG02492 | A7
HG01255 | A7
HG01258 | A7
HG002 | A7
HG03688 | A7
HG04199 | A7
HG01192 | A3
NA18608 | A12
NA18747 | A11
HG00329 | A11
HG00290 | A11
HG00358 | A11
HG02083 | A7
HG02514 | A7
HG02074 | A5
HG02027 | A7
HG02392 | A3
HG00609 | A3
HG01596 | A3
HG00621 | A3
HG02391 | A3
HG00512 | A3
HG02056 | A3
HG02071 | A3
HG02135 | A4
HG02129 | A4
HG00706 | A7
HG01952 | A7
HG01928 | A3
NA19682 | A3
HG03017 | A3
HG03732 | A3
HG04160 | A3
HG04228 | A4
HG04115 | A3
HG04204 | A3
HG02738 | A3
NA20850 | A3
HG00642 | A7
200084 | A3
200080 | A3
200085 | A3
HG00096 | A3
HG01505 | A3
NA12886 | A3
NA12877 | A3
NA12882 | A3
HG01943 | A3
HG01358 | A3
NA19650 | A3
NA20509 | A3
HG00731 | A3
HG00126 | A3
NA20752 | A3
HG01934 | A3
0.000
0.025
0.050
0.075
0.100
0.125
0.150
0.175
Y-chromosome phylogenetic branch length

**Supplementary Figure 35. Parsimony reconstruction of *AZFc* architecture under two alternative ancestral states.** Maximum-parsimony reconstruction of *AZFc* architectural states across the 98-chromosome Y phylogeny using the two equally parsimonious ancestral states. **Top,** reconstruction rooted with Architecture:1 (HG01890-like). **Bottom,** reconstruction rooted with Architecture:3 (GRCh38-like). Terminal samples are labeled by sample ID and AZFc architecture (A1–A12), with colors corresponding to the architecture classes shown in the legend. Grey branches indicate phylogenetic relationships without an inferred architecture change, whereas colored branch segments mark reconstructed transitions between AZFc architectures. Numbers above colored branches indicate the minimum number of modeled internal-NAHR events required to connect the reconstructed parental and descendant architectures. Both root reconstructions required 23 architecture-state changes corresponding to a minimum of 30 state-changing internal-NAHR events across the phylogeny. Horizontal branch lengths represent Y-chromosome phylogenetic branch length.

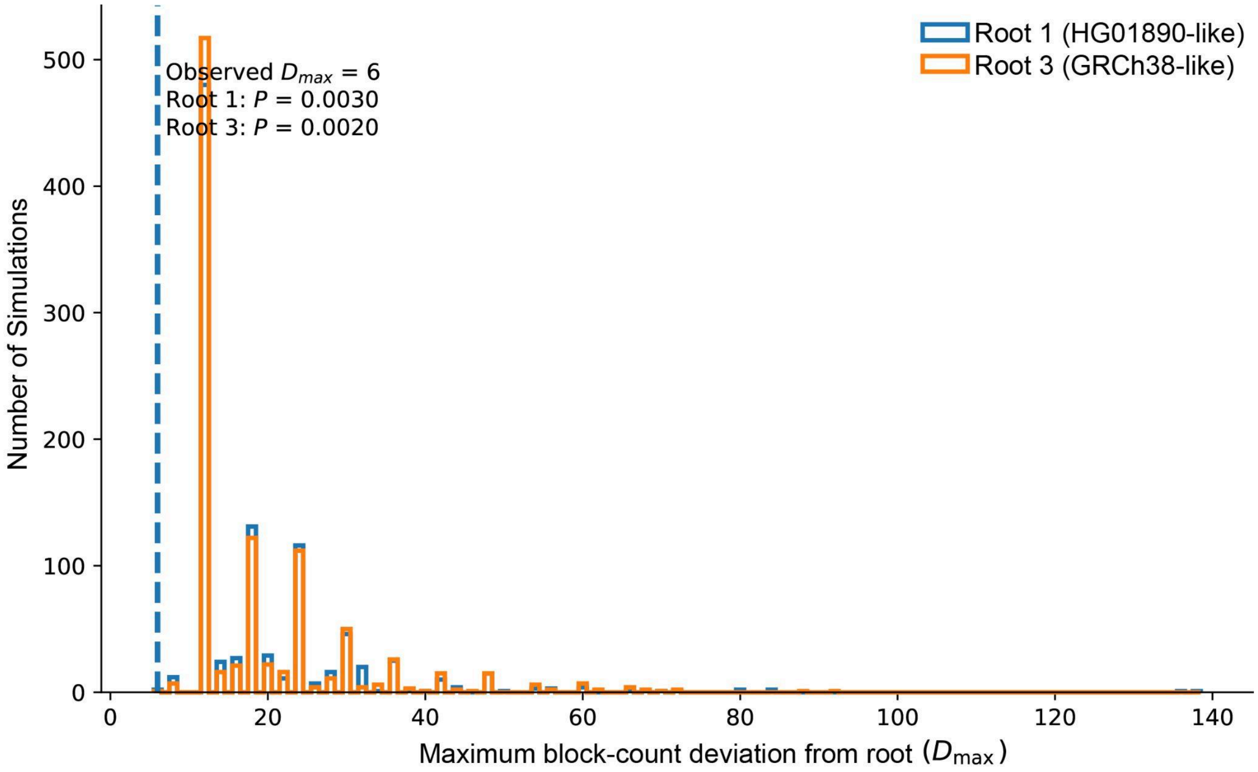


**Supplementary Figure 36. *AZFc* architectures show restricted block-copy excursion under a permissive NAHR model.** Distribution of the maximum absolute terminal block-count deviation from the inferred ancestral state ($D_{max}$) across 1,000 simulations conditioned on exactly 30 state-changing internal-NAHR events. Simulations were performed separately using the two equally parsimonious ancestral states, Architecture:1 (HG01890-like; blue) and Architecture:3 (GRCh38-like; orange), each containing 17 modeled repeat blocks. The dashed vertical line marks the observed $D_{max}$=6, corresponding to the observed range of 11–23 blocks. Only 2/1,000 simulations using Architecture:1 and 1/1,000 using Architecture:3 were as restricted as the observed data, giving one-sided Monte Carlo P=0.0030 and P=0.0020, respectively.

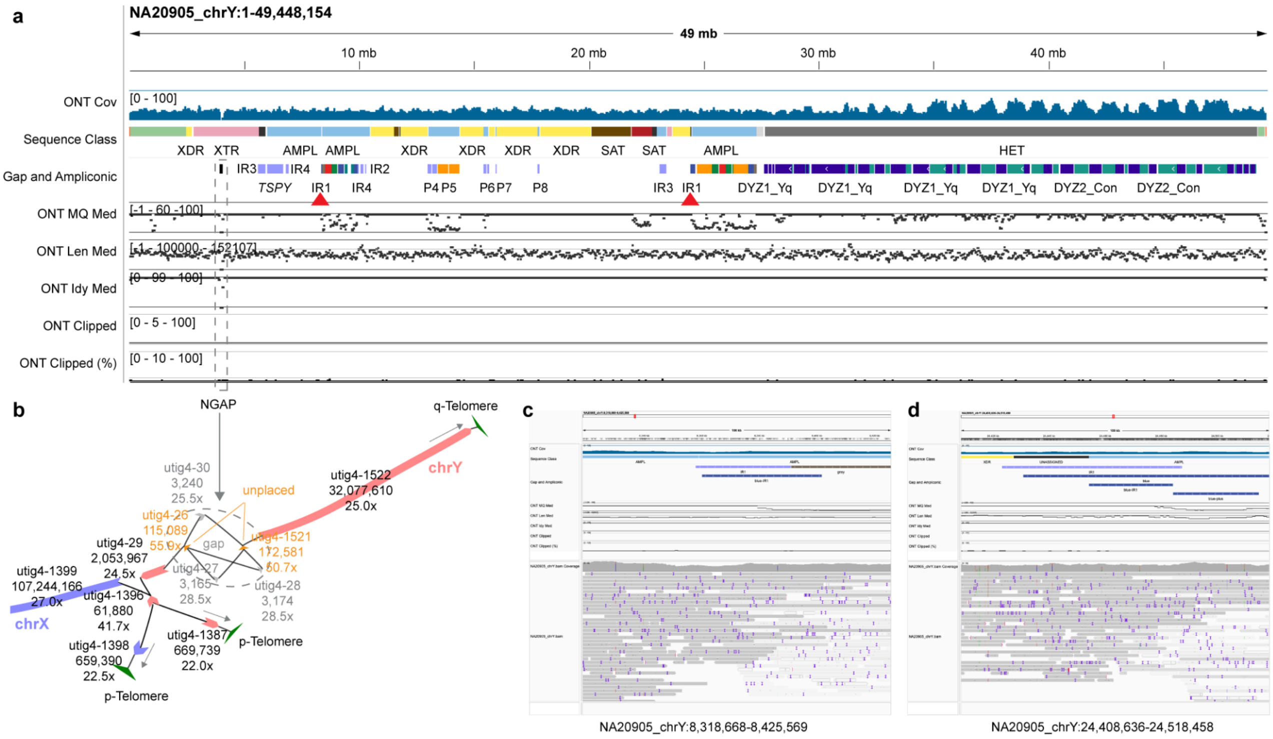


**Supplementary Figure 37. Validation of the NA20905 large inversion. (a)** IGV screen shot of the entire Y chromosome assembly of NA20905, with ONT UL read coverage shown on top. Coverage is mostly even except for the N-gap (NGAP) and Yq12 (HET) region. The Yq12 associated sequencing coverage bias was observed previously in the same DYZ2/HSat1B (green in Gap and Ampliconic track) locus in Rhie et al., 2023[21]. IR1 inversion breakpoints highlighted in **Figure 2C** are marked with red triangles. Median read mapping quality (ONT MQ), read length (ONT Len Med), mapped sequence identity to the assembled sequence (ONT Idy Med) supports even alignment with high MQ in overall. Few regions had MQ drops, including the centromere and in large palindromic arms because the shorter reads were similarly mappable between the partnering palindromic arm. Mid-line shows read length over 100 kb. The number of clipped reads were mostly under 5, less than 10% of the total reads mapped in each 1024 bp window. ONT UL bam file was converted to paf using sam2paf.sh, and read mapping tracks were generated with 'collect_summary.sh in.paf ONT full' from https://github.com/arangrhie/T2T-Polish/tree/master/coverage. **(b)** Verkko assembly graph shows the only unresolved region was a small region in XTR, due to ambiguous path walk in the highlighted NGAP region in panel **a**. **(c, d)** ONT read alignments in the highlighted IR1 breakpoints in panel **a** shows at least 5 clean ultra-long read alignments map across each of the entire IR1 repeat, supporting the integrity of the assembly. IGV screen shots are showing read alignments shaded darker for higher MQ, and visualized in 'consensus mode' hiding indels shorter than 5 bps with mismatched variants shown with allele frequency > 0.2 in the coverage. Exact coordinates of the visualized region are shown below.

### *DAZ* genes

Contributing authors: Mark Loftus, Pille Hallast, Luyao Ren, Gianni Martino, Miriam Konkel, Charles Lee

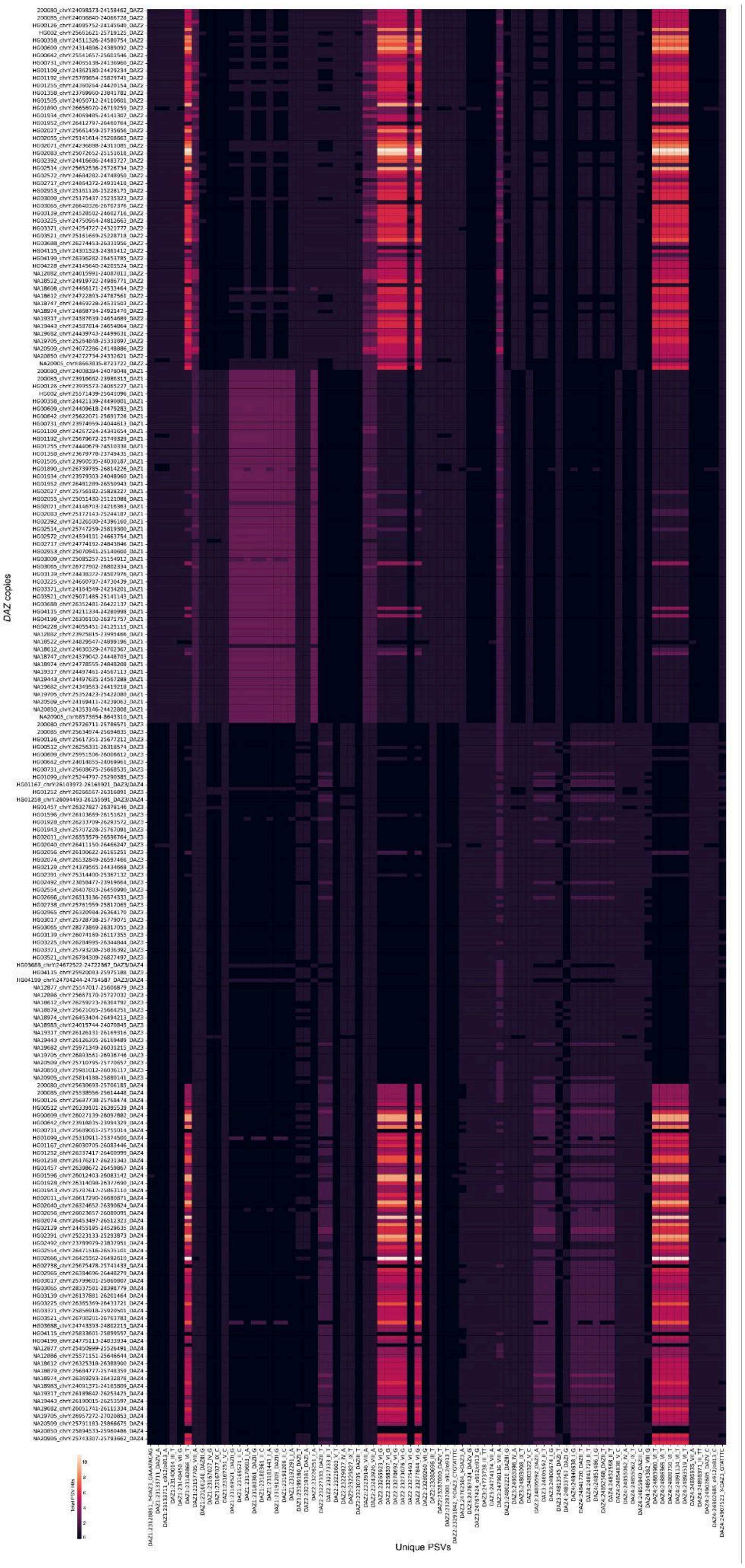

**Supplementary Figure 38. Heatmap of *DAZ* paralogous sequence variant (PSV) profiles.** Heatmap showing the number of full-length, 100% identity mappings of each PSV sequence to each *DAZ* gene copy (only alternate *DAZ* y-axis labels are shown for clarity). PSV-based *DAZ* classifications are appended to the end of each *DAZ* gene label.

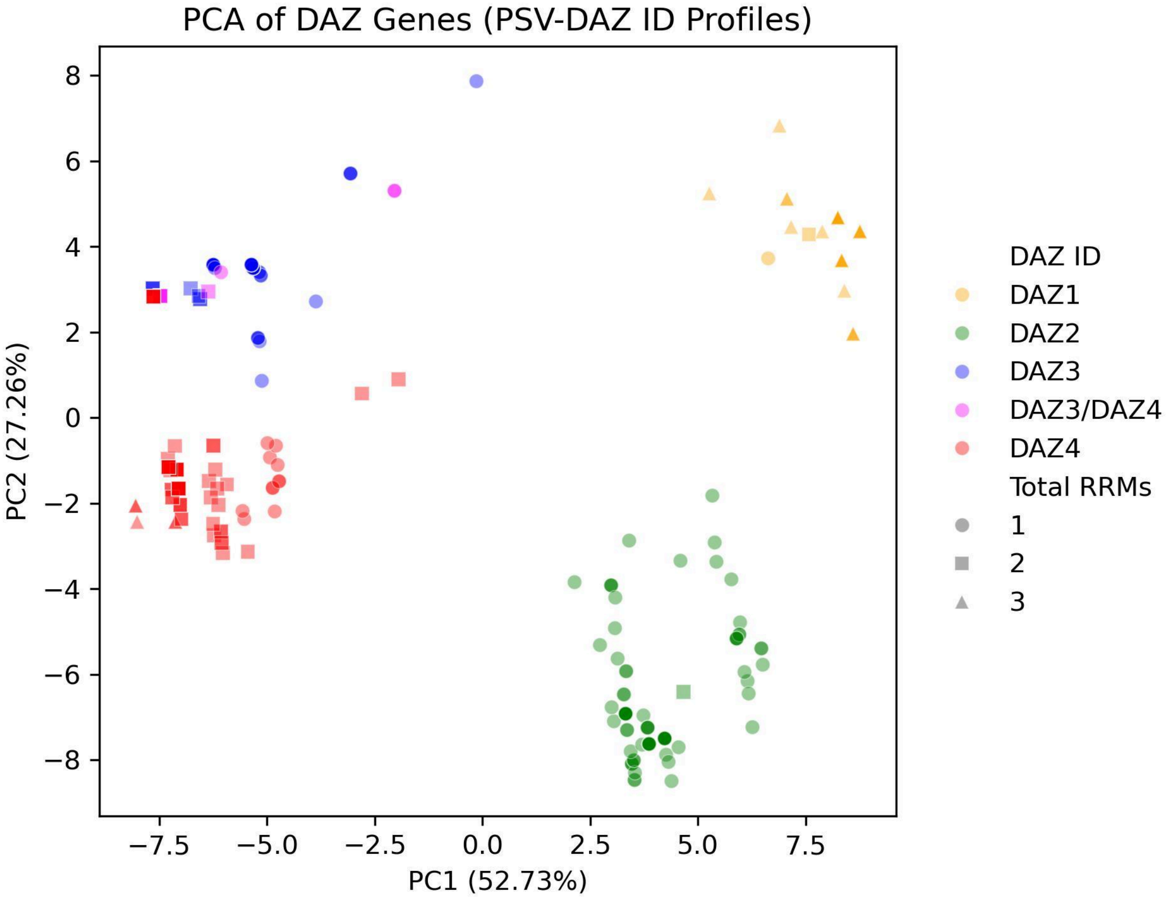


**Supplementary Figure 39. PCA of *DAZ* PSV mapping profiles distinguishes *DAZ* copies and RRM copy number.** Principal component analysis (PCA) was performed on log-transformed and scaled PSV mapping count profiles for individual *DAZ* gene copies (each point represents one *DAZ* copy). Points are colored by PSV-based *DAZ* classification (*DAZ1*, ***orange***; *DAZ2*, ***green***; *DAZ3*, ***blue***; *DAZ3/DAZ4*, ***pink***, *DAZ4*, ***red***). Point shape denotes the number of RNA-recognition motifs (RRMs) encoded by each gene copy (***circle***, 1 RRM; ***square***, 2 RRMs; ***triangle***, 3 RRMs). PC1 and PC2 explain 52.73% and 27.26% of the variance, respectively, separating *DAZ1* and *DAZ2* into distinct clusters while *DAZ3* and *DAZ4* show partial overlap with an intermediate *DAZ3/4* cluster. Hence, it is not always possible to identify which *DAZ* paralog each assembled copy belongs to (and whether it carries 1/2/3 RRMs) from PSV signatures alone.

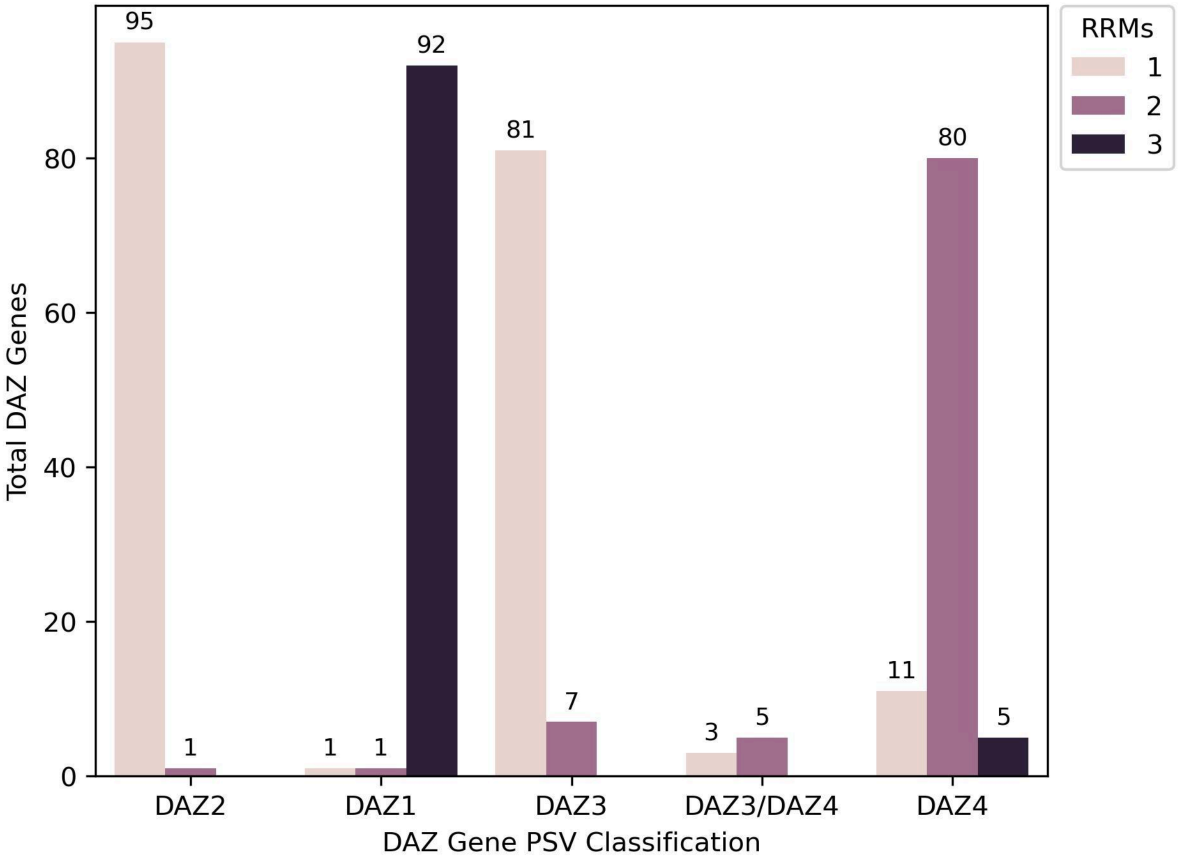


**Supplementary Figure 40. *DAZ* gene classification and RRM copy number.** Bar plot showing PSV-based classifications of all *DAZ* gene copies from Y chromosomes with QC-passed *AZFc* regions. Bars are colored by the number of RNA-recognition motifs (RRMs) encoded by each *DAZ* copy. These results show PSV classification alone does not reliably predict RRM copy number.

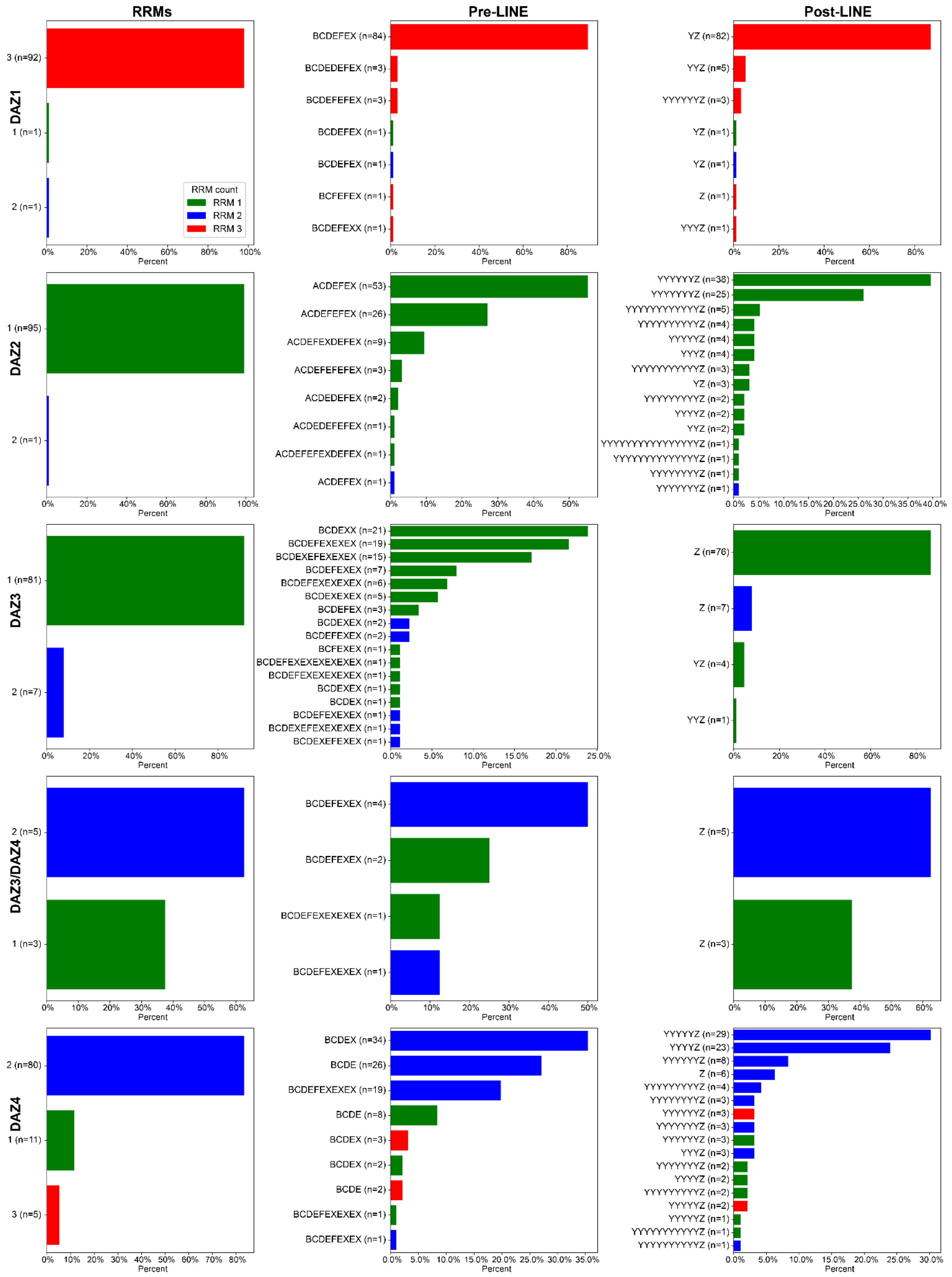


**Supplementary Figure 41. *DAZ* copy composition by PSV classification, RRM content, and flanking repeat pattern haplotypes around the intragenic LINE-1 element.** Rows are stratified by PSV-based *DAZ* gene definition (*DAZ1*, *DAZ2*, *DAZ3*, *DAZ3/DAZ4* ambiguous, and *DAZ4*). ***Left column (RRMs):*** Distribution of RRM copy number among *DAZ* copies in each PSV class. ***Middle***

***and left column (Pre-LINE and post-LINE):*** frequency of repeat pattern either before or after the LINE element. Bars show the percent of *DAZ* copies within each PSV class assigned to each pattern (pattern labels at left; counts shown as ***n***). Bar colors indicate the RRM count encoded by the *DAZ* copy (1 RRM = ***green***, 2 RRMs = ***blue***, 3 RRMs = ***red***).

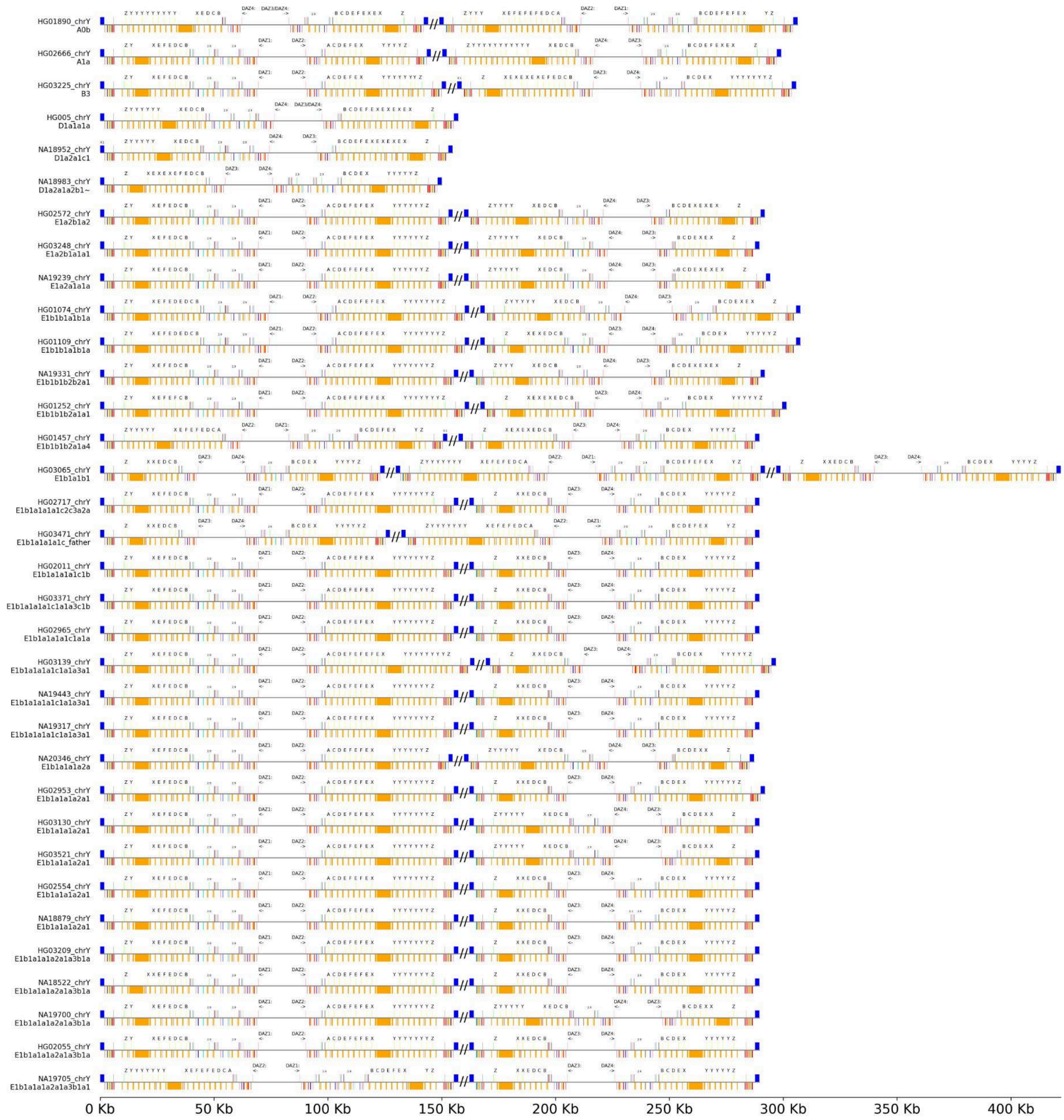
HG01890_chrY A0b
HG02666_chrY A1a
HG03225_chrY B3
HG005_chrY D1a1a1a
NA18952_chrY D1a2a1c1
NA18983_chrY D1a2a1a2b1~
HG02572_chrY E1a2b1a2
HG03248_chrY E1a2b1a1a1
NA19239_chrY E1a2a1a1a
HG01074_chrY E1b1b1a1b1a
HG01109_chrY E1b1b1a1b1a
NA19331_chrY E1b1b1b2b2a1
HG01252_chrY E1b1b1b2a1a1
HG01457_chrY E1b1b1b2a1a4
HG03065_chrY E1b1a1b1
HG02717_chrY E1b1a1a1a1c2c3a2a
HG03471_chrY E1b1a1a1a1c_father
HG02011_chrY E1b1a1a1a1c1b
HG03371_chrY E1b1a1a1a1c1a1a3c1b
HG02965_chrY E1b1a1a1a1c1a1a
HG03139_chrY E1b1a1a1a1c1a1a3a1
NA19443_chrY E1b1a1a1a1c1a1a3a1
NA19317_chrY E1b1a1a1a1c1a1a3a1
NA20346_chrY E1b1a1a1a2a
HG02953_chrY E1b1a1a1a2a1
HG03130_chrY E1b1a1a1a2a1
HG03521_chrY E1b1a1a1a2a1
HG02554_chrY E1b1a1a1a2a1
NA18879_chrY E1b1a1a1a2a1
HG03209_chrY E1b1a1a1a2a1a3b1a
NA18522_chrY E1b1a1a1a2a1a3b1a
NA19700_chrY E1b1a1a1a2a1a3b1a
HG02055_chrY E1b1a1a1a2a1a3b1a
NA19705_chrY E1b1a1a1a2a1a3b1a1
0 Kb
50 Kb
100 Kb
150 Kb
200 Kb
250 Kb
300 Kb
350 Kb
400 Kb

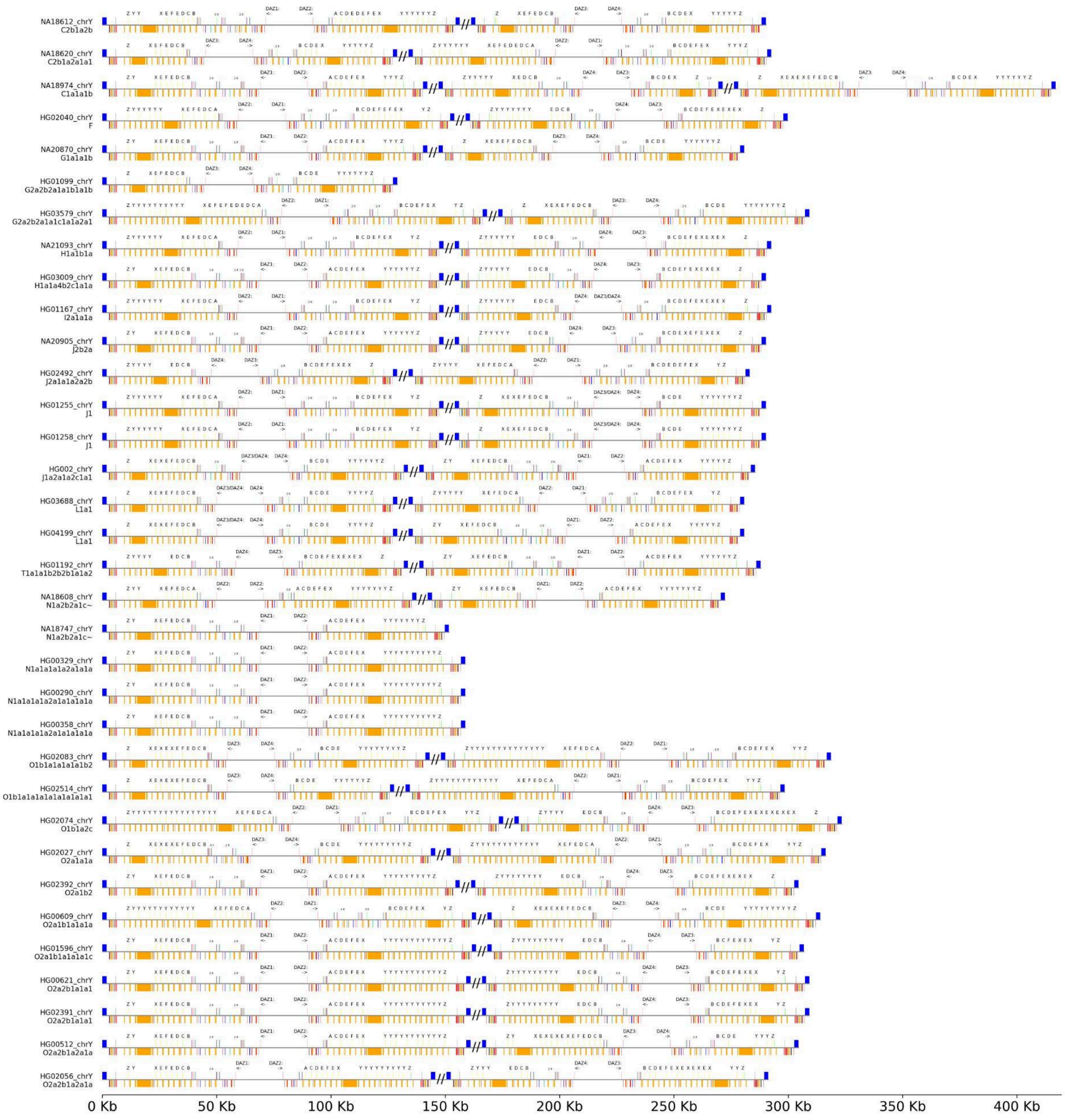
NA18612_chrY
C2b1a2b
NA18620_chrY
C2b1a2a1a1
NA18974_chrY
C1a1a1b
HG02040_chrY
F
NA20870_chrY
G1a1a1b
HG01099_chrY
G2a2b2a1a1b1a1b
HG03579_chrY
G2a2b2a1a1c1a1a2a1
NA21093_chrY
H1a1b1a
HG03009_chrY
H1a1a4b2c1a1a
HG01167_chrY
I2a1a1a
NA20905_chrY
J2b2a
HG02492_chrY
J2a1a1a2a2b
HG01255_chrY
J1
HG01258_chrY
J1
HG002_chrY
J1a2a1a2c1a1
HG03688_chrY
L1a1
HG04199_chrY
L1a1
HG01192_chrY
T1a1a1b2b2b1a1a2
NA18608_chrY
N1a2b2a1c~
NA18747_chrY
N1a2b2a1c~
HG00329_chrY
N1a1a1a1a2a1a1a
HG00290_chrY
N1a1a1a1a2a1a1a1a1a
HG00358_chrY
N1a1a1a1a2a1a1a1a1a
HG02083_chrY
O1b1a1a1a1a1b2
HG02514_chrY
O1b1a1a1a1a1a1a1a1a1
HG02074_chrY
O1b1a2c
HG02027_chrY
O2a1a1a
HG02392_chrY
O2a1b2
HG00609_chrY
O2a1b1a1a1a
HG01596_chrY
O2a1b1a1a1a1c
HG00621_chrY
O2a2b1a1a1
HG02391_chrY
O2a2b1a1a1
HG00512_chrY
O2a2b1a2a1a
HG02056_chrY
O2a2b1a2a1a
0 Kb
50 Kb
100 Kb
150 Kb
200 Kb
250 Kb
300 Kb
350 Kb
400 Kb

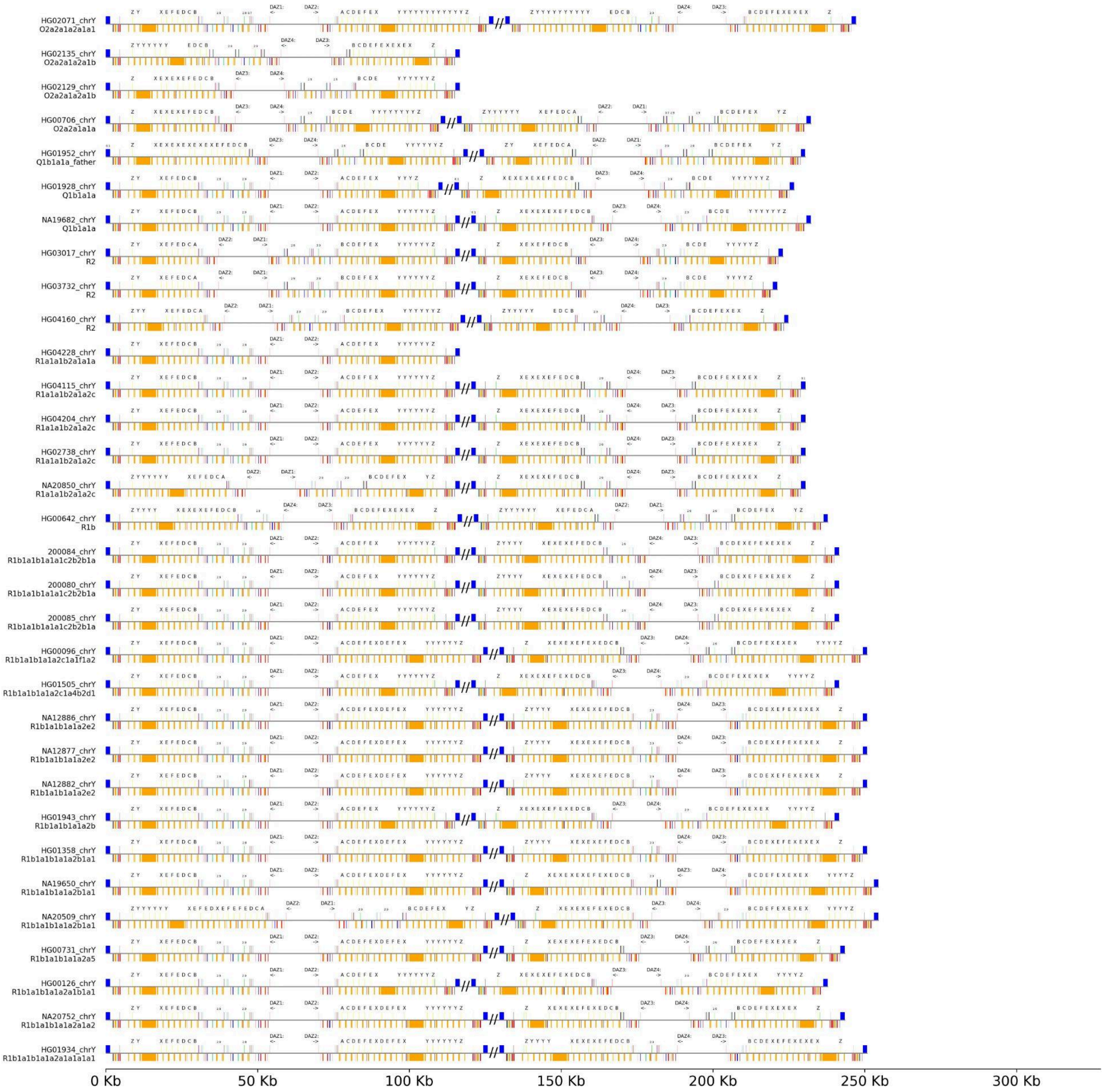


**Supplementary Figure 42. *DAZ* gene exon and repeat architecture.** The above three figures show *DAZ* exon organization and repeat-element composition across all QC-passing samples in the *AZFc* region (n=100). For each *DAZ* copy, exons are displayed along the top of each track and RepeatMasker-annotated repeats along the bottom. Exons are color-coded by type: *DAZ* repeat exons (***yellow***), 5′ UTR/first exon (***pink***), 3′ UTR/last exon (***dark blue***), and RNA-recognition motif (RRM) exons (multiple colors). Gene orientation is indicated by the direction of the arrow beneath the *DAZ* ID (PSV classification) above exon 1. Sample IDs and assigned Y haplogroups are shown on the left, and samples are ordered phylogenetically. *DAZ* repeat exons are labeled by letter following historical nomenclature. For non-*DAZ* repeat exons that deviate from the canonical Ensembl sequence, percent divergence is shown above the exon (most are 0%). To aid visualization, *DAZ* copies are plotted closer together than their true genomic spacing. Diagonal break marks (“//”) indicate regions where inter-copy distances were compressed, reflecting that the *DAZ* copies are separated in the genome and reside in distinct palindromes.

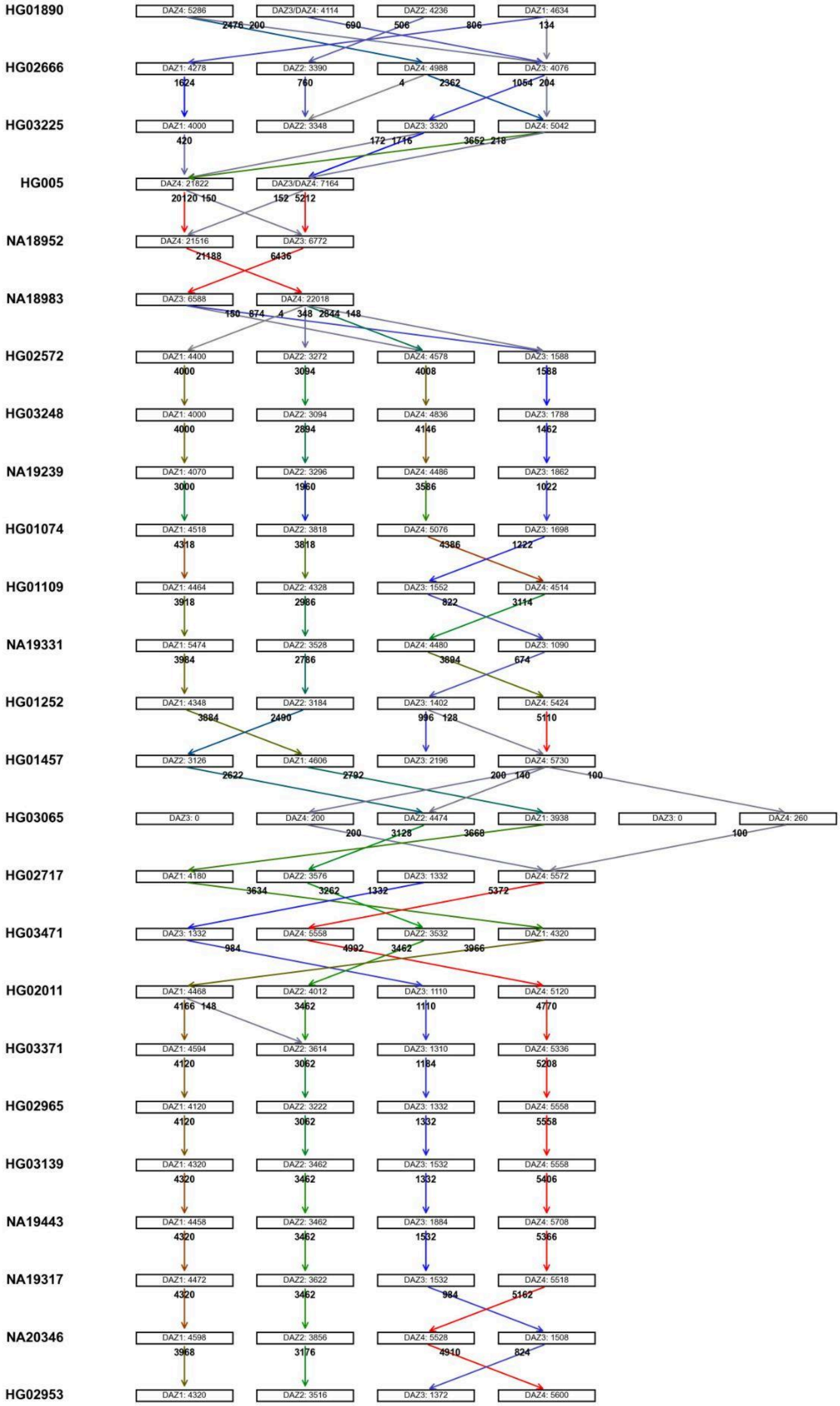
HG01890 DAZ4: 5286 DAZ3/DAZ4: 4114 DAZ2: 4236 DAZ1: 4634
2476 200 690 506 806 134
HG02666 DAZ1: 4278 DAZ2: 3390 DAZ4: 4988 DAZ3: 4076
1624 760 4 2362 1054 204
HG03225 DAZ1: 4000 DAZ2: 3348 DAZ3: 3320 DAZ4: 5042
420 172 1716 3652 218
HG005 DAZ4: 21822 DAZ3/DAZ4: 7164
20120 150 152 5212
NA18952 DAZ4: 21516 DAZ3: 6772
21188 6436
NA18983 DAZ3: 6588 DAZ4: 22018
150 874 4 348 2844 148
HG02572 DAZ1: 4400 DAZ2: 3272 DAZ4: 4578 DAZ3: 1588
4000 3094 4008 1588
HG03248 DAZ1: 4000 DAZ2: 3094 DAZ4: 4836 DAZ3: 1788
4000 2894 4146 1462
NA19239 DAZ1: 4070 DAZ2: 3296 DAZ4: 4486 DAZ3: 1862
3000 1960 3586 1022
HG01074 DAZ1: 4518 DAZ2: 3818 DAZ4: 5076 DAZ3: 1698
4318 3818 4386 1222
HG01109 DAZ1: 4464 DAZ2: 4328 DAZ3: 1552 DAZ4: 4514
3918 2986 822 3114
NA19331 DAZ1: 5474 DAZ2: 3528 DAZ4: 4480 DAZ3: 1090
3984 2786 3894 674
HG01252 DAZ1: 4348 DAZ2: 3184 DAZ3: 1402 DAZ4: 5424
3884 2490 996 128 5110
HG01457 DAZ2: 3126 DAZ1: 4606 DAZ3: 2196 DAZ4: 5730
2622 2792 200 140 100
HG03065 DAZ3: 0 DAZ4: 200 DAZ2: 4474 DAZ1: 3938 DAZ3: 0 DAZ4: 260
200 3128 3668 100
HG02717 DAZ1: 4180 DAZ2: 3576 DAZ3: 1332 DAZ4: 5572
3634 3262 1332 5372
HG03471 DAZ3: 1332 DAZ4: 5558 DAZ2: 3532 DAZ1: 4320
984 4992 3462 3966
HG02011 DAZ1: 4468 DAZ2: 4012 DAZ3: 1110 DAZ4: 5120
4166 148 3462 1110 4770
HG03371 DAZ1: 4594 DAZ2: 3614 DAZ3: 1310 DAZ4: 5336
4120 3062 1184 5208
HG02965 DAZ1: 4120 DAZ2: 3222 DAZ3: 1332 DAZ4: 5558
4120 3062 1332 5558
HG03139 DAZ1: 4320 DAZ2: 3462 DAZ3: 1532 DAZ4: 5558
4320 3462 1332 5406
NA19443 DAZ1: 4458 DAZ2: 3462 DAZ3: 1884 DAZ4: 5708
4320 3462 1532 5366
NA19317 DAZ1: 4472 DAZ2: 3622 DAZ3: 1532 DAZ4: 5518
4320 3462 984 5162
NA20346 DAZ1: 4598 DAZ2: 3856 DAZ4: 5528 DAZ3: 1508
3968 3176 4910 824
HG02953 DAZ1: 4320 DAZ2: 3516 DAZ3: 1372 DAZ4: 5600

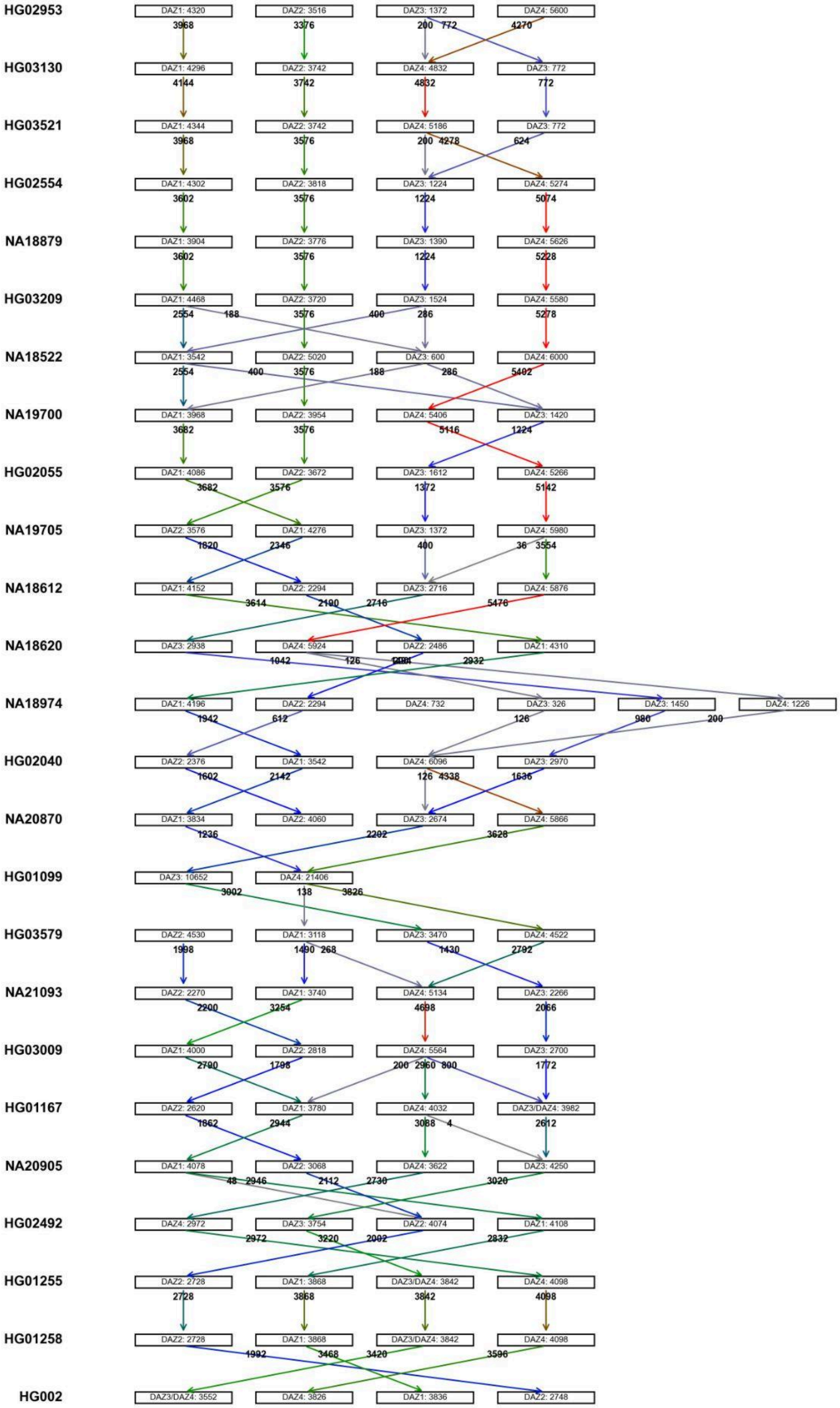

HG02953
DAZ1: 4320
DAZ2: 3516
DAZ3: 1372
DAZ4: 5600
HG03130
DAZ1: 4296
DAZ2: 3742
DAZ4: 4832
DAZ3: 772
HG03521
DAZ1: 4344
DAZ2: 3742
DAZ4: 5186
DAZ3: 772
HG02554
DAZ1: 4302
DAZ2: 3818
DAZ3: 1224
DAZ4: 5274
NA18879
DAZ1: 3904
DAZ2: 3776
DAZ3: 1390
DAZ4: 5626
HG03209
DAZ1: 4468
DAZ2: 3720
DAZ3: 1524
DAZ4: 5580
NA18522
DAZ1: 3542
DAZ2: 5020
DAZ3: 600
DAZ4: 6000
NA19700
DAZ1: 3968
DAZ2: 3954
DAZ4: 5406
DAZ3: 1420
HG02055
DAZ1: 4086
DAZ2: 3672
DAZ3: 1612
DAZ4: 5266
NA19705
DAZ2: 3576
DAZ1: 4276
DAZ3: 1372
DAZ4: 5980
NA18612
DAZ1: 4152
DAZ2: 2294
DAZ3: 2716
DAZ4: 5876
NA18620
DAZ3: 2938
DAZ4: 5924
DAZ2: 2486
DAZ1: 4310
NA18974
DAZ1: 4196
DAZ2: 2294
DAZ4: 732
DAZ3: 326
DAZ3: 1450
DAZ4: 1226
HG02040
DAZ2: 2376
DAZ1: 3542
DAZ4: 6096
DAZ3: 2970
NA20870
DAZ1: 3834
DAZ2: 4060
DAZ3: 2674
DAZ4: 5866
HG01099
DAZ3: 10652
DAZ4: 21406
HG03579
DAZ2: 4530
DAZ1: 3118
DAZ3: 3470
DAZ4: 4522
NA21093
DAZ2: 2270
DAZ1: 3740
DAZ4: 5134
DAZ3: 2266
HG03009
DAZ1: 4000
DAZ2: 2818
DAZ4: 5564
DAZ3: 2700
HG01167
DAZ2: 2620
DAZ1: 3780
DAZ4: 4032
DAZ3/DAZ4: 3982
NA20905
DAZ1: 4078
DAZ2: 3068
DAZ4: 3622
DAZ3: 4250
HG02492
DAZ4: 2972
DAZ3: 3754
DAZ2: 4074
DAZ1: 4108
HG01255
DAZ2: 2728
DAZ1: 3868
DAZ3/DAZ4: 3842
DAZ4: 4098
HG01258
DAZ2: 2728
DAZ1: 3868
DAZ3/DAZ4: 3842
DAZ4: 4098
HG002
DAZ3/DAZ4: 3552
DAZ4: 3826
DAZ1: 3836
DAZ2: 2748

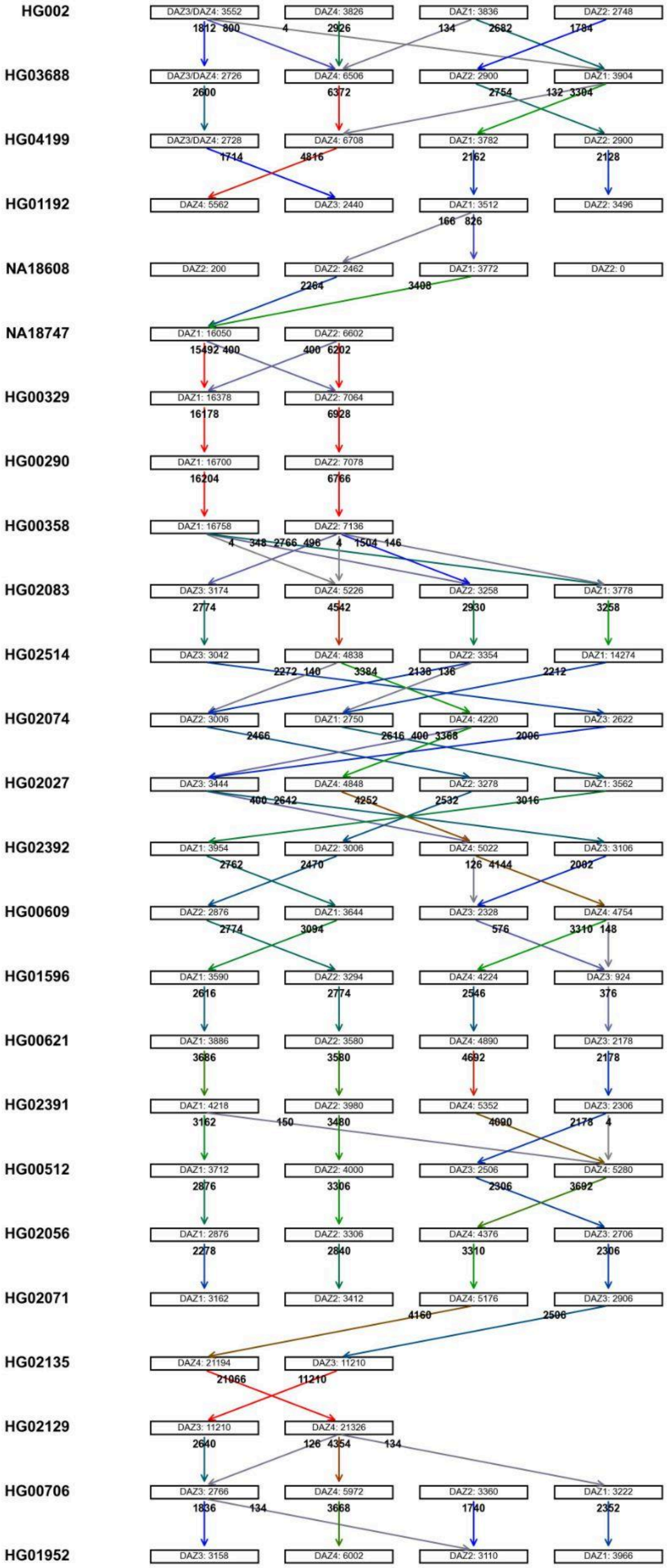
HG002
DAZ3/DAZ4: 3552
DAZ4: 3826
DAZ1: 3836
DAZ2: 2748
HG03688
DAZ3/DAZ4: 2726
DAZ4: 6506
DAZ2: 2900
DAZ1: 3904
HG04199
DAZ3/DAZ4: 2728
DAZ4: 6708
DAZ1: 3782
DAZ2: 2900
HG01192
DAZ4: 5562
DAZ3: 2440
DAZ1: 3512
DAZ2: 3496
NA18608
DAZ2: 200
DAZ2: 2462
DAZ1: 3772
DAZ2: 0
NA18747
DAZ1: 16050
DAZ2: 6602
HG00329
DAZ1: 16378
DAZ2: 7064
HG00290
DAZ1: 16700
DAZ2: 7078
HG00358
DAZ1: 16758
DAZ2: 7136
HG02083
DAZ3: 3174
DAZ4: 5226
DAZ2: 3258
DAZ1: 3778
HG02514
DAZ3: 3042
DAZ4: 4838
DAZ2: 3354
DAZ1: 14274
HG02074
DAZ2: 3006
DAZ1: 2750
DAZ4: 4220
DAZ3: 2622
HG02027
DAZ3: 3444
DAZ4: 4848
DAZ2: 3278
DAZ1: 3562
HG02392
DAZ1: 3954
DAZ2: 3006
DAZ4: 5022
DAZ3: 3106
HG00609
DAZ2: 2876
DAZ1: 3644
DAZ3: 2328
DAZ4: 4754
HG01596
DAZ1: 3590
DAZ2: 3294
DAZ4: 4224
DAZ3: 924
HG00621
DAZ1: 3886
DAZ2: 3580
DAZ4: 4890
DAZ3: 2178
HG02391
DAZ1: 4218
DAZ2: 3980
DAZ4: 5352
DAZ3: 2306
HG00512
DAZ1: 3712
DAZ2: 4000
DAZ3: 2506
DAZ4: 5280
HG02056
DAZ1: 2876
DAZ2: 3306
DAZ4: 4376
DAZ3: 2706
HG02071
DAZ1: 3162
DAZ2: 3412
DAZ4: 5176
DAZ3: 2906
HG02135
DAZ4: 21194
DAZ3: 11210
HG02129
DAZ3: 11210
DAZ4: 21326
HG00706
DAZ3: 2766
DAZ4: 5972
DAZ2: 3360
DAZ1: 3222
HG01952
DAZ3: 3158
DAZ4: 6002
DAZ2: 3110
DAZ1: 3966

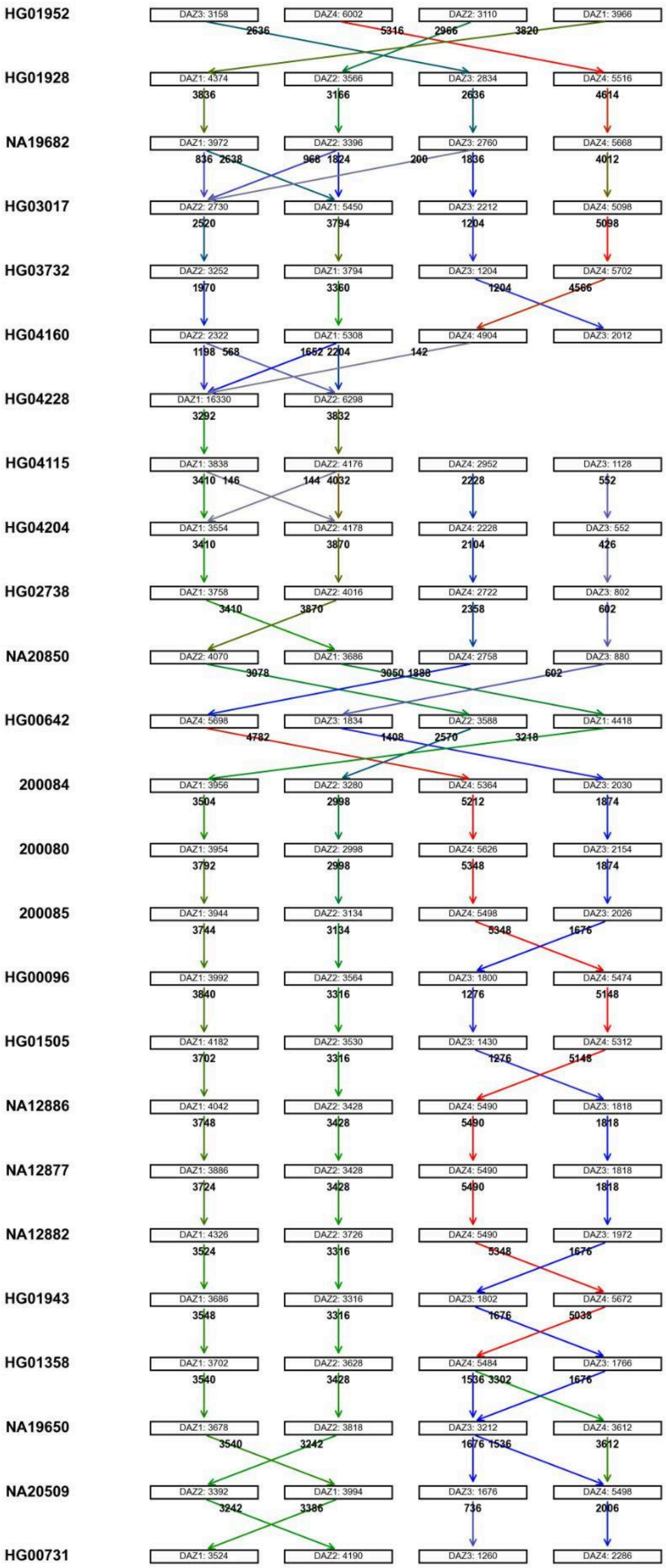
HG01952
DAZ3: 3158
DAZ4: 6002
DAZ2: 3110
DAZ1: 3966
2636 5316 2966 3820
HG01928
DAZ1: 4374
DAZ2: 3566
DAZ3: 2834
DAZ4: 5516
3836 3166 2636 4614
NA19682
DAZ1: 3972
DAZ2: 3396
DAZ3: 2760
DAZ4: 5668
836 2638 968 1824 200 1836 4012
HG03017
DAZ2: 2730
DAZ1: 5450
DAZ3: 2212
DAZ4: 5098
2520 3794 1204 5098
HG03732
DAZ2: 3252
DAZ1: 3794
DAZ3: 1204
DAZ4: 5702
1970 3360 1204 4566
HG04160
DAZ2: 2322
DAZ1: 5308
DAZ4: 4904
DAZ3: 2012
1198 568 1652 2204 142
HG04228
DAZ1: 16330
DAZ2: 6298
3292 3832
HG04115
DAZ1: 3838
DAZ2: 4176
DAZ4: 2952
DAZ3: 1128
3410 146 144 4032 2228 552
HG04204
DAZ1: 3554
DAZ2: 4178
DAZ4: 2228
DAZ3: 552
3410 3870 2104 426
HG02738
DAZ1: 3758
DAZ2: 4016
DAZ4: 2722
DAZ3: 802
3410 3870 2358 602
NA20850
DAZ2: 4070
DAZ1: 3686
DAZ4: 2758
DAZ3: 880
3078 3050 1888 602
HG00642
DAZ4: 5698
DAZ3: 1834
DAZ2: 3588
DAZ1: 4418
4782 1408 2570 3218
200084
DAZ1: 3956
DAZ2: 3280
DAZ4: 5364
DAZ3: 2030
3504 2998 5212 1874
200080
DAZ1: 3954
DAZ2: 2998
DAZ4: 5626
DAZ3: 2154
3792 2998 5348 1874
200085
DAZ1: 3944
DAZ2: 3134
DAZ4: 5498
DAZ3: 2026
3744 3134 5348 1676
HG00096
DAZ1: 3992
DAZ2: 3564
DAZ3: 1800
DAZ4: 5474
3840 3316 1276 5148
HG01505
DAZ1: 4182
DAZ2: 3530
DAZ3: 1430
DAZ4: 5312
3702 3316 1276 5148
NA12886
DAZ1: 4042
DAZ2: 3428
DAZ4: 5490
DAZ3: 1818
3748 3428 5490 1818
NA12877
DAZ1: 3886
DAZ2: 3428
DAZ4: 5490
DAZ3: 1818
3724 3428 5490 1818
NA12882
DAZ1: 4326
DAZ2: 3726
DAZ4: 5490
DAZ3: 1972
3524 3316 5348 1676
HG01943
DAZ1: 3686
DAZ2: 3316
DAZ3: 1802
DAZ4: 5672
3548 3316 1676 5038
HG01358
DAZ1: 3702
DAZ2: 3628
DAZ4: 5484
DAZ3: 1766
3540 3428 1536 3302 1676
NA19650
DAZ1: 3678
DAZ2: 3818
DAZ3: 3212
DAZ4: 3612
3540 3242 1676 1536 3612
NA20509
DAZ2: 3392
DAZ1: 3994
DAZ3: 1676
DAZ4: 5498
3242 3386 736 2006
HG00731
DAZ1: 3524
DAZ2: 4190
DAZ3: 1260
DAZ4: 2286

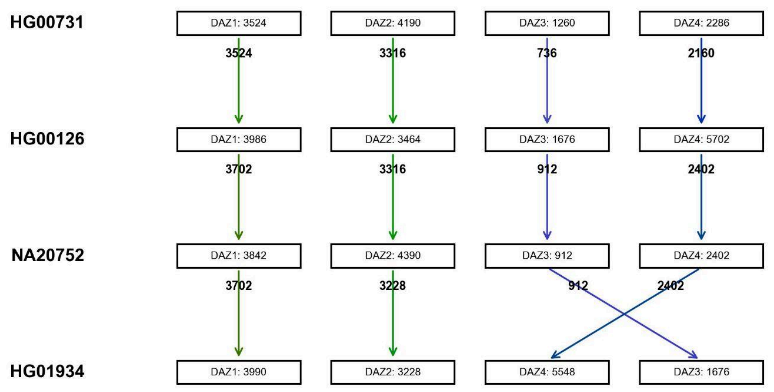


**Supplementary Figure 43. *DAZ* unique kmer trajectory.** The above figures show *DAZ* gene copies for each sample along with the number of gene-unique 100-bp k-mers. Samples are ordered phylogenetically and labeled on the left. Arrows connect *DAZ* copies between adjacent samples in the phylogeny: an arrow from an upper gene to the gene below indicates that k-mers unique to the upper gene in a sample are also found uniquely in the lower sample gene copy. Numbers within *DAZ* rectangles and on arrows report the total unique k-mers and the number shared under this criterion, respectively. Arrow color encodes the abundance of shared sample unique k-mers (low, ***gray***; low–medium, ***blue***; medium–high, ***green***; high, ***red***).

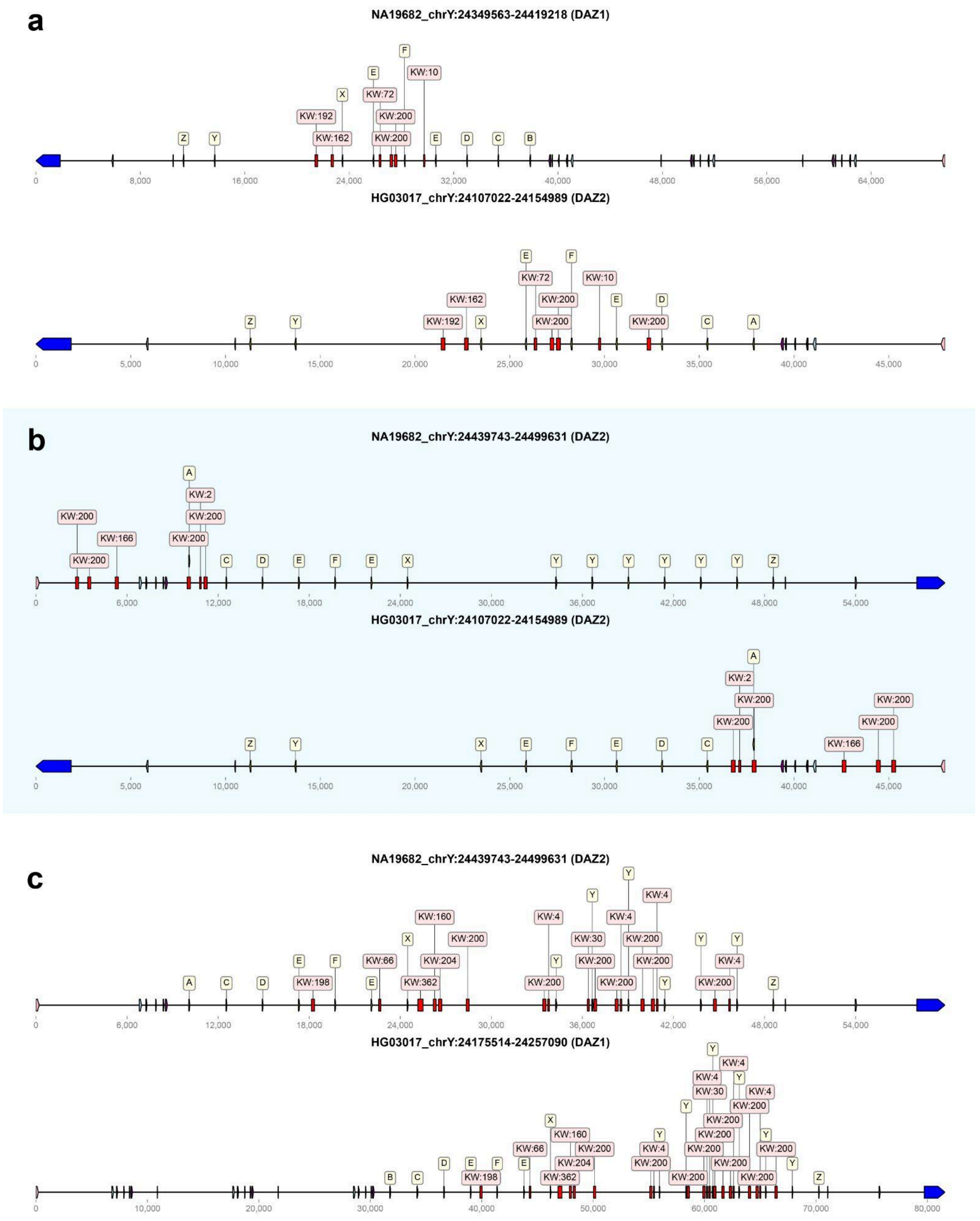


**Supplementary Figure 44. Examples of *DAZ* inversions inferred from shared gene unique k-mer windows.** Three pairwise comparisons illustrate how shared k-mer windows (KWs) can localize an inversion breakpoint between *DAZ* copies. KWs are genomic windows that contain one or more 100-bp k-mers unique to a given *DAZ* copy. KW positions are plotted along each *DAZ* gene model (exons labeled; arrow indicates transcriptional orientation). **a, *DAZ1* and DAZ2 comparison.** The two *DAZ* copies share a matching set of KWs concentrated near the RNA recognition motif

(RRM) exons. **b, *DAZ2* comparison.** The two *DAZ2* copies similarly share KWs near the RRM exons. **c, *DAZ2* and *DAZ1* comparison.** In NA19682, the *DAZ2* repeat-exon region carries the same KW pattern observed in *DAZ1* from HG03017, indicating a segment exchange consistent with an inversion. Together, these KW correspondences localize the inferred inversion breakpoint to the *DAZ* repeat-exon block between the C/D/E repeat exons.

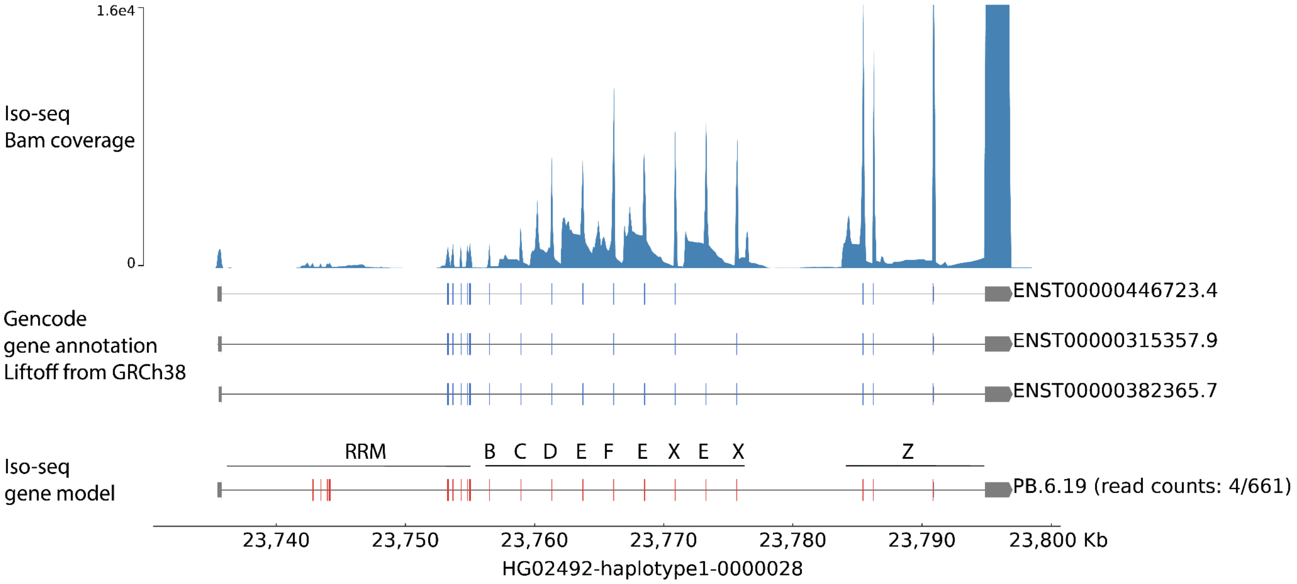


**Supplementary Figure 45. Expression of transcripts compatible with a hybrid *DAZ* architecture.** Testis Iso-seq data support expression of a hybrid *DAZ* gene in which the *DAZ3* copy acquired a second RRM motif through an inversion event involving *DAZ4*, resulting in a reciprocal *DAZ4* copy with one RRM instead of two. Shown are Iso-seq BAM coverage (top), GENCODE gene annotations (middle), and supported Iso-seq gene models (bottom), mapped to the HG02492 (haplogroup J2a1a1a2a2b) Y-chromosome assembly.

### *TSPY* and *RBMY* gene families

Contributing authors: Mark Loftus, Pille Hallast, Tomoya Kanno, Gianni Martino, Miriam Konkel, Mindy Shi

#### *TSPY* array and genes

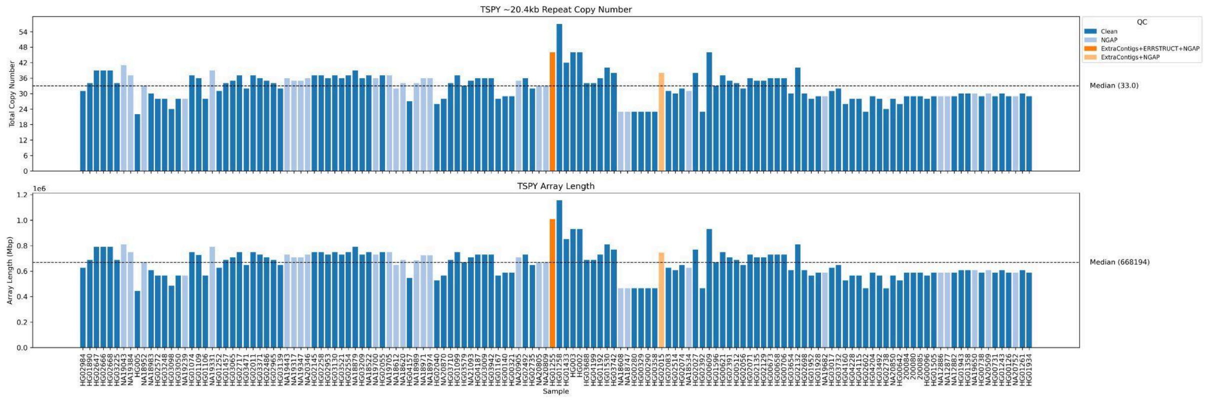


**Supplementary Figure 46. *TSPY* array size and QC status across samples.** Bar plots summarize the TSPY tandem array in 140 Y chromosome assemblies. **Top:** estimated copy number of the 20.4 kb *TSPY* repeat unit per sample (median = 33). **Bottom:** corresponding assembled *TSPY* array length (median = 668,194 bp). Bars are colored by QC status: ***blue***, clean arrays with no QC flags; ***light blue***, arrays containing an internal gap of Ns (NGAP); ***dark orange***, arrays with NGAP plus an ERRSTRUCT structural QC flag and additional *TSPY* copies detected on non-primary contigs (EXTRACONTIGS); ***light orange***, arrays with NGAP and extra-contig *TSPY* copies but no ERRSTRUCT flag. Samples are ordered phylogenetically, with African lineages beginning on the left. Overall, most samples have 30-40 *TSPY* repeats, but a subset show unusually large arrays and/or assembly/QC flags suggesting fragmented or uncertain array reconstruction.

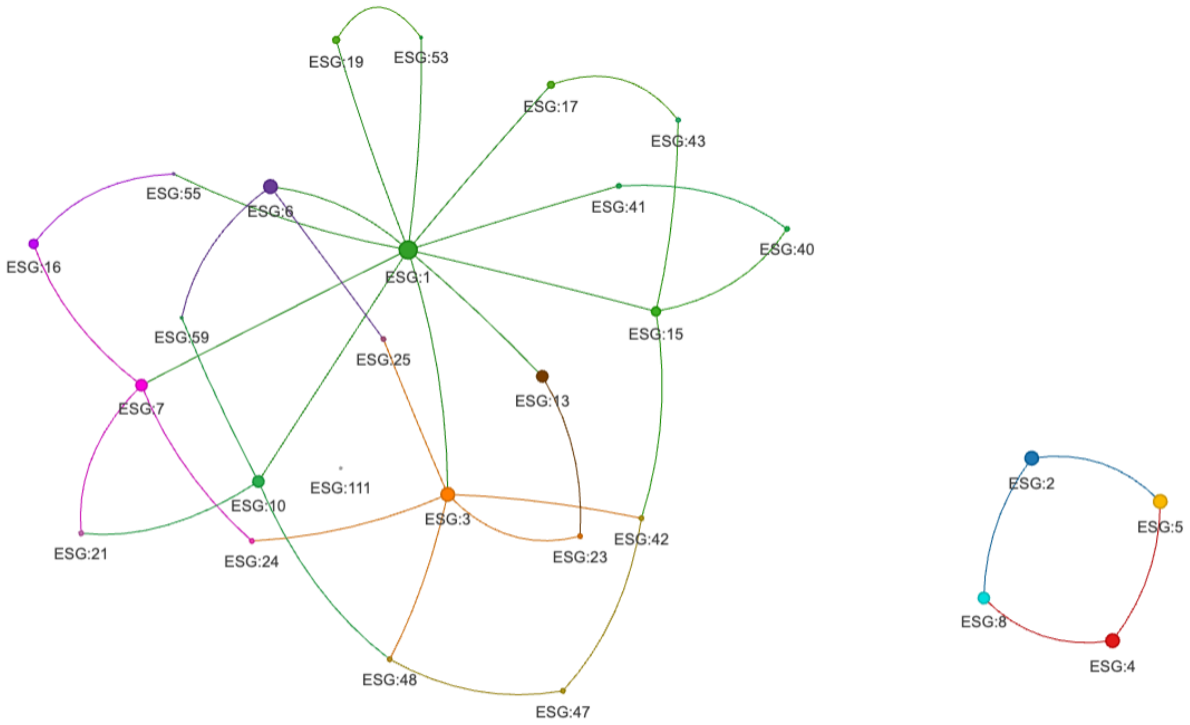


**Supplementary Figure 47. The *TSPY* amino acid sequence network.** Nodes represent unique *TSPY* amino-acid Exact Sequence Groups (ESGs). Undirected edges connect each ESG to its nearest neighbor(s) in sequence space, defined as the ESG(s) with minimum Hamming distance (ties allowed). The nine highest-degree ESGs were designated hub nodes and assigned unique colors; all other nodes are colored by hub assignment based on shortest path proximity. Node size is proportional to the total number of *TSPY* copies associated with each ESG across all samples. ESG identifiers are arbitrary labels and carried forward from earlier analyses and do not indicate rank or inferred evolutionary order. Therefore, *TSPY* protein sequences form a few common “hub” variants that account for many copies, surrounded by many rarer variants that differ by only a small number of amino acid changes.

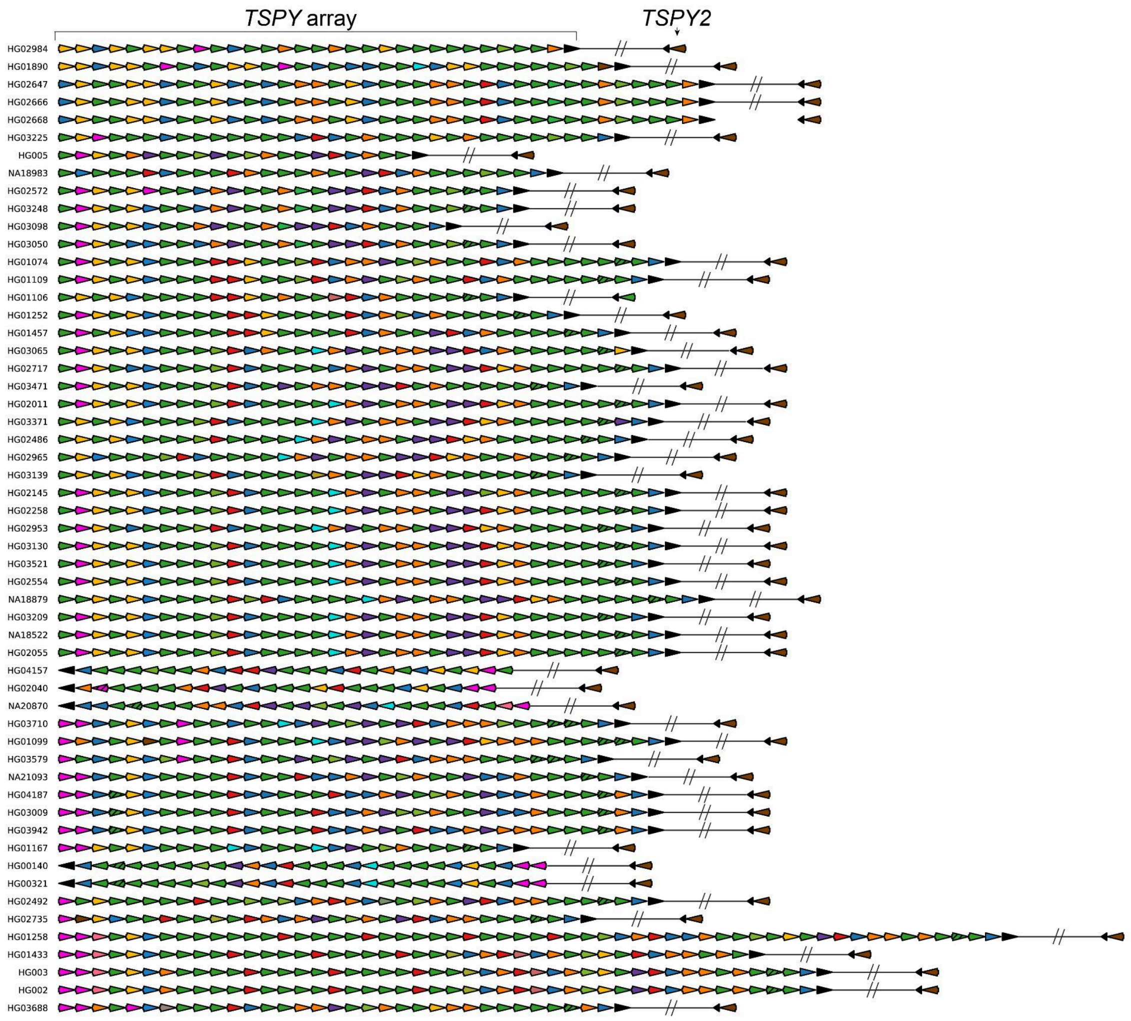

TSPY array
TSPY2
HG02984
HG01890
HG02647
HG02666
HG02668
HG03225
HG005
NA18983
HG02572
HG03248
HG03098
HG03050
HG01074
HG01109
HG01106
HG01252
HG01457
HG03065
HG02717
HG03471
HG02011
HG03371
HG02486
HG02965
HG03139
HG02145
HG02258
HG02953
HG03130
HG03521
HG02554
NA18879
HG03209
NA18522
HG02055
HG04157
HG02040
NA20870
HG03710
HG01099
HG03579
NA21093
HG04187
HG03009
HG03942
HG01167
HG00140
HG00321
HG02492
HG02735
HG01258
HG01433
HG003
HG002
HG03688

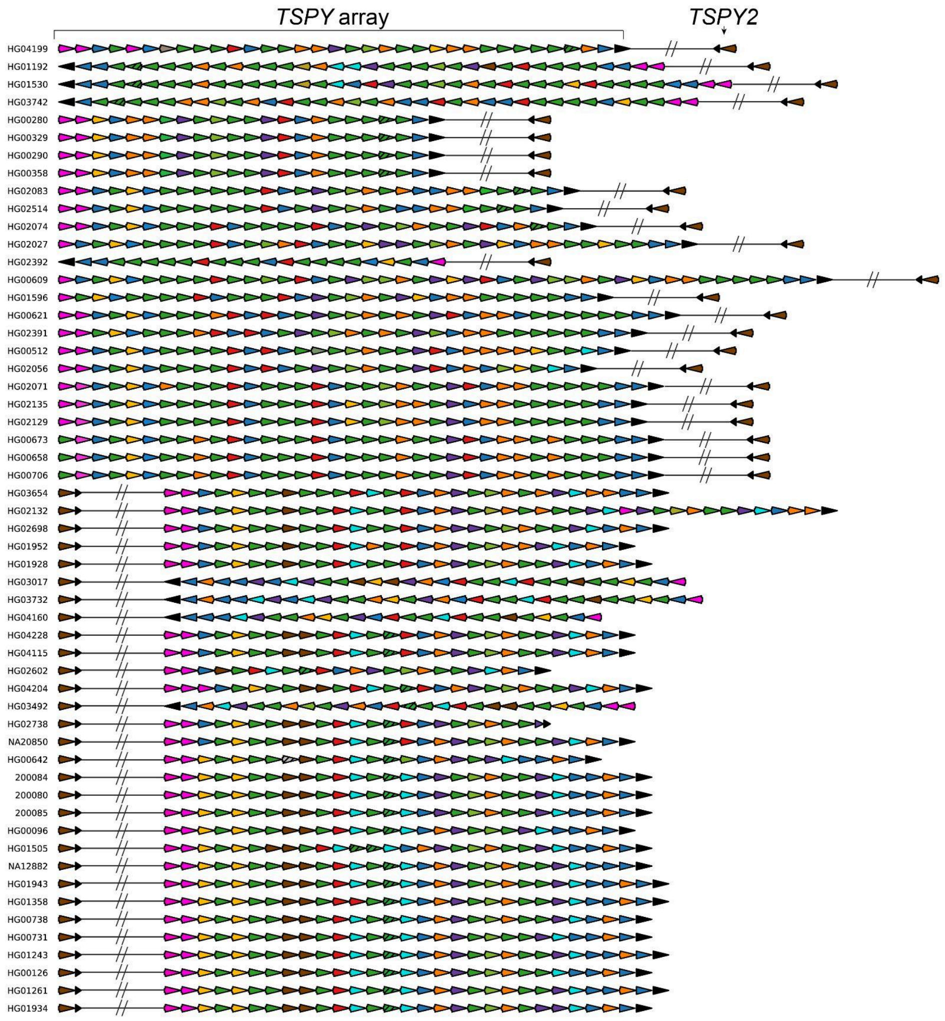


**Supplementary Figure 48. Organization and coding diversity of *TSPY* genes.** For each QC-passing sample (n=110; rows ordered by phylogeny with haplogroup A at top), arrows depict individual *TSPY* gene copies in assembly order within the tandem *TSPY* array (***left***) and the single *TSPY2* copy located outside the array (*right*). Arrow direction indicates gene orientation. Copy colors denote amino acid Exact Sequence Group (ESG) membership as defined in the *TSPY* sequence network analysis. Copies containing premature stop codons (putative pseudogenes) are indicated with black striped overlays. The slash "//" separating *TSPY2* from the *TSPY* array denotes a distance >100,000 kb (not to scale). Across individuals, the overall tandem-array architecture is broadly conserved, while the amino-acid sequence composition of copies varies substantially within and between haplogroups.

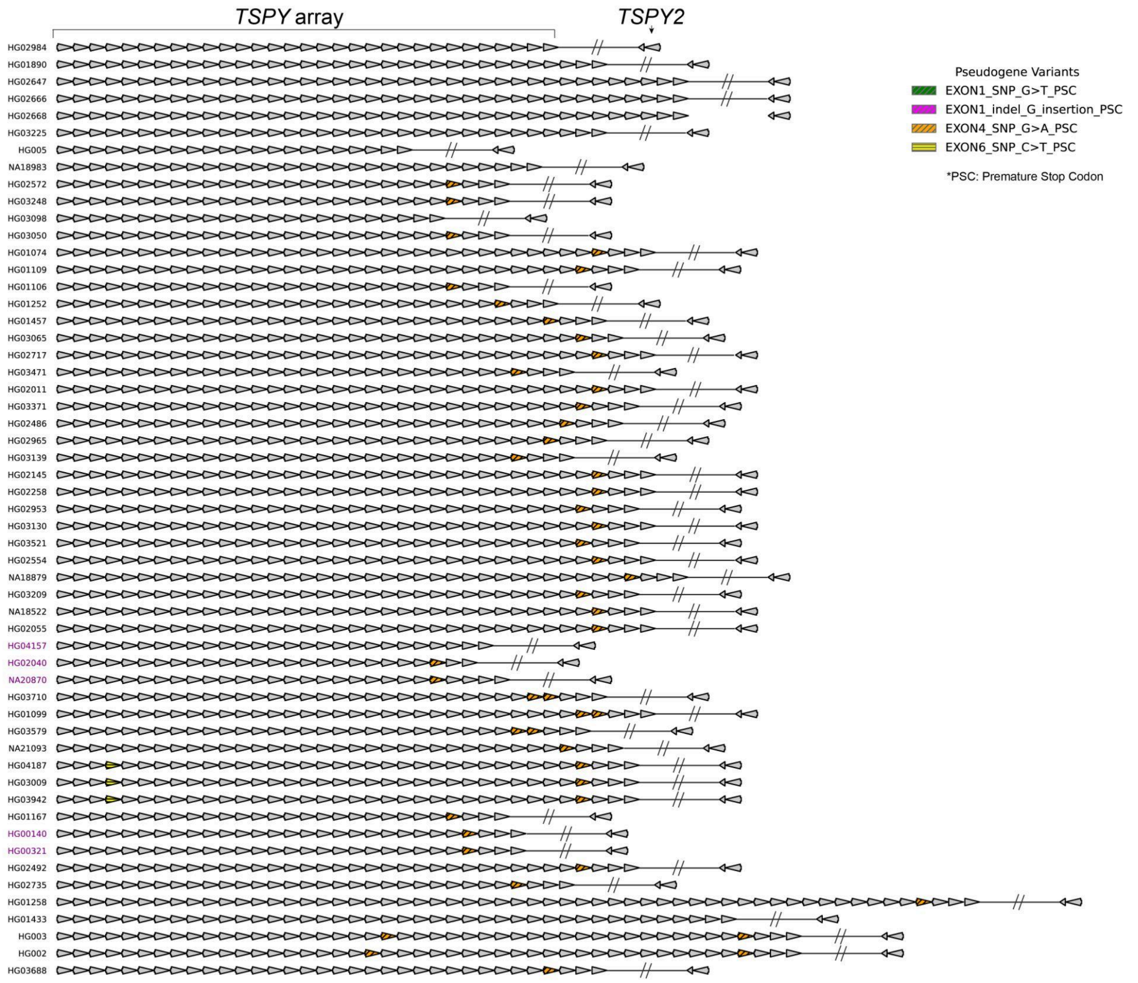
TSPY array
TSPY2
HG02984
HG01890
HG02647
HG02666
HG02668
HG03225
HG005
NA18983
HG02572
HG03248
HG03098
HG03050
HG01074
HG01109
HG01106
HG01252
HG01457
HG03065
HG02717
HG03471
HG02011
HG03371
HG02486
HG02965
HG03139
HG02145
HG02258
HG02953
HG03130
HG03521
HG02554
NA18879
HG03209
NA18522
HG02055
HG04157
HG02040
NA20870
HG03710
HG01099
HG03579
NA21093
HG04187
HG03009
HG03942
HG01167
HG00140
HG00321
HG02492
HG02735
HG01258
HG01433
HG003
HG002
HG03688
Pseudogene Variants
EXON1_SNP_G>T_PSC
EXON1_indel_G_insertion_PSC
EXON4_SNP_G>A_PSC
EXON6_SNP_C>T_PSC
*PSC: Premature Stop Codon

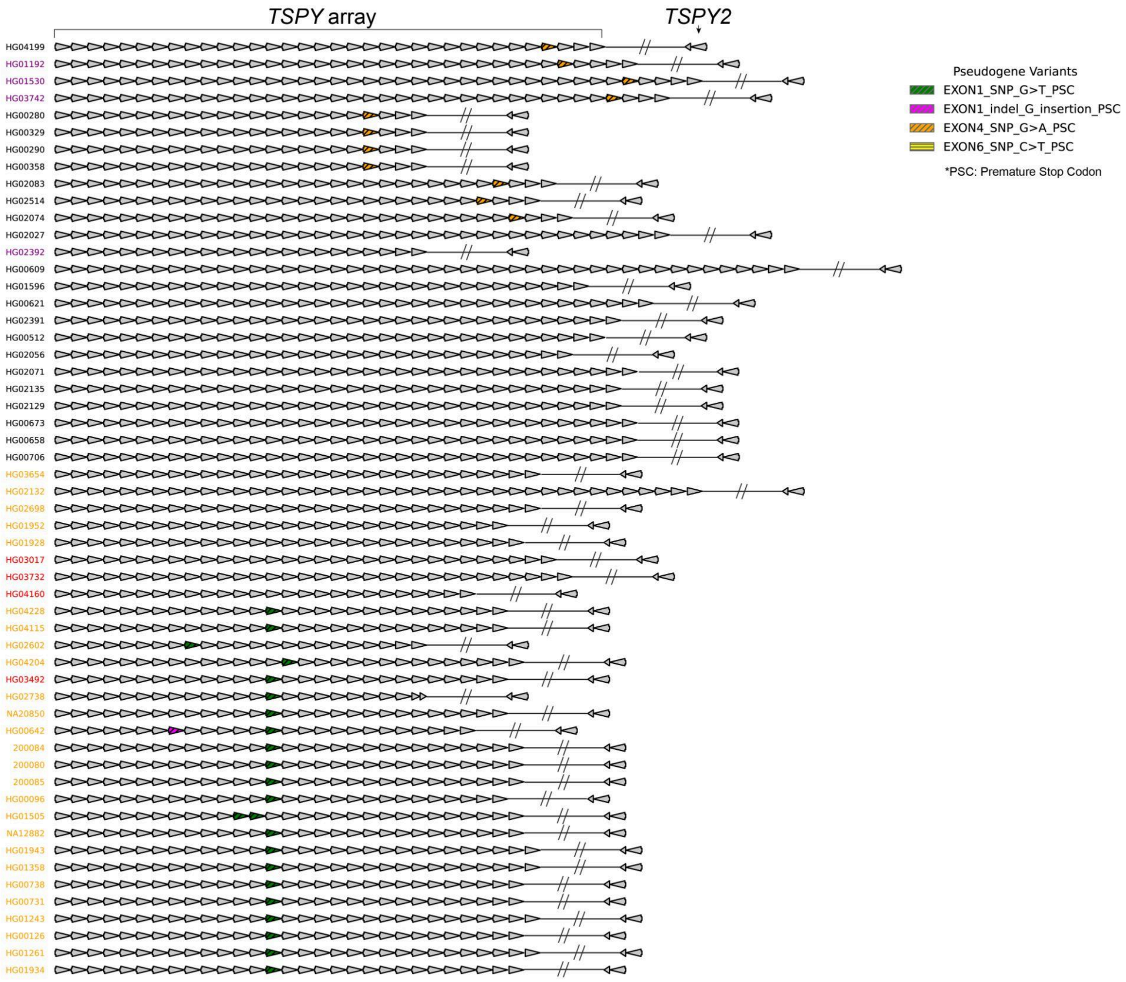


**Supplementary Figure 49. Putative *TSPY* pseudogene copies within the *TSPY* array.** Each row shows *TSPY* genes for one QC-passing sample (ordered phylogenetically). Individual *TSPY* copies are drawn as arrows in assembly order. Arrowheads indicate gene orientation after re-orienting arrays to a common display direction. Gene copies predicted to be functionally intact are shown in ***gray*.** Putative pseudogene copies are highlighted and colored by the feature supporting loss of coding potential status (legend; e.g., premature stop - introducing SNPs or indels in specific exons). The detached arrows to the right represent the single *TSPY2* copy located outside the tandem array. The slash "//" indicates a genome separation > 100 kb (not to scale). Sample IDs are color-coded to indicate IR1 orientation state (***red***: array and *TSPY2* flipped relative to reference, +IR1; ***yellow***: *TSPY2* matches +IR1; ***purple***, *TSPY2* matches −IR1. Among the *TSPY1* arrays there were four variants which produce a premature stop codon (PSC): 1) a SNV (G>T) in exon 1 (green pseudogene), 2) an indel (G insertion) in exon 1 (pink pseudogene), 3) a SNV (G>A) in exon 4 (orange pseudogene), and 4) a SNV (C>T) in exon 6 (yellow pseudogene, produces a PSC though not as likely as it is in the last exon near the UTR).

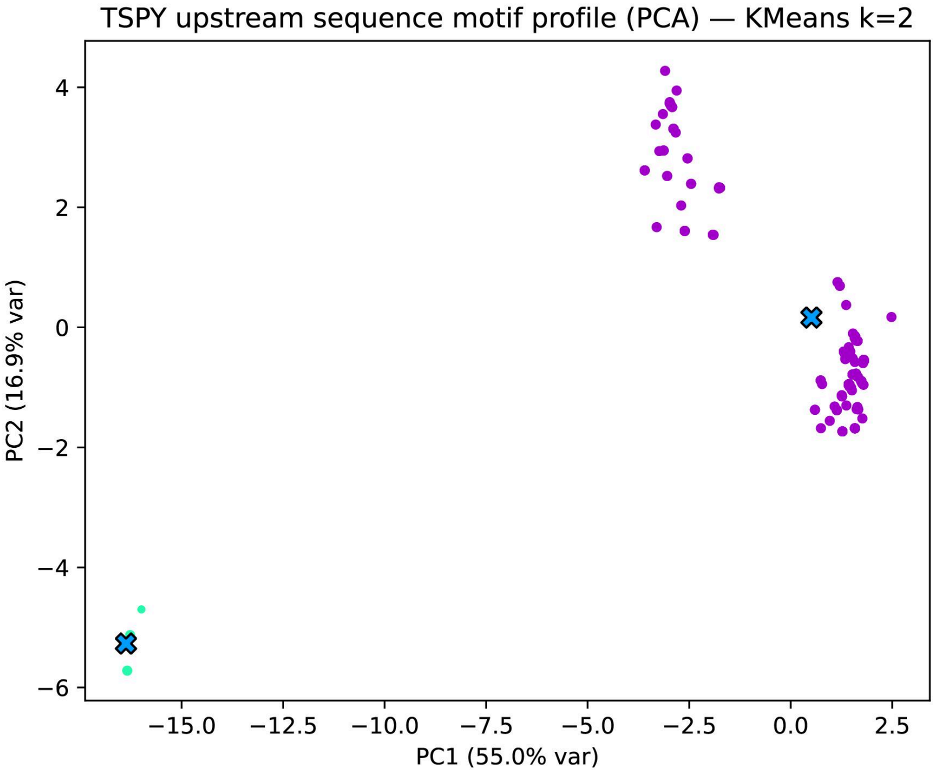


**Supplementary Figure 50. Two major classes of upstream TF-motif profiles among *TSPY/TSPY2* copies:** Principal component analysis (PCA) was performed on transcription factor (TF) motif counts computed from the 2 kb upstream regions of individual *TSPY* and *TSPY2* gene copies. Each plotted observation represents one gene copy positioned by its scores on PC1 (55% variance) and PC2 (16.9% variance). Gene copies with identical TF-motif profiles have identical PCA scores and therefore overlap exactly; consequently, a single visible point may represent multiple gene copies. K-means clustering (k=2) partition copies into two groups with distinct motif compositions (***Purple*** and ***Teal***), and cluster centroids are marked with an "X." Thus, *TSPY* upstream regions fall into two main promoter-like motif patterns rather than forming a single continuum.

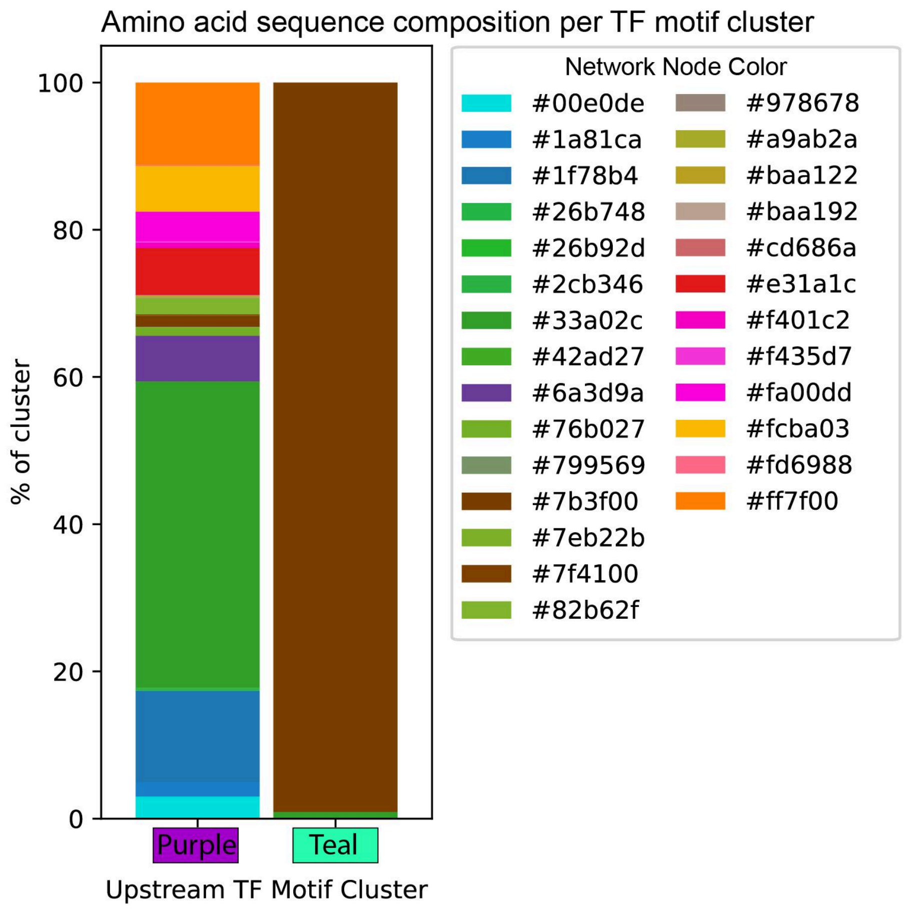


**Supplementary Figure 51. Amino acid sequence group composition of *TSPY* copies across upstream TF motif clusters.** Stacked bar plots show for each upstream TF motif profile cluster (***Purple*** and ***Teal***; defined by k-means clustering of motif counts in upstream regions), the proportion of *TSPY/TSPY2* gene copies assigned to each amino-acid Exact Sequence Group (ESG) from the *TSPY* protein sequence network. Segment colors correspond to ESG network node colors. Results show that the TF motifs are different between copies of *TSPY* within the array (***Purple***) versus the *TSPY2* copy outside the array (***Teal***). The two TF motif clusters differ in their mixture of protein sequence groups, indicating that promoter-like motif profiles are associated with specific *TSPY* amino acid variants.

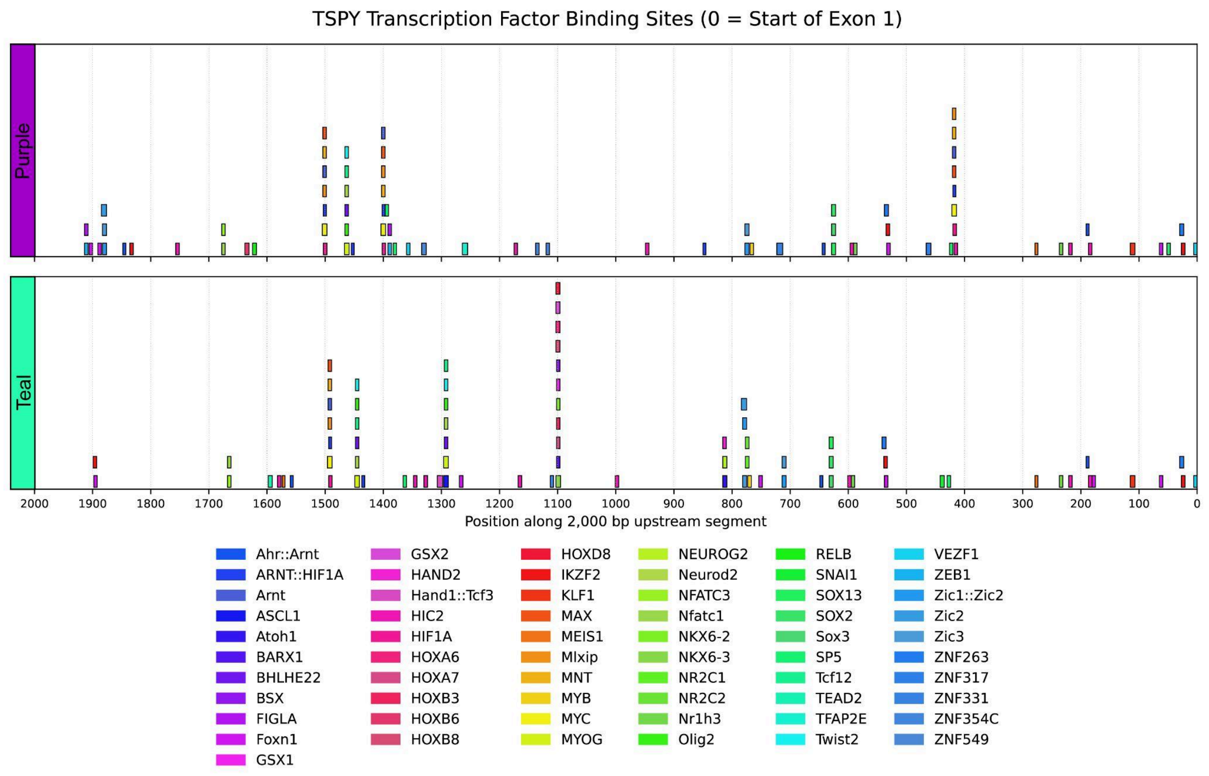


**Supplementary Figure 52. Predicted transcription factor locations in the 2 kb upstream region of *TSPY* exon 1.** Predicted transcription factor (TF) motif matches are shown across the 2 kb region of exon 1 (x-axis; distance in bp with 0 marking the start of exon 1) for one randomly selected representative *TSPY* copy from each upstream TF motif cluster (***Purple*** and ***Teal***). Colored rectangles indicate the positions of motif matches for the indicated TFs. Stacked rectangles indicate overlapping motif matches at the same locus. Unlike the *RBMY* TF motif clusters, the two *TSPY* TF motif clusters (purple and teal) varied much more in the TF motifs they contained. The first cluster (purple) was comprised of 53 TF motifs of which 20 (GSX1, Hand1::Tcf3, BARX1, ASCL1, GSX2, NR2C2, Atoh1, BSX, HOXA6, HOXB8, NKX6-2, TEAD2, HOXA7, SNAI1, HAND2, NEUROG2, HOXB3, HOXD8, NKX6-3, NR2C1) were unique compared to cluster two (teal). The teal cluster had 41 TF motifs of which 8 ('TFAP2E', 'RELB', 'ZNF317', 'FIGLA', 'ZEB1', 'ZNF549', 'ZNF331', 'HOXB6') were unique compared to the purple cluster. Cluster 1 (purple) gene array TF motifs GO term biological processes were linked more towards spermatogenesis, developmental morphogenesis, and Sertoli/immune signaling. In contrast, Cluster 2 (teal), which was *TSPY2*-specific, was enriched for GO biological process terms related to brain, central nervous system, and olfactory development.

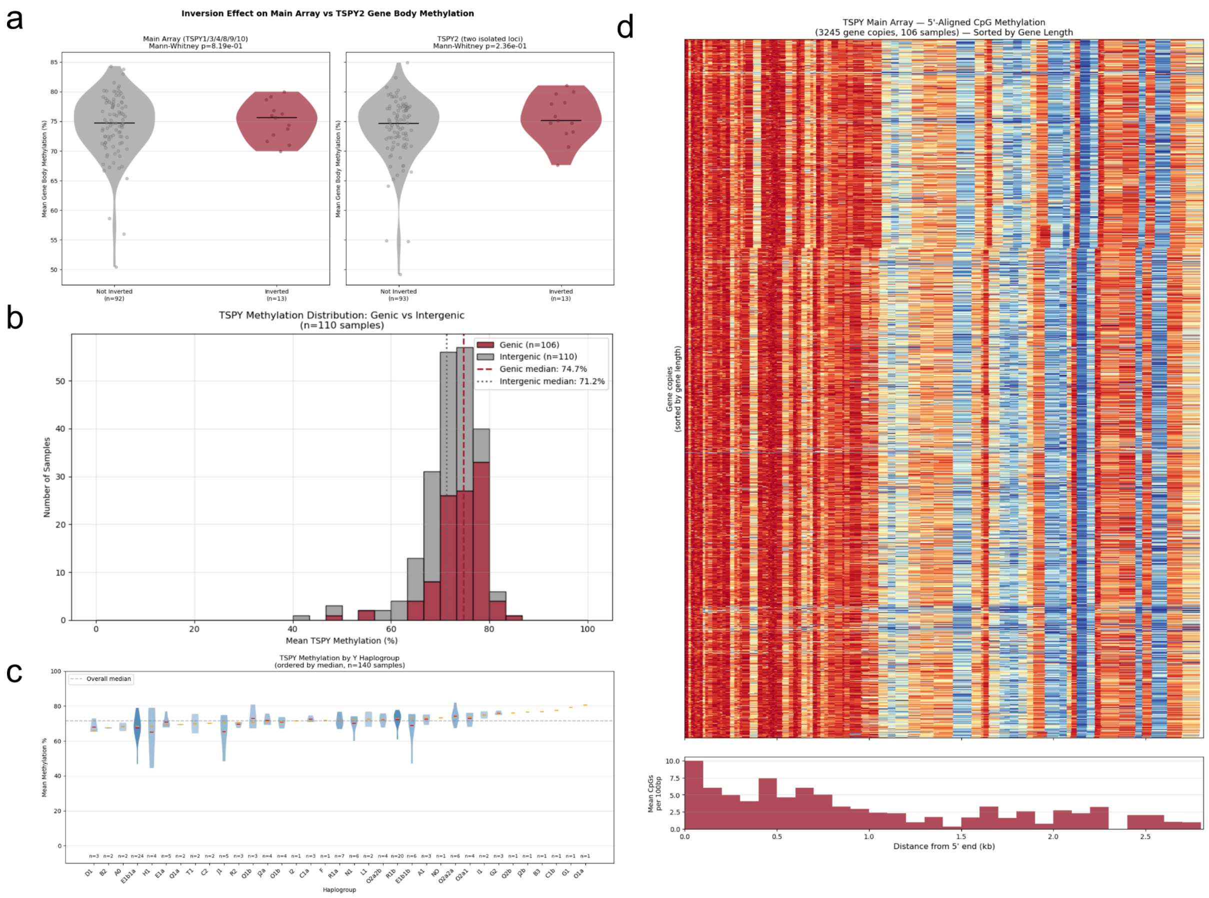


**Supplementary Figure 53. *TSPY* array CpG methylation signals. a)** *TSPY* methylation by inversion status. Tandem array *TSPY* genes and *TSPY2* show no significant methylation difference between inverted and non-inverted samples (Mann-Whitney U test, $p > 0.05$). **b)** Distribution of mean *TSPY* methylation in genic versus intergenic regions. Gene bodies show higher methylation than intergenic spacers, consistent with expected CpG methylation patterns. **c)** *TSPY* methylation by Y chromosome haplogroup. No strong haplogroup-driven differences in methylation were observed. Haplogroups ordered by median methylation. Sample sizes indicated below each group. **d)** *TSPY* main array gene methylation aligned by 5' ends shows consistent pattern across all samples, with hypermethylation at the 5' end of the gene body, consistent with transcription-associated gene body methylation.

### *RBMY* genes

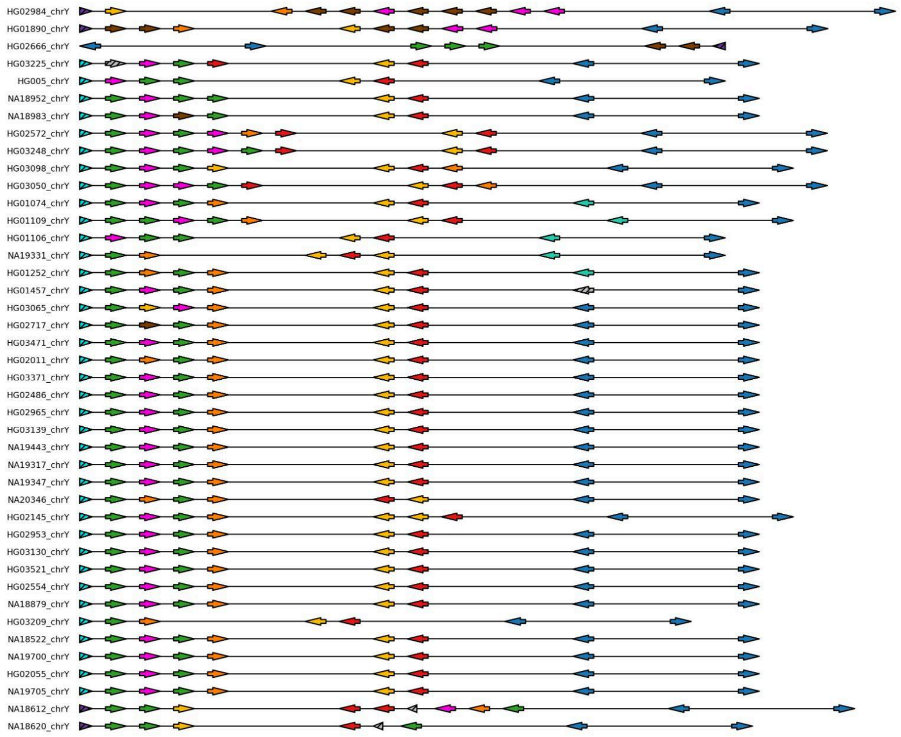

HG02984_chrY
HG01890_chrY
HG02666_chrY
HG03225_chrY
HG005_chrY
NA18952_chrY
NA18983_chrY
HG02572_chrY
HG03248_chrY
HG03098_chrY
HG03050_chrY
HG01074_chrY
HG01109_chrY
HG01106_chrY
NA19331_chrY
HG01252_chrY
HG01457_chrY
HG03065_chrY
HG02717_chrY
HG03471_chrY
HG02011_chrY
HG03371_chrY
HG02486_chrY
HG02965_chrY
HG03139_chrY
NA19443_chrY
NA19317_chrY
NA19347_chrY
NA20346_chrY
HG02145_chrY
HG02953_chrY
HG03130_chrY
HG03521_chrY
HG02554_chrY
NA18879_chrY
HG03209_chrY
NA18522_chrY
NA19700_chrY
HG02055_chrY
NA19705_chrY
NA18612_chrY
NA18620_chrY


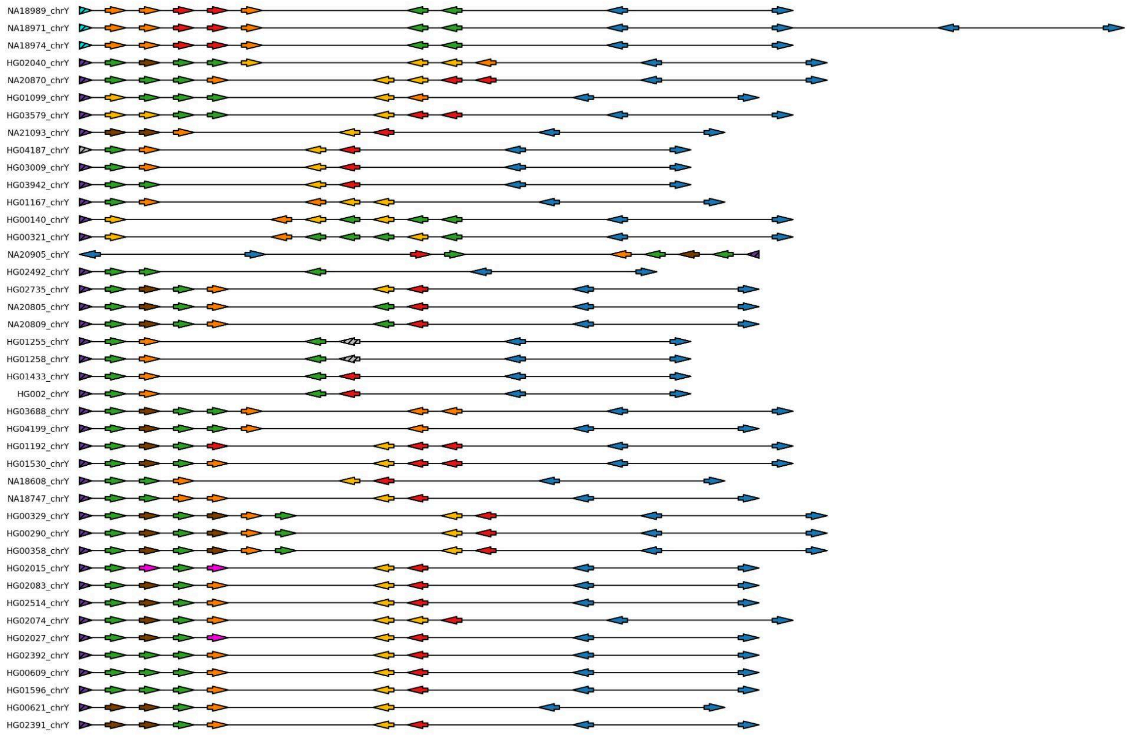

NA18989_chrY
NA18971_chrY
NA18974_chrY
HG02040_chrY
NA20870_chrY
HG01099_chrY
HG03579_chrY
NA21093_chrY
HG04187_chrY
HG03009_chrY
HG03942_chrY
HG01167_chrY
HG00140_chrY
HG00321_chrY
NA20905_chrY
HG02492_chrY
HG02735_chrY
NA20805_chrY
NA20809_chrY
HG01255_chrY
HG01258_chrY
HG01433_chrY
HG002_chrY
HG03688_chrY
HG04199_chrY
HG01192_chrY
HG01530_chrY
NA18608_chrY
NA18747_chrY
HG00329_chrY
HG00290_chrY
HG00358_chrY
HG02015_chrY
HG02083_chrY
HG02514_chrY
HG02074_chrY
HG02027_chrY
HG02392_chrY
HG00609_chrY
HG01596_chrY
HG00621_chrY
HG02391_chrY

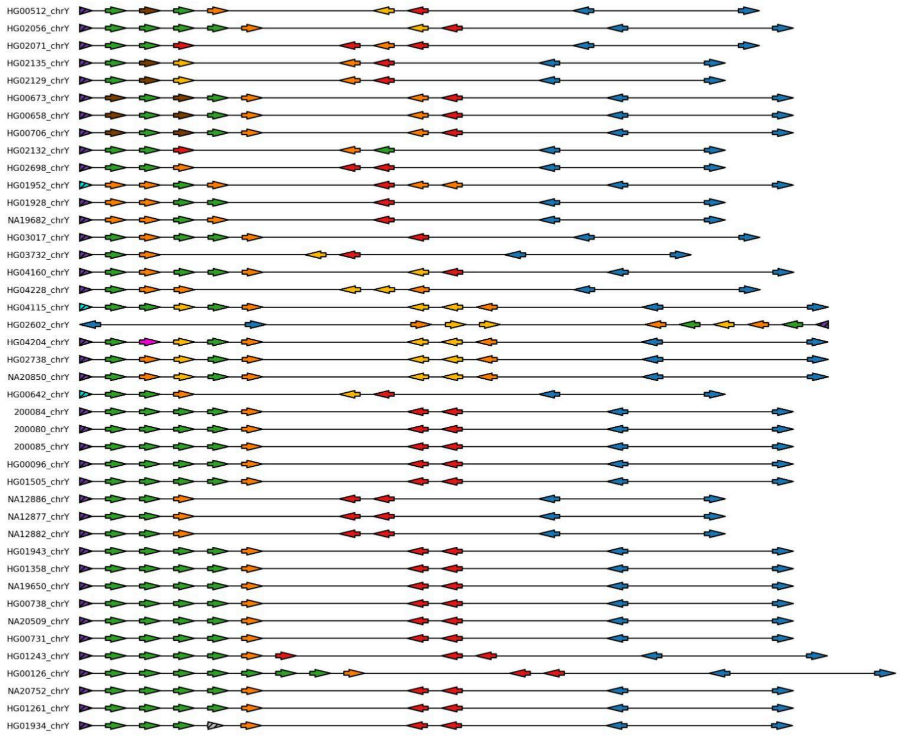


**Supplementary Figure 54. Organization and coding sequence diversity of *RBMY* genes.** For each QC-passing sample (n=126; rows), *RBMY* gene copies are displayed in assembly order. Each arrow represents one *RBMY* copy, with arrow direction indicating gene orientation. Gene copies are colored by amino-acid Exact Sequence Group (ESG) assignment from the *RBMY* protein sequence network analysis. Copies containing premature stop codons (putative pseudogenes) are indicated with black striped overlays. The break between *RBMY* copy groups denotes a genomic separation >100,000 base pairs (not to scale) and distinguishes the primary *RBMY* array from *RBMY* copies located outside the array.

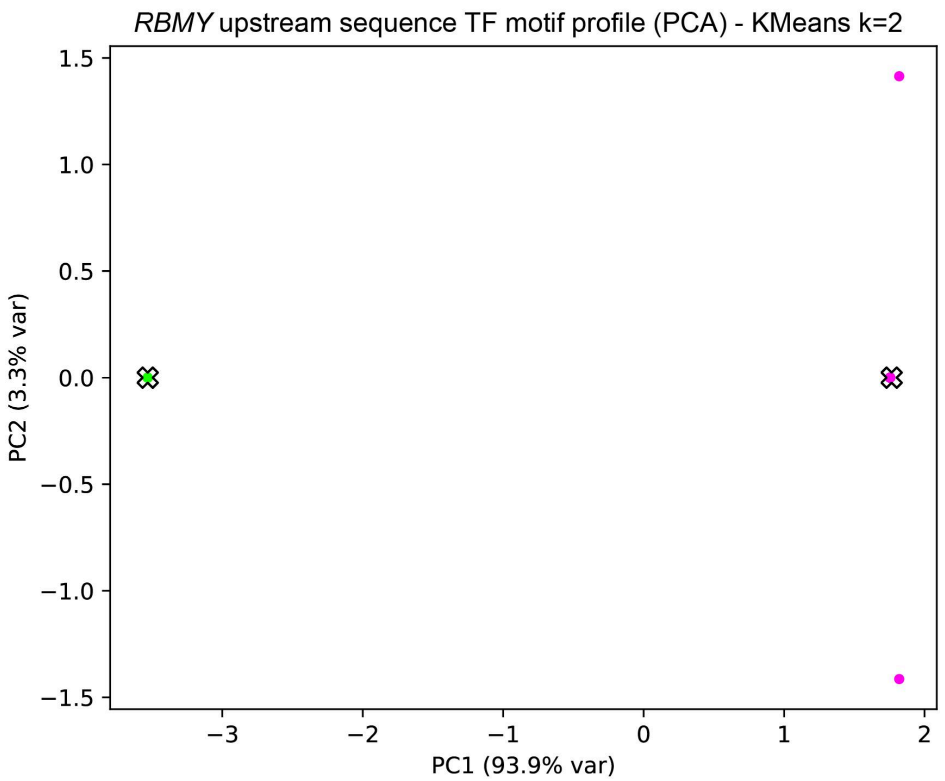


**Supplementary Figure 55. Two upstream TF motif profile groups among *RBMY* gene copies.** Principal component analysis (PCA) was performed on transcription factor (TF) motif counts computed from the 1 kb upstream regions of individual *RBMY* gene copies. Each plotted observation represents one *RBMY* gene copy positioned by its scores on PC1 (93.9% variance) and PC2 (3.3% variance). Multiple *RBMY* copies share identical TF-motif profiles and therefore occupy identical PCA coordinates, causing them to overlap exactly; consequently, a single visible point may represent multiple gene copies. K-means clustering (k=2) partitions copy into two motif profile groups (***Lime*** and ***Pink***), and cluster centroids are marked with an "X".

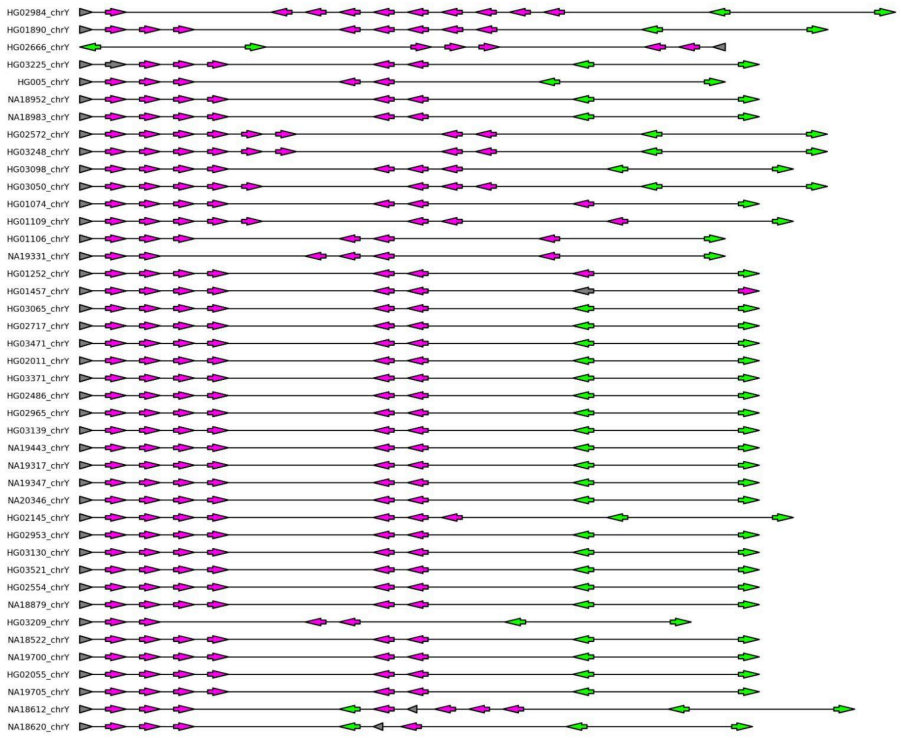
HG02984_chrY
HG01890_chrY
HG02666_chrY
HG03225_chrY
HG005_chrY
NA18952_chrY
NA18983_chrY
HG02572_chrY
HG03248_chrY
HG03098_chrY
HG03050_chrY
HG01074_chrY
HG01109_chrY
HG01106_chrY
NA19331_chrY
HG01252_chrY
HG01457_chrY
HG03065_chrY
HG02717_chrY
HG03471_chrY
HG02011_chrY
HG03371_chrY
HG02486_chrY
HG02965_chrY
HG03139_chrY
NA19443_chrY
NA19317_chrY
NA19347_chrY
NA20346_chrY
HG02145_chrY
HG02953_chrY
HG03130_chrY
HG03521_chrY
HG02554_chrY
NA18879_chrY
HG03209_chrY
NA18522_chrY
NA19700_chrY
HG02055_chrY
NA19705_chrY
NA18612_chrY
NA18620_chrY

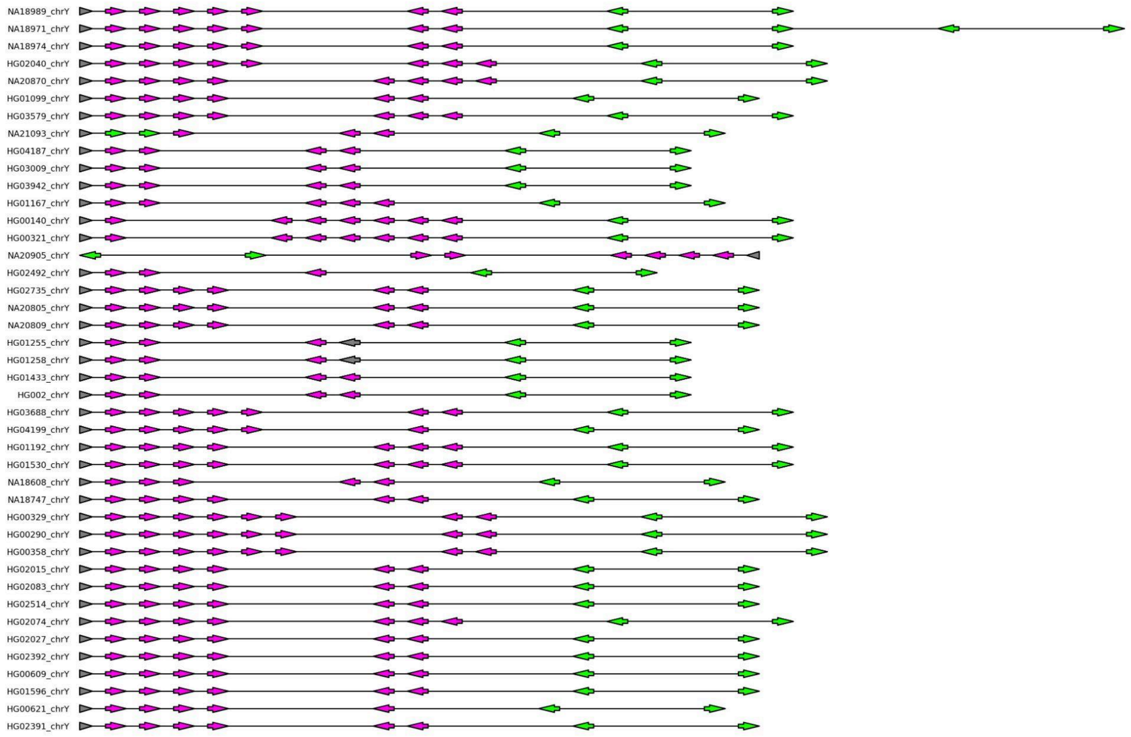
NA18989_chrY
NA18971_chrY
NA18974_chrY
HG02040_chrY
NA20870_chrY
HG01099_chrY
HG03579_chrY
NA21093_chrY
HG04187_chrY
HG03009_chrY
HG03942_chrY
HG01167_chrY
HG00140_chrY
HG00321_chrY
NA20905_chrY
HG02492_chrY
HG02735_chrY
NA20805_chrY
NA20809_chrY
HG01255_chrY
HG01258_chrY
HG01433_chrY
HG002_chrY
HG03688_chrY
HG04199_chrY
HG01192_chrY
HG01530_chrY
NA18608_chrY
NA18747_chrY
HG00329_chrY
HG00290_chrY
HG00358_chrY
HG02015_chrY
HG02083_chrY
HG02514_chrY
HG02074_chrY
HG02027_chrY
HG02392_chrY
HG00609_chrY
HG01596_chrY
HG00621_chrY
HG02391_chrY

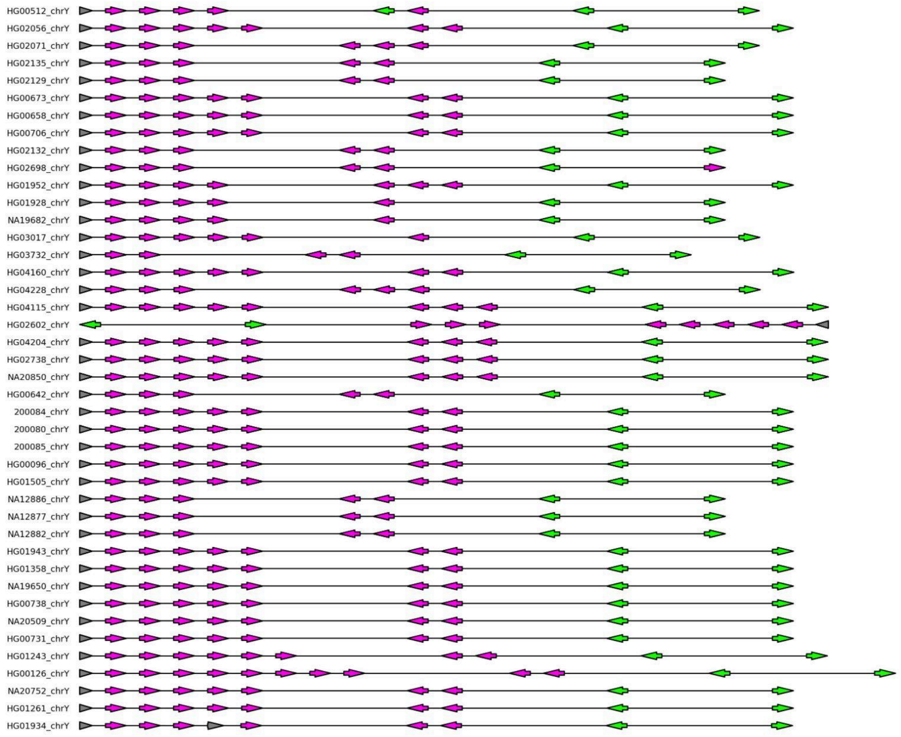


**Supplementary Figure 56. *RBMY* arrays annotated by upstream TF-motif cluster assignment.** *RBMY* gene copies are shown in assembly order for each QC-passing sample (rows; ordered phylogenetically). Each arrow represents one *RBMY* copy and arrow direction indicates gene orientation. Arrow color indicates the upstream TF-motif cluster assigned to that copy (cluster 1 - ***pink***, cluster 2 - ***lime***; defined by PCA/k-means of TF motif counts in the upstream region).Thus, *RBMY* copies within the array fall into two distinct upstream TF motif clusters, and most samples show a consistent mix of these promoter types across their duplicated gene copies.

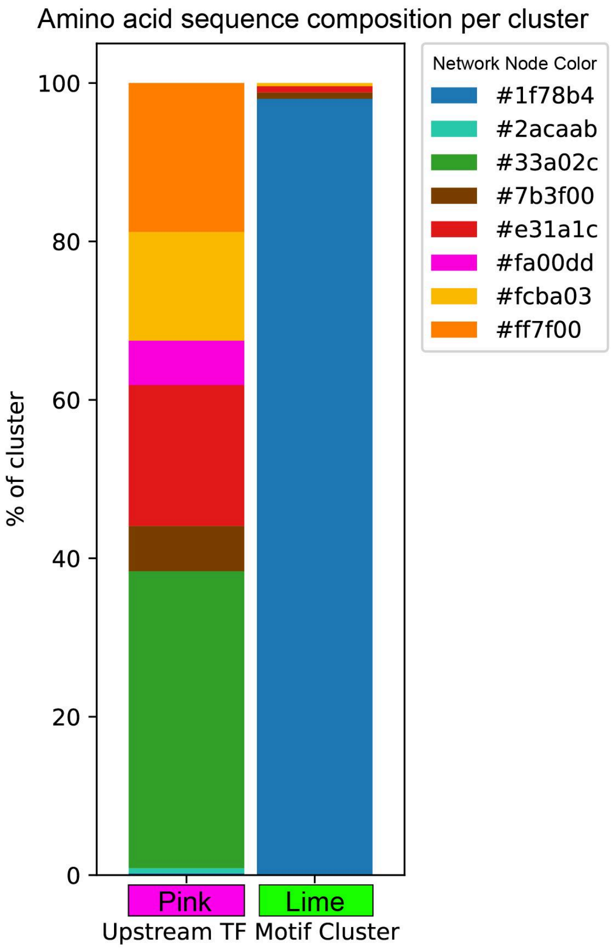


**Supplementary Figure 57. Protein sequence group composition of *RBMY* copies across upstream TF motif clusters.** Stacked bar plots show for each upstream TF motif profile cluster (***Pink*** and ***Lime***; defined by k-means clustering of TF motif counts in *RBMY* upstream regions), the proportion of *RBMY* gene copies assigned to each amino-acid Exact Sequence Group (ESG) from the *RBMY* protein sequence network. Bar segments are colored by their network ESG node color (HEX codes). The two upstream TF motif clusters differ in their mixture of *RBMY* protein sequence groups, suggesting an association between promoter-like motif profiles and specific *RBMY* variants.

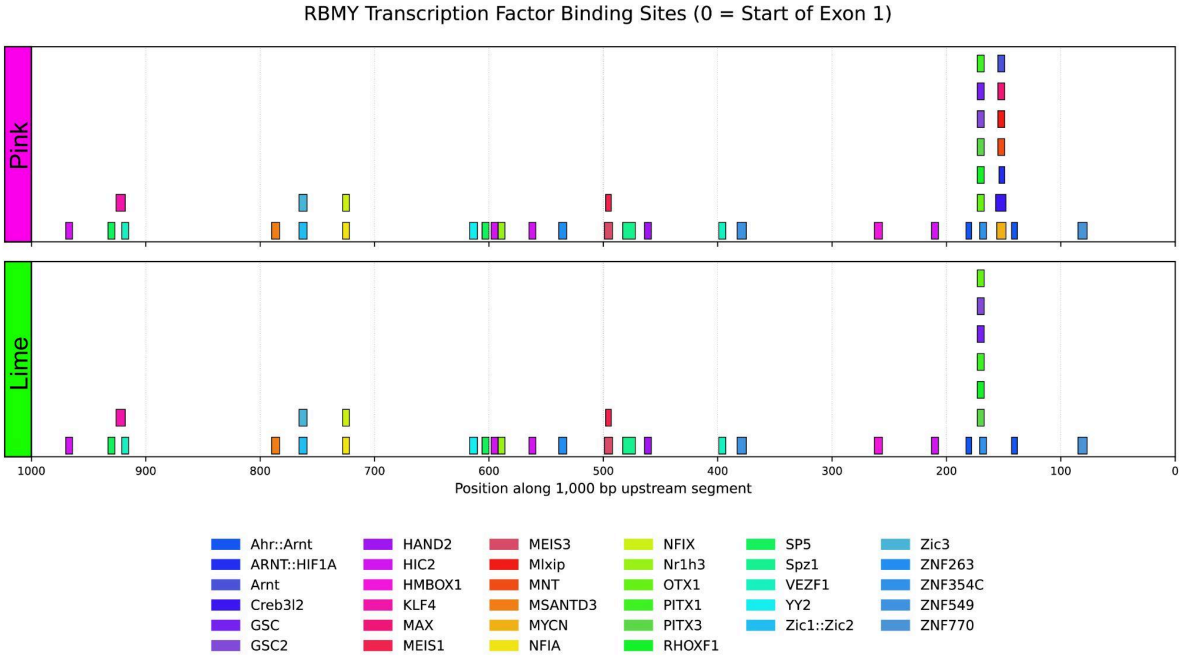


**Supplementary Figure 58. Predicted TF motif positions in the 1 kb upstream region of *RBMY* exon 1.** Predicted transcription factor (TF) motif matches are shown across the 1 kb region upstream of exon 1 (x-axis; distance in bp with 0 marking the start of exon 1) for one randomly selected representative *RBMY* copy from each upstream TF motif cluster (***Pink*** and ***Lime***). Colored rectangles indicate motif match positions. Stacked rectangles denote overlapping motif matches. Motif architectures are broadly similar between the two examples, with several motif matches present in the ***pink*** example, but not in the ***lime*** example (including Arnt, ARNT::HIF1A, MAX, MYCN, MNT, Mlxip, and Creb3l2).

### Centromere

Contributing authors: Shenghan Gao, Carolina Montano, Keith Oshima, Mark Loftus, Pille Hallast, Glennis Logsdon

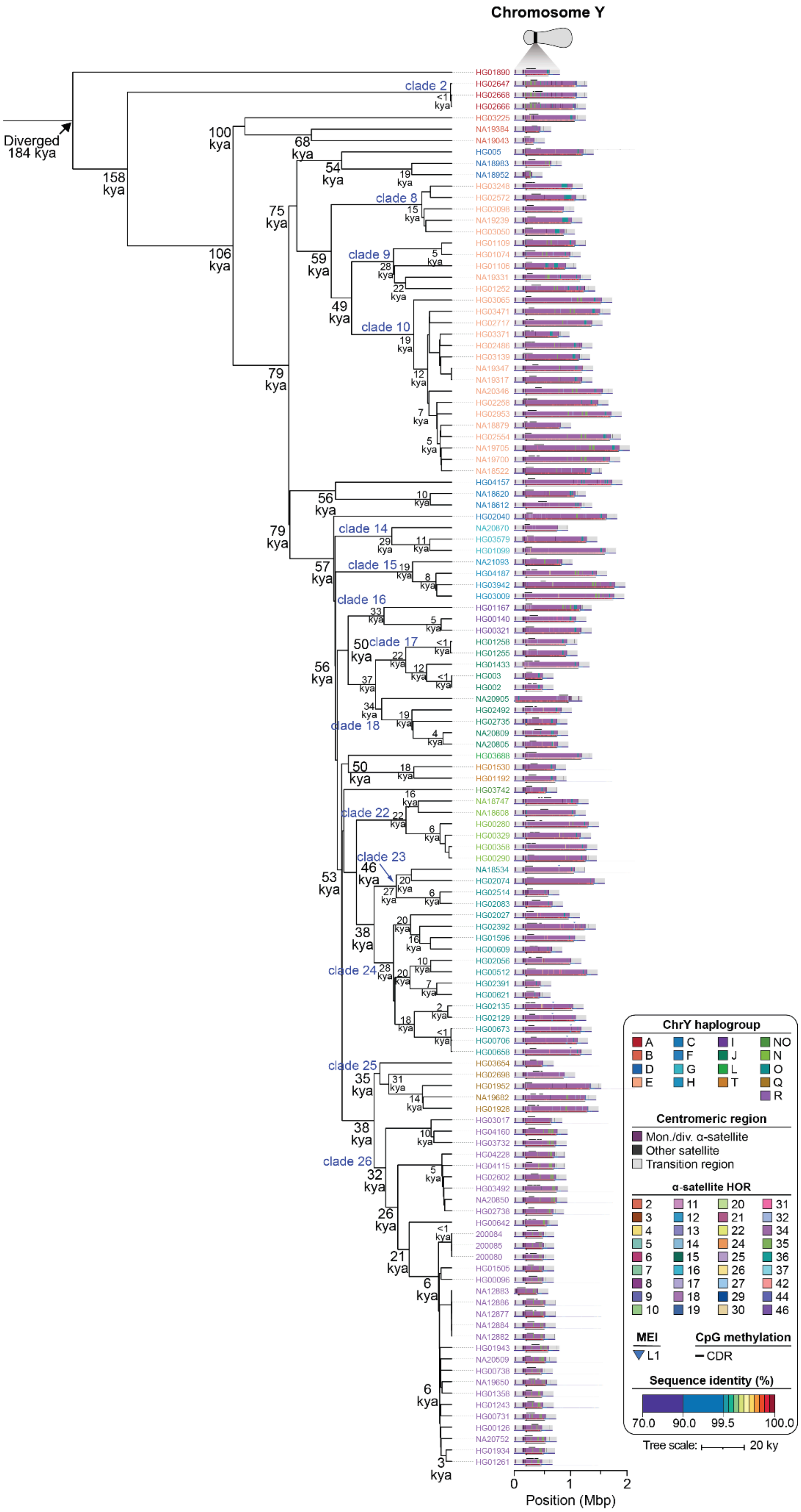
Chromosome Y
Diverged 184 kya
158 kya
106 kya
100 kya
79 kya
75 kya
68 kya
59 kya
54 kya
49 kya
clade 2
clade 8
clade 9
clade 10
clade 14
clade 15
clade 16
clade 17
clade 18
clade 22
clade 23
clade 24
clade 25
clade 26
ChrY haplogroup
A B D E C F G H I J L T NO N O Q R
Centromeric region
Mon./div. α-satellite
Other satellite
Transition region
α-satellite HOR
MEI
L1
CpG methylation
CDR
Sequence identity (%)
70.0 90.0 99.5 100.0
Tree scale: 20 ky
0 1 2
Position (Mbp)

**Supplementary Figure 59. Phylogenetic reconstruction of chromosome Y centromere haplotypes.** Phylogenetic tree (**Methods**) depicting the topology of the chromosome Y *DYZ3* α-satellite HOR array topology along with estimated divergence times. The clades labelled in blue were used for centromeric mutation rate estimates (**Suppl. Figure 62, Suppl. Table 25**).

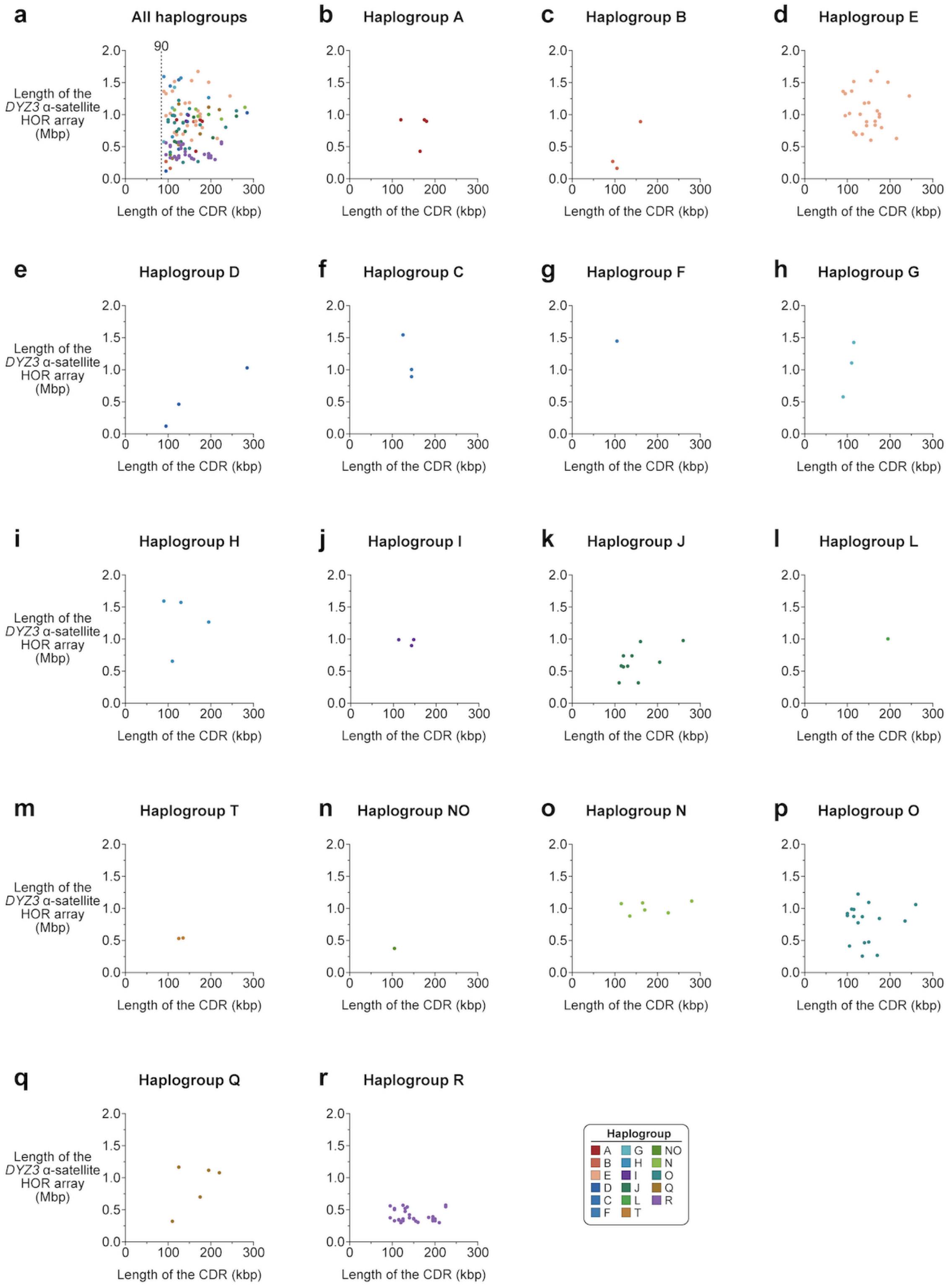


**Supplementary Figure 60. Chromosome Y putative kinetochore sites are always at least 90 kb and do not correlate with the length of the *DYZ3* α-satellite HOR array. a-r)** Plots showing the length of the putative kinetochore site, defined by the presence of a centromere dip region (or CDR[34]), and the length of the *DYZ3* α-satellite HOR array on chromosome Y for **a)** all haplogroups and **b-r)** select haplogroups.

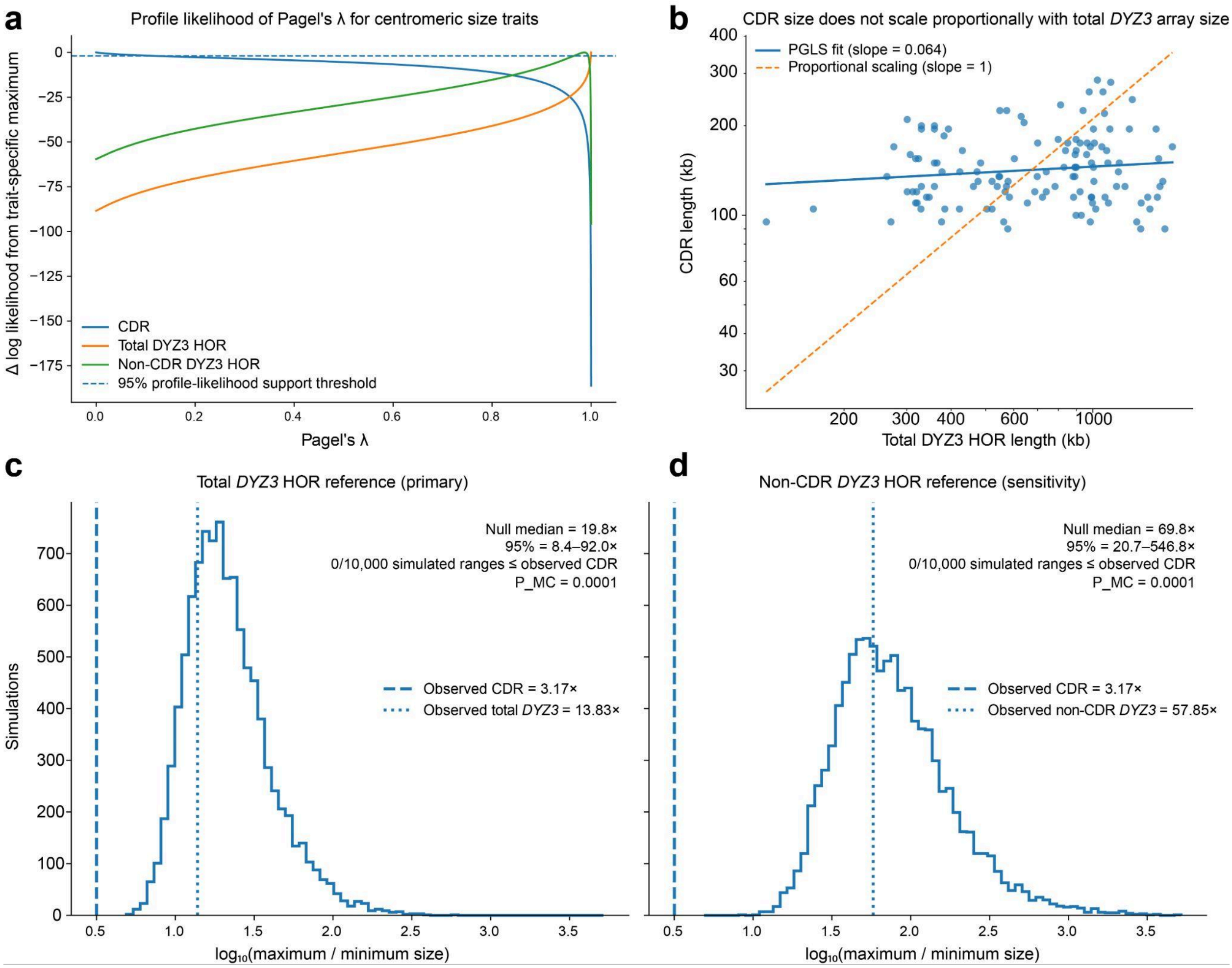


**Supplementary Figure 61. CDR size is decoupled from phylogenetic and structural variation in the underlying *DYZ3* α-satellite HOR array. (a)** Profile-likelihood analysis of Pagel's λ for CDR length, total *DYZ3* α-satellite HOR array length, and non-CDR *DYZ3* HOR array length across 123 Y centromeres. Total *DYZ3* HOR array length showed strong phylogenetic signal (λ = 1.00), whereas CDR length showed essentially no detectable phylogenetic signal (λ = 0). Curves show the change in log likelihood relative to the trait-specific maximum, and the dashed horizontal line indicates the 95% profile-likelihood support threshold. **(b)** Relationship between total *DYZ3* HOR array length and CDR length. Points represent individual Y centromeres. The solid line shows the fitted phylogenetic generalized least-squares (PGLS) relationship on the log–log scale (slope = 0.064, 95% CI = −0.024 to 0.153; $P$ = 0.153 for slope = 0), whereas the dashed line shows proportional scaling (slope = 1), which was strongly rejected ($P$ = $4.96e^{42}$). **(c)** Distribution of fold ranges from 10,000 simulations under the phylogenetic process fitted to total *DYZ3* HOR length. The observed CDR range (3.17-fold; dashed line) was smaller than all simulated ranges (median = 19.8-fold; 95% simulated interval = 8.4–92.0-fold; Monte Carlo $P$ = $1e^{-4}$). The dotted line indicates the observed 13.83-fold range in total *DYZ3* HOR length. **(d)** Sensitivity analysis using the phylogenetic process fitted to non-CDR *DYZ3* HOR array length. The observed CDR range was again smaller than all 10,000 simulated ranges (median = 69.8-fold; 95% simulated interval = 20.7–546.8-fold; Monte Carlo $P$ = $1e^{-4}$); the dotted line indicates the observed 57.85-fold range in non-CDR *DYZ3* HOR array length.

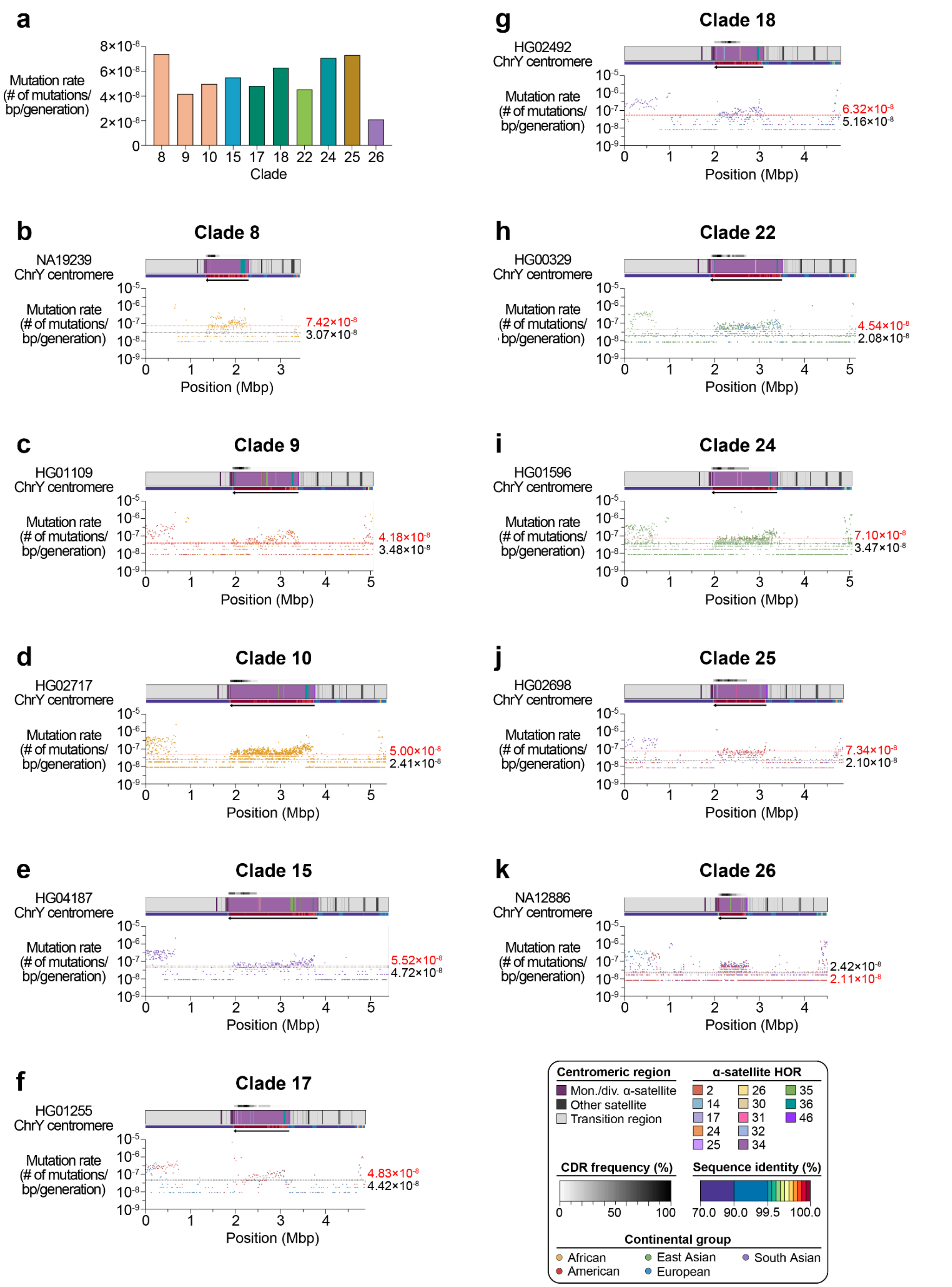


**Supplementary Figure 62. Estimated mutation rates of major centromere haplogroups from chromosome Y. a)** Estimated mean mutation rate across the *DYZ3* active α-satellite HOR array for major phylogenetic clades. **b-k)** Plots showing the estimated mutation rate across the chromosome Y centromeric regions, separated by clade. Individual data points from 10-kbp pairwise sequence alignments are shown. Mean mutation rates across the active α-satellite HOR array and flanking non-satellite sequences are shown as dotted red and black lines, respectively. See **Suppl. Fig. 59** for clade locations on the Y phylogeny.

### Yq12 and *DYZ19*

Contributing authors:

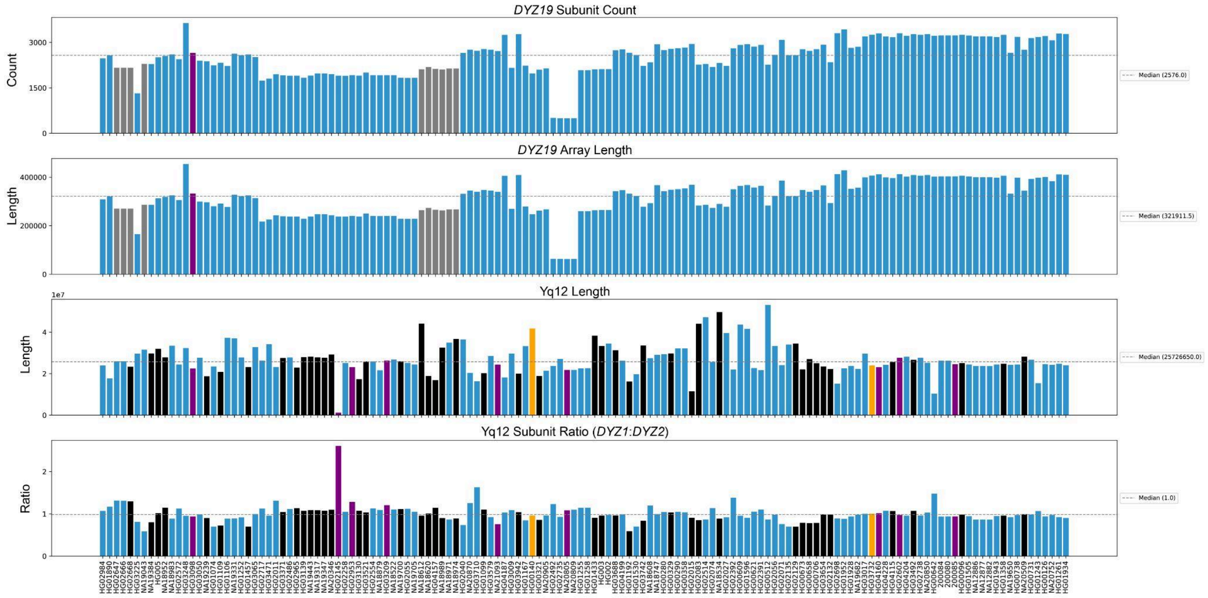


**Supplementary Figure 63. *DYZ19* and Yq12 repeat-array metrics across Y chromosome assemblies.** Four panels show repeat-array metrics for two Y-chromosome regions (*DYZ19* and Yq12) across samples. Panel 1 shows the total number of *DYZ19* subunit repeats per array, and panel 2 shows the corresponding *DYZ19* array length (bp). Panel 3 shows the assembled Yq12 region length (bp), and panel 4 shows the Yq12 subunit ratio (*DYZ1* subunit counts / *DYZ2* subunit counts). For each panel, the median is indicated by a horizontal line and annotated to the right of the plot. Bar colors denote assembly context: ***purple***, repeat subunits detected only on non-primary (non-main contig) sequence; ***gray***, deviation from an expected perfect tandem structure; ***black***, unresolved gap of Ns within the region; ***orange***, ERRSTRUCT QC flag within the region; ***blue***, no QC flags (good). Both *DYZ19* and Yq12 vary in size across individuals, and some assemblies show QC features (gaps or structural flags) that can affect repeat array estimates.

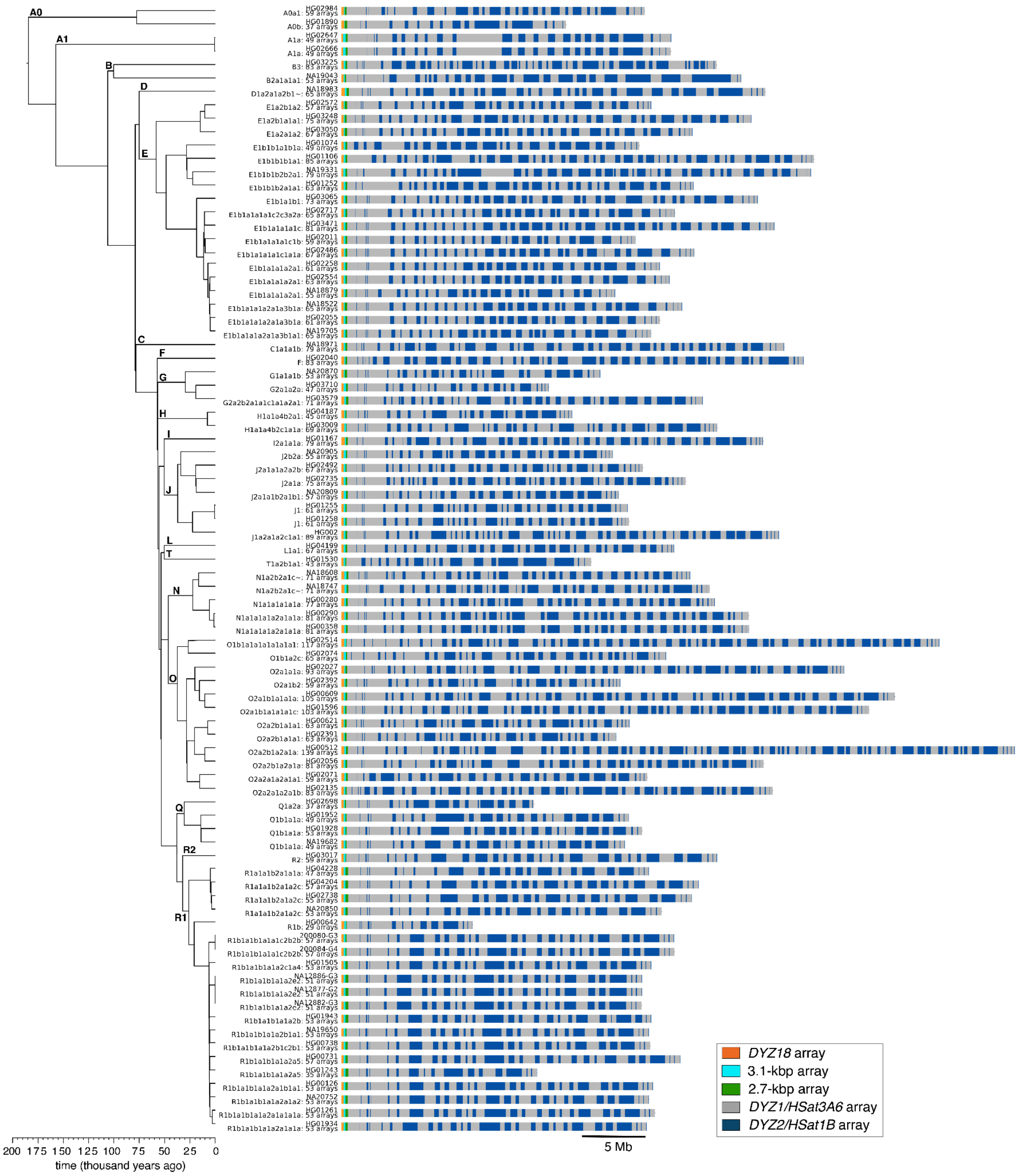


**Supplementary Figure 64. Yq12 size and repeat composition across a dated Y chromosome phylogeny.** A time-calibrated Y phylogeny (left; time in thousands of years ago) is shown alongside the repeat array composition of the Yq12 region for 84 QC-passing assemblies. The linear arrangement of annotated repeat subarrays across the Yq12 region, with colored blocks denoting distinct repeat array classes (legend; *DYZ18* array, 3.1 kbp array, 2.7 kbp array, *DYZ1/HSat3A6a* array, and *DYZ2/HSat1B* array). Only QC-passing Yq12 regions are shown (no unresolved N gaps, no ERRSTRUCT QC flags in the region, and no Yq12 repeat motifs detected on non-primary contigs). Across the dated Y phylogeny, the Yq12 region varies widely in both total size and the order/amount of its repeat subarrays, showing substantial lineage-specific structural diversity rather than a single conserved Yq12 structure.

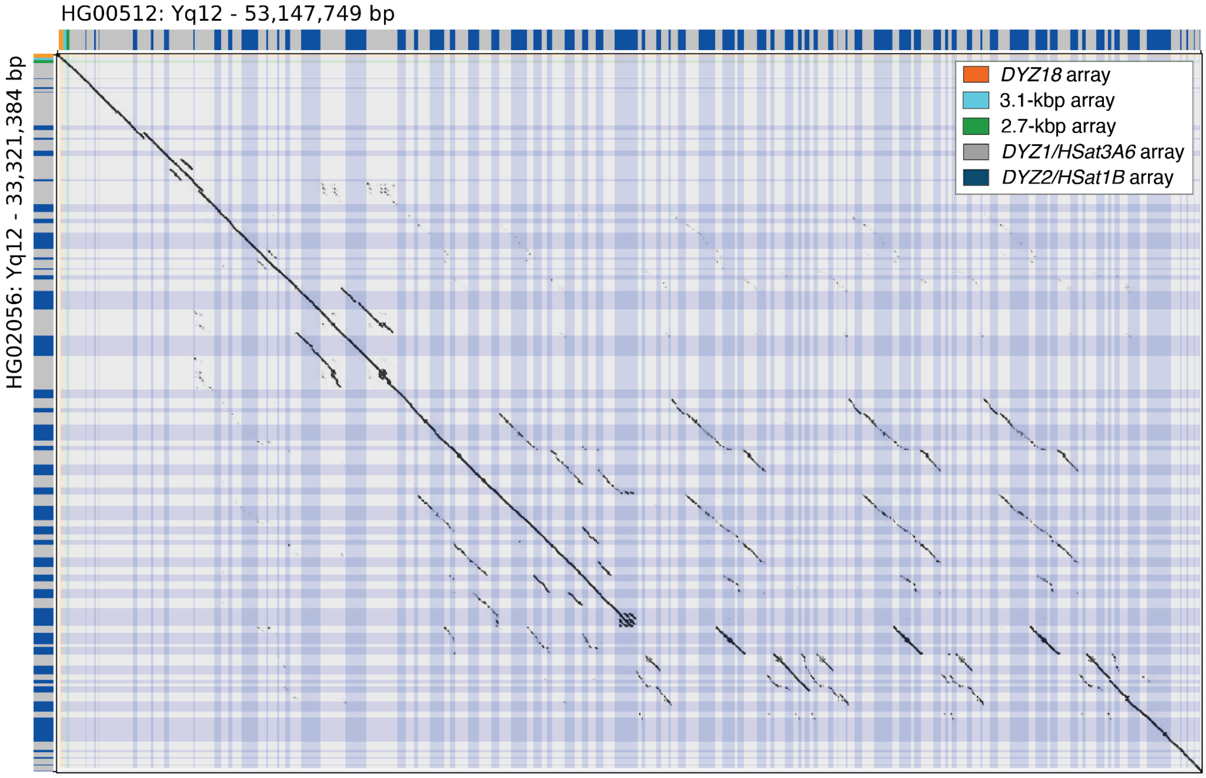


**Supplementary Figure 65. The largest assembled Yq12 region shows extensive rearrangement relative to its closest phylogenetic neighbor.** Dotplot comparison of the Yq12 assemblies from HG00512 (53,147,749 bp; ***top axis***) and its closest relative in the Y phylogeny, HG02056 (33,321,384; ***left axis***), using a 2-kb word size. Diagonal matches indicate collinear sequence, while prominent off-diagonal alignments highlight rearrangements and repeat block reorganization associated with the expanded >53 Mb Yq12 in HG00512. Colored tracks along the axes denote annotated Yq12 repeat subarrays (legend). The time to the most recent common ancestor (TMRCA) of HG00512 and HG02056 is ~10,300 years ago (95% HPD: 8,400-12,300 years). Therefore, Yq12 can expand dramatically over a short timescale, with the >53 Mb HG00512 Yq12 arising through multiple repeat block rearrangements relative to its closest phylogenetic neighbor.

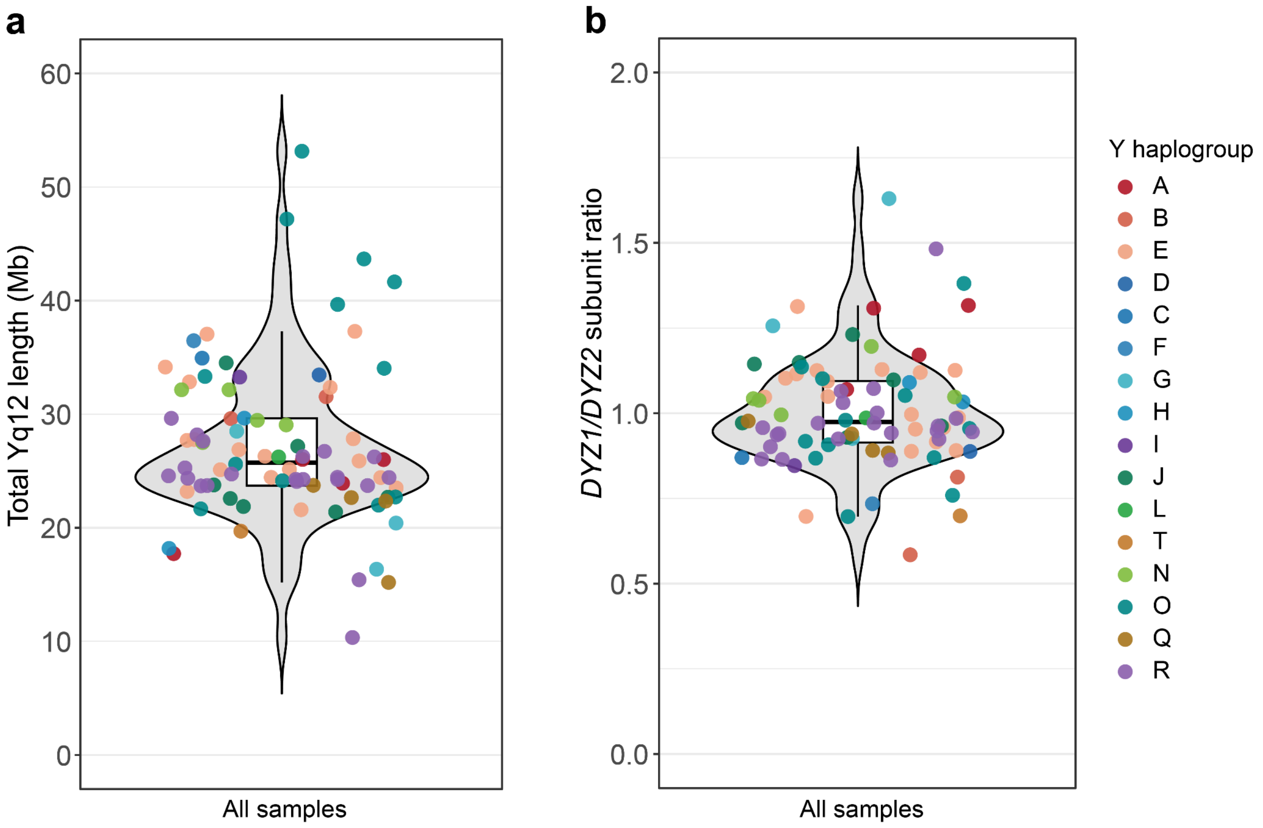


**Supplementary Figure 66. Yq12 length and *DYZ1*:*DYZ2* subunit ratio across fully assembled samples.** Violin plots summarize variation in (***left***) total assembled Yq12 length (Mb) and (***right***) the Yq12 subunit composition measured as the *DYZ1*/*DYZ2* repeat count ratio across 84 samples with QC-passing Yq12 assemblies (no unresolved N gaps within Yq12 and no Yq12 repeat motifs detected on no-primary contigs). Points represent individual samples and are colored by Y chromosome haplogroup (hg). Thus, Yq12 length varies widely across samples, while the *DYZ1*:*DYZ2* ratio clusters near 1 with modest haplogroup-specific shifts.

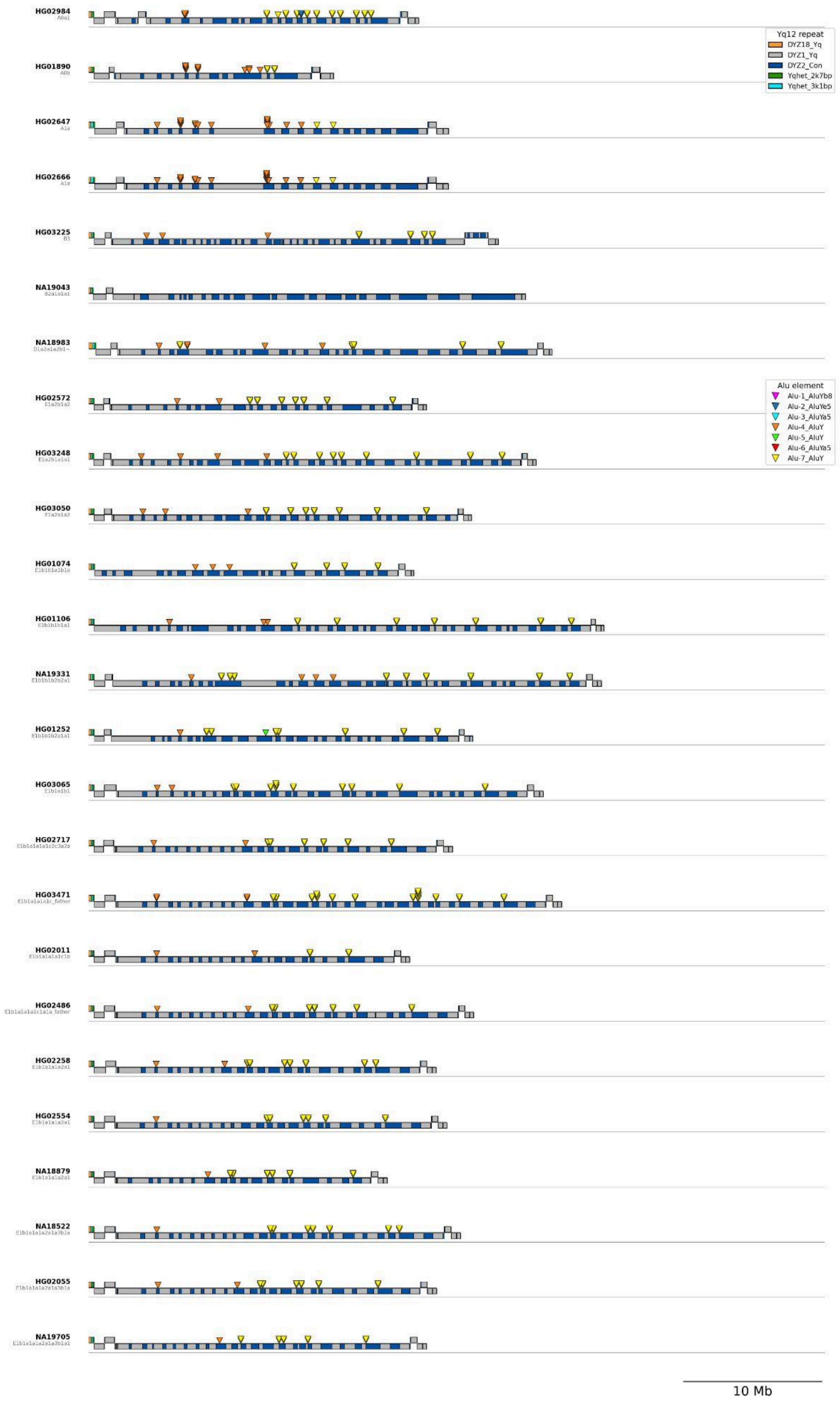
HG02984
HG01890
HG02647
HG02666
HG03225
NA19043
NA18983
HG02572
HG03248
HG03050
HG01074
HG01106
NA19331
HG01252
HG03065
HG02717
HG03471
HG02011
HG02486
HG02258
HG02554
NA18879
NA18522
HG02055
NA19705
Yq12 repeat
DYZ18_Yq
DYZ1_Yq
DYZ2_Con
Yqhet_2k7bp
Yqhet_3k1bp
Alu element
Alu-1_AluYb8
Alu-2_AluYe5
Alu-3_AluYa5
Alu-4_AluY
Alu-5_AluY
Alu-6_AluYa5
Alu-7_AluY
10 Mb

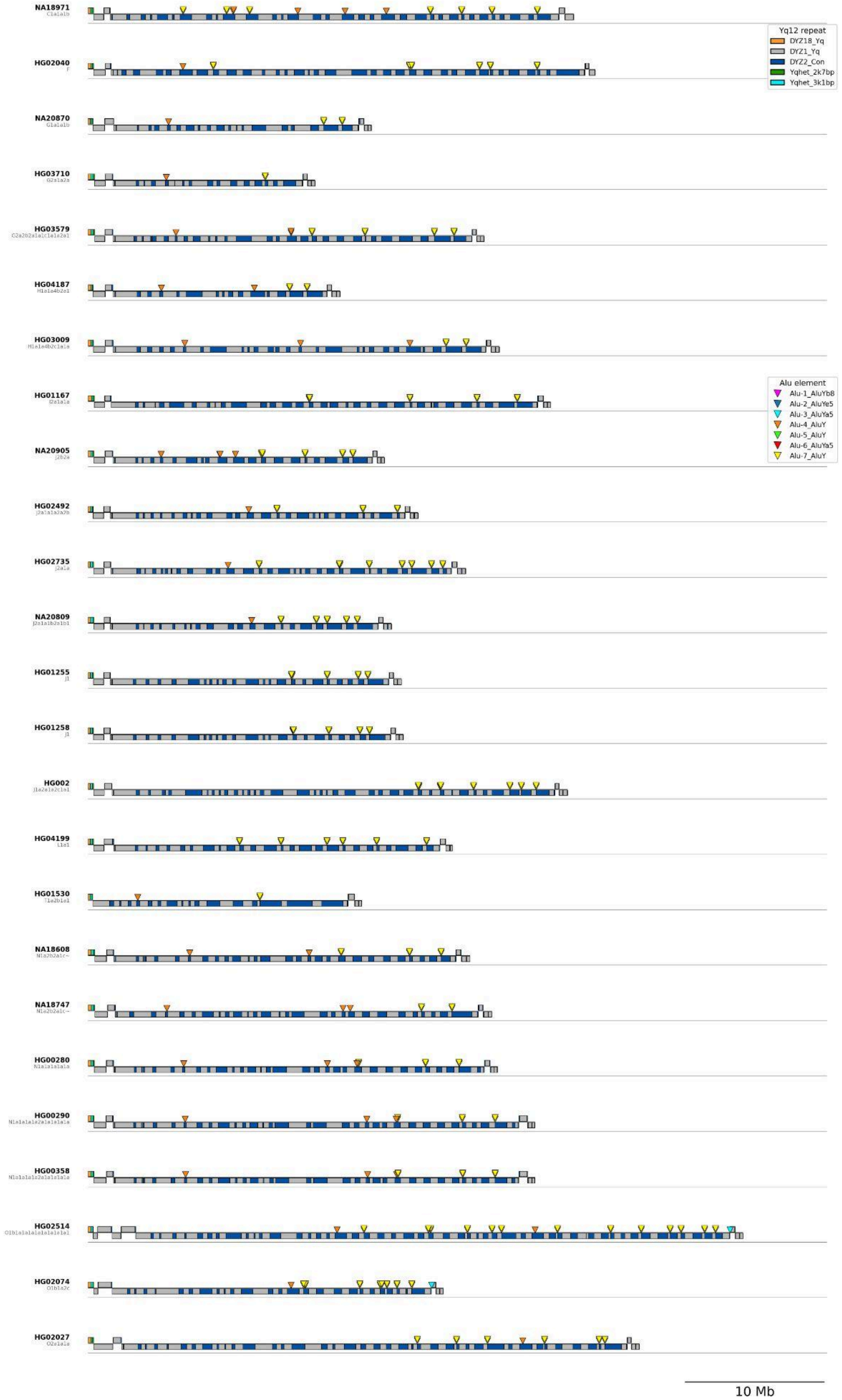
NA18971
HG02040
NA20870
HG03710
HG03579
HG04187
HG03009
HG01167
NA20905
HG02492
HG02735
NA20809
HG01255
HG01258
HG002
HG04199
HG01530
NA18608
NA18747
HG00280
HG00290
HG00358
HG02514
HG02074
HG02027
Yq12 repeat
DYZ18_Yq
DYZ1_Yq
DYZ2_Con
Yqhet_2k7bp
Yqhet_3k1bp
Alu element
Alu-1_AluYb8
Alu-2_AluYe5
Alu-3_AluYa5
Alu-4_AluY
Alu-5_AluY
Alu-6_AluYa5
Alu-7_AluY
10 Mb

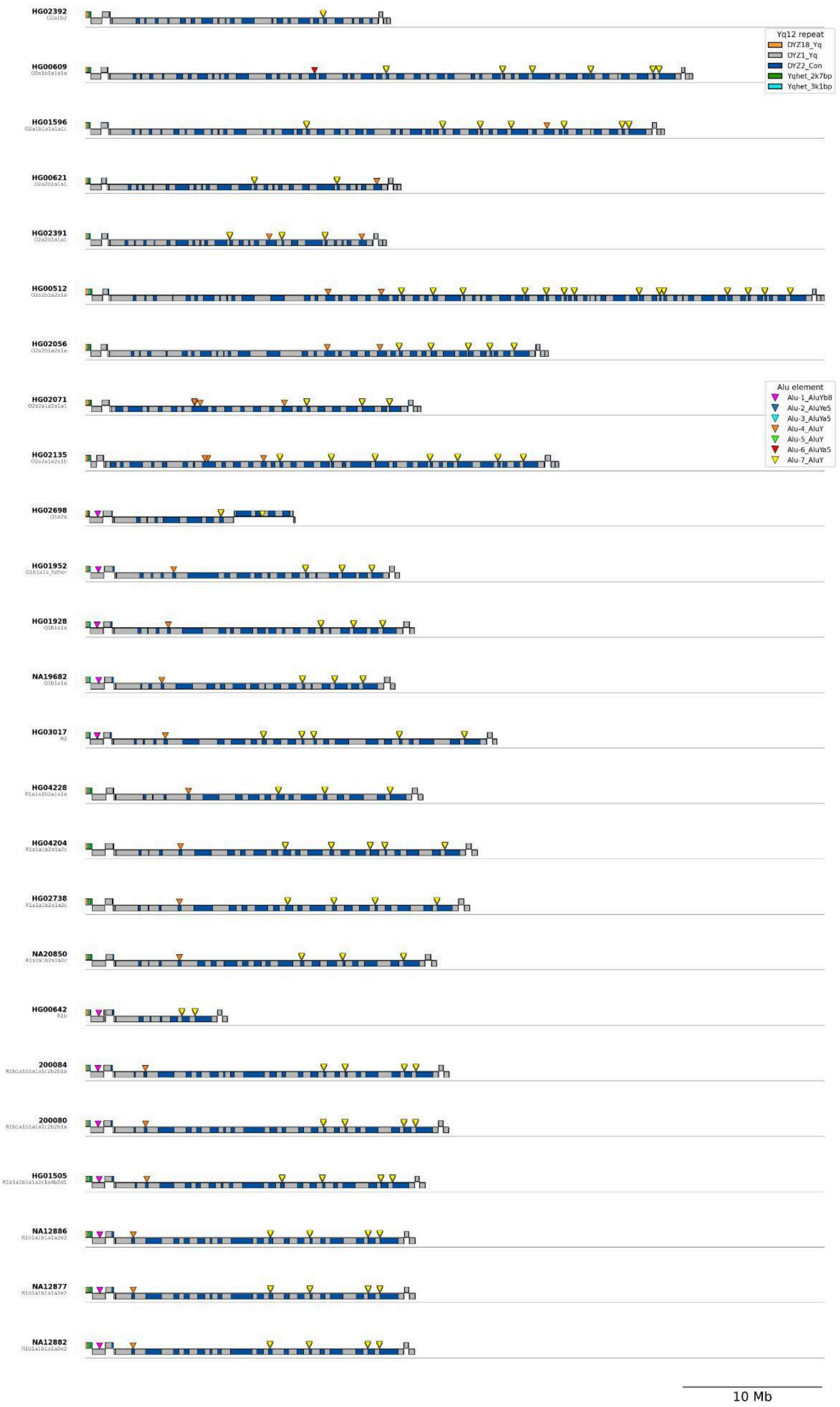
HG02392
HG00609
HG01596
HG00621
HG02391
HG00512
HG02056
HG02071
HG02135
HG02698
HG01952
HG01928
NA19682
HG03017
HG04228
HG04204
HG02738
NA20850
HG00642
200084
200080
HG01505
NA12886
NA12877
NA12882
Yq12 repeat
DYZ18_Yq
DYZ1_Yq
DYZ2_Con
Yqhet_2k7bp
Yqhet_3k1bp
Alu element
Alu-1_AluYb8
Alu-2_AluYe5
Alu-3_AluYa5
Alu-4_AluY
Alu-5_AluY
Alu-6_AluYa5
Alu-7_AluY
10 Mb

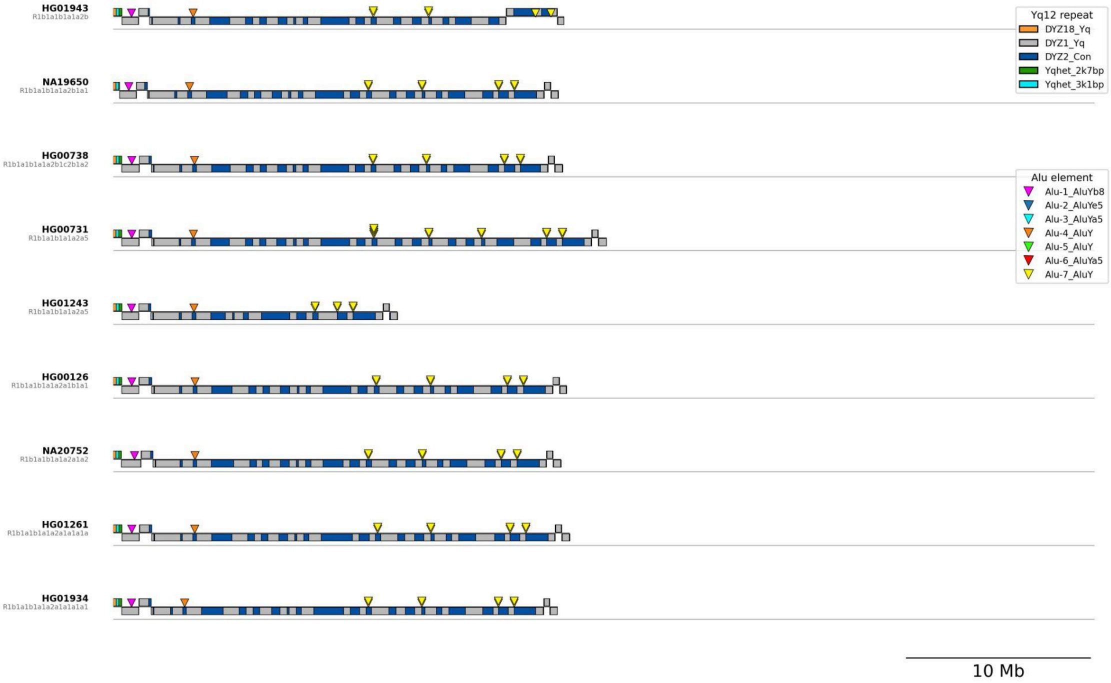


**Supplementary Figure 67. Yq12 structural organization and repeat subunit composition across fully assembled samples.** Shown are 84 QC-passing Yq12 assemblies (no unresolved N gaps, no ERRSTRUCT QC flags in the region, and no Yq12 repeat motifs detected on non-primary contigs). Samples are ordered by a Y chromosome SNV-based phylogeny (**Suppl. Fig. 1**, see **Methods**) with haplogroup A at the top. For each sample (row), colored blocks depict the linear arrangement of Yq12 repeat subarrays (legend) along the assembled Yq12 region. Blocks are mirrored about a central midline to indicate orientation (sense/+orientation above the midline; antisense/-orientation below). Additionally, the location of seven unique *Alu* element insertions, and their subsequent duplications, are shown as triangles (*Alu* subfamily shown in legend).

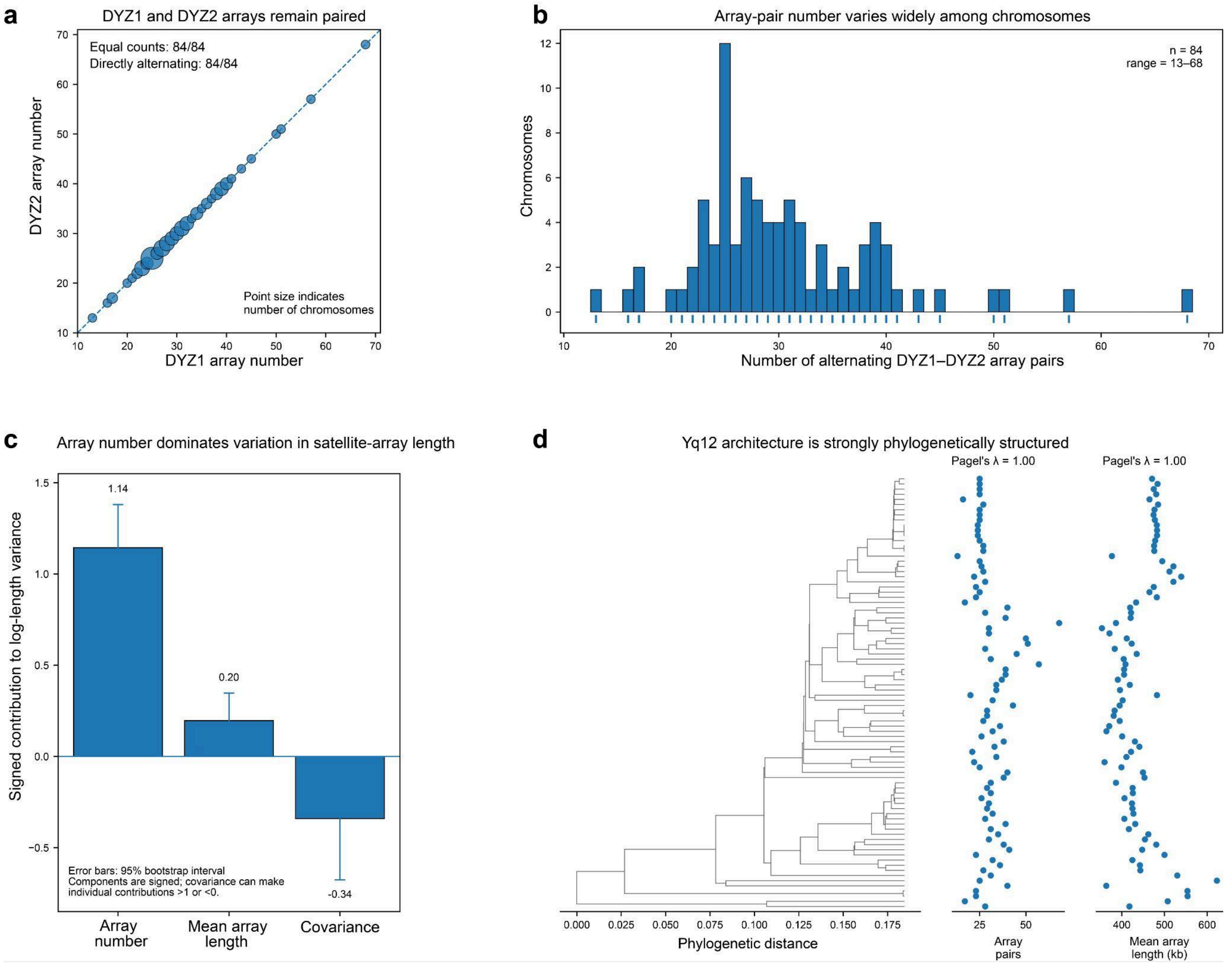


**Supplementary Figure 68. Yq12 size variation occurs within a conserved and phylogenetically structured satellite architecture. a,** Relationship between the number of *DYZ1* and *DYZ2* arrays across the 84 QC-passing Yq12 assemblies. All chromosomes contained equal numbers of *DYZ1* and *DYZ2* arrays, and all arrays were arranged in directly adjacent alternating order. Point size indicates the number of chromosomes sharing an identical *DYZ1–DYZ2* array-count combination; the dashed line indicates a 1:1 relationship. **b,** Distribution of the number of alternating *DYZ1–DYZ2* array pairs per Y chromosome. Total array-pair number varied from 13 to 68 across the 84 assemblies; rug marks indicate individual chromosomes. **c,** Variance decomposition of log-transformed total *DYZ1/DYZ2* satellite-array length into contributions from array number, mean physical array length and their covariance. Array number was the dominant component (variance fraction = 1.144, 95% bootstrap interval 0.967–1.381), whereas mean array length contributed substantially less (0.197, 0.118–0.348). The covariance between the two components was negative (−0.341, −0.677 to −0.126), indicating that chromosomes with more arrays tended in part to contain smaller individual arrays. Components are signed and therefore individual contributions can exceed 1 or be negative when covariance is present. Error bars indicate 95% bootstrap intervals. **d,** Y-chromosome phylogeny of the 84 QC-passing assemblies aligned with the number of alternating array pairs and mean physical *DYZ* array length for each chromosome. Both architectural traits showed strong phylogenetic structure (Pagel's λ = 1.00 for both total array number and mean array length).

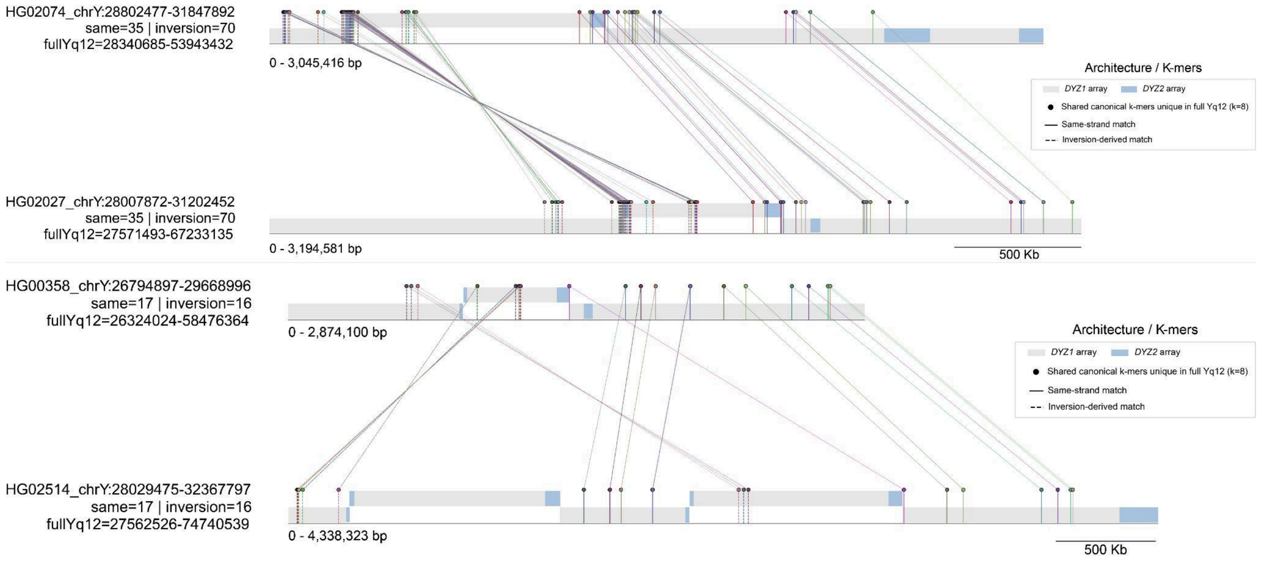


**Supplementary Figure 69. Kmer signatures within Yq12 proximal inversions.** Paired Yq12 subregions from phylogenetically closely related Y chromosome assemblies were compared using canonical k-mers (k = 8) that are unique within each sample's full Yq12 and shared between regions. Tracks display subregion coordinates with underlying Yq12 arrays (*DYZ1*, ***grey***; *DYZ2*, blue) plotted strand-aware (sense: ***top***; antisense: ***bottom***). Shared k-mers are shown as colored lollipops marking their positions. Solid lines indicate same-strand matches, whereas dashed lines denote inversion-derived matches, with connectors linking identical k-mers across regions. Only k-mers occurring exactly once in both the full Yq12 and each subregion are retained. Results highlight inversions within the *proximal* (centromeric side) inversion region, with breakpoints existing within *DYZ1* (***grey***) subunits.

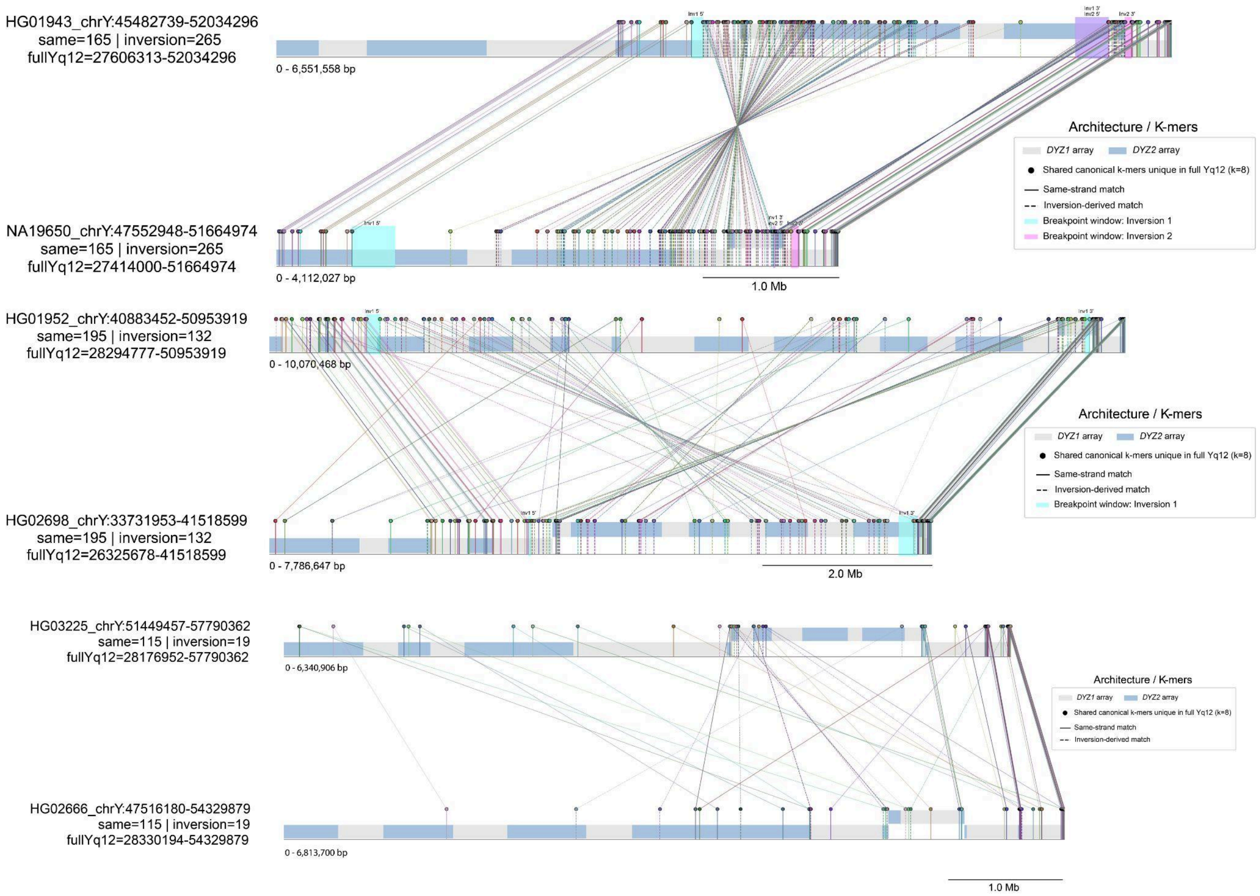


**Supplementary Figure 70. Kmer signatures within Yq12 distal expanded inversions.** Paired Yq12 subregions from two closely related Y chromosome assemblies (nearest neighbors in the phylogeny) were compared using canonical k-mers (k = 8) that are unique within each sample's full Yq12 and shared between regions. Tracks display subregion coordinates with underlying Yq12 arrays (*DYZ1*, ***grey***; *DYZ2*, ***blue***) plotted strand-aware (sense: ***top***; antisense: ***bottom***). Shared k-mers are shown as colored lollipops marking their positions. Solid lines indicate same-strand matches, whereas dashed lines denote inversion-derived matches, with connectors linking identical k-mers across regions. Only k-mers occurring exactly once in both the full Yq12 and each subregion are retained. Results show hidden inversions within the 'expanded' canonical *distal* (PAR2 side) inversion region of three samples (HG01943, HG02698, and HG03225), with breakpoints existing within *DYZ1* (***grey***) subunits. For HG01943 and HG02698, breakpoint intervals were defined and are highlighted by cyan and magenta overlays.

### *De novo* germline mutations

Contributing authors: Pille Hallast, Mark Loftus

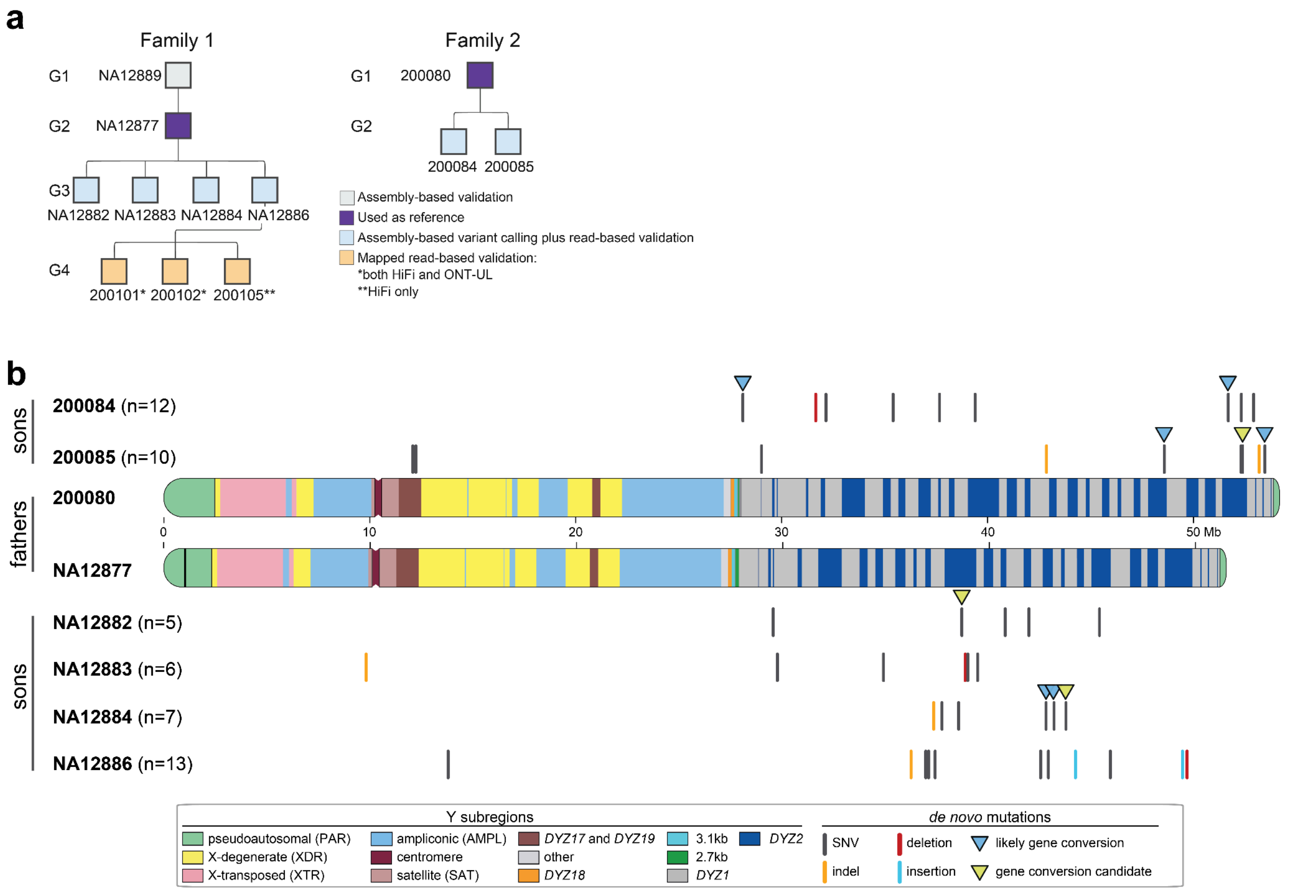


**Supplementary Figure 71. CEPH pedigrees analyzed and distribution of identified DNMs across the Y chromosome. a,** Structure of the two CEPH pedigrees analyzed, indicating how each sample was used. **b,** Structures of the paternal Y chromosomes, shown in the middle, with DNMs identified in each son plotted either above (200080 pedigree) or below (NA12877 pedigree). For *de novo* SNVs in Yq12, the gene-conversion classification is also indicated by triangles.

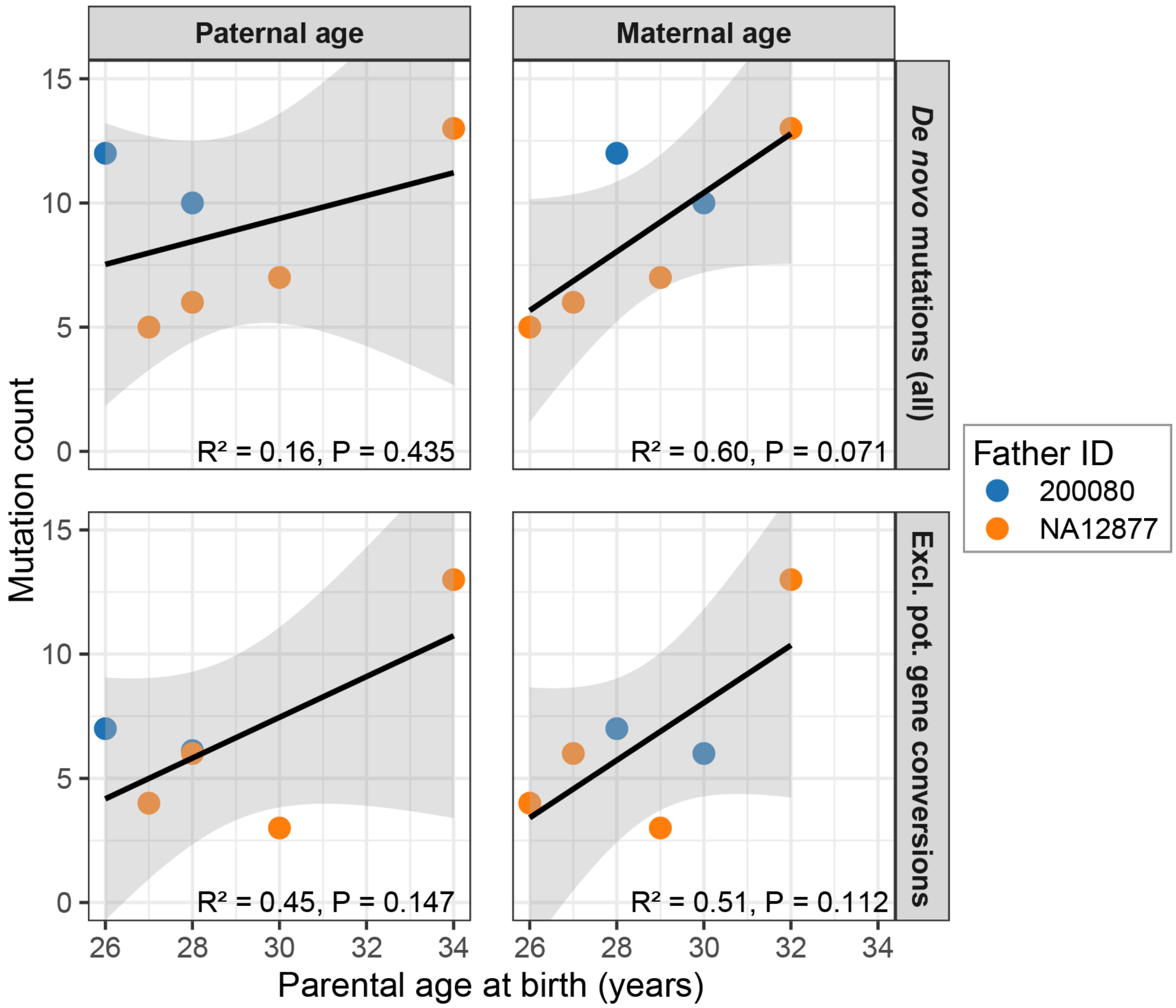


**Supplementary Figure 72. Relationship between parental age and the number of *de novo* mutations.** Scatter plots showing the relationship between paternal age (***left***) or maternal age (***right***) at birth and the number of *de novo* mutations identified per individual: total *de novo* mutations (***top***) and *de novo* mutations excluding SNVs classified as likely or potential gene-conversion events (***bottom***). Each point represents one offspring; sons of 200080 are shown in blue and sons of NA12877 in orange. Solid lines indicate linear regression fits, and shaded regions indicate the 95% confidence intervals. $R^2$ and two-sided P values shown in each panel were obtained from the corresponding simple linear regression model.

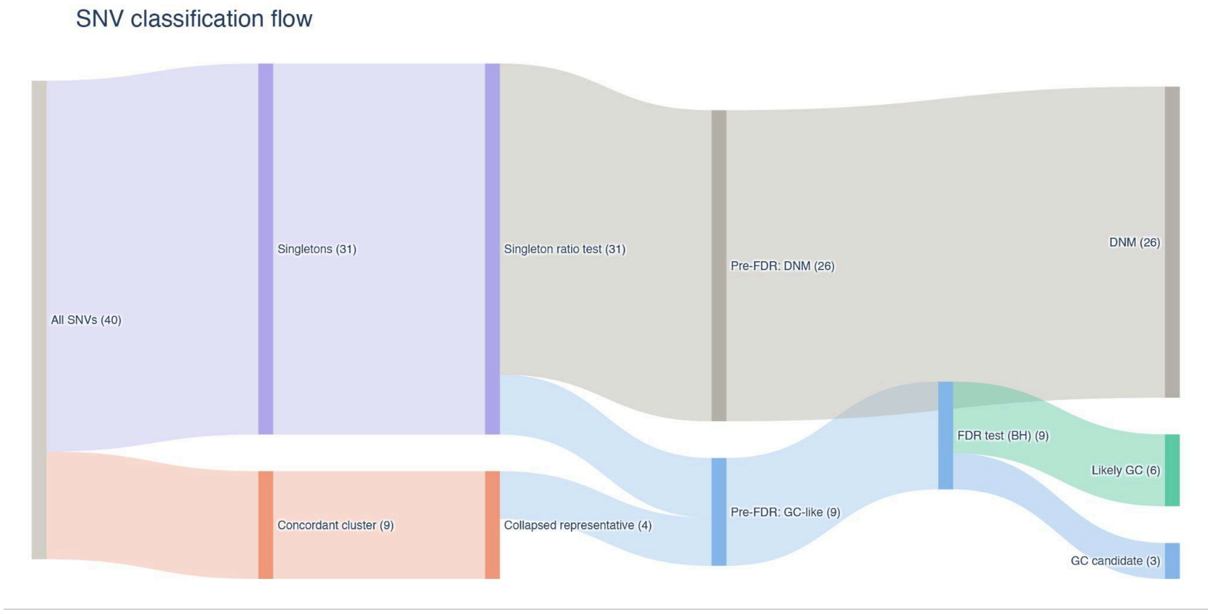


**Supplementary Figure 73. Classification of 40 candidate Yq12 *de novo* SNVs through the donor-discovery pipeline.** Sankey diagram showing the flow of 40 candidate SNVs in the *DYZ1* and *DYZ2* arrays through distance clustering (grouping), deduplication, and final assignment. SNVs entered the pipeline as singletons and were assigned as members of multi-SNV clusters defined at a 300-bp distance threshold. Fully concordant clusters were collapsed to single representative events, with 9 clustered SNVs reduced to 4 representatives and thereby decreasing the total from 40 initial SNVs to 35 nonredundant events. Gene conversion (GC) candidate events were subjected to Benjamini–Hochberg correction, with 2 of 5 single SNV FDR-tested events promoted to 'Likely_GC'. Together with the 4 representatives of fully concordant clusters, this produced final classification counts of 26 DNMs, 6 'Likely_GC' events, and 3 'GC_candidate' events. Ribbon widths are proportional to event number, and the narrowing between the initial and post-deduplication stages reflects cluster collapse rather than event loss.

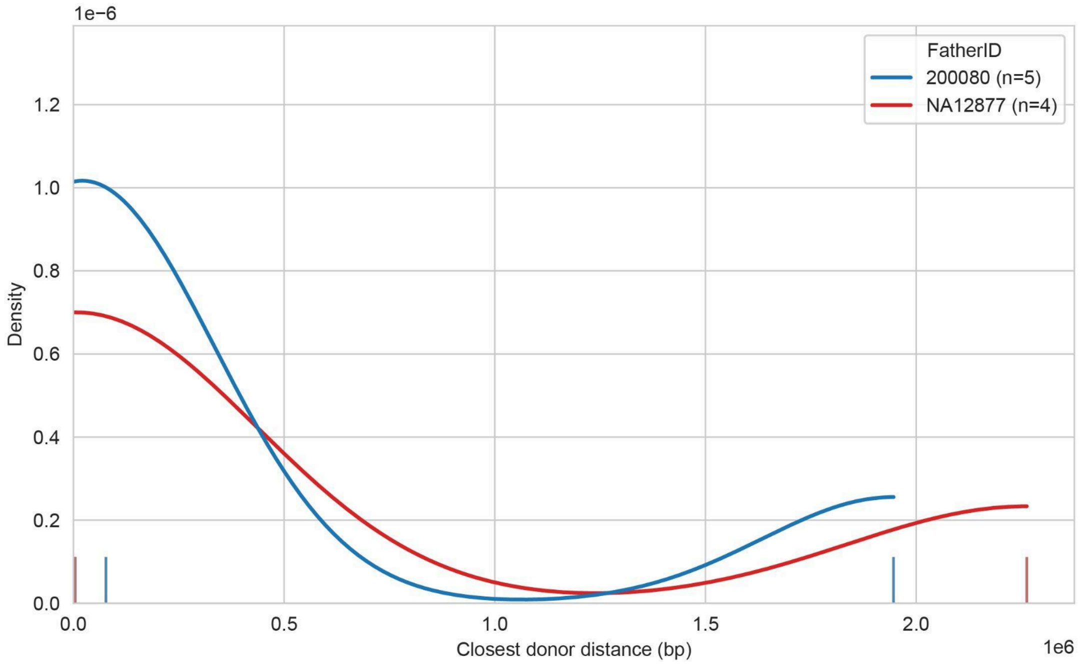


**Supplementary Figure 74. Distribution of distances to the closest homologous donor across fathers.** Kernel density plots show the distribution of closest donor distances from the DNMs for 'GC_candidate 'and 'Likely_GC' events stratified by father. Distances were calculated as the genomic distance (bp) from each DNM event to its nearest homologous donor. Rug marks along the x-axis indicate events classified as 'Likely_GC', shown in lighter shades matched to the corresponding father. The number of events contributing to each father-specific density curve is indicated in the legend. Density estimates were generated independently for each father (common_norm = False) using a bandwidth adjustment of 0.5.

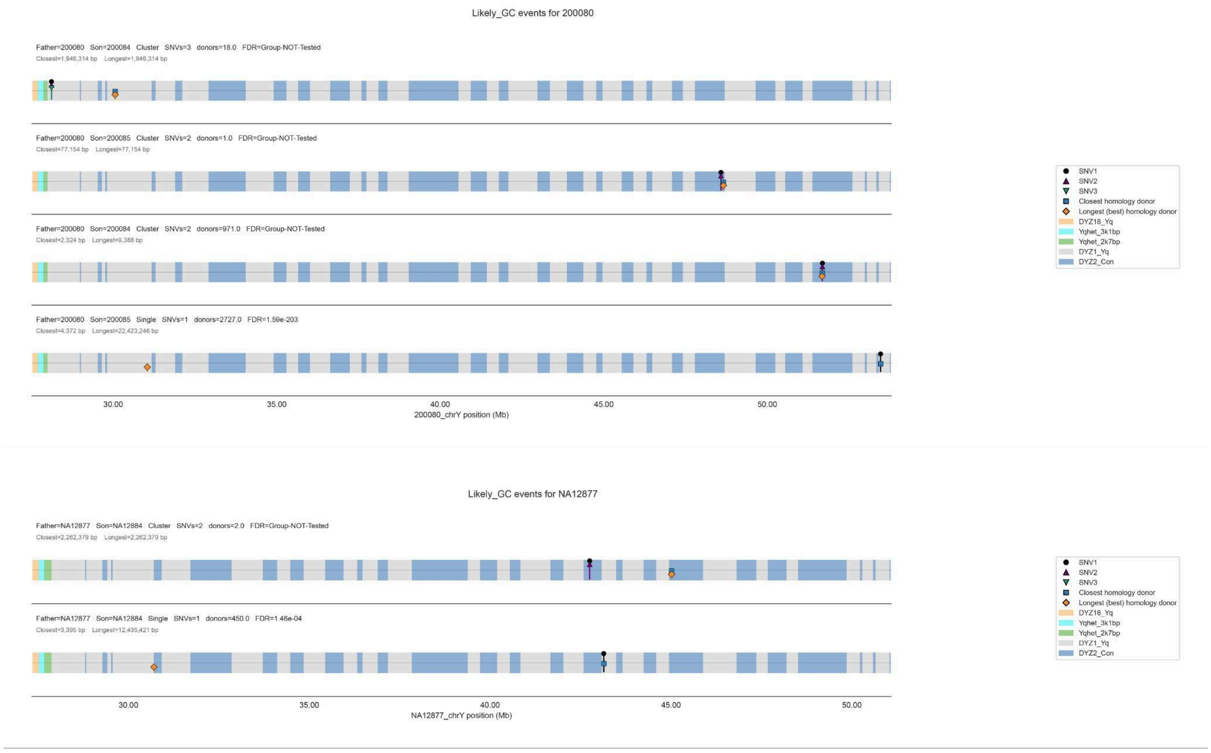


**Supplementary Figure 75. Genomic positions of likely gene conversion events and homologous donor sequences within Yq12.** For each father, likely gene conversion events (Likely_GC) are shown as separate horizontal tracks across the corresponding paternal Y-chromosomal interval. Colored background blocks indicate the local Yq12 repeat architecture, including *DYZ1*, *DYZ2*, and other annotated satellite/repeat classes. Lollipop markers denote the focal DNM SNV positions within each event: SNV1 is shown as a black circle, SNV2 as a purple triangle, and SNV3 as a green inverted triangle. The closest homologous donor on the father's assembly is shown as a blue square, and the longest/best homologous donor is shown as an orange diamond, each plotted at the midpoint of the donor interval. Track labels report the father–son pair, event group size, number of SNVs shown, number of matching donors, FDR-adjusted gene-conversion p-value, and distances to the closest and longest homologous donors. Coordinates are shown in megabases along the paternal Y chromosome.

### Repeat architecture shapes epigenetic organization across palindromic and ampliconic sequence

#### Palindromes

Contributing authors: Karol Pal and Kateryna D. Makova

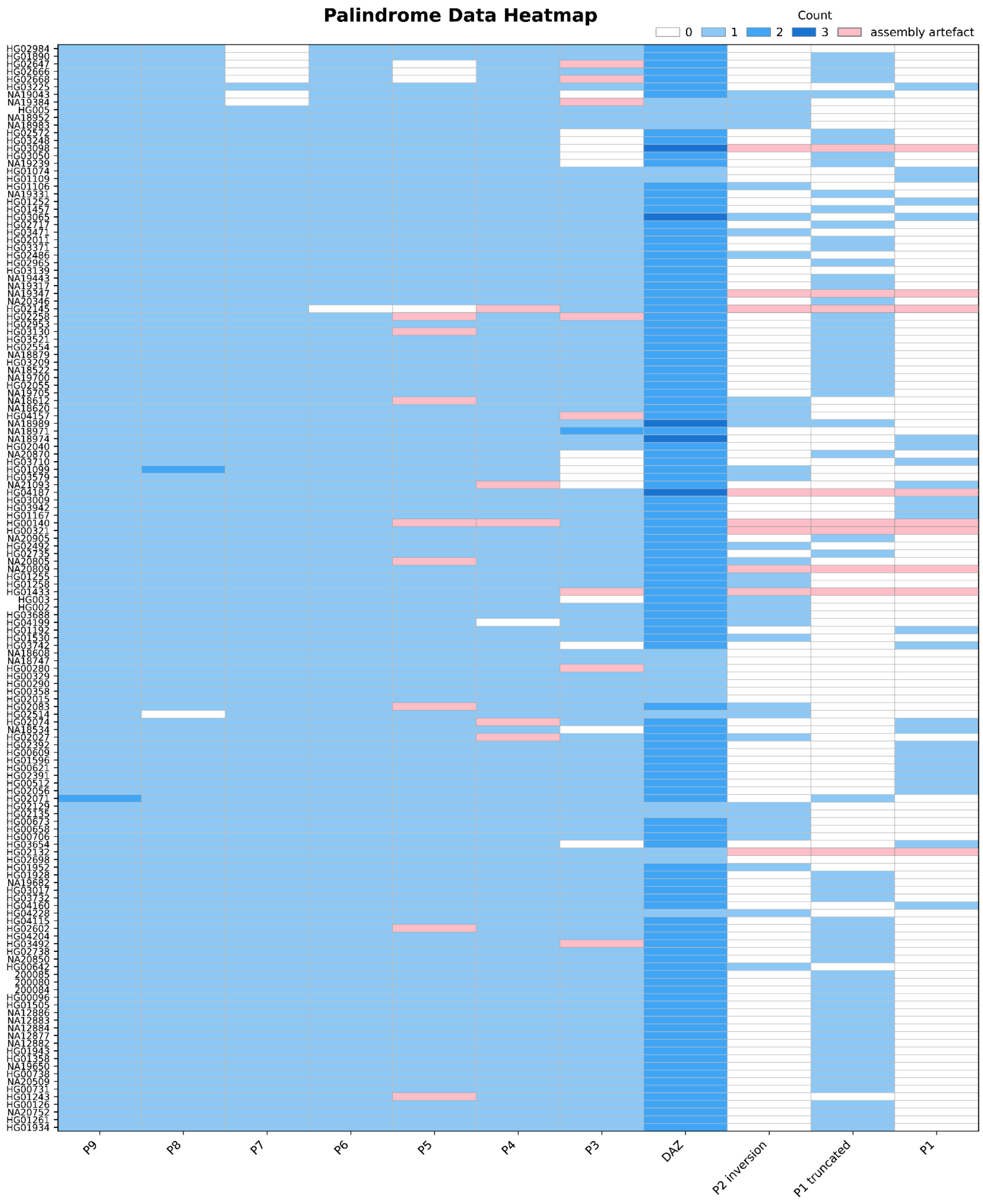

**Supplementary Figure 76. Heatmap of palindromes occurrences.** Heatmap displaying the number of palindrome copies identified at each of ten Y chromosome palindrome loci (P1–P9 and *DAZ*). Rows ordered based on phylogeny. Color intensity indicates copy number. Palindrome P1 comprises two disconnected components, because inter-arm homology is often broken by a repetitive region resulting in a truncated palindrome. Pink color marks palindrome sequences which partially map to unplaced contigs which prevents their proper identification by PALINDROVER[35]. The rest of the white cells are mostly caused by biological variability such as indels in the arms of palindromes preventing proper palindrome identification or shorter palindromes arms preventing assignment to proper cluster.

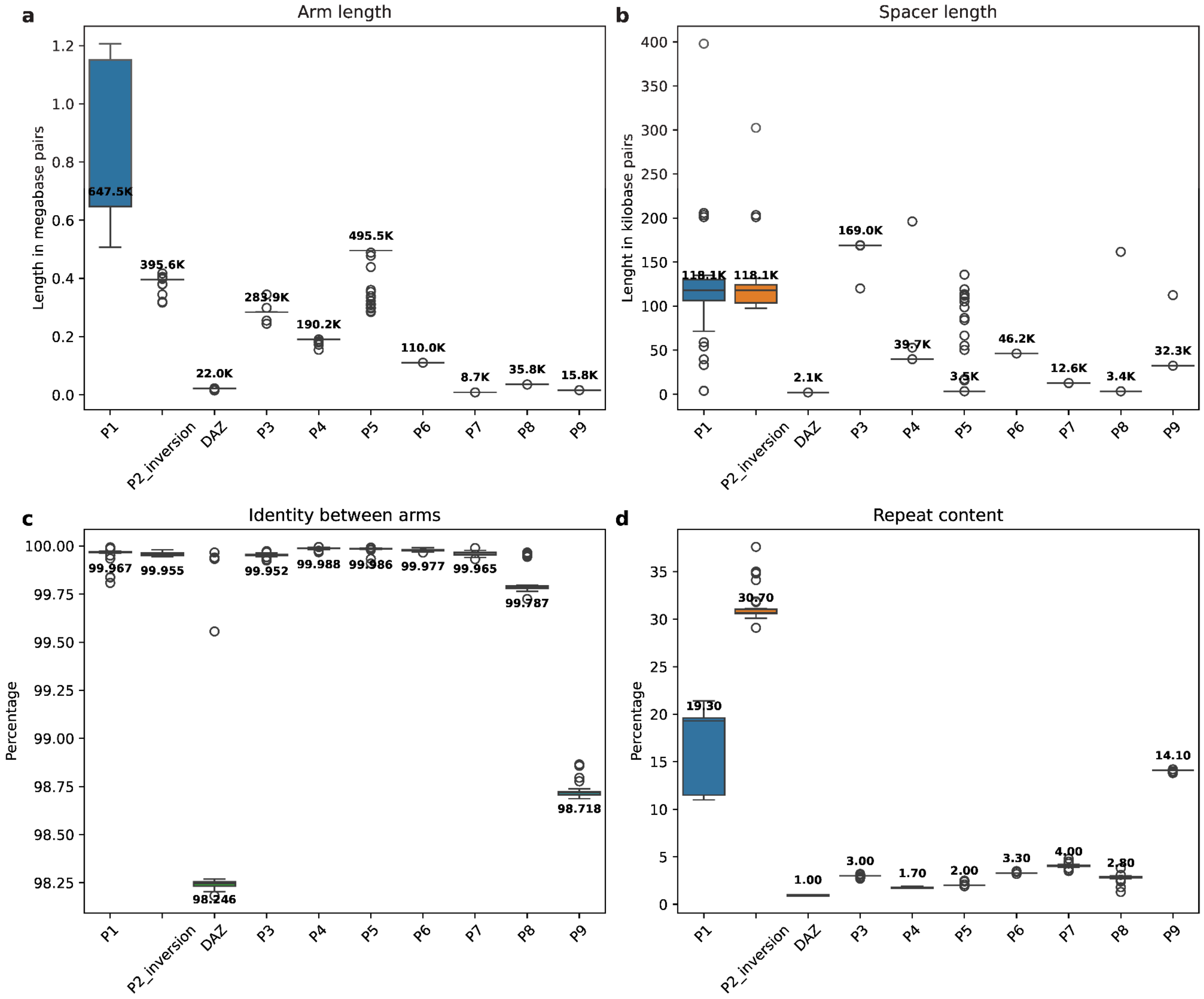


**Supplementary Figure 77. Summary statistics of palindrome groups.** Four-panel boxplot with summary statistics of Y chromosome palindromes at ten loci (P1, P2_inversion, *DAZ*, P3–P9). **a,** Arm length (kb). **b,** Spacer length (kb). **c,** Sequence identity between palindrome arms (%). **d,** Repeat content of palindrome arms (%). Statistics calculated based on the following counts per palindrome group; P1: 89, P2_inversion: 35, *DAZ*: 271, P3: 120, P4: 136, P5: 130, P6: 141, P7: 135, P8: 142, P9 (Q6): 143.

### Gene conversion

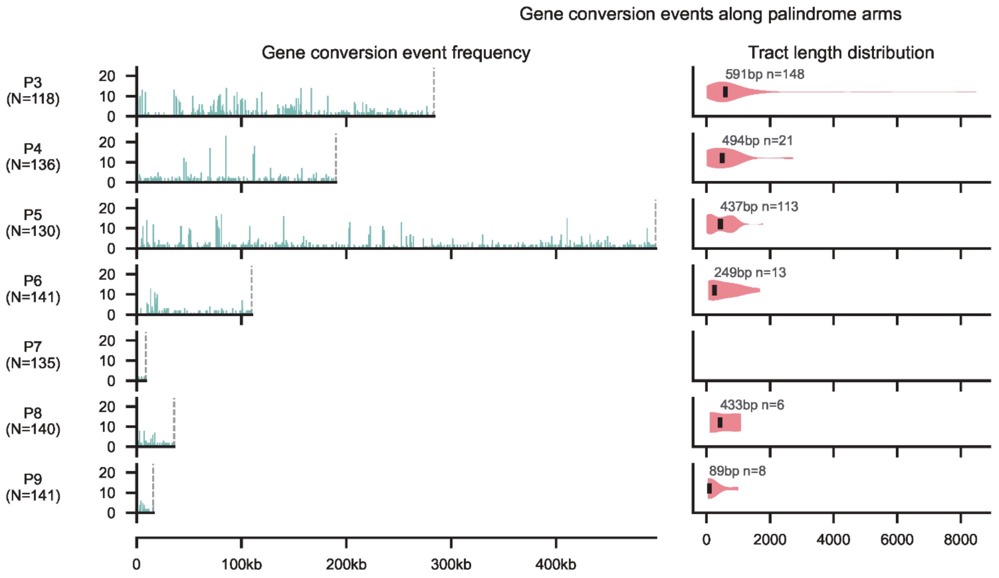


Summary statistics

| | events | events/kb | reversions | fixations | GC bias (%) |
|---|---|---|---|---|---|
| **P3** | 1133 | 4.00 | 242 | 188 | 60 |
| **P4** | 460 | 2.42 | 148 | 138 | 57 |
| **P5** | 1169 | 2.36 | 387 | 353 | 60 |
| **P6** | 225 | 2.05 | 62 | 59 | 61 |
| **P7** | 18 | 2.08 | 8 | 7 | 43 |
| **P8** | 113 | 3.17 | 48 | 47 | 57 |
| **P9** | 121 | 7.71 | 10 | 13 | 56 |
| **Sum** | **3239** | **2.84** | **905** | **805** | **59** |

**Supplementary Figure 78. Gene conversion summary.** Left column: frequency of independent gene conversion events per position along each palindrome arm. Middle column: distribution of estimated gene conversion tract lengths per palindrome, defined as clusters of ≥2 gene conversion events on the same ancestral node within 1 kb; median tract length (bp) annotated above each distribution. Right column: per-palindrome summary statistics including total gene conversion events, event rate (events/kb), number of gene conversion events resolving pseudoheterozygosity toward the ancestral allele (reversions) or derived allele (fixations), GC-bias expressed as (AT to GC) / (AT to GC) + (GC to AT), and number of putative conversion tracts. Summary row reports pooled values across all palindromes.

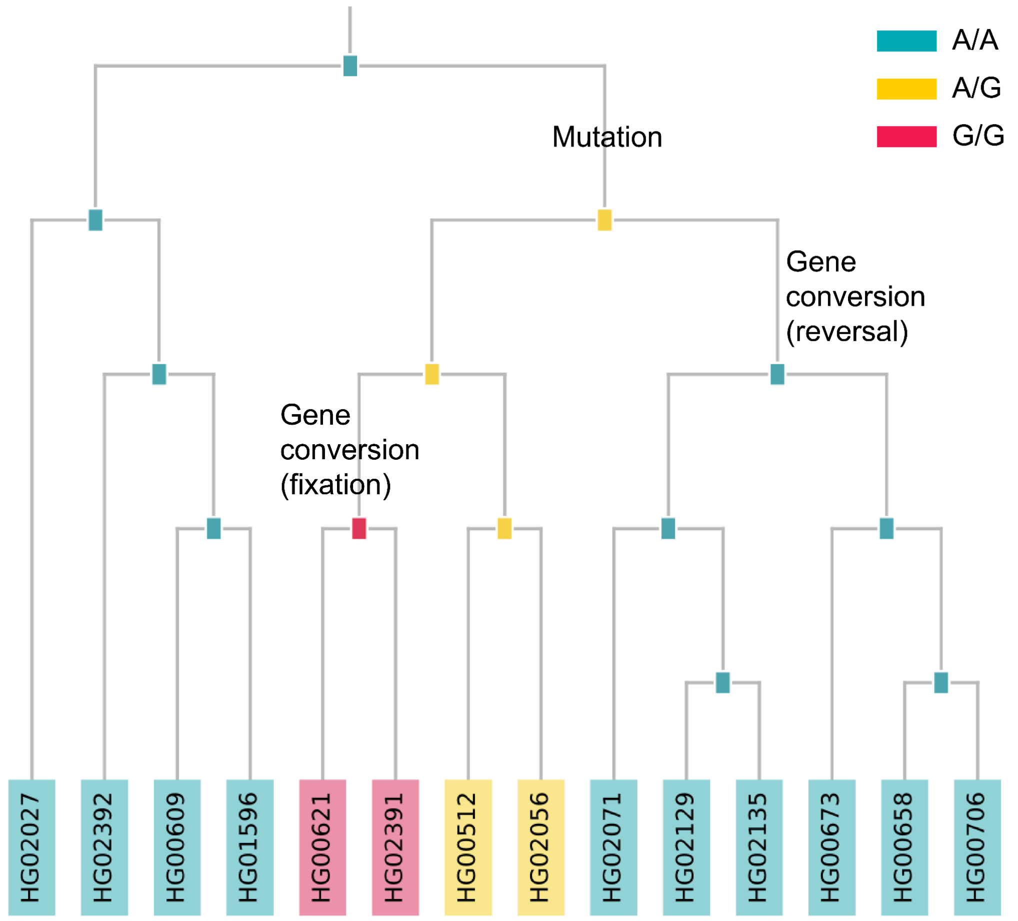


**Supplementary Figure 79. Example of gene conversion.** A subtree of a parsimony-reconstructed phylogeny for a single variant site (location 47999 on arm A of palindrome P6 in the HG002 sample), illustrating two independent gene conversion outcomes at the same site. A single mutation event gives rise to pseudoheterozygosity in an ancestral lineage. Two subsequent gene conversion events resolve this pseudoheterozygosity in independent descendant lineages: one converts the pseudoheterozygous state back to the ancestral allele while the other fixes the derived allele. Nodes are coloured by inferred genotype; internal nodes represent ancestral states reconstructed by the Fitch parsimony state reconstruction method[36]; leaf nodes represent observed haploid genotypes.

### Methylation on palindromes and ampliconic genes

Contributing authors: Mariateresa Mazzetto and Monika Cechova

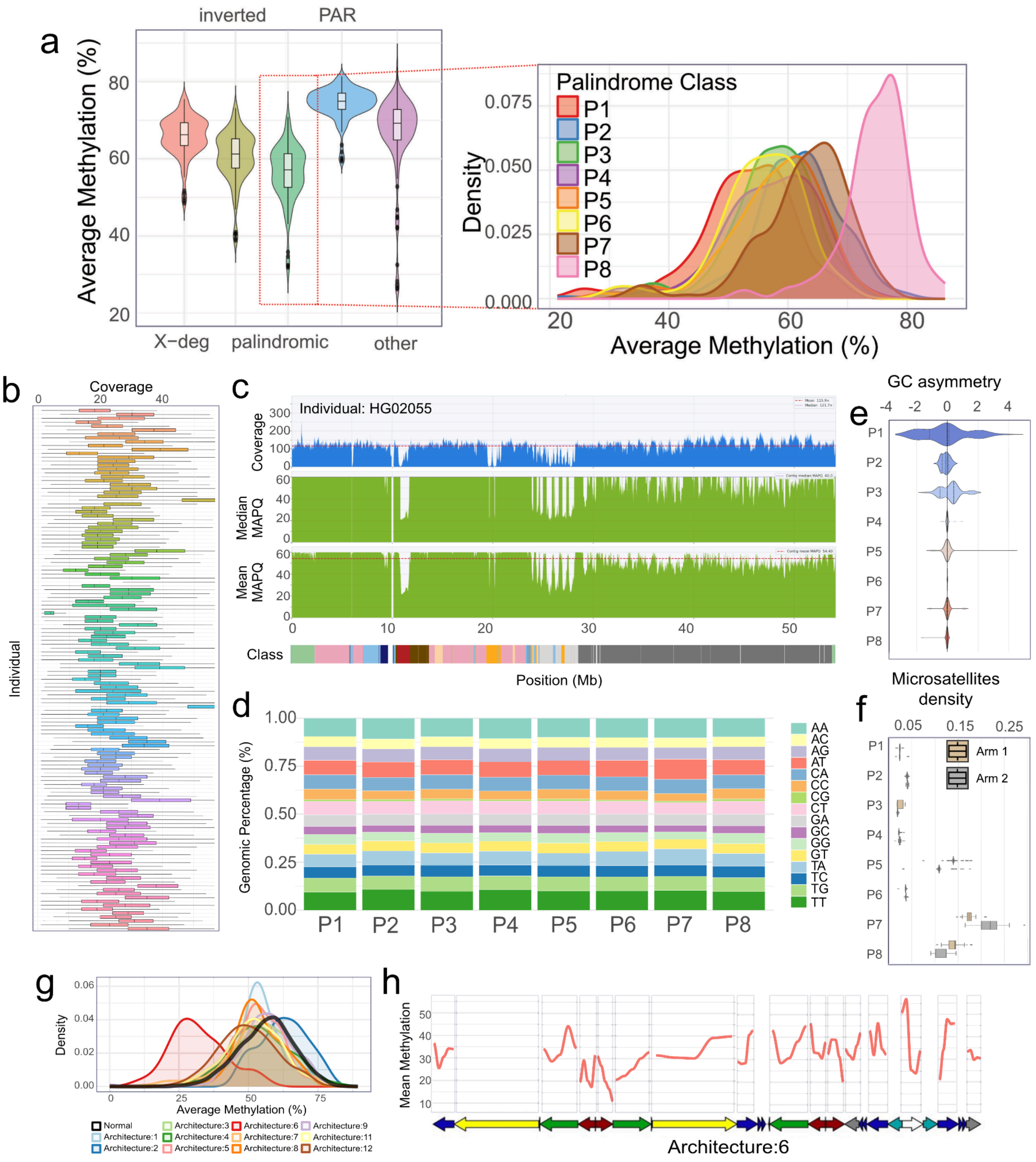


**Supplementary Figure 80. a, Left Panel:** Distribution of per-individual mean CpG methylation (%) across Y chromosome sequence classes. For each individual, methylation values were averaged across all assayed CpGs within each class. Violins show the across-individual density and the embedded box indicates the median and interquartile range. **Right Panel:** Per-individual mean methylation (%) for each palindrome (P1-P8), shown as kernel density estimates to highlight differences among palindrome classes. **b,** Methylation coverage on chromosome Y across individuals. Coverage was measured as the total number of bases; individuals are color-coded. **c,** Coverage and MAPQ variations along chromosome Y in sample HG02055. MAPQ were measured along the genomic area by windows of 5kb with a sliding window of 2.5 kb using raaqa v0.1.1. Both median and mean MAPQ are visualized. Sequence classes defined by Rhie, Nurk, Cechova, Hoyt et al. are also displayed on the lower part of the figure. **d,** Dinucleotide composition of palindromic regions across individuals. Relative frequency of all 16 dinucleotide combinations was plotted for all 8 palindromes. **e,** Distribution of GC content differences between paired palindrome arms (arm1 − arm2) across palindromes (P1–P8). Values are centered at zero, where positive or negative shifts indicate asymmetric GC composition between arms. The violin plots illustrate the variability and magnitude of GC asymmetry across individuals for each palindrome. ANOVA p-value < $2e^{-16}$ **f,** Microsatellite density (fraction of bases annotated as tandem repeats) is shown for each palindrome (P1–P8), separated by arm. Boxplots summarize the distribution of microsatellite density across individuals, highlighting differences in repeat content both between palindromes and between corresponding arms. **g,** Methylation patterns on AZFc regions characterized by unique architectures (**architectures shown in Supplementary Figure 25. 'Distinct AZFc haplotypes'**). Methylation of AZFc regions with different unique architectures (on the x-axis), and methylation on typical architectures is displayed as well. **h,** Methylation pattern on AZFc regions with architecture:6. Methylation was summarized in 5 bins per color block before plotting. AZFc haplotype structure is included below. No methylation signal was detected on the first yellow block on the individual displaying Arch:6.

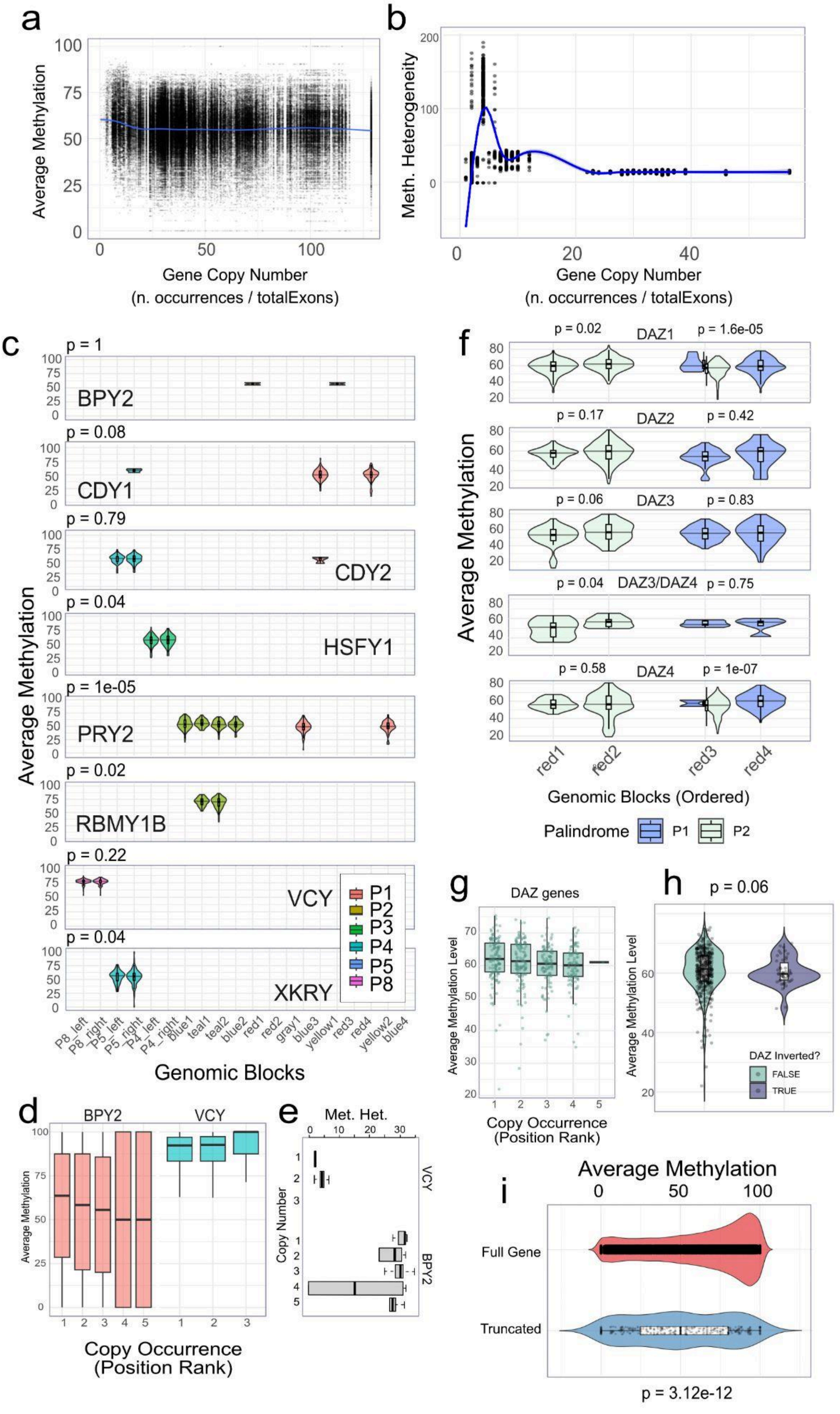

a
Average Methylation
Gene Copy Number
(n. occurrences / totalExons)
b
Meth. Heterogeneity
Gene Copy Number
(n. occurrences / totalExons)
c
Average Methylation
p = 1
BPY2
p = 0.08
CDY1
p = 0.79
CDY2
p = 0.04
HSFY1
p = 1e-05
PRY2
p = 0.02
RBMY1B
p = 0.22
VCY
p = 0.04
XKRY
P1
P2
P3
P4
P5
P8
Genomic Blocks
f
Average Methylation
p = 0.02 DAZ1 p = 1.6e-05
p = 0.17 DAZ2 p = 0.42
p = 0.06 DAZ3 p = 0.83
p = 0.04 DAZ3/DAZ4 p = 0.75
p = 0.58 DAZ4 p = 1e-07
red1 red2 red3 red4
Genomic Blocks (Ordered)
Palindrome P1 P2
d
BPY2 VCY
Average Methylation
Copy Occurrence
(Position Rank)
e
Met. Het.
Copy Number
VCY
BPY2
g
DAZ genes
Average Methylation Level
Copy Occurrence
(Position Rank)
h
p = 0.06
Average Methylation Level
DAZ Inverted?
FALSE
TRUE
i
Average Methylation
Full Gene
Truncated
p = 3.12e-12

**Supplementary Figure 81. a-b, Methylation summary metrics as a function of ampliconic gene copy abundance. a,** Scatterplot of mean CpG methylation (%) versus exon copy number count per gene. Each point represents a gene instance in an individual. The ***blue*** smoothed trend line summarizes the average methylation as a function of the X-axis copy metric. **b,** Methylation heterogeneity standard deviation versus the same copy number abundance metric, with a LOESS smoother (***blue***). **c,** Ampliconic gene methylation varies by ampliconic block and palindrome context. For each ampliconic gene family (panels ordered alphabetically), violin plots show the distribution of per-individual mean CpG methylation (%) measured within the gene body (or gene-overlapping sequence) stratified by the ampliconic block / palindromic segment in which the gene copy occurs (x-axis, blocks ordered along the ampliconic region). Colors indicate the palindrome (P1-P8) associated with each block. P-values shown in each panel summarize overall difference across blocks for that gene. Overall tests performed using ANOVA with Bonferroni correction for multiple group comparisons. Pairwise comparisons used the t-test method. **d-e,** Copy number specific methylation and methylation heterogeneity for BPY2 and VCY. **d,** Distributions of per-individual mean CpG methylation (%) for BPY2 and VCY stratified by paralog copy rank, where copies are ordered by genomic position along the ampliconic region (copy rank 1 = most proximal, increasing rank = more distal). **e,** Methylation heterogeneity for BPY2 and VCY stratified by copy number. Together, these plots show that methylation levels (and/or heterogeneity) can differ among paralog copies of the same ampliconic gene family. **f-h,** Copy number and structure-associated methylation differences across DAZ gene copies. **f,** DAZ copy methylation by ampliconic block. Per-individual mean CpG methylation (%) for DAZ gene copies stratified by ampliconic block (x-axis; palindromes correspond to colored segments ordered along the DAZ-containing ampliconic region). Violin plots show the across-individual distribution. Embedded boxplots indicate the median and interquartile range. Palindrome class if indicated by color (P1 vs P2). P-values were obtained using the T-test for pairwise comparisons, and ANOVA for non-pairwise comparisons. **g,** DAZ methylation by copy genomic position. Mean CpG methylation (%) for all DAZ family gene copies grouped by copy rank (copies ordered by genomic location within the DAZ ampliconic region). Points represent individuals and boxplots summarize the distribution across individuals. **h,** DAZ methylation by inversion status. Comparison of DAZ methylation between individuals/assemblies with or without a DAZ region inversion. Violin plots show the distribution of per-individual mean methylation across DAZ copies in each inversion class. Together, these panels show that DAZ methylation varies across copies and is associated with local genomic context (copy rank) and structural state (inversion). **i,** DAZ methylation by deletion status. Comparison of DAZ methylation between individuals/assemblies with or without a DAZ region deletion. Violin plots show the distribution of per-individual mean methylation across DAZ copies in each class. Together, these panels show that DAZ methylation is associated with structural integrity (deletion).

Non-canonical structures and G4-motifs

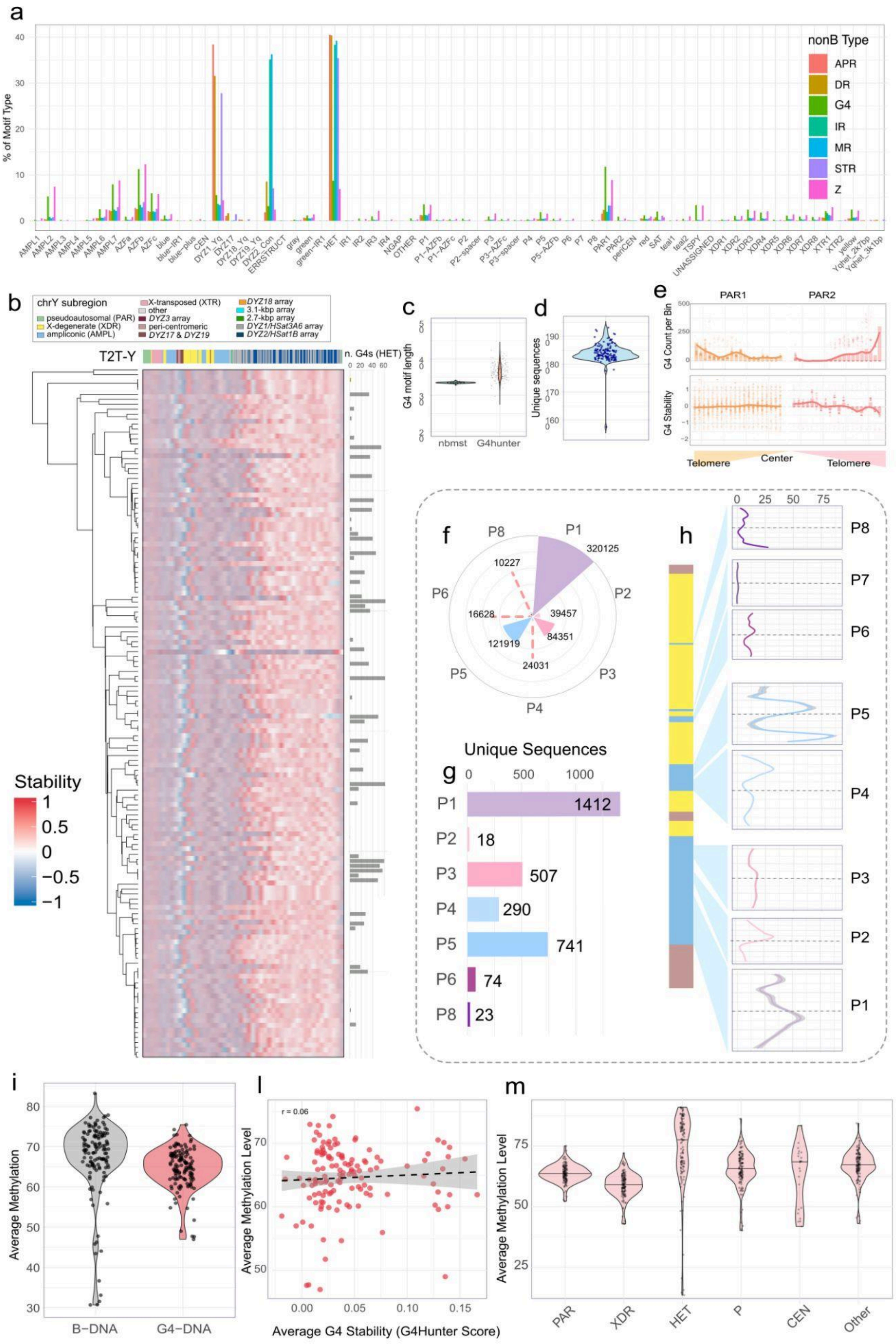

**Supplementary Figure 82. Distribution and structural organization of G-quadruplex (G4) motifs across the human Y chromosome. a,** Relative abundance of predicted non-B DNA motifs across Y chromosome sequence classes and repeat families. Bars represent the percentage contribution of each motif type (APR, DR, G4, IR, MR, STR, Z) within genomic subregions, highlighting strong enrichment of specific structures in repetitive elements and heterochromatic regions. **b,** Heatmap of G4 stability scores (G4Hunter) across telomere-to-telomere Y chromosome assemblies (T2T-Y), clustered by phylogenetic similarity. Rows correspond to individual haplotypes and columns to genomic bins. Top annotation indicates sequence classes, and the right panel shows total G4 counts per haplotype, revealing both conserved patterns and inter-individual variability. The number of G4 motifs on heterochromatin regions (HET) is shown. **c,** Average length of G4 motifs per sample using nbmst and G4hunter. Results are shown without filtering for stability. **d,** Distribution of G4 counts along pseudoautosomal regions (PAR1 and PAR2), revealing distinct positional enrichment patterns along the telomere–center axis rather than a uniform distribution. **e,** G4 stability profiles along PAR1 and PAR2. Values represent normalized G4 stability scores across genomic bins, illustrating regional variation in predicted structural stability. **f,** Radial plot of G4 counts across palindromic regions (P1–P8), illustrating differential enrichment of G4 motifs among palindrome arms. **g,** Number of unique G4-forming sequences identified within each palindrome, indicating substantial variability in sequence diversity across palindromic regions. **h,** Structural profiles of G4 density along representative palindrome arms (P1–P8) after filtering for stability>1. For each palindrome (P1-P8), the numbers of G4s were summarized along the left and right arms after aligning arms by normalized position (0-100% of palindrome length). At each position bin, the values were summarized across individuals using 10 quantiles. Lines show quantile trajectories for the left and right arms. The schematic below indicates the relative order of the palindromes along the Y chromosome inside ampliconic regions (lightblue) and near X-degenerate regions (yellow). **i,** Boxplot with the comparison of average DNA methylation levels between canonical B-DNA regions and G4-forming regions, showing distinct methylation profiles associated with non-B DNA structures. **l,** Relationship between G4 stability and average DNA methylation. Each point represents a genomic region, with the trend line indicating a weak but consistent association between structural propensity and methylation level. **m,** Distribution of average DNA methylation across Y chromosome sequence classes (PAR, XDR, heterochromatic (HET), palindromic (P), centromeric (CEN), and other regions), highlighting differences in epigenetic profiles across genomic contexts.

### Polishing Experiments

Contributing authors: Jana Ebler, Tobias Marschall

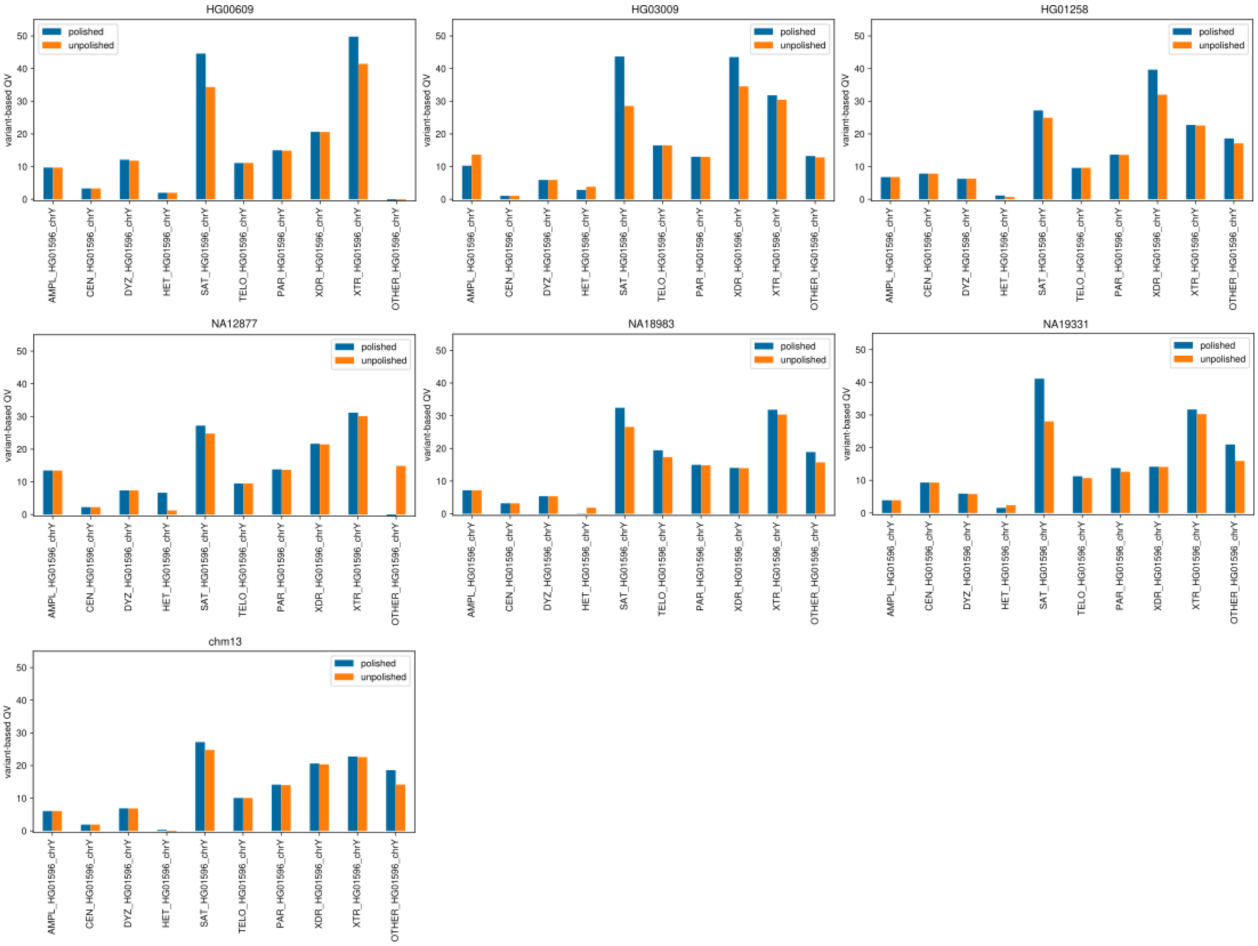


**Supplementary Figure 83. Variant-based QV estimates computed across different annotation regions along Chromosome Y sequences before and after polishing sample HG01596.** The chromosome Y sequence of sample HG01596 was reconstructed based on short reads and low-coverage ONT data with a polishing pipeline initialized with chromosome Y assemblies of six different samples as well as CHM13. Included were the phylogenetically closest sample HG00609, as well as five samples from different continental groups: HG03009 (SAS), HG01258 (AMR), NA12877 (EUR), NA18983 (EAS) and NA19331 (AFR). Additionally, the T2T-CHM13v2Y Chromosome (HG002) is included. The plots show variant-based QVs for different regions that were computed by comparing the polished sequence to the chromosome Y assembly of HG01596.

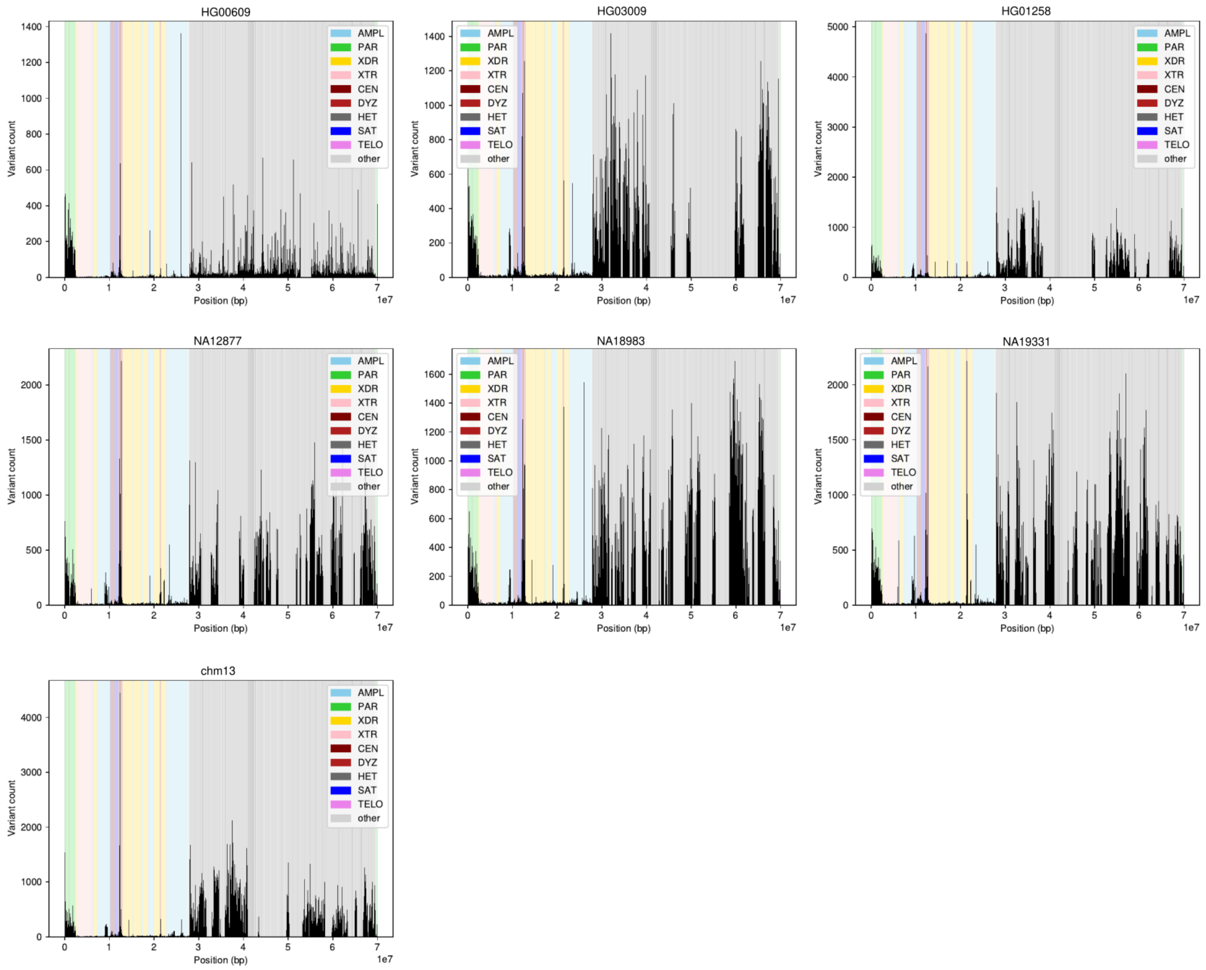


**Supplementary Figure 84. Distribution of variant call counts before polishing.** Shown are variant call counts resulting from the alignment of Y chromosome assemblies of six samples and the CHM13-Y, to the Y chromosome assembly of HG01596. Included were the phylogenetic closest sample HG00609, as well as five samples from different continental groups: HG03009 (SAS), HG01258 (AMR), NA12877 (EUR), NA18983 (EAS) and NA19331 (AFR).

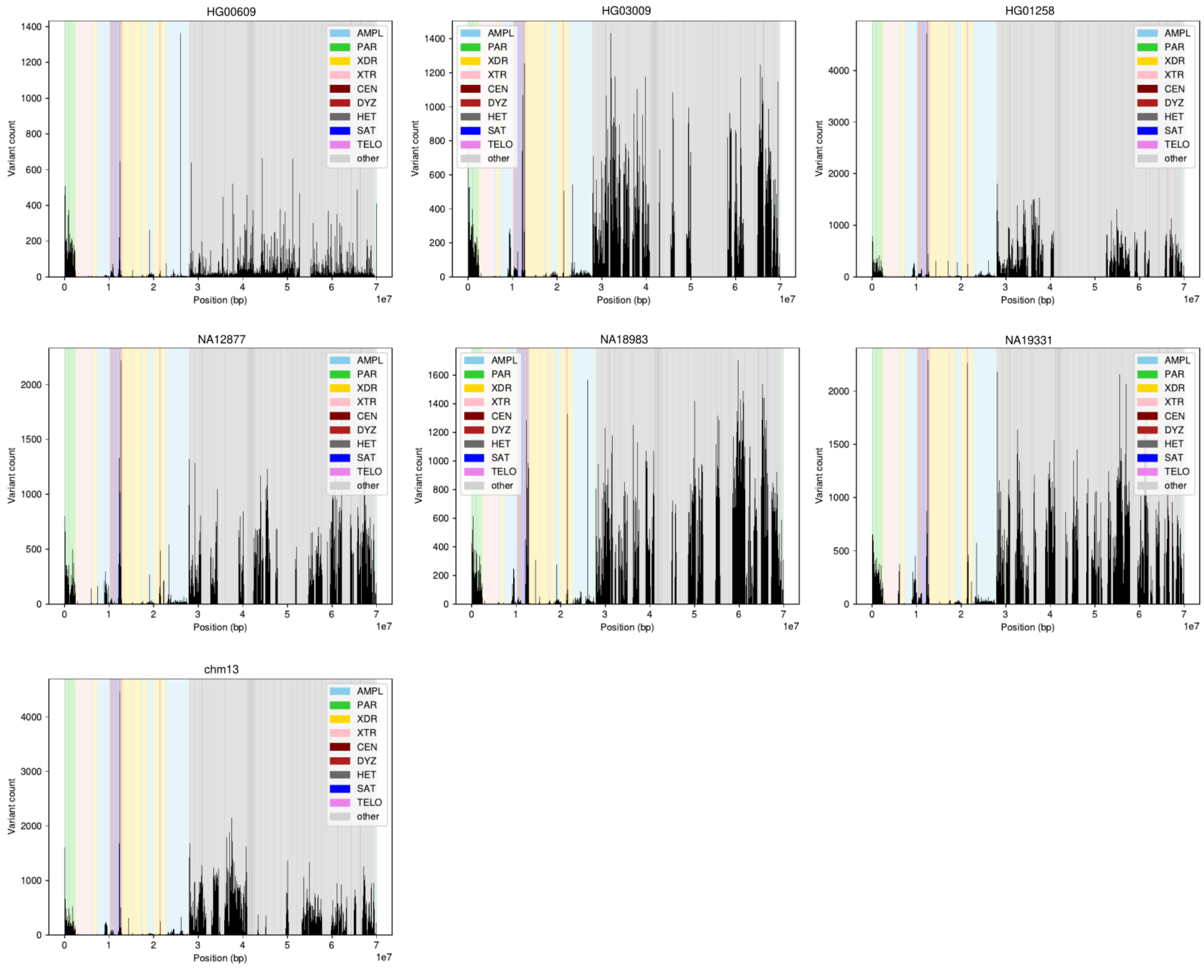


**Supplementary Figure 85. Distribution of variant call counts after polishing.** The Y chromosome sequence of sample HG01596 was reconstructed based on short reads and low-coverage ONT data with a polishing pipeline initialized with Y chromosome assemblies of different samples. Included were the phylogenetic closest sample HG00609, as well as five samples from different continental groups: HG03009 (SAS), HG01258 (AMR), NA12877 (EUR), NA18983 (EAS) and NA19331 (AFR). Additionally, the CHM13-Y (HG002) was included. The plots show variant call counts along these reconstructed haplotypes called from alignments to the chromosome Y assembly of HG01596.

### *RBMY*, *TSPY* and *DAZ* gene expression

Contributors: Gianni V. Martino, Miriam Konkel

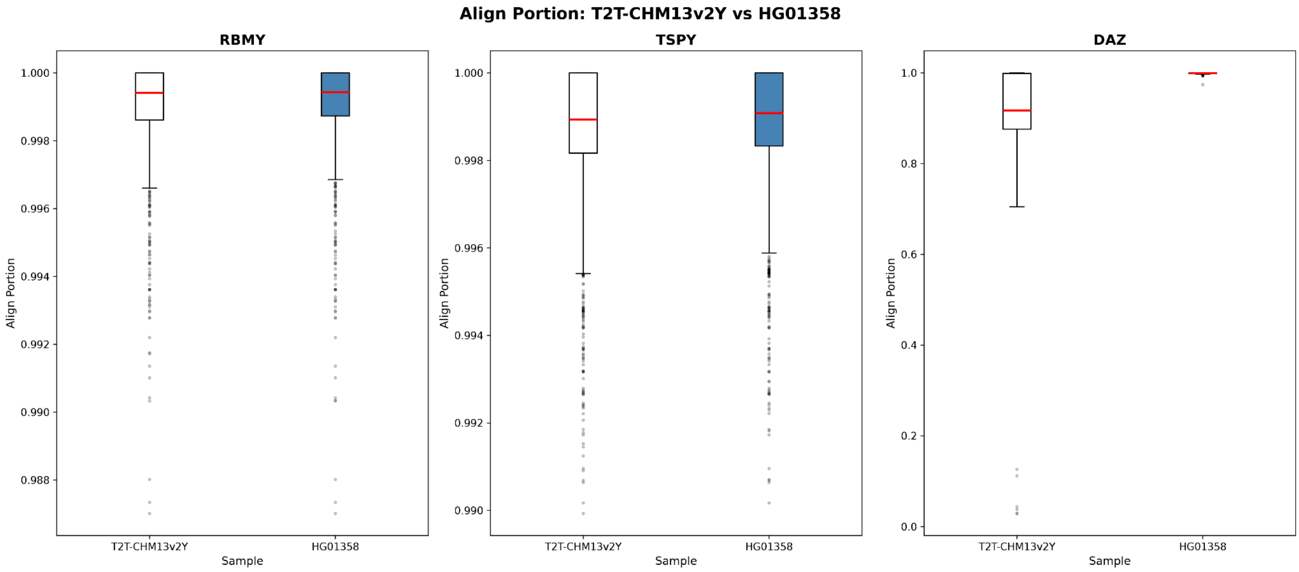


**Supplementary Figure 86. Comparison of T2T-CHM13v2Y vs best assembly alignments.** A box plot illustrating the overall improvement in read alignment quality when utilizing the optimal assembly vs T2T-CHM13v2Y reference for *RBMY*, *TSPY*, and *DAZ* ampliconic genes.

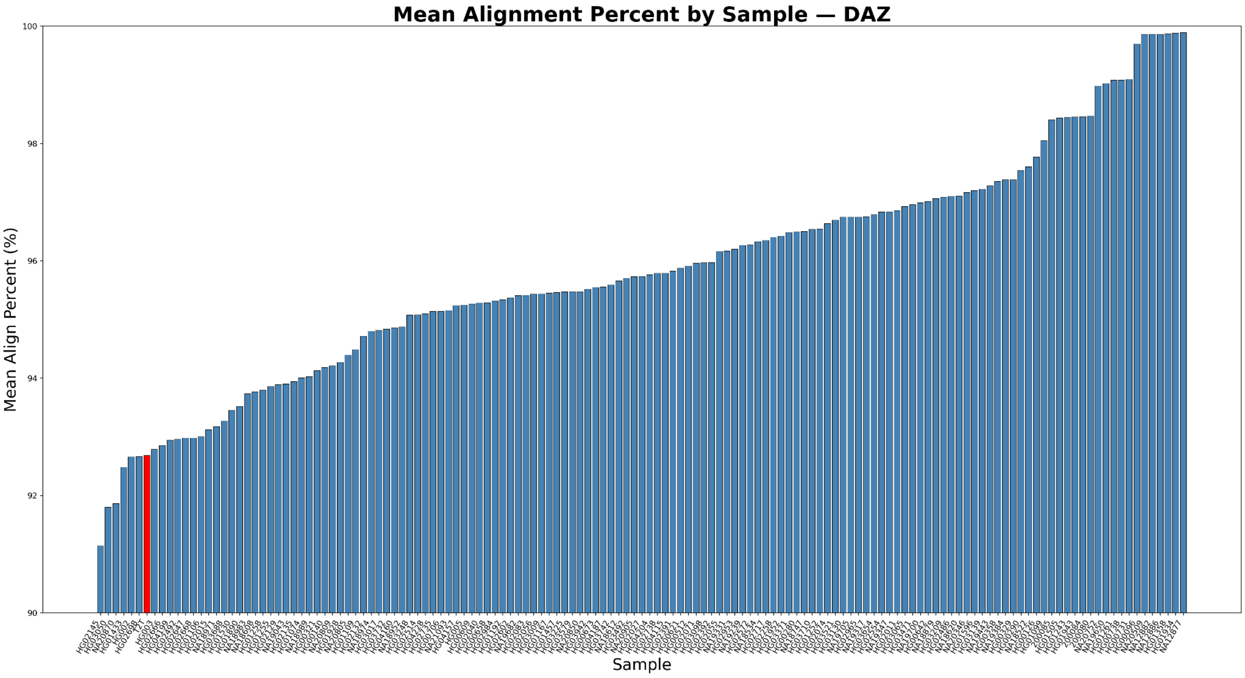


**Supplementary Figure 87. *DAZ* alignment quality distribution by genome.** A comparison of the average percent of DAZ read nucleotides mapped when aligning to 140 individual Y chromosomes and the T2T-CHM13v2Y reference genome. The red bar highlights the position of the T2T-CHM13v2Y reference genome within the curve.

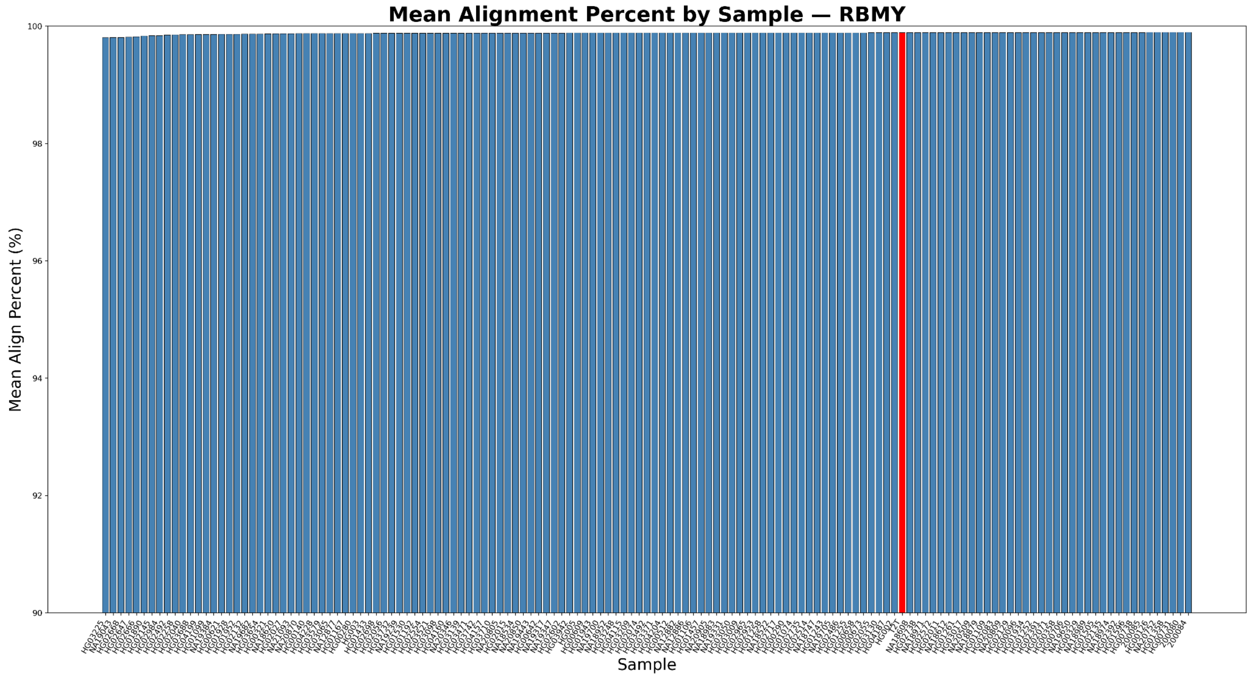


**Supplementary Figure 88. *RBMY* alignment quality distribution by genome.** A comparison of the average percent of *RBMY* read nucleotides mapped when aligning to 140 individual Y chromosomes and the T2T-CHM13v2Y reference genome. The red bar highlights the position of the T2T-CHM13v2Y reference genome within the curve.

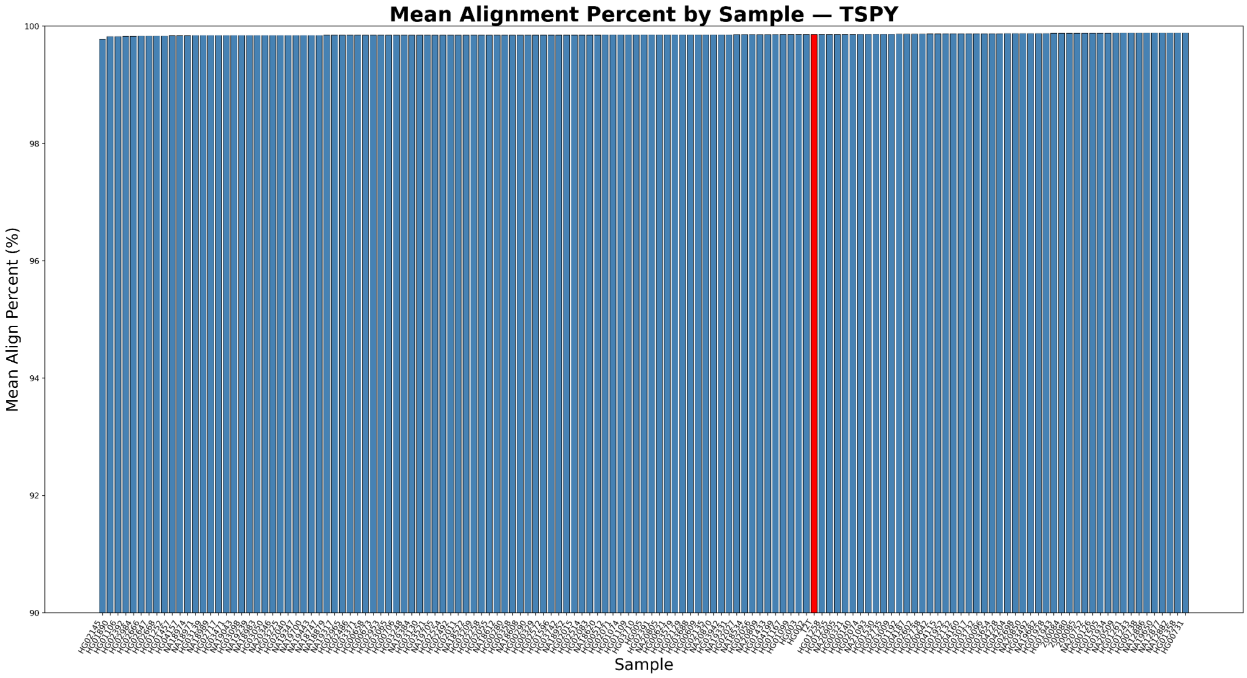


**Supplementary Figure 89. *TSPY* alignment quality distribution by genome.** A comparison of the average percent of *TSPY* read nucleotides mapped when aligning to 140 individual Y chromosomes and the T2T-CHM13v2Y reference genome. The red bar highlights the position of the T2T-CHM13v2Y reference genome within the curve.

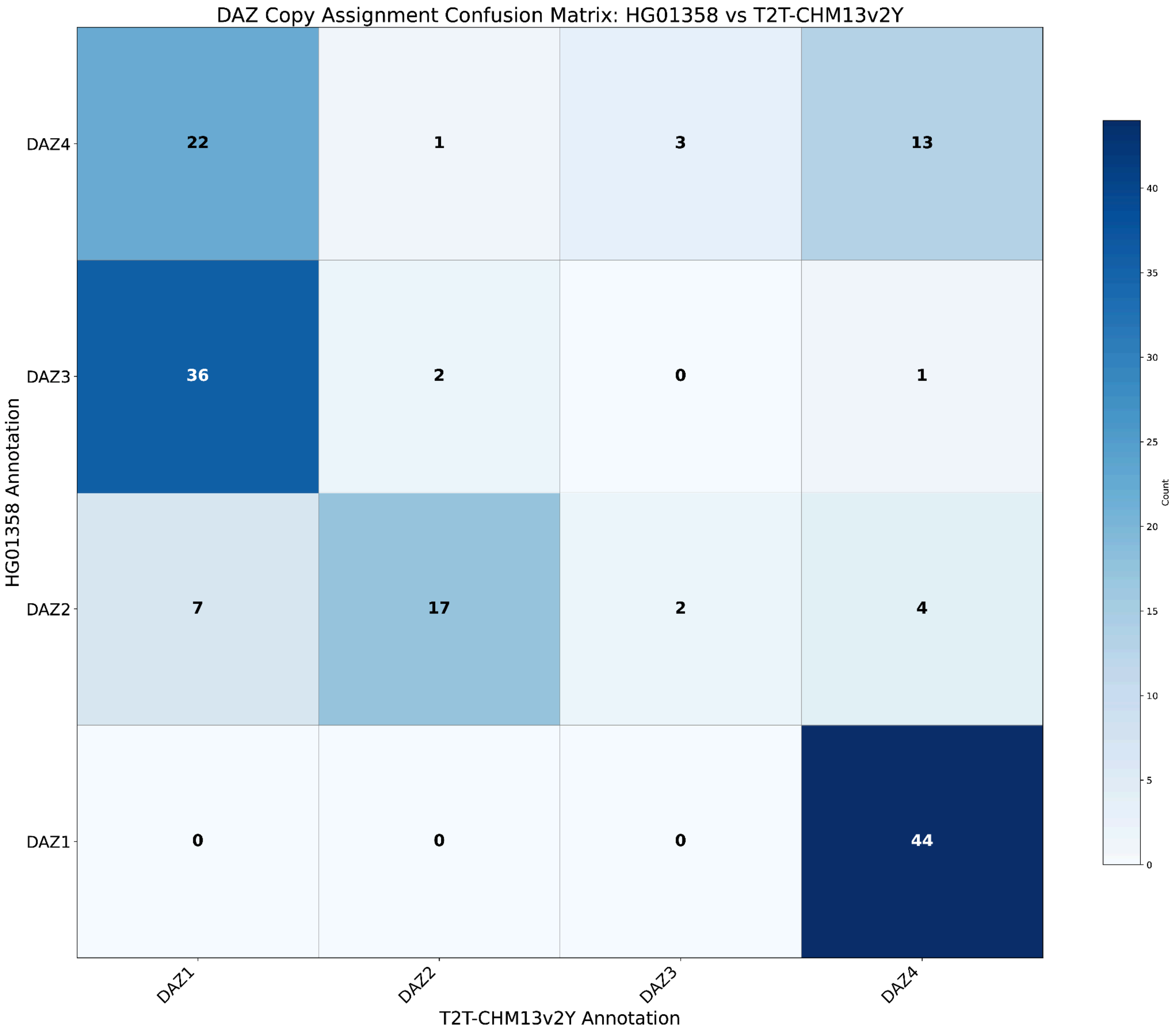


**Supplementary Figure 90. T2T-CHM13v2Y vs best assembly *DAZ* copy annotation.** A confusion matrix portraying the difference in reads aligned to ampliconic gene copies when mapping to T2T-CHM13v2Y vs the assembly producing the best alignment quality out of 140 individuals.

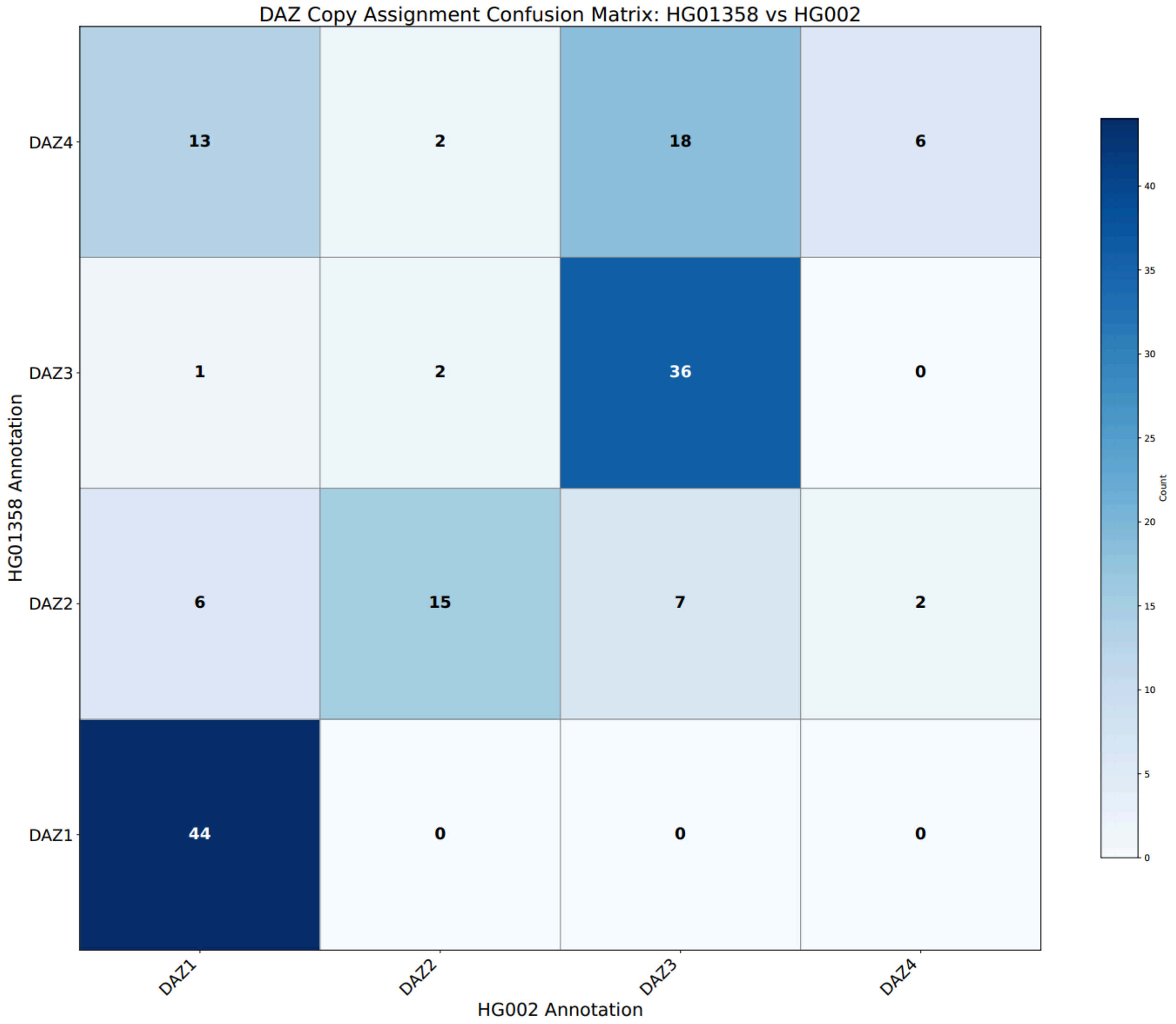


**Supplementary Figure 91. HG002 vs best assembly *DAZ* copy annotation.** A confusion matrix portraying the difference in reads aligned to ampliconic gene copies when mapping to the HG002 assembly vs HG01358.

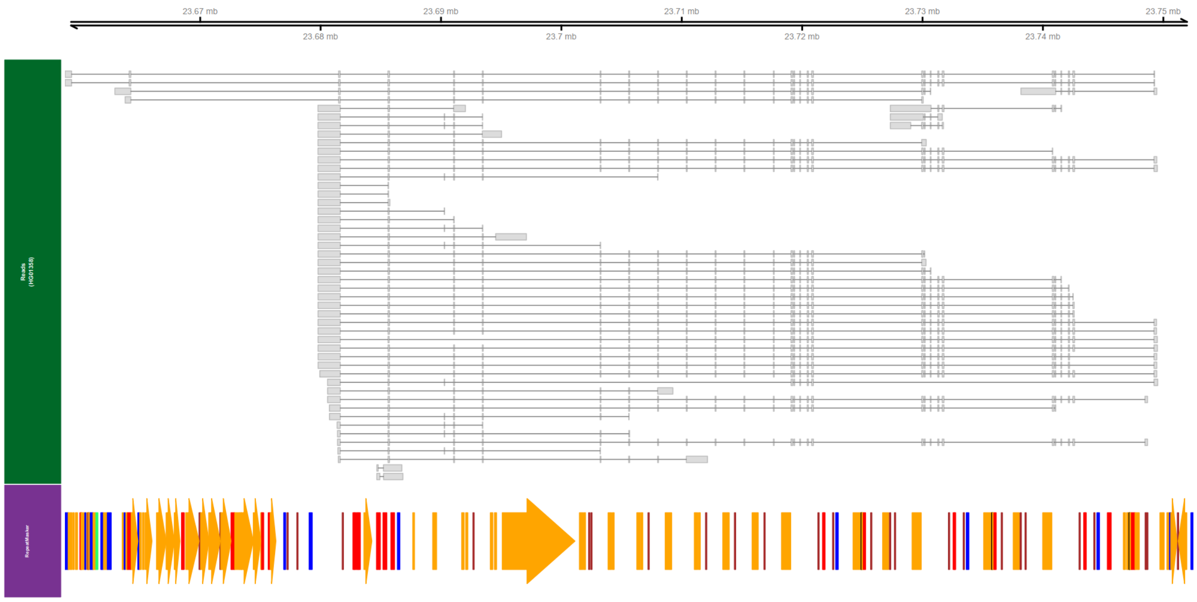


**Supplementary Figure 92. ENCODE expression snapshot - *DAZ1*.** An overview of all unique splicing *DAZ1* transcripts expressed in testis from a single individual (SRR31360662) visualized in the coordinate space of the best mapping assembly (HG01358) of the 140 individuals. Repeats are represented with directionality as arrows: orange (LINE), red (SINE), blue (DNA transposon), brown (simple repeat), and green (LTR retrotransposon). Alternative transcription start sites can be observed within LINE elements as well as the presence of unreported exons.

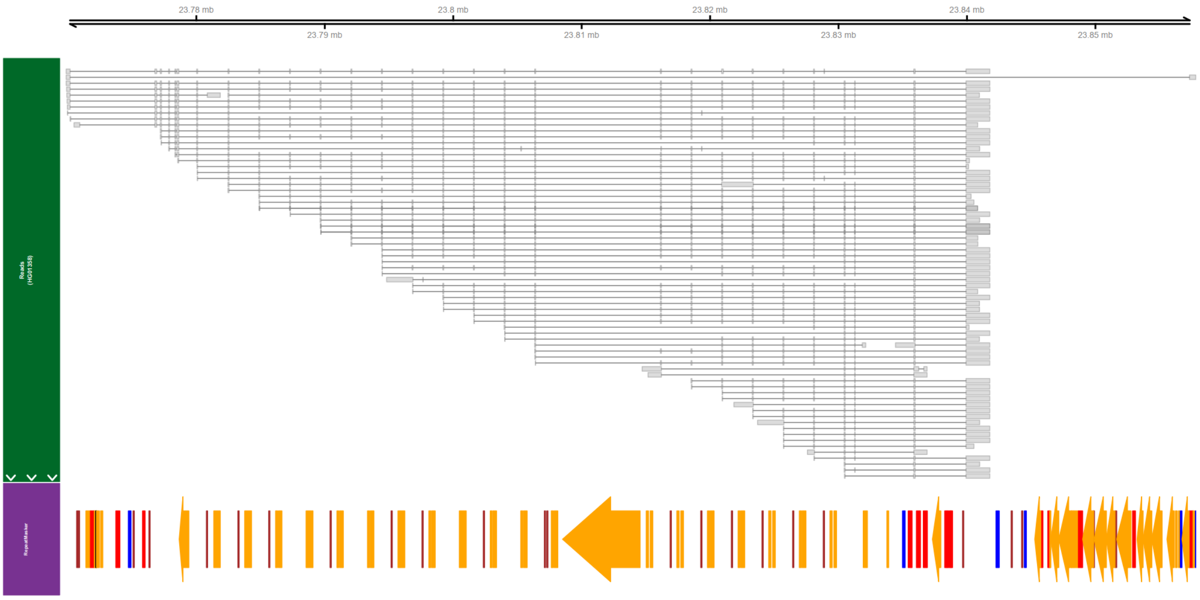


**Supplementary Figure 93. ENCODE expression snapshot - *DAZ2*.** An overview of all unique splicing *DAZ2* transcripts expressed in testis from a single individual (SRR31360662) visualized in the coordinate space of the best mapping assembly (HG01358) of the 140 individuals. Repeats are represented with directionality as arrows: orange (LINE), red (SINE), blue (DNA transposon), brown (simple repeat), and green (LTR retrotransposon). Alternative transcription start sites can be observed within LINE elements as well as the presence of unreported exons.

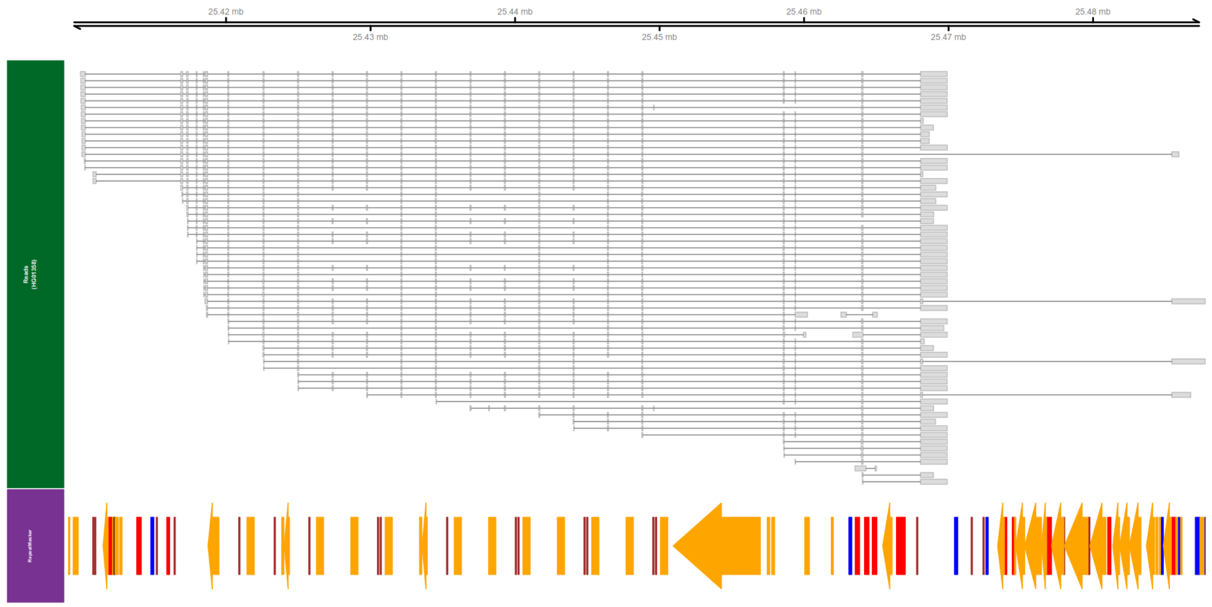


**Supplementary Figure 94. ENCODE expression snapshot - *DAZ3*.** An overview of all unique splicing *DAZ3* transcripts expressed in testis from a single individual (SRR31360662) visualized in the coordinate space of the best mapping assembly (HG01358) of the 140 individuals. Repeats are represented with directionality as arrows: orange (LINE), red (SINE), blue (DNA transposon), brown (simple repeat), and green (LTR retrotransposon). Inclusion of an unreported exon 29 can be observed.

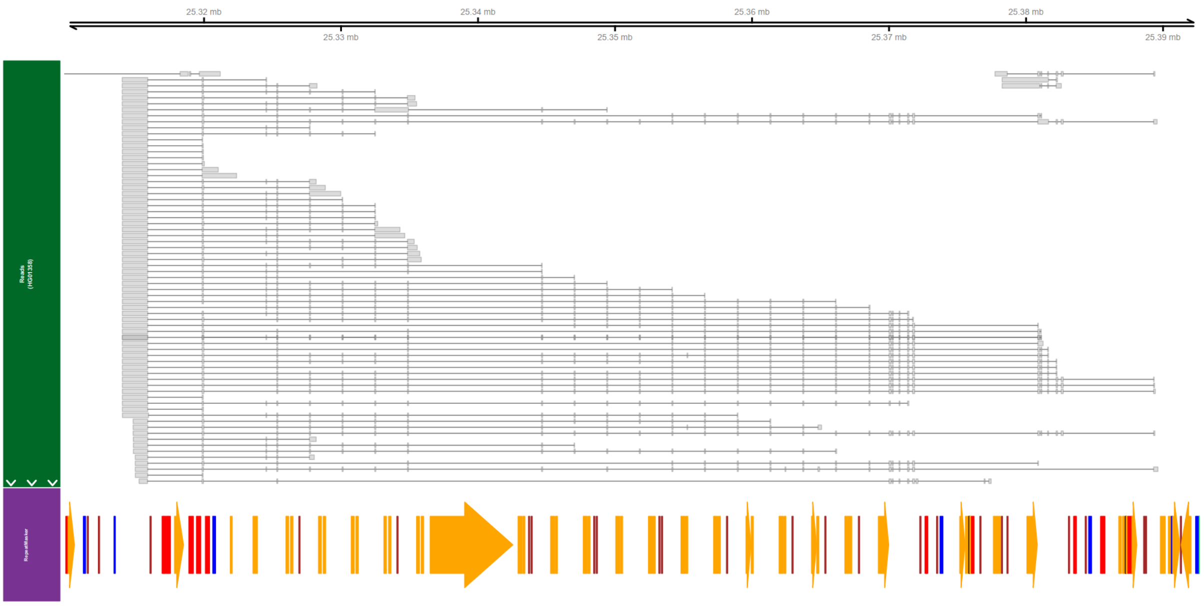


**Supplementary Figure 95. ENCODE expression snapshot - *DAZ4*.** An overview of all unique splicing *DAZ4* transcripts expressed in testis from a single individual (SRR31360662) visualized in the coordinate space of the best mapping assembly (HG01358) of the 140 individuals. Repeats are represented with directionality as arrows: orange (LINE), red (SINE), blue (DNA transposon), brown (simple repeat), and green (LTR retrotransposon). Alternative transcription start sites can be observed within LINE elements as well as the presence of unreported exons.

### *TSPY* gene expression and copy number variation

Contributors: Rebecca Siford, Brendan J. Pinto, Melissa A. Wilson

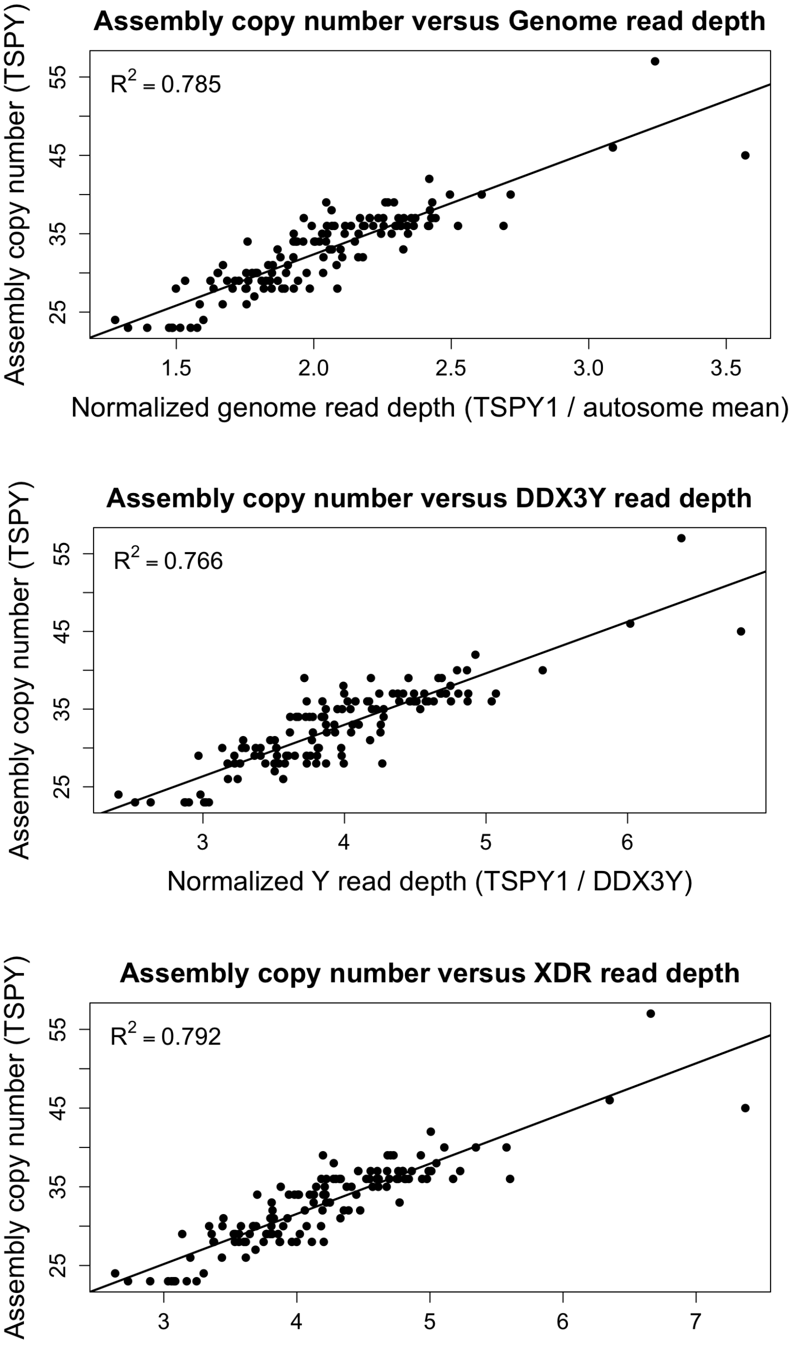


**Supplementary Figure 96. Assembly copy number versus read depth normalization approaches for *TSPY*.** Scatterplots show the HPRC/assembly derived *TSPY* copy number on the y axis versus short read depth based estimates from 30x whole genome alignments on the x axis.

Read depth was computed as the mean depth across the annotated *TSPY1* region and normalized by 3 methods: autosomal mean depth (chr1-22), a single copy Y gene (*DDX3Y*), and mean depth across the X-degenerate (XDR) regions on the Y chromosome. Each point is one individual. The fitted lines show a linear regression used for calibration, and the reported R^2 indicates model fit for each normalization method; Genome read depth (R^2 = 0.785), *DDX3Y* (R^2 = 0.766), and XDR (R^2 = 0.792). Mean depth across the autosomes is used as the primary regression model for later analyses due to similar fit between models.

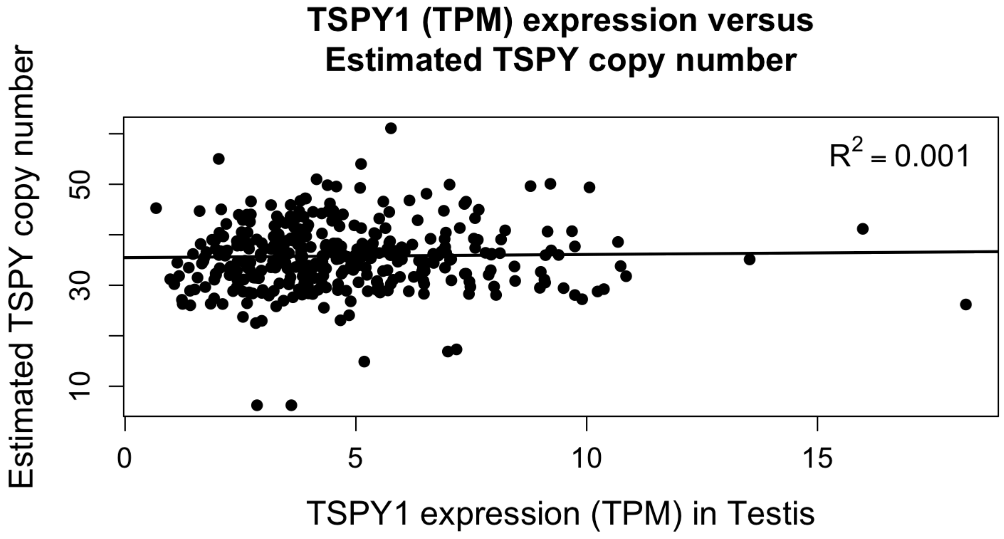


**Supplementary Figure 97. *TSPY1* expression versus estimated copy number in GTEx testis.** Each point represents a GTEx testis RNA seq sample (n=367). *TSPY1* expression (TPM) is plotted on the x axis against estimated *TSPY* copy number on the y axis. Estimated *TSPY1* copy number was computed by converting short read depth normalized across the autosomes to copy number using a linear fit to the HPRC assembly. The R^2 summarizes the association between expression and estimated copy number in testis (R^2 = 0.001).

### Y callable regions

Contributors: Rebecca Siford, Nancy F. Hansen, Melissa A. Wilson

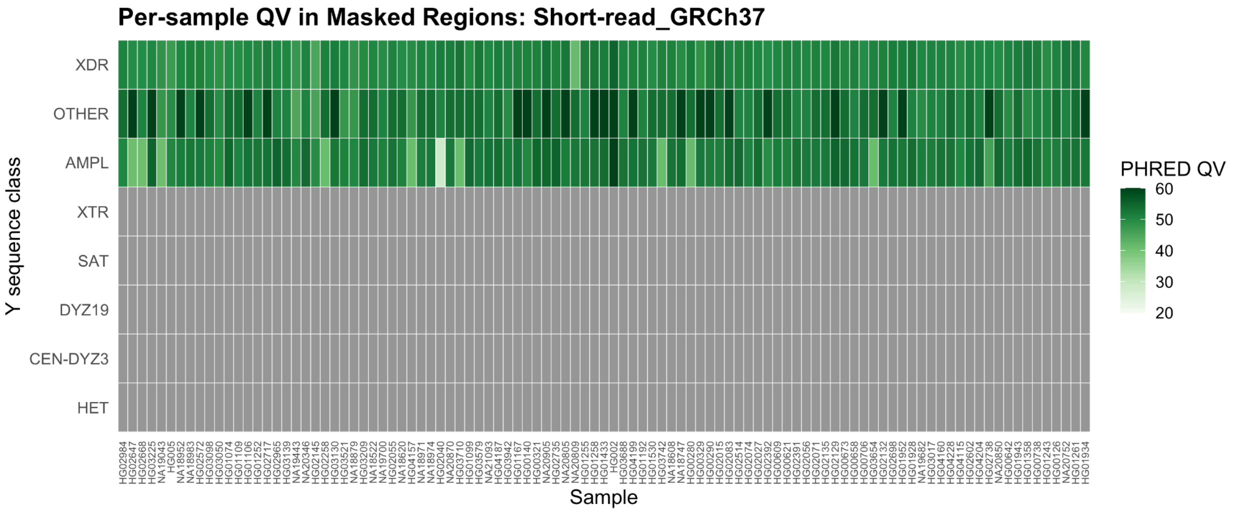


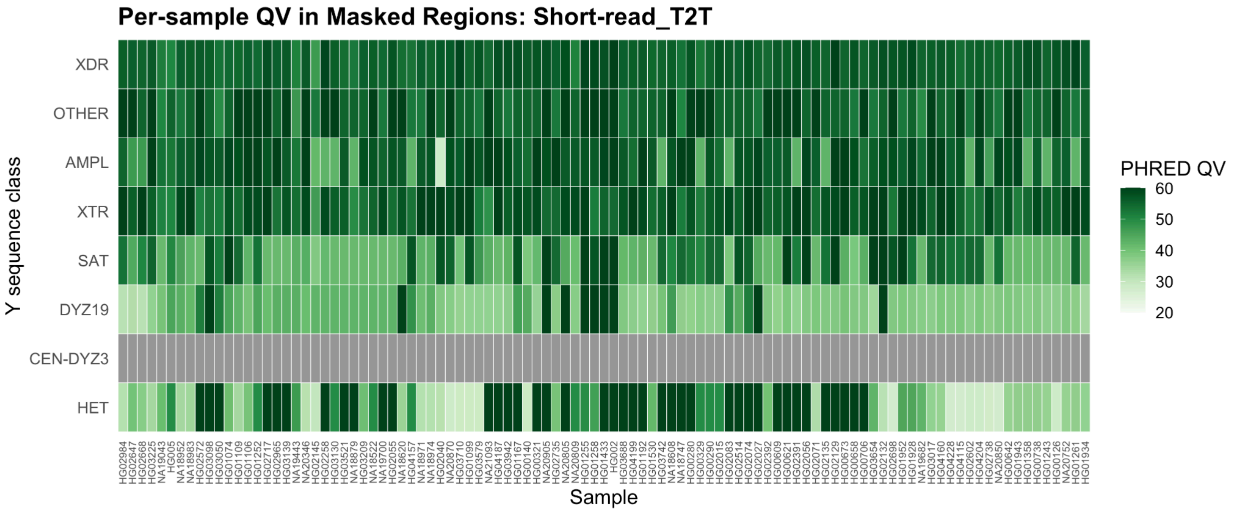


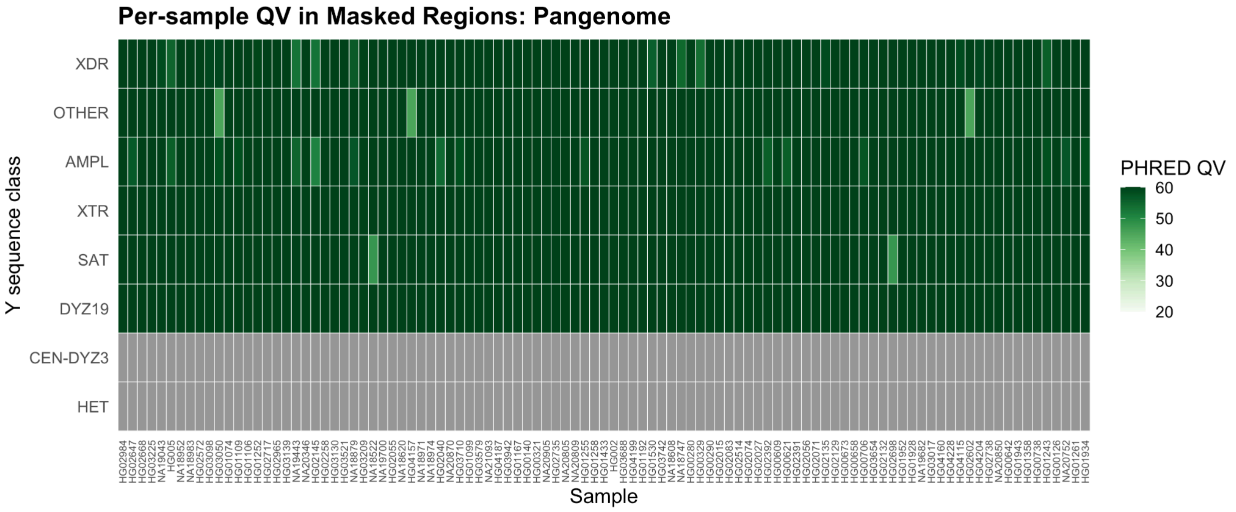

**Supplementary Figure 98. Per sample QV within masked Y chromosome sequence classes.** Heatmaps show phred scaled quality values (QV) for each sample across Y chromosome sequence classes after intersecting with bases within each global callable region mask. Panels correspond to the Short read GRCh37, short read T2T (T2T-CHM13v2Y), and Pangenome (**Methods**) masks. QV was calculated from the number of discrepancies between the variant constructed genome and the corresponding Verkko Y assembly within each masked sequence class. Sequence classes were defined using T2T-CHM13v2Y based annotations. Samples are ordered by Y chromosome phylogeny. Grey tiles indicate where no masked QV could be calculated because no bases were called.
